# Exploring intra- and intergenomic variation in haplotype-resolved pangenomes

**DOI:** 10.1101/2024.06.05.597558

**Authors:** Eef M. Jonkheer, Dick de Ridder, Theo A. J. van der Lee, Jorn R. de Haan, Lidija Berke, Sandra Smit

**Affiliations:** Bioinformatics Group, Wageningen University & Research, Droevendaalsesteeg 1, 6708PB, Wageningen, The Netherlands; Biointeractions and Plant Health, Wageningen Plant Research, Wageningen, 6708PB, The Netherlands; Genetwister Technologies B.V., Nieuwe Kanaal 7b, 6709 PA, Wageningen, The Netherlands

## Abstract

With advances in long-read sequencing and assembly techniques, haplotype-resolved (phased) genome assemblies are becoming more common, also in the field of plant genomics. Computational tools to effectively explore these phased genomes, particularly for polyploid genomes are currently limited. Here we describe a new strategy adopting a pangenome approach. To analyze both intra- and intergenomic variation in phased genome assemblies, we have made the software package PanTools ploidy-aware by updating the pangenome graph representation and adding several novel functionalities to assess synteny and gene retention, profile repeats and calculate synonymous and nonsynyonymous mutation rates. Using PanTools, we constructed and analyzed a pangenome comprising of one diploid and four tetraploid potato cultivars, and a pangenome of five diploid apple species. Both pangenomes show high intra- and intergenomic allelic diversity in terms of gene absence/presence, SNPs, indels and larger structural variants. Our findings show that the new functionalities and visualizations are useful to discover introgressions and detect likely misassemblies in phased genomes. PanTools is available at https://git.wur.nl/bioinformatics/pantools.

## I. Introduction

Until recently, obtaining chromosome-scale genome assemblies, let alone haplotype-phased genomes, required tremendous effort. As a result, genome assemblies of diploid or polyploid organisms used in genomic analyses are typically haploid representations, where the multiple copies of a chromosome are collapsed into a single sequence with a mosaic of alleles. Such haploid representations ignore the intragenomic variation in gene content and organization. However, differences between haplotypes may provide novel insights, not only into the evolutionary history but also in explaining certain phenotypes [1]. Including intragenomic variation in reference genomes would greatly facilitate their use for genetics and breeding but requires comprehensive methodology to define haplotypes [2], [3]. Due to various advances in sequencing technology [4], [5] and assembly algorithms [6], [7], the different haplotypes of genomes with two or more chromosome sets can now be accurately resolved. Exploring such fully phased genomes allows for more comprehensive assessment of the genetic variation, and enables new types of analyses such as gene-dosage analysis or detection of allele-specific expression (when combined with other -omics data).

In recent years, there has been an increasing number of available (partially) phased genome assemblies for fungi [8], [9], plants [10]–[13], and animals [14]–[16]. In these recent publications, the haplotype-resolved genomes are generally analyzed through custom pipelines that consist of various tools not specifically designed to handle (partially) phased genome assemblies. To enable efficient comparison of multiple phased genomes, we propose pangenome representations and related tools as a possible solution. These approaches already serve to identify small and large-scale variation in large genome collections.

We have developed an extensive toolkit for pangenome analysis, called PanTools. It stores a distinctive hierarchical graph structure in a Neo4j database, including a compacted De Bruijn graph (DBG) to represent sequences. Structural annotation nodes are linked to their respective start and stop positions in the DBG. Homology relationships function as an additional layer to connect annotation nodes. The heterogeneous graph can be queried through Neo4j’s Cypher query language or by various PanTools methods for gene-level and phylogenetic analyses [17]. To enable the analyses of haplotype-resolved genomes, we now made PanTools ploidy aware by including new functionality to perform comparative analyses within and between genomes.

In this study, we demonstrate the new PanTools functionality on two datasets: tetraploid potatoes (*Solanum tuberosum*) and diploid apples (*Malus* spp.). These two crops were selected due to the availability of multiple high-quality haplotype-resolved assemblies [11], [12], [18]–[22], along with a hypothesised high intragenomic variation among haplotypes. This variation is thought to arise from the occurrence of multiple ancient whole-genome duplication (WGD) events within these lineages. Potato and apple share three WGD events and each has its own lineage-specific event [23]–[25]. Moreover, the more recent domestication of these two crops involved extensive selective breeding and hybridization [26], [27].

We show the usefulness of our pangenome representation and functionalities, emphasizing the previously hidden intragenomic variation within haploid reference genomes. We describe a universal approach for constructing both pangenome graphs and characterizing gene content both inter- and intragenomically, and explore different approaches to establish phylogenetic relationships between sequences. With our newly developed visualization methodology, we create visualizations that provide insights into the variation in genomic organization. Finally, we utilized our pangenome approach to identify new allelic variation of *StCDF1*, a key regulator of maturation and tuberisation in potato.

## II. Results AND DISCUSSION

### A. Ploidy-aware pangenome analyses

PanTools has a hierarchical pangenome representation, linking divergent genomes not only through a sequence variation graph but also through structural and functional annotations and homology. To enable the analysis of haplotype-resolved assemblies, we introduced new annotations to label haplotypes, updated existing functionalities, and introduced a new set of command-line tools.

PanTools allows for the incorporation of the haplotype information to control which sequences or features are compared. Pangenomes are constructed from collections of genome FASTA files; a genome layer is formed with *genome* nodes that connect to *sequence* nodes (representing contigs/scaffolds/pseudomolecules), which in turn link to the start and stop *nucleotide* nodes in a compacted De Bruijn graph (DBG) representation of the DNA sequences. The database scheme underlying the graph is shown in Supplementary Fig. S3. A *sequence* node can be annotated with a chromosome number (1, 2, .. .) and haplotype phase (A, B, .. .). By combining the two, each sequence node has a unique haplotype identifier within a genome (e.g. 1A, 1B, 2A, etc.). Sequences in a genome with the same haplotype (letter) are considered to be a subgenome. From a biological standpoint this concept of a subgenome does not exist, as there is no genetic or physical linkage between assembled chromosomes. However, defining the subgenomes in this way enables the assessment of gene presence among multiple haplotypes within a specific chromosome (e.g. 1A 1B 1C 1D). Nevertheless, it is worth to note that it is not meaningful to compare for instance, the “C” subgenome between two genomes, given the random composition of chromosomes within the subgenome. Finally, sequences that lack phasing information are all labeled as chromosome 0 (and do not receive a unique haplotype identifier). A schematic overview of the terminology is included as Supplementary Fig. S1.

Every pangenome analysis starts with collecting genome and sequence nodes, to determine which sequences and features will be compared. Phasing information enables the collection of specific sequences to perform more targeted analyses - for example, comparison among genomes, specific subgenomes or homeologous chromosomes. In genome assemblies where the chromosomal organization is still unknown, phylogenetic relatedness is determined from the number of shared *k*-mers in the DBG. This distance method was updated to count *k*-mers per sequence instead of per genome, allowing the identification of homoeologous chromosomes.

We updated several PanTools methods to work at both sequence level and genome level. Single-copy genes function as ideal markers for phylogeny inference [28]. In phased genomes, single-copy genes may have a copy per subgenome, drastically reducing the number of genes that can be detected when simply looking for a single copy using the current methods at genome level. To address this issue, we allow for one gene copy per subgenome. In this way applications that rely on single-copy genes, such as BUSCO and the core phylogeny, can work at a subgenome level rather than the bulked genome.

PanTools’ gene classification method identifies shared genes between genomes and can describe a pangenome’s gene content as core (present in all genomes), accessory/dispensable (present in some but not all), and cloud (present in one genome) (Supplementary Fig. S2). We updated the term ‘unique’ to ‘cloud’ because the former suggests a gene is found only once, while it may actually have multiple (allelic) copies. With haplotype information incorporated in the graph, gene presence/absence can now be established for every subgenome of a genome. Accordingly, we further characterize gene content by presence in number of subgenomes (1, 2, .. .).

To further explore the information in fully phased genomes, new functionalities have been developed in PanTools, integrating bioinformatics methods into the pangenome representation. First, synteny (collinear gene blocks) between genomes, as detected by MCScanX [29], can be added to the pangenome (Supplementary Fig. S3). From the synteny block information we calculate gene retention and visualize fractionation patterns across chromosomes. We speed up the minimap2 [30] whole-genome alignments analysis, by making use of haplotype and chromosome information to compare only homeologous chromosomes, thereby avoiding computationally intensive all-vs-all comparisons. With PAL2NAL [31] we calculate synonymous and nonsynyonymous mutation rates on aligned sequences in homology groups or syntenic gene pairs, allowing to study evolutionary rates in a species or population. Finally, we developed a novel graphical representation of genomic structure in which we combine the newly integrated genomic features such as synteny, repeat density and subgenome presence of genes.

### B. Pangenome design choices and construction

To showcase PanTools’ updated and novel functionalities, we built two pangenomes exhibiting different ploidy levels. A species-level pangenome was constructed of five *S. tuberosum* (potato) cultivars: DM1-3 516 R44 (DM) [20], Atlantic [12], Castle Russet (CR) [12], Otava [18] and Cooperation-88 (C88) [19]. DM is a doubled haploid and therefore represented as a haploid assembly, whereas the other accessions were tetraploid with fully resolved haplotypes, leading to 17 (1 + 4 × 4) subgenomes in total. All assemblies were chromosome-scale and had 12 or 48 pseudomolecules, matching the base chromosome number of 12 in *S. tuberosum* [20]. We found a clear dichotomy in assembly statistics as a result of different sequencing and assembly approaches. The Otava and C88 assemblies, of size 3.1-3.2 Gb, were considerably larger than CR and Atlantic at 2.5-2.7 Gb. This large difference was further reflected in the total numbers of genes annotated: 150,853-152,835 in Otava and C88 and 105,449-114,021 in Atlantic and CR. These notable discrepancies suggests the two latter assemblies have a greater number of collapsed regions or suffer from other assembly/annotation challenges.

Benchmarking universal single-copy orthologs (BUSCO) [32] has become the standard for assessing genome assembly quality. Unlike technical metrics, such as the number of reads mapping back to the genome or the N50 value, BUSCO is biologically meaningful and based on informed expectations of the single-copy gene content. Completeness of the *S. tuberosum* genomes according to BUSCO ranged from 93.6-99.5% (Supplementary Fig. S4). All four haplotype-resolved genomes showed high levels of duplication, especially C88 in which nearly all (98.2%) BUSCO genes were duplicated. Because we suspected that most duplicates were a result of haplotype phasing, BUSCO was run separately on each of the subgenomes (Supplementary Fig. S5).

This approach successfully removed gene duplicates, but also revealed large differences in subgenome completeness: 59.2-62.9% in CR, 70.2-77.3% in Atlantic, 88.9-90.9% in Otava and 95.6-96.6% in C88. BUSCO applied on only unsorted/unphased sequences showed minimal completeness (*<* 1%) in Otava and C88, but substantial completeness in Atlantic and CR (*>* 31.0%). This again suggests haplotypes of the latter two genomes were not fully phased. The C88 subgenomes had the highest completeness, resulting in 4,870 BUSCO genes detected as single-copy on all four subgenomes (Supplementary Fig. S6-S9). In comparison, Atlantic had only 1,307 single-copy genes found in all four subgenomes. These contrasts in statistics reflect clear differences in phasing quality between genomes.

Inferring homology is fundamental to gene-based pangenome analyses. *S. tuberosum* proteomes were clustered with different settings (so-called ‘relaxation modes’), where increasing relaxation modes indicate lower clustering stringency. The most critical parameter is the minimum required sequence similarity of pairwise alignments, starting at 95% and lowered by 10% in each subsequent mode. Between relaxation mode 2 and 8, a nearly 7-fold decrease in the number of homology groups was observed (Fig. 1A). The availability of different relaxation modes allows calibration to different data sets, but obviously raises the question of what the optimal setting is.

**Fig. 1.**
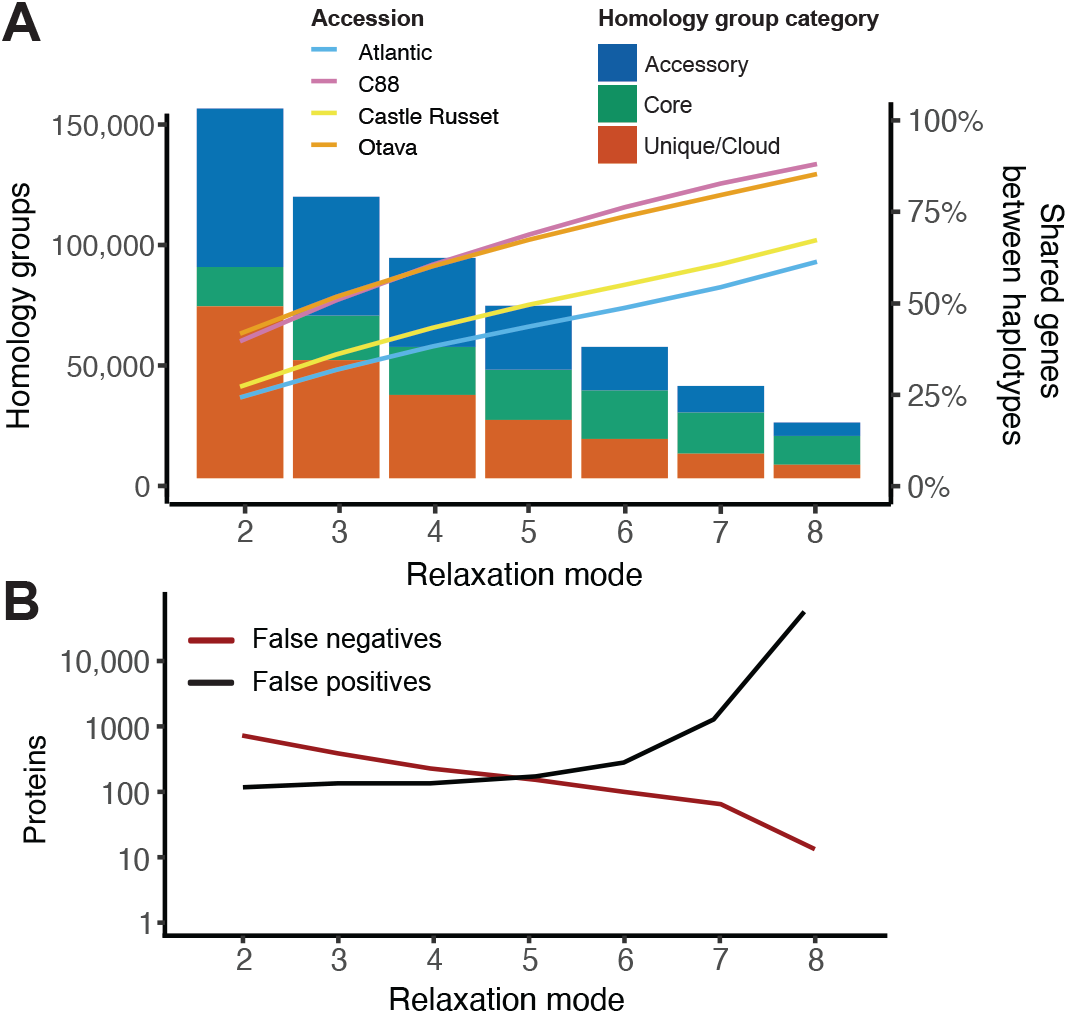
(A) The effect of increasing relaxation modes (lowering clustering stringency) on the *S. tuberosum* pangenome composition, in terms of total number of homology groups (bar charts) and average percentage of genes shared between subgenomes (line graphs). (B) PanTools’ BUSCO benchmark results of the seven homology grouping settings (relaxation modes).

We used BUSCO genes to assess each homology grouping and found a clear trade-off between recall and precision (Fig. 1B). The highest *F*_1_-scores (not shown), combining recall and precision, were obtained with mode 5 and 6, corresponding to ≥ 55% or ≥ 45% sequence similarity thresholds. After mode 6 the number of false positives rapidly increases. In addition to this benchmark, we calculated the average percentage of genes shared between subgenomes as a metric for selecting a suitable grouping (line graphs in Fig. 1A). The clear division between the two pairs of genomes is in line with the earlier BUSCO completeness assessment: CR and Atlantic subgenomes share less because of lower phasing quality. As the difference in *F*_1_-scores between mode 5 and 6 was negligible, mode 6 was selected based on the higher overlap in gene content between subgenomes.

Besides potato, a genus-level pangenome was constructed from five diploid *Malus* (apple) accessions: *M. domestica* cv. Gala (Gala), *M. domestica* ‘Golden Delicious’ GDDH13, *M. sieversii, M. sylvestris* and *M. baccata*. Haplotypes were resolved for the Gala, *M. sieversii* and *M. sylvestris* genomes. All assemblies except *M. baccata* were arranged into 17 or 34 whole-chromosome pseudomolecules, correctly representing the 17 chromosomes of the *Malus* genus [21].

For constructing the *Malus* pangenome, we followed the same BUSCO clustering approach. BUSCO indicated very high completeness for all genomes, as well as for the separate subgenomes. The analysis supports the existence of a recent Maleae-specific whole-genome duplication (WGD), as nearly one third of the gene content in all *Malus* (sub)genomes was marked as duplicated. A more detailed description of the Malus pangenome construction is provided in Supplementary Analysis 1. Following a universal strategy, we built the *S. tuberosum* and *Malus* pangenomes of datasets that were distinct in terms of genome size, ploidy level, phasing quality, and evolutionary divergence between genomes (Table I).

**Table I.** Genome assembly and pangenome statistics.

|  | *S. tuberosum* | *Malus* |
| --- | --- | --- |
| Genomes | 5 | 5 |
| Subgenomes | 17 | 8 |
| Sequences | 34,050 | 158,662 |
| Total input size | 12.2 Gb | 5.3 Gb |
| Protein-coding genes | 723,497 | 361,864 |
| *K*-mers | 1,117.7M | 690.0M |
| Compressed *k*-mers | 221.6M | 103.2M |
| Homology groups | 52,240 | 68,751 |
| Singleton groups | 6,638 | 15,635 |

### C. Gene content and relatedness of (sub)genomes

Pangenomes are studied by classifying genes as shared between (subsets) of genomes, i.e. which genes are core, which are accessory and which are cloud. Given phased assemblies, gene content can also be assessed within and between subgenomes: where are these core/accessory/cloud genes located, and to what extent are they found in different subgenomes? Such analyses shed light on the organization and evolution of subgenomes, albeit subject to assembly and annotation quality. With homology relationships established and integrated into the graph, we next characterized the genetic composition of the *S. tuberosum* pangenome. The *Malus* analysis is included as Supplementary Analysis 2.

The protein-coding genes of the *S. tuberosum* genomes clustered in 52,240 homology groups, 37.1% of which were core, 33.0% accessory and 29.9% cloud (Fig. 2A). Nearly half of the cloud groups are present in only single subgenome. Notably, 80% of these subgenome exclusive groups are singleton groups, and do not show any homology to another sequence, causing suspicion about their realness. Interestingly, cloud genes were more abundant in the higher quality Otava and C88 genomes. On the other end of the spectrum, core genes require occurrence in a minimum of 5 subgenomes (from different accessions), but are generally found in 11-17 subgenomes. When we analyse gene content of individual genomes, we see the majority of genes was characterized as core (Fig. 2B).

**Fig. 2.**
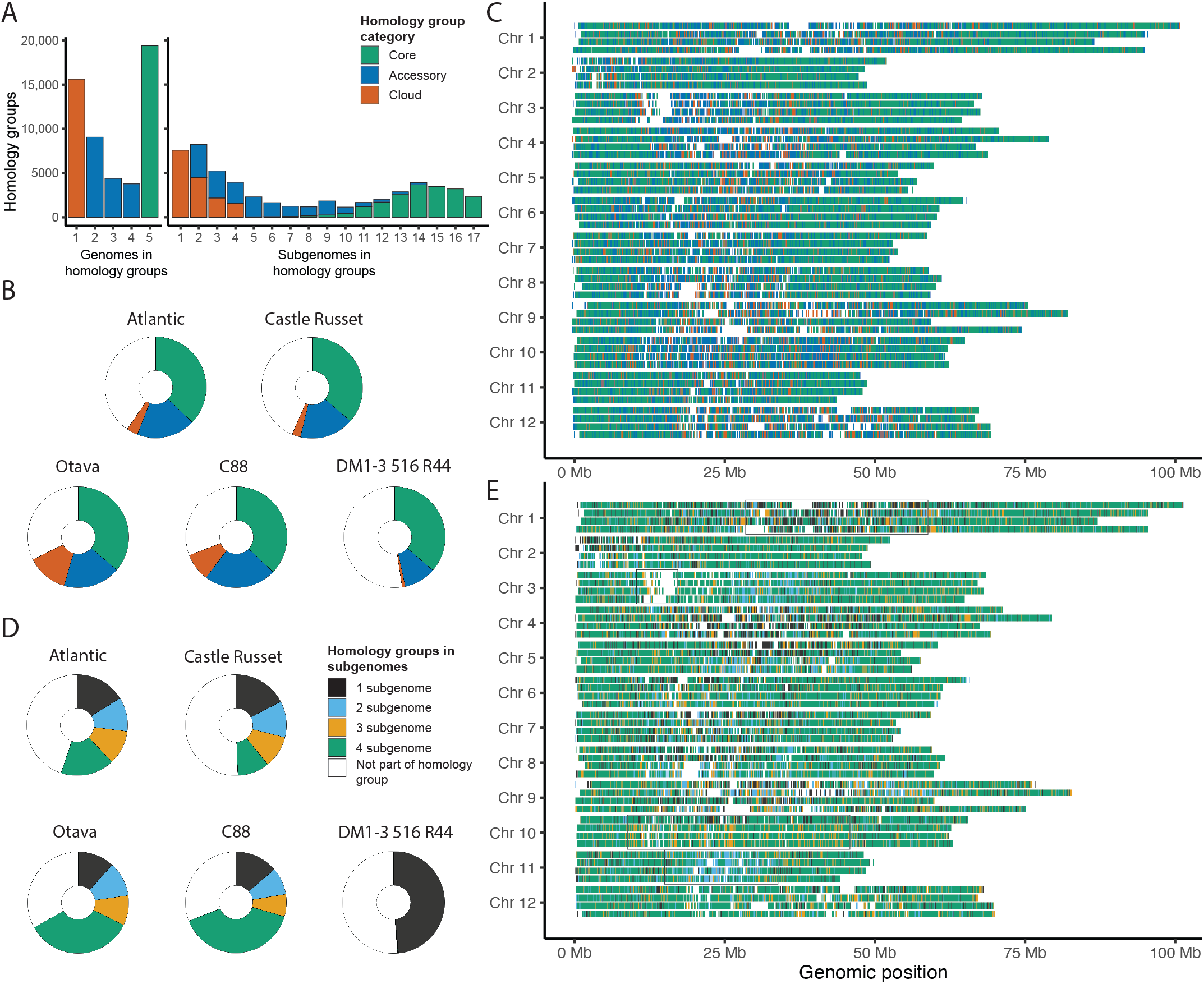
Characterization of the *S. tuberosum* pangenome gene content. (A) Number of genomes (left) and subgenomes (right) in the 52,240 homology groups. (B) Pie chart slices representing the proportion of groups being classified as core, accessory, or cloud. Each circle represents the pangenome’s 52,240 homology groups. (C) Gene regions of C88 colored based on intergenomic variation, the pangenomic gene classification of five genomes. (D) Pie charts where the slices show number of homology groups with genes in total number of subgenomes. The proportion of white in these circle is slightly larger compared to plot B, because certain genes are not part of a subgenome; they are located on unphased sequences. (E) C88 gene regions colored by intragenomic variation, their presence in 1 to 4 subgenomes.

The genomic distribution of C88’s genes revealed an interesting pattern (Fig. 2C). Most accessory and cloud genes are positioned in the pericentromeric region of the chromosome, whereas core genes generally lie in chromosome arms. The observed localization of cloud genes was most prominent in the Otava and C88 genomes but was also seen in the other genomes (Supplementary Fig. S10-S13). Clear differences in patterns between these visualizations suggest that due to the higher quality, a larger number of genes were likely annotated in high repetitive regions. The high frequency of cloud genes in pericentromeric regions may be partly attributed due to the lack of recombination in the heterochromatic centromere [33], [34]. Another contributing factor might be the accumulation of deleterious mutations in potato genomes, disrupting open reading frames and altering the protein sequences [35].

As an alternative to grouping *S. tuberosum* genes by their intergenomic presence in the pangenome, we can now also characterize genes by intragenomic presence in (1 to 4) subgenomes. In Otava and C88 nearly half of the groups have genes present on all four subgenomes, whereas in Atlantic and CR this was around a fifth of the groups (Fig. 2D). We again visualized C88’s gene regions but now colored them according to the subgenome characterization (Fig. 2E). Core genes in the chromosome arms are strongly correlated to presence in all four haplotypes. We briefly discuss four chromosomal sections with distinctive patterns:

- All four haplotypes in the middle of Chr 1 show a stretch of gene regions occurring in a single haplotype, indicating a distinct gene set on each haplotype.
- The absence of genes within the same region of four Chr 3 haplotypes is indicative of the centromere, characterized by its high repeat content.
- Chr 10 shows one disparate haplotype, indicated in orange (3 occurrences) in the remaining haplotypes.
- The pericentromeric regions of Chr 11 predominantly show two divergent alleles present on two haplotypes, suggesting two distinct sets of alleles.

This visualization demonstrates the haplotype diversity in the C88 genome. Additionally, it shows clear mosaic patterns that indicate large genomic regions can exist in one to four copies. The uneven distribution is clearly exemplified in the last two patterns: Chr 10 and 11 have differently observed relationships (2:2 versus 3:1). As the C88 potato variety derived from two distinct cultivars, backcrossing is the likely cause of this observable genomic architecture [19].

We generated genome visualization of the other four *S. tuberosum* genomes as well (Supplementary Fig. S10-S13). Otava was comparable to C88, having multiple chromosomes with a single disparate haplotype and also a chromosome with bi-allelic regions. In contrast, patterns in both CR and Atlantic mostly indicate limited phasing. Overall, these results demonstrate how subgenome-level pangenomics can help explore differences in terms of gene content, provided genome assemblies are of sufficient quality.

### D. Establishing the evolutionary history in the pangenome

Inferring an accurate phylogeny is crucial for understanding the evolutionary history. The heterogenous pangenome graph allows us to infer phylogeny from different types of genetic variation, such as *k*-mers, SNPs, or genes. The choice of input data and algorithm depends on the research objective. In this study, we were interested in comparing different methods to identify a reliable method for haplotype-resolved assemblies.

Here, we describe phylogenetic relationships in the *S. tuberosum* pangenome; the phylogenetic analysis of *Malus* is described in Supplementary Analysis 3. Establishing a genome-level SNP tree on single-copy genes (occurring only once in every genome) is the most commonly used method for resolving plant phylogeny. It was not possible to directly obtain these high-resolution SNP trees for either of the haplotype-resolved pangenomes, as *S. tuberosum* had only 7 identified single-copy homology groups, whereas *Malus* had 17 groups.

Complete subgenomes cannot be directly compared, as the haplotype assignments per chromosome are ambiguous when genomes are assembled without parental data. Therefore, rather than making a phylogeny of complete subgenomes we inferred separate trees per chromosome. PanTools includes a range of phylogenomic methods that were adjusted to this end. Two methods use a distance based on either the number of shared *k*-mers or genes. Two more sophisticated methods, a core genome SNP tree and a consensus tree of core gene trees, were strongly hampered by the phasing quality as they require core gene content.

Here, we explore the tree topologies of *S. tuberosum* Chr 1 as a representative example for the other chromosomes. Tree topologies of the 11 remaining potato chromosomes show similar trends as Chr 1. As input for the Chr1 core SNP tree, we collected homology groups with one gene copy per haplotype. Genes of the individual Chr1 haplotypes cluster into 2,155-4,010 homology groups, but only 296 were shared among all haplotypes, with 52 groups identified as single-copy. From these single-copy groups, a low-resolution SNP tree with high ambiguity was inferred. A Splitstree representation [36] of this phylogeny in Fig. 3A reveals clear conflicting signals. The consensus tree (Supplementary Fig. S14A) was inferred on the 296 groups present in every (Chr 1) haplotype. Equivalent to the SNP tree, the consensus tree shows minimal branch support.

**Fig. 3.**
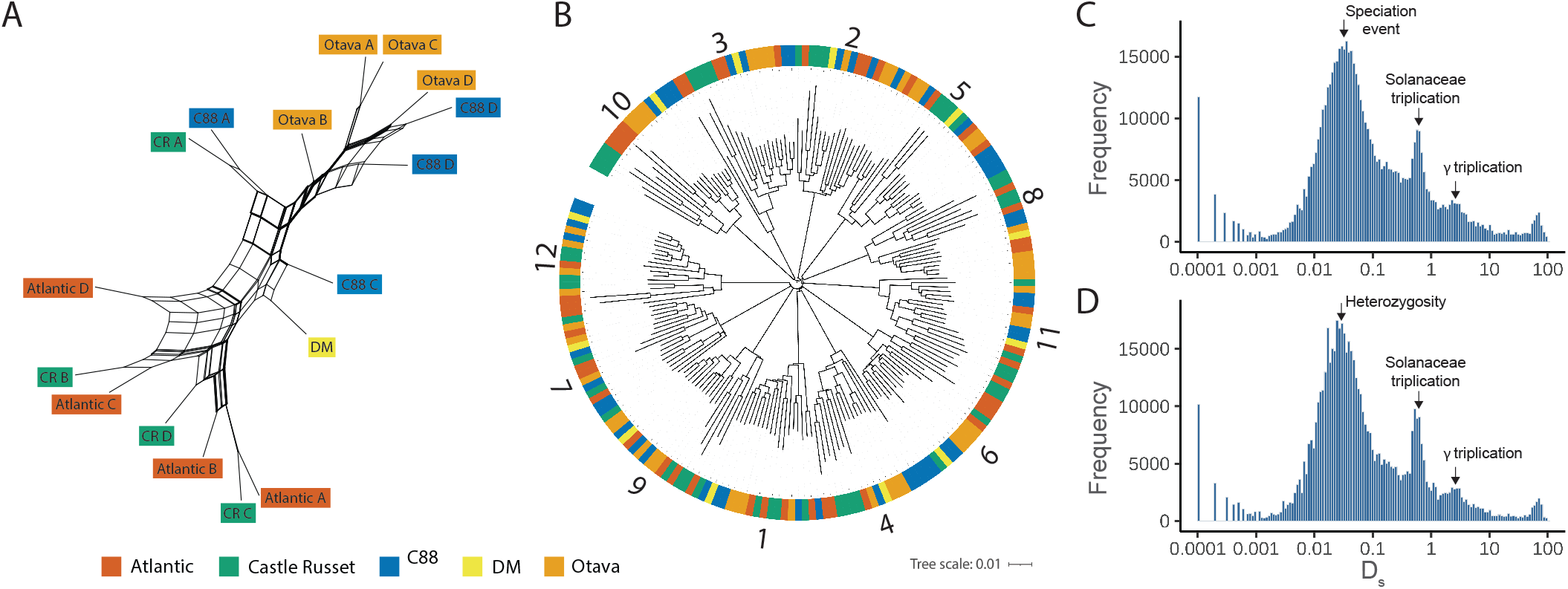
Evolutionary history of *S. tuberosum*. (A) Splits graph of *S. tuberosum* Chr 1 core SNP phylogeny. Tree labels are colored by accession name. (B) *K*-mer distance phylogenetic tree of *S. tuberosum* sequences with at least 100 gene annotations. Clades are marked by a chromosome number. The tree is rooted at midpoint. (C, D) Distribution of synonymous substitution rates (D_s_) derived from homologous sequences between Otava and C88 (C), and within the single C88 genome (D). Evolutionary events are highlighted by arrows and labels.

To obtain the *k*-mer and gene trees, we utilized the pangenome graph to extract shared entities and used these to calculate pairwise distances between sequences. From shared *k*-mers in the DBG we established a sequence-level *k*-mer distance tree. The tree shows 12 clades (Fig. 3B), corresponding to the number of *S. tuberosum* chromosomes. None of the chromosome numbers assigned to the sequences conflicted with another, supporting a correct topology. The fourth tree was based on gene absence/presence identified from homology groups (Supplementary Fig. S14B). This tree stands out as it shows that all Chr 1 haplotypes of a genome cluster together.

The hybrid origin of *S. tuberosum* explains the extensive conflict observed in the phylogenetic relationships [37]. Removing the lower quality genomes will certainly improve the phylogenetic signal. Nevertheless, truly resolving the potato taxonomy calls for more complex models such as phylogenetic networks that consider reticulation events [38].

To explore the WGD history of *S. tuberosum* we calculated synonymous (D_s_) substitution rates of homologous genes, between and within genomes (Supplementary Fig. S15). The distribution of the intergenomic synonymous substitution rates between Otava and C88 reveal three clearly visible peaks that can be linked to evolutionary events (Fig. 3C). The youngest and highest peak (D_s_ 0.001-0.01) derived from orthologous pairs indicates the speciation time of the two *S. tuberosum* species. Paralogous regions that originated in the *Solanaceae* aleohexaploidy appear as a second peak (D_s_ 0.6-0.9). A third, weak peak (D_s_ 2-3) provides evidence of the eudicot paleohexaploidy (γ) event.

The distribution of D_s_ substitutions is generally only reported between genomes. Therefore, we were interested in observing the intragenomic mutation patterns, comparing haplotypes within a single genome. The intragenomic D_s_ distribution of C88 (Fig. 3D) shows three peaks that look nearly identical to the distribution between C88 and Otava. This triple-peak pattern of mutation rates was not specific to C88 but was found in all intragenomic comparisons of the phased genomes (Supplementary Fig. S15). This was notable because in unphased (haploid) genome assemblies, synonymous substitution plots reveal only whole-genome duplication events. A plausible explanation for the visibility of the youngest peak (D_s_ 0.001-0.01) is, that as a result of phasing, alleles located on the different haplotypes are now aligned whereas in unphased assemblies only duplicated genes are aligned. Although the first peak in intergenomic comparisons is associated to speciation, in intragenomic haplotype-resolved assemblies it reflects heterozygosity between subgenomes. These results demonstrate that pangenomic evolutionary analyses offer far more insight when performed at the subgenome level.

### E. Extensive haplotype-specific variation revealed in ancient polyploids

Comparing the genomic organization of organisms uncovers the genomic conservation and rearrangements, which provide insights into evolutionary dynamics of genomes. The homology grouping, together with established phylogenetic relationships in the pangenome, serves as a framework to analyze genome organization. Here, we discuss several PanTools methods to examine structural conservation and changes among *Malus* chromosomes; in Supplementary Analysis 4, we apply these same approaches to the potato pangenome.

Through synteny analyses we determined pairwise conserved collinearity between all sequences in the pangenome. The macrosynteny suggests high collinearity, with blocks spanning nearly entire chromosomes. However, at the microsynteny-level, between syntenic blocks additional genes were not collinear but rather just fall between two syntenic anchors. To this end, we developed a visualization of both large-scale genome structure analysis (macrosynteny) and conservation of local gene content and order (microsynteny). Retention is calculated in sliding windows based on homology (conserved gene sequence) or synteny (conserved gene order). Two examples on apple demonstrate the use of these visualization methods and show the high diversity between the haplotypes and reveal the fate of duplicated chromosomes post-polyploidization.

Retention patterns based on *M. sylvestris* Chr 11A (Fig. 4A) as query were representative for the majority of *Malus* visualizations; one chromosome pair (green lines) showing high retention of syntenic genes, with another pair (red lines) having lost most collinear gene pairs. The two most similar haplotype sequences represent chromosomes homeologous to the selected query. Less retained sequences are remnants of the most recent WGD. Overall, gene retention w.r.t. the homeologous chromosomes was high, although with strong local fluctuations. The strongest loss of synteny was observed in *M. sieversii* Chr 11A around genomic position 10Mb (region i), where collinearity was fully lost. The observable retention pattern of the WGD-duplicated regions relative to the reference is highly similar between all three retention plots. Two prominent exceptions of regions displaying disparate synteny levels are Gala 10-13 Mb (region ii) and *M. sieversii* 7-9 MB (region iii), as no syntenic gene pairs were found within these regions. The visualizations support hypotheses of fractionation initially quickly degrades duplicated regions, but further advances at a diminishing rate [39].

**Fig. 4.**
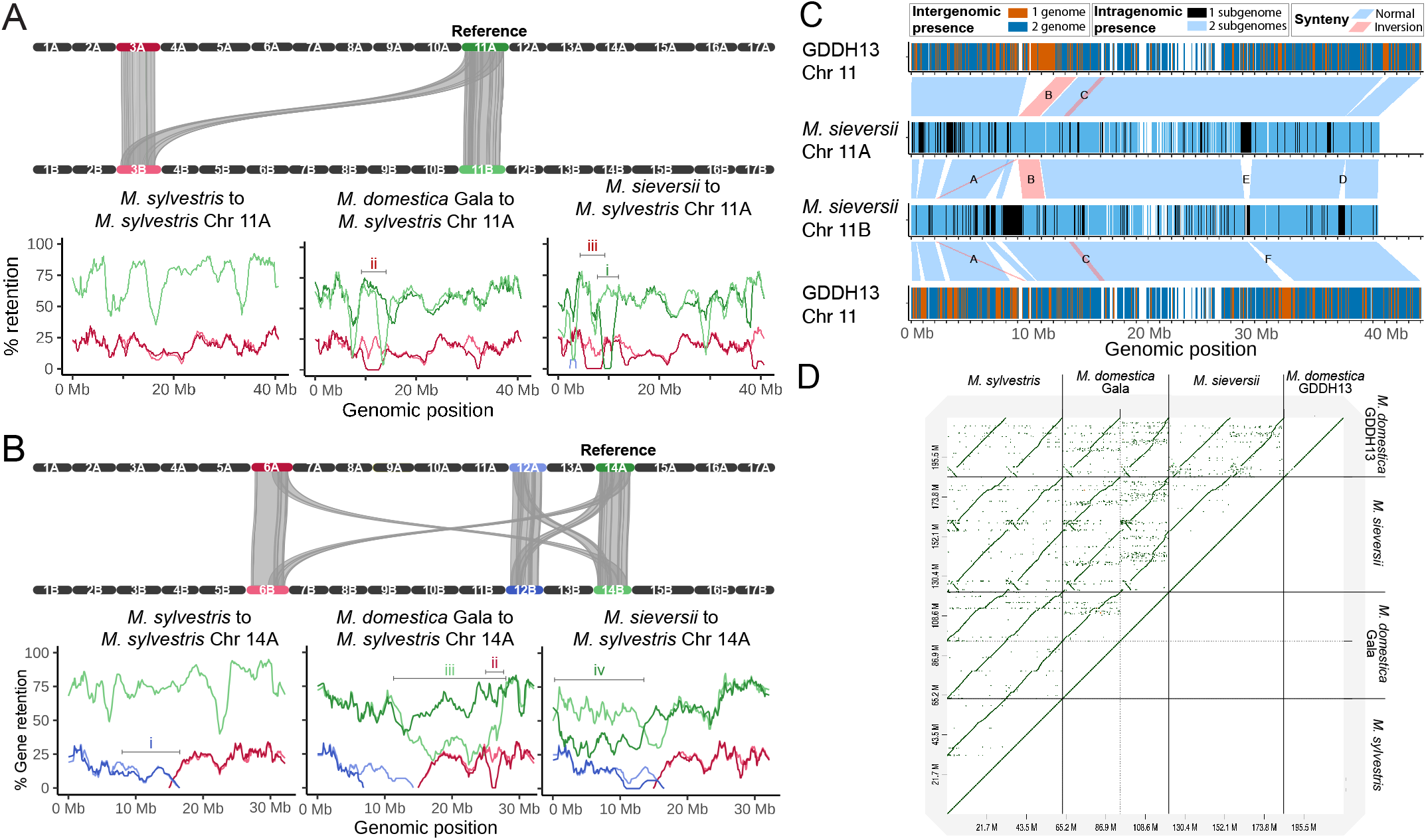
Structural variant visualizations on the *Malus* pangenome. (A, B) Syntenic gene retention of *M. sylvestris* Chr 11A (A), 14A (B) to every sequence of the pangenome. Some regions showing a divergent retention pattern were numbered and are discussed in the text. A schematic representation of macrosynteny between *M. sylvestris*’s chromosomes above each trio of retention plots shows the underlying genomic organization. Synteny relations are drawn between the selected reference sequence and all collinear regions and the corresponding chromosome in the other haplotype. (C) Genetic and structural variation in *M. sieversii* Chr 11, in relation to GDDH13. There are three different types of annotation bars. Starting from the top: GDDH13 Chr 11 gene regions shared with *M. sieversii* 11A (blue) or not (red); syntenic blocks; and gene occurrence in one (black) or both (blue) haplotypes. (D) Dot plot visualization of *Malus* Chr 1 alignments.

In a second example, we explore the retention plots using *M. sylvestris* Chr 14A as query. The WGD-duplicated regions were fragmented and located on two different chromosomes, indicated by the red and blue lines in Fig. 4B. Such rearrangements are frequently seen in WGDs, after which polyploids may gradually return to a diploid state [40]. Different models involving diploidization have been proposed to explain the chromosomal organization of diploid *Malus* species [41], [42]. The level of gene retention in these WGD-derived segments were highly similar across genomes and displayed just two outlying patterns. Only *M. sylvestris* retained the collinearity of duplicated genes on Chr 12 (region i, 10-15 Mb) on both haplotypes. Conversely, Gala Chr 6B (region ii, 25-26 Mb) was the only region that lost all genes syntenic with the reference sequence. Aside from this rearrangement, there was substantial fractionation in the homeologous chromosomes (green lines) illustrated by a high loss of synteny in nearly half the chromosome in both Gala (region iii) and *M. sieversii* (region iv). In Supplementary Analysis 5 we examine another *Malus* chromosome query that showed the strongest reduction of duplicated regions and also revealed a translocated region.

We developed a novel visualization functionality that combines genomic and pangenomic features extracted from the graph database. In Fig. 4C we show an example of Chr 11 from *M. sieversii* and GDDH13 where we combine intra- and intergenomic gene absence/presence variation with synteny annotations. *M. sieversii*’s Chr 11 was selected because of the high number of intragenomic rearrangements. Earlier, Chr 11 of *M. domestica* GDDH13 was used to assist phasing the *M*.*sieversii* assembly [11]; therefore, we included it here to place the variation in the context of a reference. The visualization shows the large blocks of unshared genes correlate to the intra- and intergenomic synteny breakpoints. Synteny blocks further reveal three major inversions. The leftmost inversion (marked by letter A) was specific to sequence 11B and appeared to be translocated as well. The second haplotype-specific inversion (B) in 11A (9-11Mb, 130 genes) was the longest identified structural variation between any *Malus* chromosomes. The third inversion (C) was intergenomic and therefore only visible (around position 13-14Mb) between *M. sieversii* and GDDH13.

Aside from inversions, synteny relationships display multiple breakpoint regions. All synteny breakpoints were due to haplotype-specific insertions/deletions, emphasizing the importance of intragenomic variation. Synteny was lost around position 37 Mb (D) because of a nearly 1Mb sized region in which no gene is shared. Perhaps even more intriguing is the two-sided breakpoint (E) identified at position 28Mb, where both haplotypes show a distinct region of only non-homologous genes. The second half of this non-syntenic region (F) on Chr 11B was the only block of haplotype-specific genes that broke synteny to the GDDH13 reference. Possibly, this could be a genomic fragment introgressed into the *M. sieversii* genome, or it could have been lost from all other haplotypes. Considering all other haplotype-specific regions in *M. sieversii* are syntenic to GDDH13, it is most likely these synteny breakpoints are the result of collapsed haplotypes.

With PanTools’ visualization utility, we created an image for each haplotype-resolved chromosome set of apple and potato to provide a comprehensive view of the intragenomic variation (Supplementary data). These visualizations display gene regions with coloring according to the presence in number of subgenomes, with synteny relationships drawn between the chromosomes. Each image offers a clear overview of gene-absence variation and gene order conservation in sets of two (apple) or four (potato) homologous chromosomes.

Another perspective on intragenomic and intergenomic variation across a set of chromosomes is offered by a multi-genome dot plot. Dot plots are popular visualizations to identify large scale deletions, inversions, and repeats. Using the earlier established phylogenetic relationships, only homeologous chromosomes were aligned to another. In Fig. 4D. we show the dot plot visualization of all *Malus* Chr 1 haplotypes in the pangenome. Notably, both GDDH13 as *M. sieversii* have an 5MB inversion in regard to the other *Malus* genomes. This inverted region in *M. sieversii* displays another small inversion that is most clearly visible against *M. sylvestris*’s chromosomes.

Graphically representing the genomic organization with structural variation supports a better understanding of the complexity of genomes. The introduction of new PanTools features allows for users to create both novel and more traditional plots to display chromosomal rearrangements. The presented examples demonstrated extremely high variation between haplotypes. Regardless of whether the observed variations reflect true biology or assembly artefacts, these visualizations can provide valuable support for comparative genomic analyses.

### F. Exploring allelic diversity

A desirable feature of pangenomes is the ability to identify all allelic variants of genes for functional selection and breeding. We demonstrate novel functionalities for this purpose on the potato gene *StCDF1. S. tuberosum* originates from a region close to the equator and its adaption to short-day growing conditions prevents tuber formation in the long-day conditions during spring and summer in higher latitude locations. The transcription factor CYCLING DOF FACTOR 1 (*StCDF1*) is a key regulator for reaching maturity and tuber formation [43]. Potato plants adapted to longer day lengths have specific *StCDF1* allelic variants [43].

The pangenome database was utilized to find all allelic variation in *StCDF1* among the potato cultivars. First, *StCDF1* was identified in DM Chr 5, where it clustered in a homology group of 22 proteins. The hierarchal pangenome annotations allowed the extraction of not only the protein sequences but also the encoding gene, transcript and CDS sequences. The 22 proteins derived from 17 genes. Manual BLAST against the genome assemblies verified the presence of 17 *StCDF1* loci. Fig. 5A provides an overview of protein sequences within the homology group, showing which alleles are present in each subgenome.

**Fig. 5.**
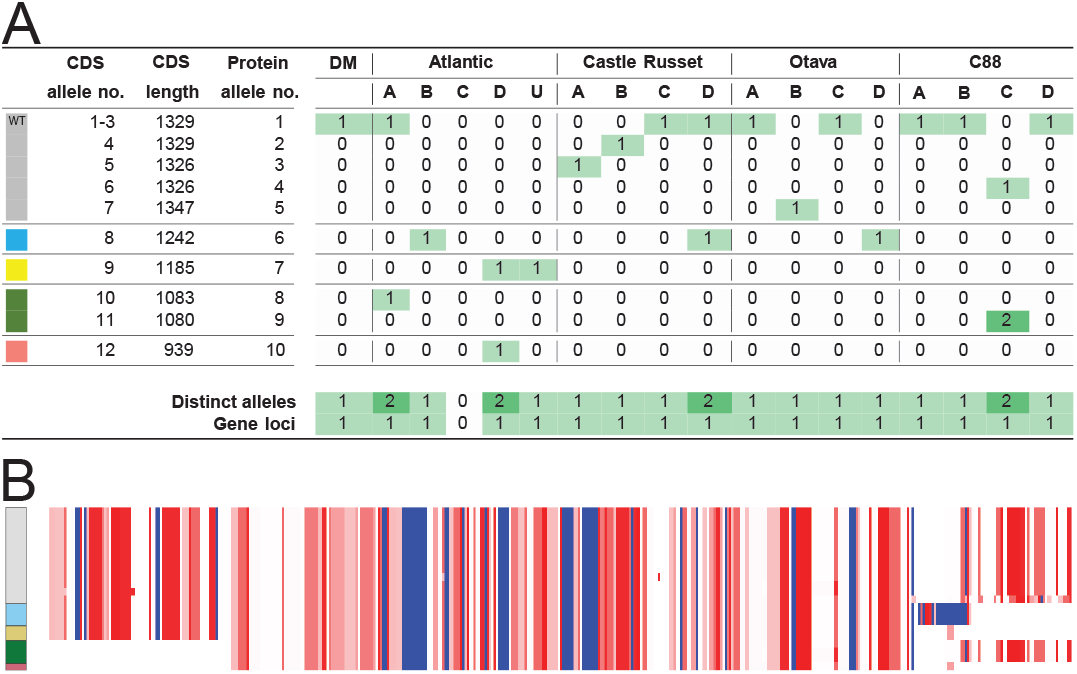
(A) Occurrence of *StCDF1* alleles in *S. tuberosum* subgenomes. Below each column the total number of unique alleles and genes in a subgenome is given. (B) Alignment of 22 *StCDF1* protein sequences (visualized via https://alignmentviewer.org). Amino acid residues are colored by hydrophobicity, where (dark) red is most hydrophobic and blue is most hydrophilic. Assigned groups (colors) based on truncations and insertions are shown on the left of the alignment.

Through protein sequence alignment five major *StCDF1* allelic groups were distinguished based on truncations and insertions (Fig. 5B). The wild-type (WT) allele (*StCDF1*.*1*, protein allele no. 1) in the grey group coding for full-length *StCDF1* proteins was the most abundant and found at least once per genome. In the C88 cultivar the WT is present on 3 subgenomes, while in Atlantic it was limited to one. Blue and yellow groups encode proteins with a 3’ (C-terminal) truncation; blue group genes had new coding sequences inserted. The green group was characterized by a 5’ (N-terminal) truncation, and consists of one Atlantic gene and two C88 genes. Lastly, the pink group gene in Atlantic had both 5’ and 3’ truncations.

Apart from the wild-type allele *StCDF1*.*1*, alleles *StCDF1*.*2* to *StCDF1*.*5* have been identified earlier at the locus [44]. These alleles carry specific insertions, leading to FKF domain truncation in the 3’ region, thereby avoiding ubiquitination [43]. Even though *StCDF1* is intensively studied, to our knowledge, so far no *StCDF1* alleles were reported with a truncated 5’ region. While our overview shows a total of 3 truncated 5’ alleles, we remain cautious as the prediction of gene models is highly complex and often results in incorrect annotations. The Atlantic and C88 genomes, in which these transcripts were found, were annotated following comprehensive strategies supported with sufficient transcript evidence [12], [19]. Upon inspecting the gene models we found that the 3 alleles with 5’ truncations are protein isoforms. These isoforms arose from variations in the exon–intron boundary predictions of gene regions, resulting in different models. As follow-up, the gene-models were examined in the Spud DB Jbrowse instance (http://spuddb.uga.edu). Mapped leaf and tuber ONT cDNA reads showed very minimal support for alternative splicing in the two Atlantic genes (Supplementary Fig. S16). Moreover, C88 lacked transcript/cDNA data for validation.

A potato plant reaches maturity when its tubers are fully developed and ready for harvest, the timing of which varies strongly between cultivars. Early maturity of the Atlantic cultivar is attributed to through multiple 3’ truncated alleles [12], not necessarily the 5’ truncation. The C88 variety, despite having four *StCDF1*.*1* alleles, can still reach maturity in long-day conditions [45], [46]. C88 has two non-WT alleles. One is still considered *StCDF1*.*1* and shows a 3 bp deletion outisde the FKF domain that is unlikely to affect gene functionality. This leaves the truncated 5’ allele as a potential candidate variant for C88’s long-day acclimation.

To conclude, PanTools helped identify allelic diversity in *StCDF1*, distinguished by major truncations in the 3’ and 5’ regions. Further validation of the diversity found is necessary, requiring greater sequencing depth or more extensive experimental validation of the gene.

## III. Concluding DISCUSSION

Genomic analyses provide an important foundation for our understanding of biology. Our current ability to resolve genomes at the haplotype level provides a representation that is far more accurate than the collapsed genomes studied thus far. However, methodology to easily analyse collections of such genomes is still scarce. We updated the PanTools pangenomics platform and added new functionalities to represent phased genome assemblies and enable identification of intragenomic variation. We demonstrated these functionalities on tetraploid potato and diploid apple pangenomes, showing both practical applications and the potential for plant breeding (e.g. maturity in potato).

The most critical factor for accurate pangenome analysis is the quality of genome assemblies. We showed that BUSCO completeness should be assessed at subgenome-level to assess phasing quality, and that visualization of subgenomes with polymorphisms is essential to decide whether certain variation actually reflects biology or results from an assembly/phasing artifact. The defined subgenomes lack any biologically linkage, making them only applicable to assess gene presence and frequencies in random haplotype combinations. Our analyses further revealed high heterozygosity in potato and apple, characterized by gene absence/presence variation and structural rearrangements. Overall, our results demonstrate the usefulness of combining a pangenome representation with state-of-the-art bioinformatics tools for detailed intra- and intergenomic analyses.

In the past decade, we have seen a rise of many pangenomic software toolkits [47]. PanTools distinguishes itself through its hierarchical pangenome graph and by providing an extensive set of comparative genomics functionalities, partially through a connection to existing tools. PanTools facilitates comprehensive pangenome analyses, from the initial quality checks to the downstream analyses. In this study, we introduced a new set of functionalities specific for haplotype-resolved assemblies. These were based on typical comparative genomic analyses, but we adopted a pangenomic approach for their implementation.

A pangenome represents all variation found in a population and provides a valuable overview of all available alleles. The application of pangenomics methodology as currently implemented in PanTools to haplotype-resolved genomes is already promising, but further development will allow fuller exploration of this rich source of data. More interactive visualization methods will facilitate visual analytics, i.e. user-guided exploration of (sub)genomic architecture and structural variations. This approach should allow users to choose any chromosome as reference and zoom into specific regions of interest, enabling the visualization of complex genetic patterns.

We can also use the current pangenomic representation framework for analysing additional sources of data. PanTools was designed to easily include such diverse data types to enable the study of complex biological systems. We observed that only half of the genes in genomes are found on all subgenomes, often with high variation among alleles. Our pangenome representation can overcome limitations imposed by a reference bias and facilitate analyses hampered by a reference bias, such as the identification of allele-specific expression.

PanTools’ hierarchical pangenome graph holds potential for exploring the evolutionary history. In this study we observed extensive fractionation after whole-genome duplications events in two autopolyploids pangenomes. In contrast, allopolyploids have highly divergent parental genomes, often leading to one subgenome becoming dominant over the other [48]. PanTools can currently identify biases in gene content and fractionation; however, studying subgenome dominance requires the integration of expression and epigenetic data [48]. The advent of more haplotype-resolved genomes will drive the methodological development needed to perform such analyses, which will help obtain an increasingly detailed picture of pangenome content, organization and evolution.

## IV. Methods

### A. Genome and annotation data collection

We focus on two use cases, a potato (*Solanum tuberosum*) and apple (*Malus*) pangenome. Both use cases were built upon publicly available datasets. For the potato use case, all data was downloaded directly from Spud DB (http://spuddb.uga.edu) in January 2022. Two phased Atlantic and Castle Russet assemblies were obtained from the study of Hoopes et al. [12]. Genomic data of Otava were derived from the Sun et al. [18] paper. C88 data was accompanied by the publication from Bao et al. [19]. The only unphased potato genome was the most recent (v6.1) assembly of DM 1-3 516 R44 created by Pham et al. [20]. For apple, all three haplotype-resolved genomes *M. domestica* (v1.0), *M. sieversii* (v1.0) and *M. sylvestris* (v1.0) were obtained from the study of Sun et al. [11]. Two additional phased genomes were obtained; the apple reference genome *M. domestica* GDDH13 (v1.1) collected from the publication by Daccord et al. [21] and a wild apple genome *M. baccata* assembly (v1.0) by Chen et al. [22]. Original GFF annotation files of *M. baccata* and DM were invalid and were updated with the AGAT toolkit (agat_convert_sp_gxf2gxf.pl) [49] to include missing ‘gene’ features.

### B. Pangenome construction and annotation

Potato and apple pangenomes were built and analyzed using the ‘phased_pangenomics’ development branch in the PanTools repository. Here, we briefly discuss the PanTools functions and their respective arguments used to perform the analysis. For more detailed explanations of the underlying algorithms, we refer to the online manual (https://pantools.readthedocs.io). The pangenomes were constructed with the ‘build_pangenome’ function and a *k*-mer size *k* of 19. Structural annotations were connected to the De Bruijn graph (DBG) with ‘add_annotations’. The gene, transcript and protein sequences created by PanTools were compared to the extracted sequences of AGAT (agat_sp_extract_sequences.pl) [49]. Chromosome and haplotype information was added to the database with ‘add_phasing’. Species names were included as metadata using ‘add_phenotypes’. Transposable element annotations derived from EDTA (v2.0.0) [50] with default settings were included into the pangenome database through the ‘add repeats’ function.

### C. Determining the optimal homology grouping

PanTools’ ‘busco_protein’ function assessed completeness of the pangenome using BUSCO (v5.3.2) [32] with the most specific lineage datasets; *Solanales* odb10 for potato and eudicots odb10 for the apple genomes. We assessed the completeness of genomes and separate subgenomes. First, ‘busco_protein’ was run without any additional arguments to evaluate the entire proteome of each genome assembly. Second, by setting the ‘–phasing’ and ‘–longest-transcripts’ arguments, BUSCO was performed against proteome subgenome subsets only including the longest protein-coding transcripts of genes.

Proteins were clustered seven times with different strictness with the ‘optimal_grouping’ [51] functionality. Clustering strictness is altered by changing the minimum required (normalized) similarity score between two sequences and tweaking the parameters controlling MCL (Markov clustering) [52]. We calculated the *F*_1_ score (the harmonic mean of precision *p* and recall *r, F*_1_ = 2(*p·r*)*/*(*p*+*r*)) for the seven groupings based on BUSCO-identified singlecopy genes that are present in every subgenome. Assuming these genes are truly single-copy, a perfect clustering would place them in a separate homology group with one representative protein per subgenome. Following this assumption, we scored the grouping based on the BUSCO genes that actually cluster together and whether other genes cluster with them.

As a second measure to assess the protein clustering, we quantified the degree of overlap among gene sets between haplotypes of the same subgenome. First, a Jaccard index is calculated from shared genes within homology groups for the possible sequence combinations in a subgenome. Subsequently, the average distance in the subgenome was calculated from all combinations, followed by taking the mean of the individual averages.

### D. Phylogenetic analyses

PanTools facilitates multiple methods to represent genomic distances among genomes or individual sequences in the pangenome. A *k*-mer distance tree was created for all chromosome-length sequences in the pangenome using the ‘kmer_classification’ method. MASH distance [53] is calculated between two sequences by counting the shared *k*-mers in the DBG. The pairwise distances were stored in a matrix and served as input for inferring a Neighbor-Joining (NJ) tree using ape (v5.0) [54]. The tree topology was validated by checking if chromosome numbers (included in genome FASTA headers) conflict within clades.

The ‘core_phylogeny’ method was used to create one sequence-level tree per chromosome. Single-copy groups were identified through ‘gene_classification’, in which only sequences belonging to a specific chromosome were included. The sequences of single-copy groups were aligned with MAFFT (v7.453) [55] and trimmed to avoid noisy regions near the end of the sequence. The multiple sequence alignment (MSA) was performed in two steps. After an initial protein alignment, the longest start and end gaps were used to trim the nucleotide sequences. These trimmed sequences were input for the second alignment. A concatenated sequence of parsimony-informative single-nucleotide polymorphisms (SNPs) was created per haplotype. IQ-tree (v1.6.12) [56] was applied to the collection of concatenated sequences with 10,000 bootstrap replications.

With the ‘consensus_tree’ function, one tree per chromosome was generated that summarizes all gene trees associated to a chromosome. Homology groups shared by all sequences of a specific chromosome were first identified as input. To obtain gene trees, sequences were aligned as described in ‘core_phylogeny’. FastTree (2.1.10) was applied to the homology group MSAs using default parameters. The gene trees were combined in a file, from which ASTRAL-Pro (version March 2022) [57] with preset configurations estimated a consensus tree.

Gene distance calculated by ‘gene_classification’ is the third type of distance to create a sequence tree for a single chromosome. Jaccard distances were obtained by counting shared genes and total genes between two genomes using the homology groups. Only unique elements were considered, and additional gene copies were ignored. Gene distance matrices were visualized as NJ trees created by ape (v5.0) [54]. All phylogenetic tree visualizations were created with iTOL v6 [58].

### E. Gene based analyses

PanTools’ ‘gene_classification’ function with the ‘–phasing’ argument was used to characterize the pangenome gene content. The homology grouping serves as the foundation for all analyses of this section. Genes were intergenomically categorized as follows: core genes were present in all genomes, accessory genes were absent in some genomes, and cloud (formerly called unique) genes were found in a single genome. For the intragenomic characterization we estimate if gene presence is in line with the ploidy of an organism, and count its occurrence in each chromosome set (1 to 2 in apple and 1 to 4 in potato). Furthermore, separate countings were performed for the individual chromosomes. For instance, a gene is found in 2 out of 4 Otava (potato) Chr 1 haplotypes. This intragenomic counting required incorporated haplotype information via ‘add_phasing’. Several examples on how these classification rules were applied are included as Supplementary Fig. S2.

Homology groups were further used to determine the frequency of genes and alleles. Gene copies were directly counted from these groups. For the allele frequency we considered every nucleotide or amino acid polymorphism within a group to represent a distinct allele. The nucleotide and protein sequences of a group were collected in two separate sets from which the unique elements were counted.

### F. Synteny estimation and graph integration

Synteny blocks were computed with the ‘calculate_synteny’ function. First, the GFF and homology input files required for MCScanX (version October 2020) [29] were generated. The (highly simplified) GFF files only contain the sequence identifiers together with gene start and stop coordinates, belonging to a single sequence. The homology files state which genes are homologous to another between two sequences, and were created for every possible sequence combination. Iterating over the input files, MCScanX was employed in parallel using default settings. Subsequently, separate output files were combined into a single collinearity file, which was included into the database using ‘add_synteny’. Genes part of the same syntenic block were connected in the graph through ‘synteny’ nodes, syntenic gene pairs gained a direct ‘is_syntenic_with’ relationship to each other.

### G. Estimate synonymous substitutions rates

Synonymous (D_s_) and nonsynonymous (D_n_) mutation rates were calculated of homology group MSAs. The alignment was performed in two rounds, as described in the “Phylogenetic analyses” section above. PAL2NAL (v14) [31] was used to convert protein alignments into corresponding codon alignments, codons with gaps and inframe stop codons were excluded. Sequences shorter than 30 amino acids were excluded to minimize artifacts caused by short alignments. D_n_ and D_s_ values were calculated in the codon alignments with codeML (part of PAML package [59]).

### H. Gene retention visualizations

The retention pattern visualization were created with PanTools’ ‘gene_retention’ function. To calculate retention of all sequences to a selected query sequence, the following steps are performed. First, all gene nodes of the query sequence are collected and ordered based on their genomic position. Then, a sliding window of 100 genes moves over the nodes in steps of 10. The window stops when it no longer can move 10 genes to the right, resulting in the visualization of full-sized windows only. At each window position, the percentage of retention is calculated based on shared homologs and syntelogs between the query to every other sequence. Genes are considered homologs when part of the same homology group, while syntelogs are required to be part of the same synteny block and form a syntenic pair. To ensure retention does not exceed 100%, syntenic depth is ignored, as it is highly influenced by gene duplications. Window positions were transformed into genomic coordinates of the query sequence, at which the retention values were plotted with ggplot2 [60].

### I. Whole genome alignment

Sequences belonging to the same chromosome (number) were aligned with minimap2 (v2.24) [30] using the ‘-x asm5’ parameter. Variants were called between two sequences with paftools.js and filtered out when the quality was below 10. The genome alignments were visualized with D-GENIES (v1.4) [61], where up to the 100,000 best matches per alignment were plotted.

### J. Sequence visualizations

The chromosome visualizations based on various annotations were created with PanTools ‘sequence_visualization’. The function can generate five types of annotation bars. First, gene regions were colored by their presence in other genomes, in concordance to the homology group category: core, accessory or cloud. Second, gene regions were colored based on their presence in the number of subgenomes within a single genome. Third, gene regions were colored gray if a gene was found on any other chromosome. Still, gene copies on the same chromosome (number) but different haplotype do not allow the region to be coloured grey. In the fourth annotation bar, a line graph plots the coverage (percentage) of the repeat and gene regions (annotations) in 1Mb windows. Repeat coverage of 100% indicates every nucleotide in the window overlapped with at least one repeat annotation. The fifth and final annotation represents synteny blocks that allow to connect two sequences. Annotation bars were plotted individually with ggplot2 [60] and stacked horizontally using the cowplot R package (https://github.com/wilkelab/cowplot).

## Supporting information

Supplementary Analyses

## V. Authors’ CONTRIBUTIONS

EJ and SS designed the study. EJ and SS were responsible for extending the functionality of PanTools, and EJ carried out the computational analyses. Result and interpretation, shaping the manuscript, were discussed with the domain experts: EJ, TL, LB, JH, SS. The manuscript was written by EJ and DR participated in revising the manuscript. All authors critically reviewed the manuscript.

## VI. Acknowledgements

We would like to thank Richard Finkers for the helpful feedback and for testing the new PanTools functionalities. We also want to thank Christian Bachem for the insightful discussions.

## VII. Supplemental INFORMATION

**Supplementary figures:** Supplementary figures S1-S16 supporting the main text.

**Supplementary analyses:** Supplementary analyses 1-5.

**Supplementary data:** Files for reproducibility and PanTools’ ‘sequence_visualization’ output for the haplotype-resolved assemblies.

### Data availability

PanTools v4 is available at https://git.wur.nl/bioinformatics/pantools, released under the GNU GPLv3 license. The novel functionalities were developed in the ‘phased_pangenomics’ branch, which was merged into ‘pantools_v4’ at commit d5eca936 (https://git.wur.nl/bioinformatics/pantools/-/commits/) and released in v4.3.0 (https://git.wur.nl/bioinformatics/pantools/-/releases/v4.3.0). Instructions for downloading the publicly available datasets and reproducing the experiments, are in Supplementary data.

