## Supplementary Figures for "Exploring intra- and intergenomic variation in haplotype-resolved pangenomes"

Supplementary figures belonging to Jonkheer et al. "Exploring intra- and intergenomic variation in haplotype-resolved pangenomes"

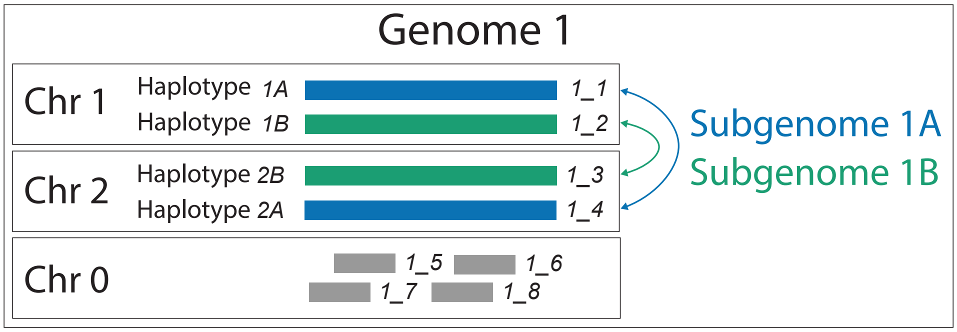


Supplementary Figure S1. Sequence terminology illustrated based on an example genome of 8 sequences. PanTools tracks sequences via a genome number combined with a sequence number (identifiers in italic). Haplotype identifiers consist of the chromosome number with haplotype phase. Haplotype information was included for sequence 1 to 4. A subgenome corresponds to all haplotypes with a certain letter within a genome.

| **Homology group** | Phenotype 1 | | | | | Phenotype 2 | | | | **Definition** |
|---|---|---|---|---|---|---|---|---|---|---|
| ***K*-mer** | Genome 1 | Genome 2 | | | | Genome 3 | | | | |
| **Function** | - | 1A | 1B | 2A | 2B | 1A | 1B | 2A | 2B | |
| 1 | 1 | 0 | 0 | 0 | 1 | 1 | 0 | 0 | 0 | Core |
| 2 | 1 | 1 | 0 | 0 | 0 | 0 | 0 | 0 | 0 | Accessory |
| 3 | 1 | 0 | 0 | 0 | 0 | 0 | 0 | 0 | 0 | Cloud (Unique) |
| 4 | 1 | 1 | 2 | 1 | 1 | 0 | 0 | 0 | 0 | Phenotype specific |
| 5 | 1 | 1 | 0 | 0 | 0 | 1 | 0 | 0 | 0 | Phenotype shared |
| 6 | 1 | 0 | 0 | 0 | 0 | 0 | 0 | 0 | 0 | Phenotype exclusive |
| 7 | 0 | 0 | 0 | 0 | 0 | 1 | 0 | 0 | 0 | 1/2 haplotypes |
| 8 | 0 | 0 | 0 | 0 | 0 | 1 | 1 | 0 | 0 | 2/2 haplotypes |
| 9 | 0 | 0 | 0 | 0 | 0 | 1 | 0 | 1 | 0 | 1/2 haplotypes & every chromosome |
| 10 | 0 | 0 | 0 | 0 | 0 | 1 | 1 | 1 | 1 | 2/2 haplotypes & every chromosome |
| 11 | 0 | 0 | 1 | 0 | 0 | 1 | 1 | 0 | 0 | 2/2 haplotypes & extra haplotype compared to G2 |

Supplementary Figure S2. Classification rules for genes, $k$-mers and functional annotations. In this example, genome 2 and 3 have phasing information added, both genomes have two chromosomes that consist of two phases. Genome 1 and 2 share a phenotype and genome 3 has its own. The core, accessory, cloud and phenotype categories are defined at the genome level. Thus, having additional copies or being located on multiple subgenomes has no influence on the classification of a gene. Examples 7-11 require chromosome numbers and haplotype letters to be assigned to sequences of the pangenome. Genomes without phasing information were ignored in these examples.

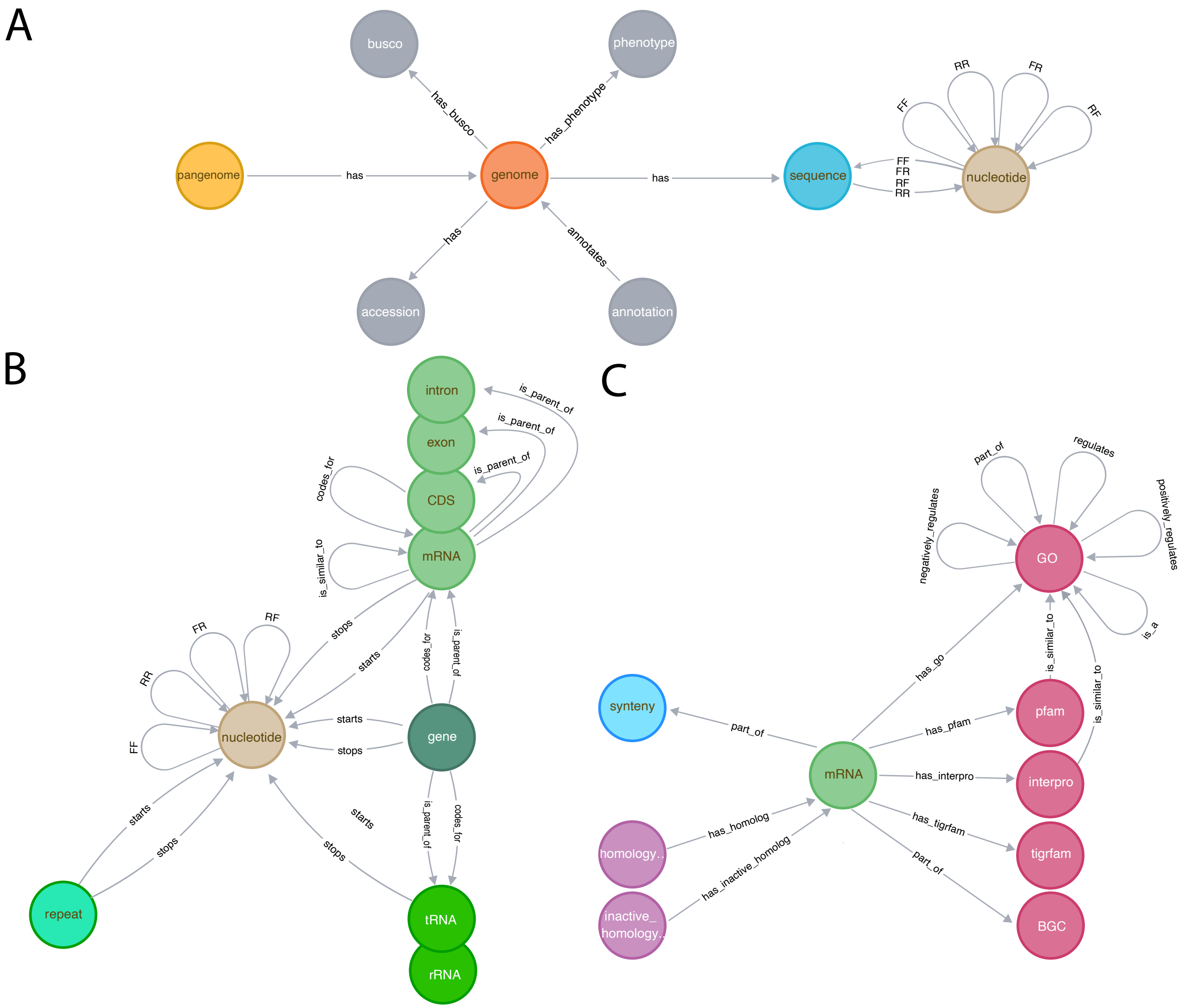


Supplementary Figure S3. The PanTools graph database scheme. The scheme is split into three panels for better interpretability; the nucleotide node connects panel A with B, the mRNA node connects panel B with C. (A) Representation of genomes, annotations, sequences and the compressed de Bruijn graph (cDBG). (B) Structural annotations are built on the DBG. To prevent overlapping edges in this panel, similar annotation nodes are stacked and redundant 'starts' and 'stops' edges are not shown. (C) mRNA nodes can be connected through 3 types of relationships: homology, synteny and functional annotation.

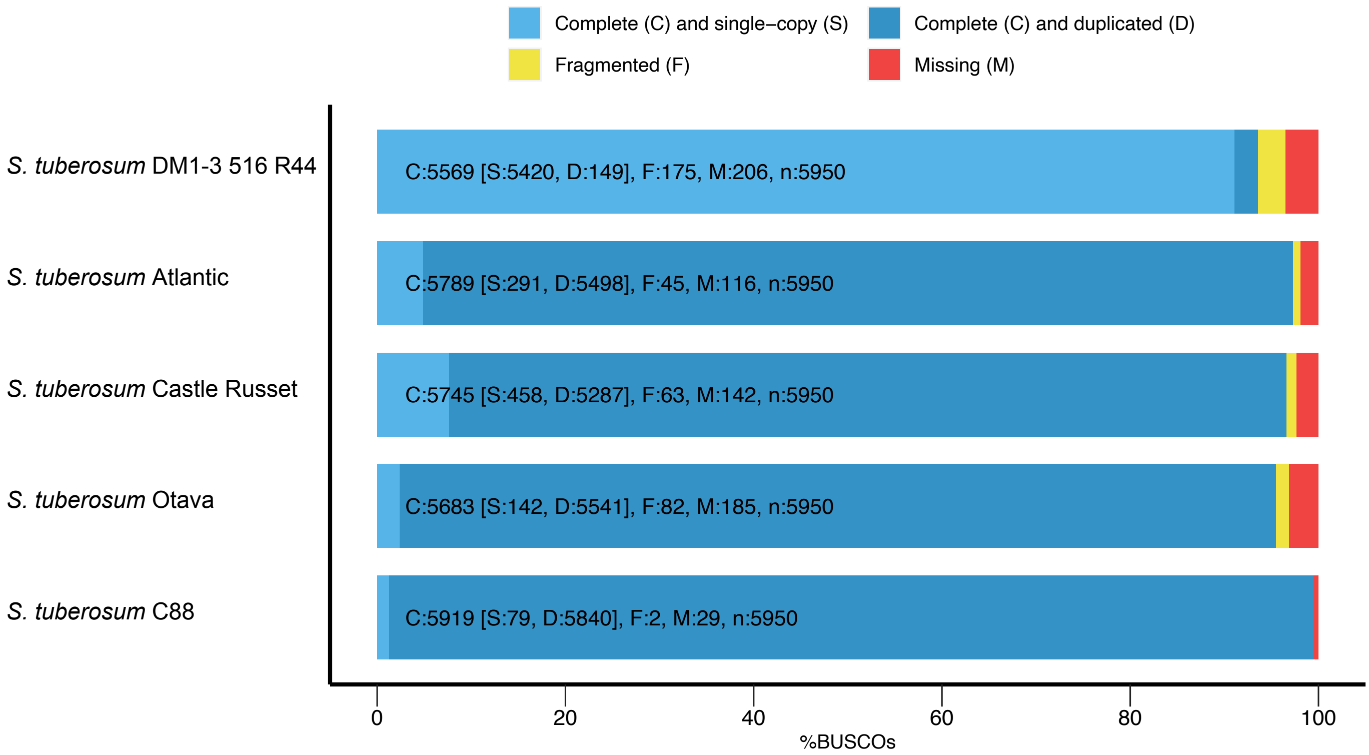


Supplementary Figure S4. BUSCO analysis of five complete *S. tuberosum* proteomes.

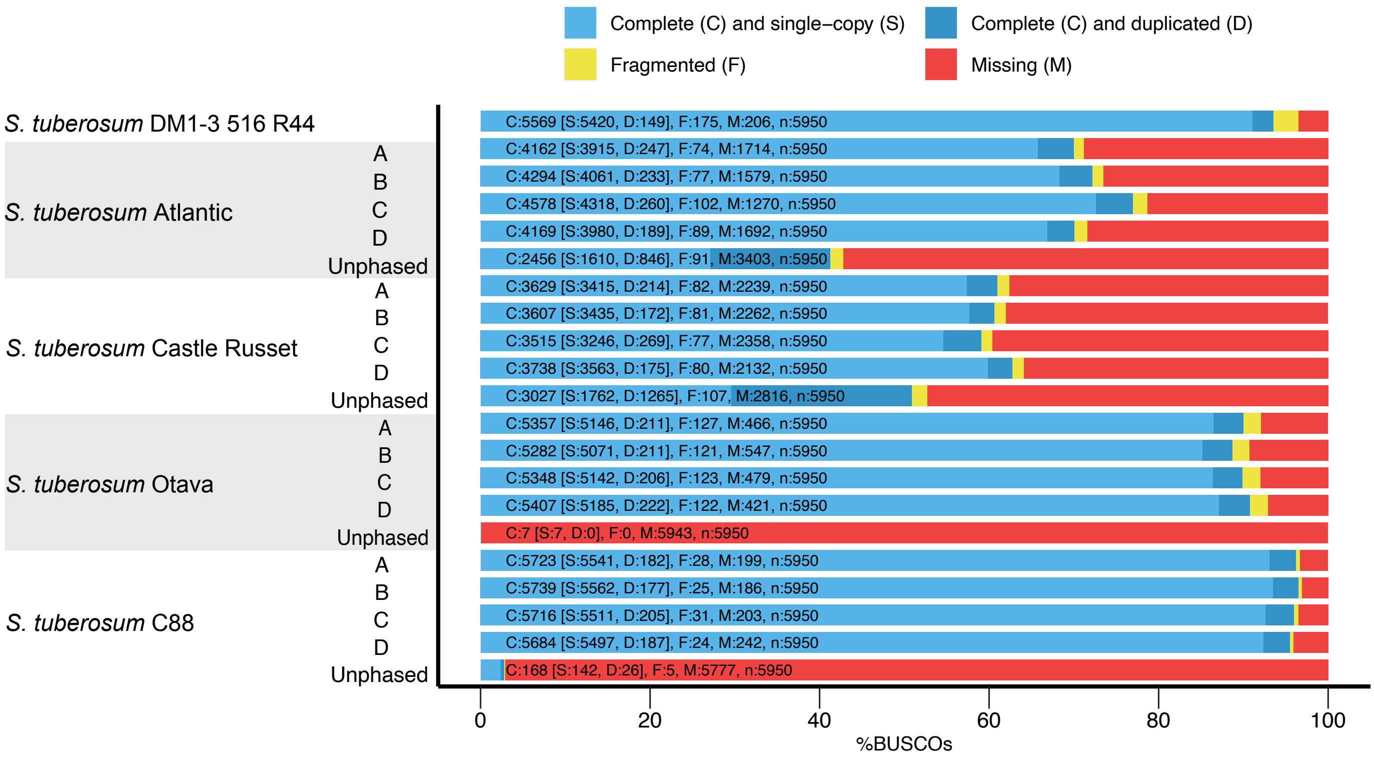


Supplementary Figure S5. BUSCO analysis of subgenome subsets from five *S. tuberosum* proteomes.

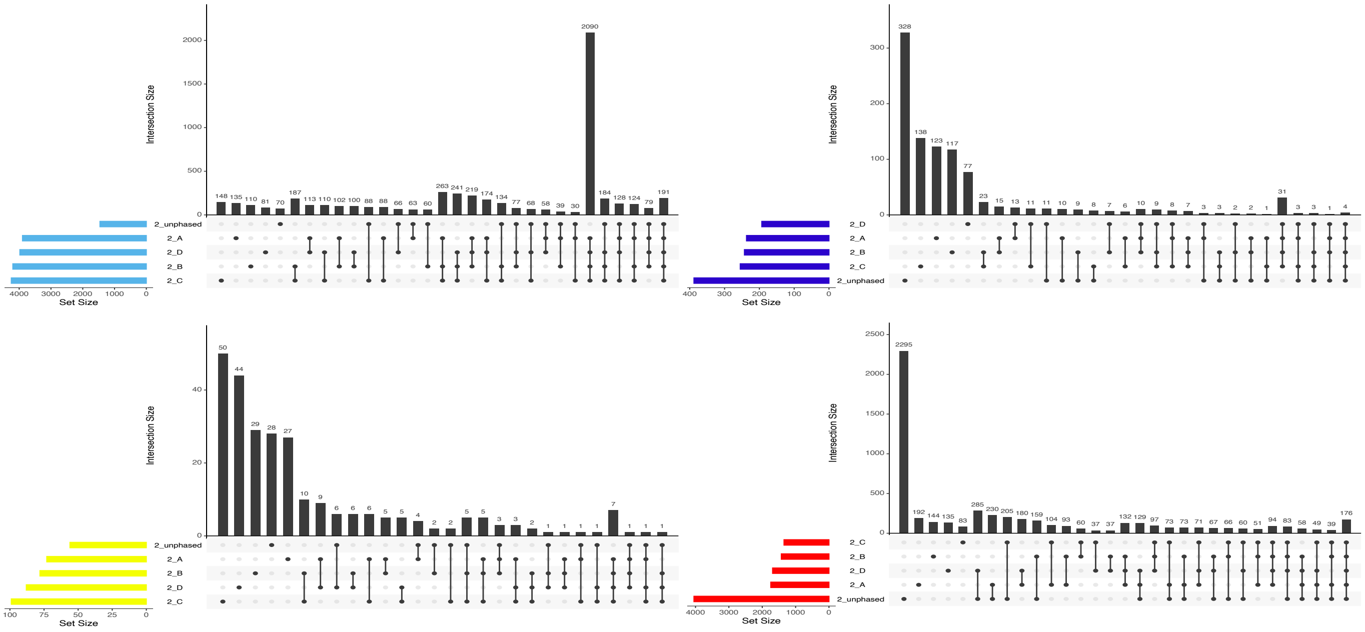


Supplementary Figure S6. BUSCO analysis of *S. tuberosum* cv Atlantic with the solanales odb10 dataset of 5,950 single-copy genes.

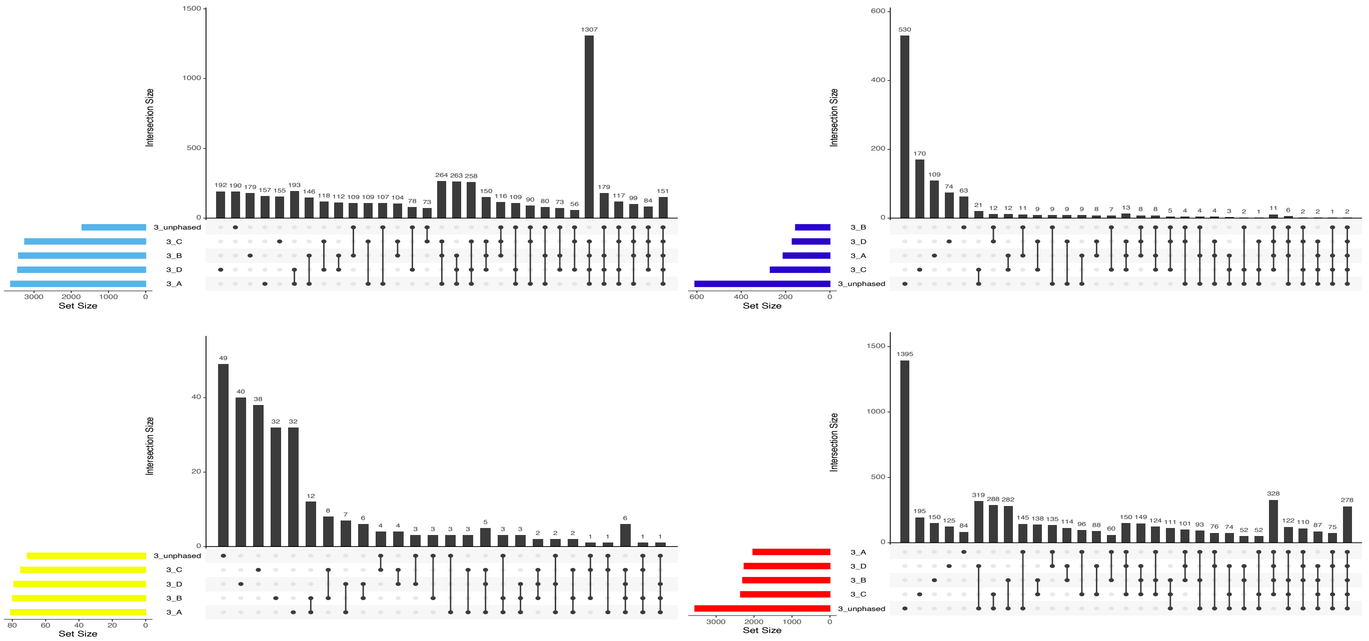


Supplementary Figure S7. BUSCO analysis of *S. tuberosum* cv Castle Russet with the solanales odb10 dataset of 5,950 single-copy genes.

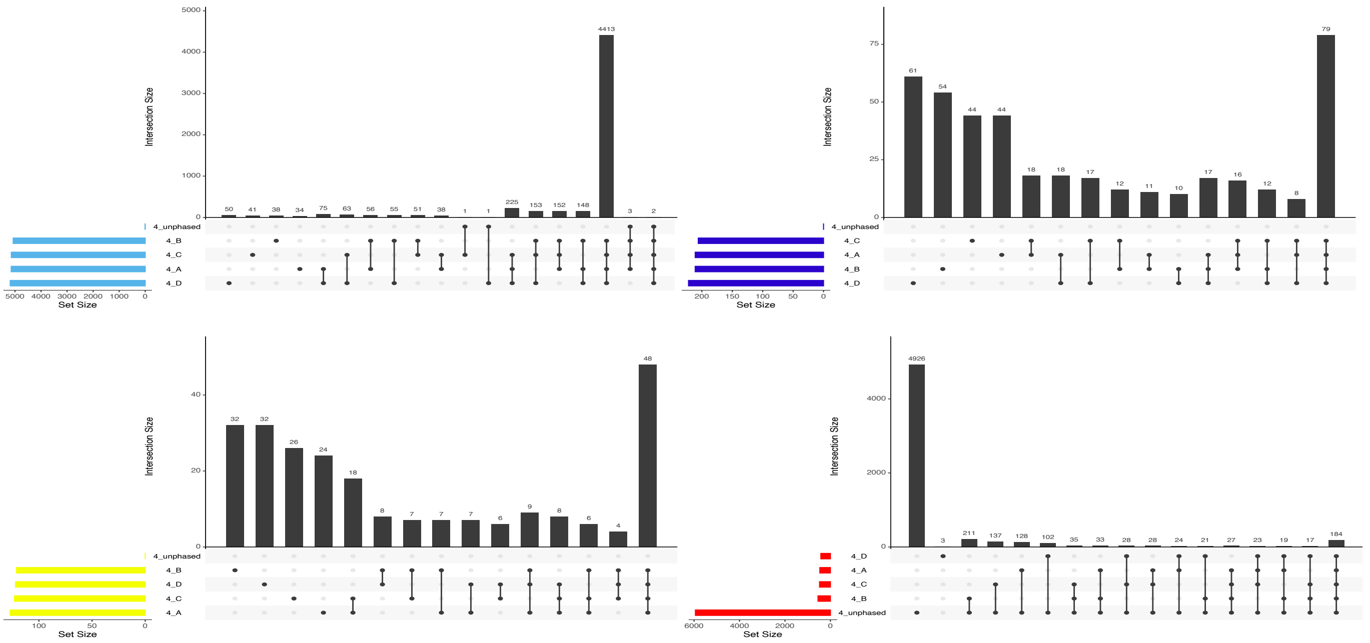


Supplementary Figure S8. BUSCO analysis of *S. tuberosum* cv Ottava with the solanales odb10 dataset of 5,950 single-copy genes.

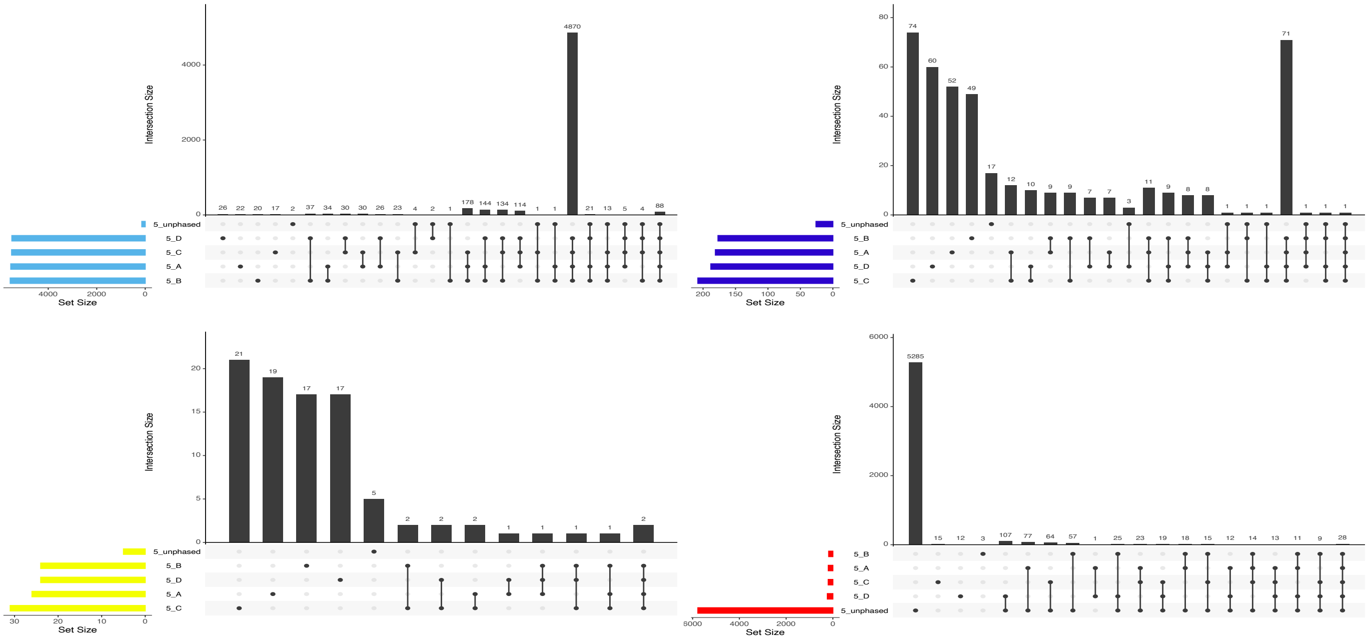


Supplementary Figure S9. BUSCO analysis of *S. tuberosum* C88 with the solanales odb10 dataset of 5,950 single-copy genes.

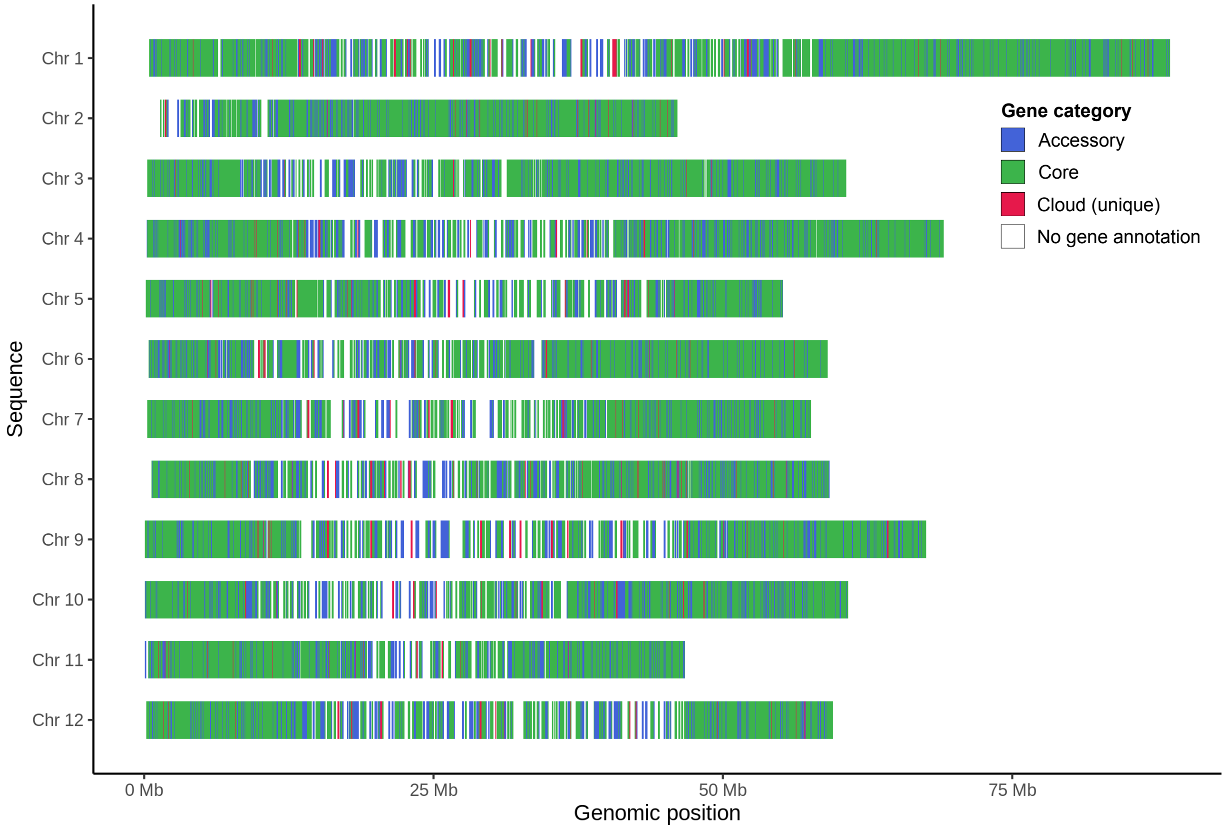


Supplementary Figure S10. Gene regions of *S. tuberosum* DM1-3 516 R44 colored by gene presence shared between five *S. tuberosum* genomes.

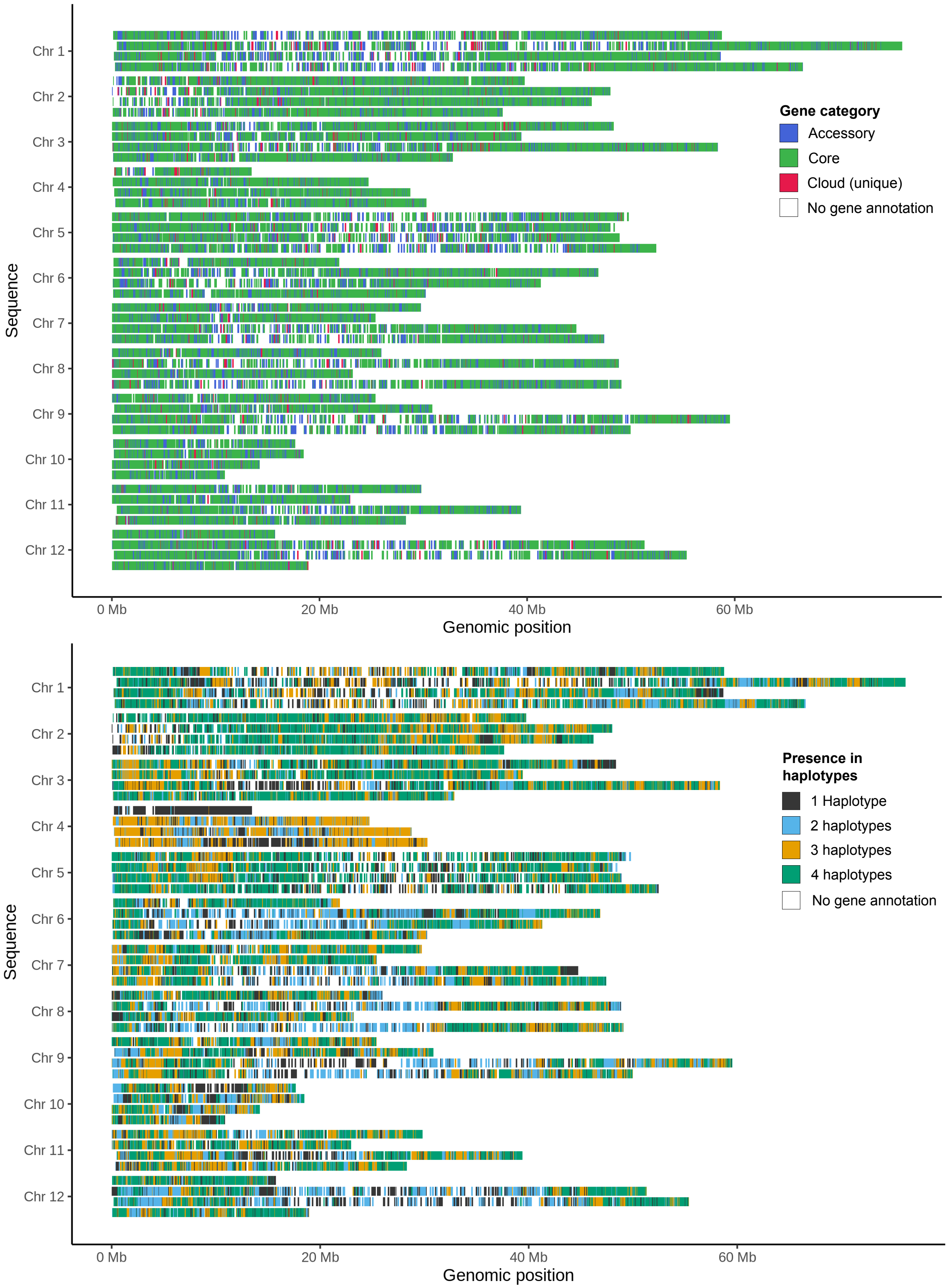


Supplementary Figure S11. Gene regions of *S. tuberosum* Atlantic colored by gene presence shared between five *S. tuberosum* genomes (top figure). Gene regions colored by gene presence in number of haplotypes (bottom figure).

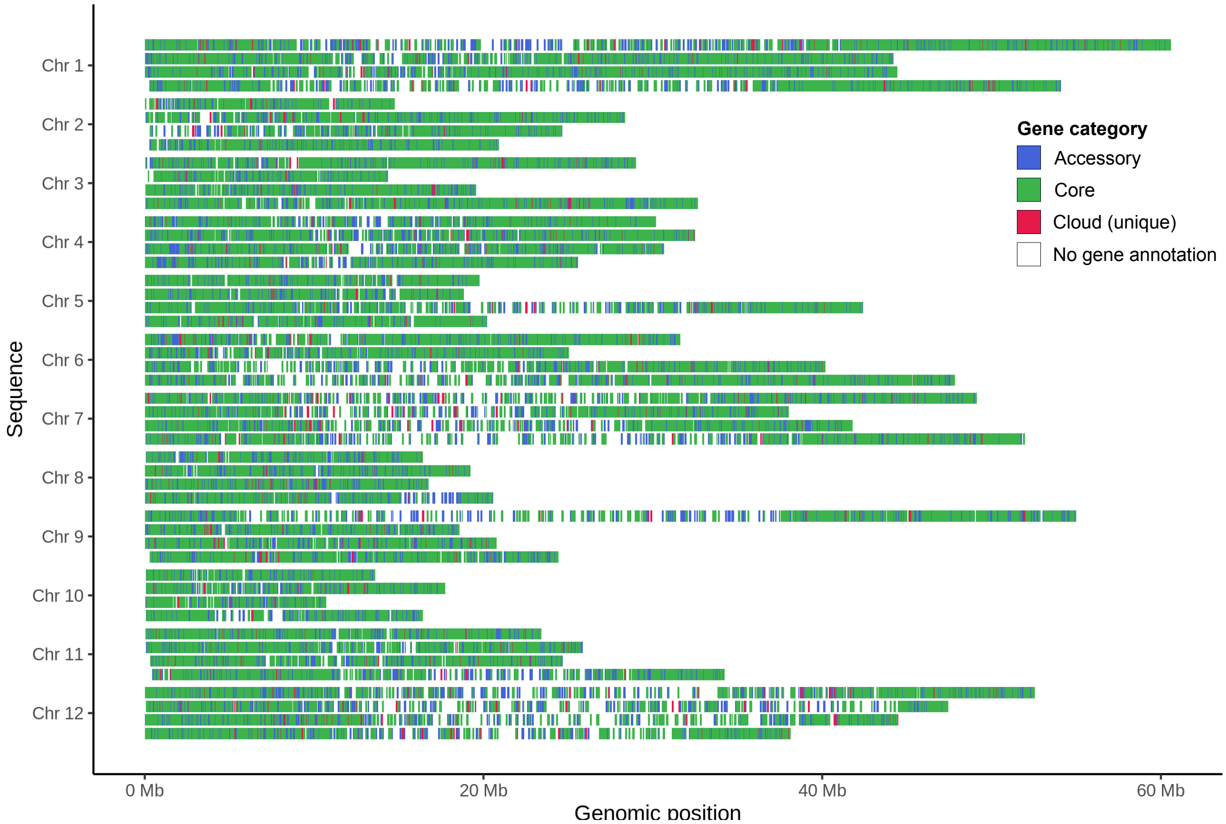


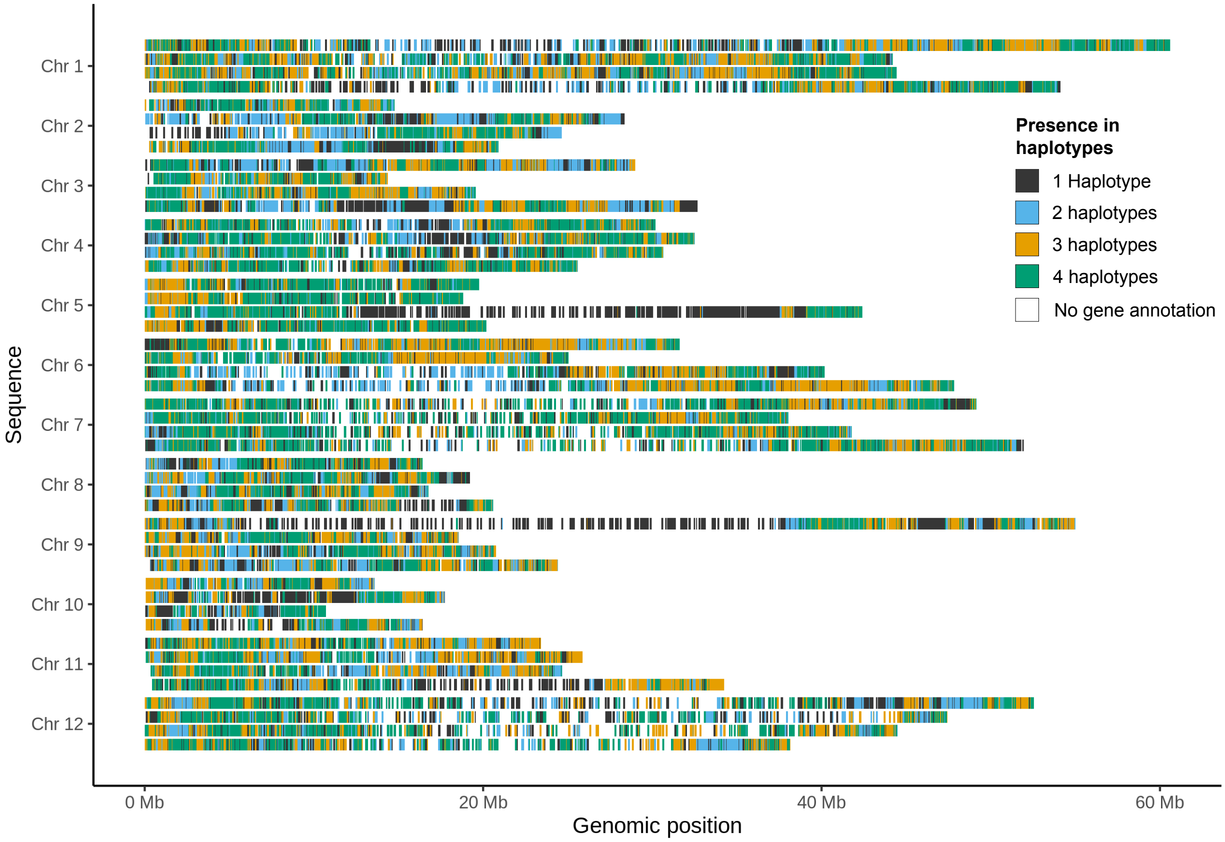


Supplementary Figure S12. Gene regions of *S. tuberosum* Castle Russet colored by gene presence shared between five *S. tuberosum* genomes (top figure). Gene regions colored by gene presence in number of haplotypes (bottom figure).

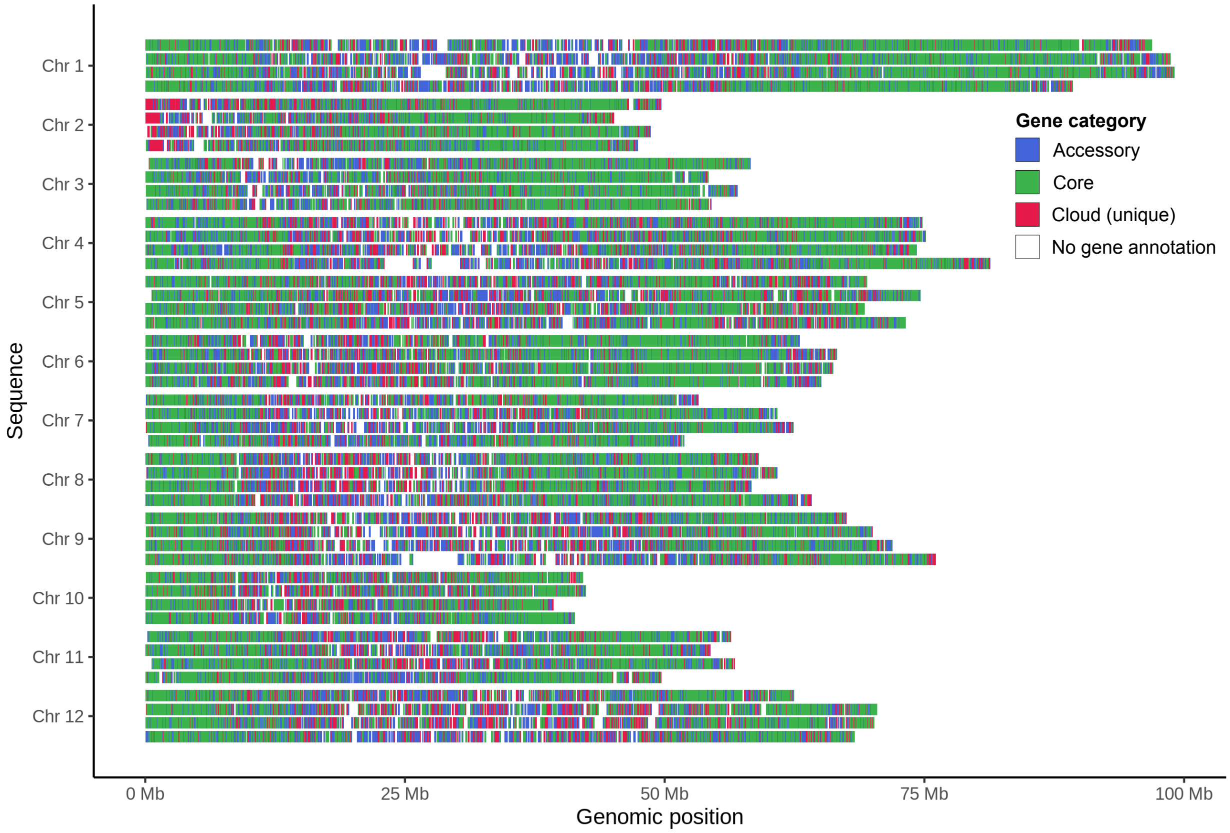


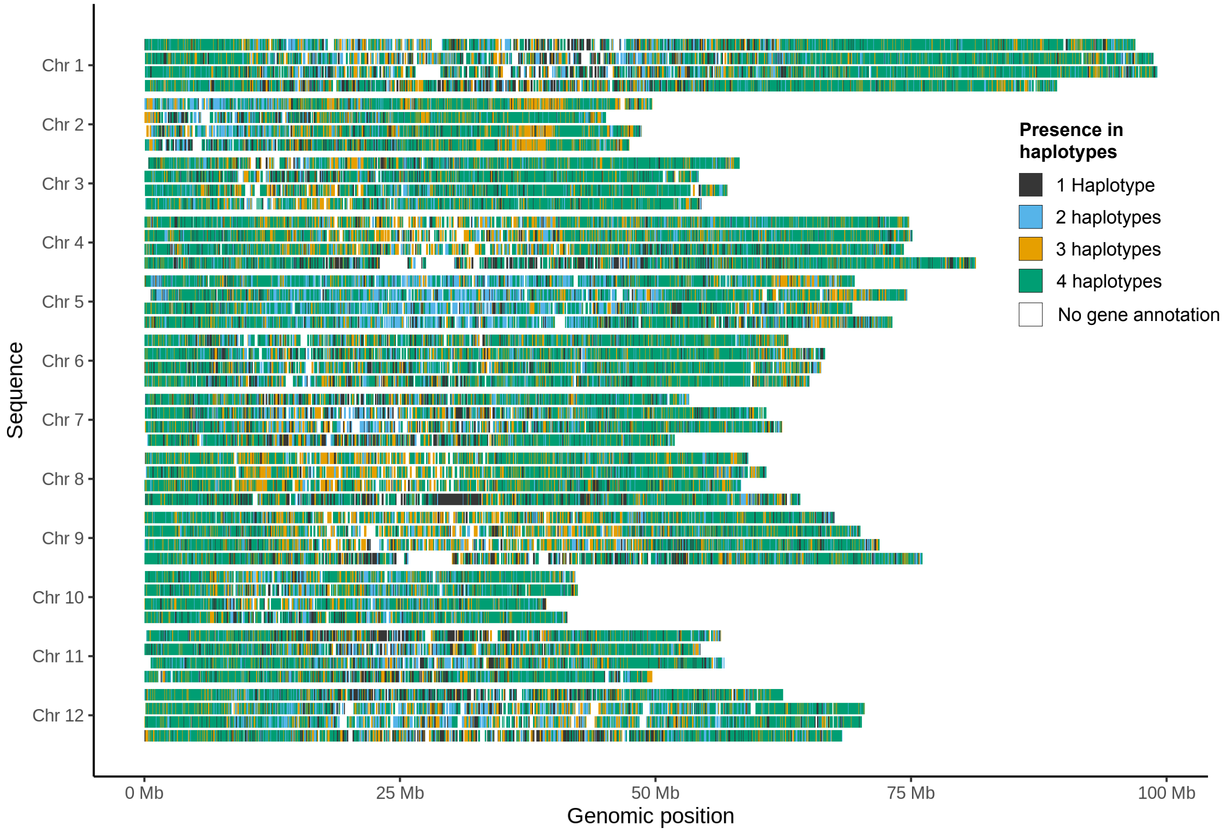


Supplementary Figure S13. Gene regions of *S. tuberosum* Otava colored by gene presence shared between five *S. tuberosum* genomes (top figure). Gene regions colored by gene presence in number of haplotypes (bottom figure).

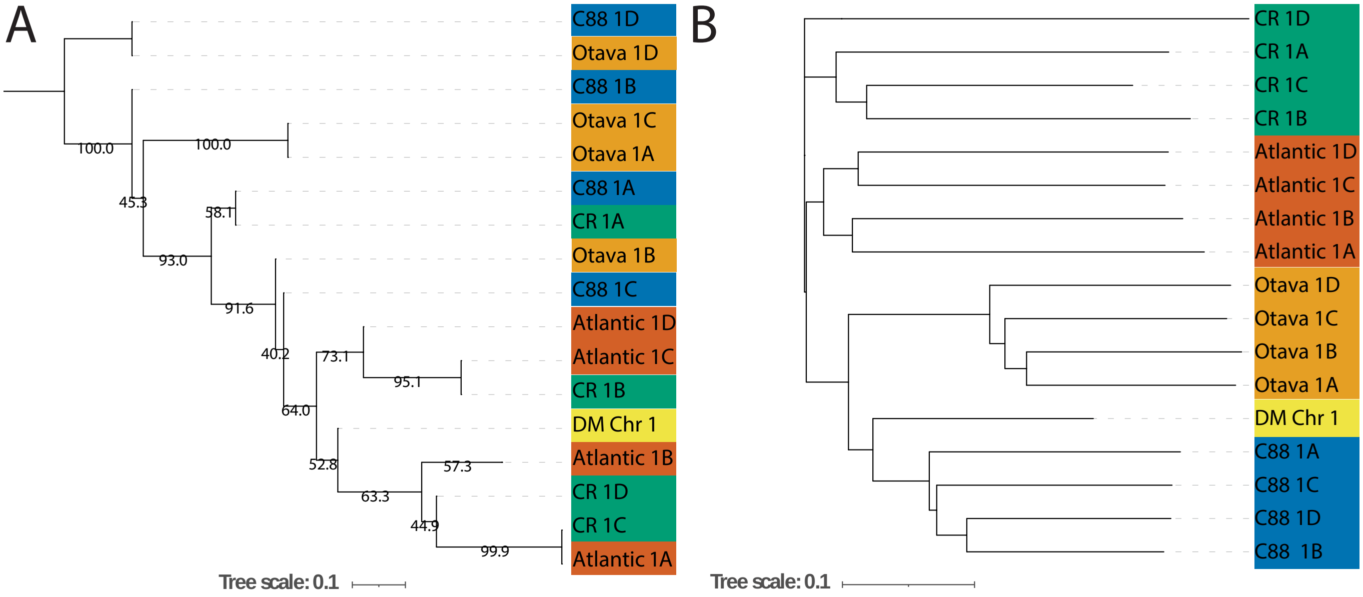


Supplementary Figure S14. *S. tuberosum* Chr 1 phylogenetic trees. (A) Consensus tree of 296 Chr 1 core homology group gene trees. (B) Gene distance tree of Chr 1 haplotypes based on gene absence/presence in 9,012 homology groups. Both trees are rooted at midpoint.

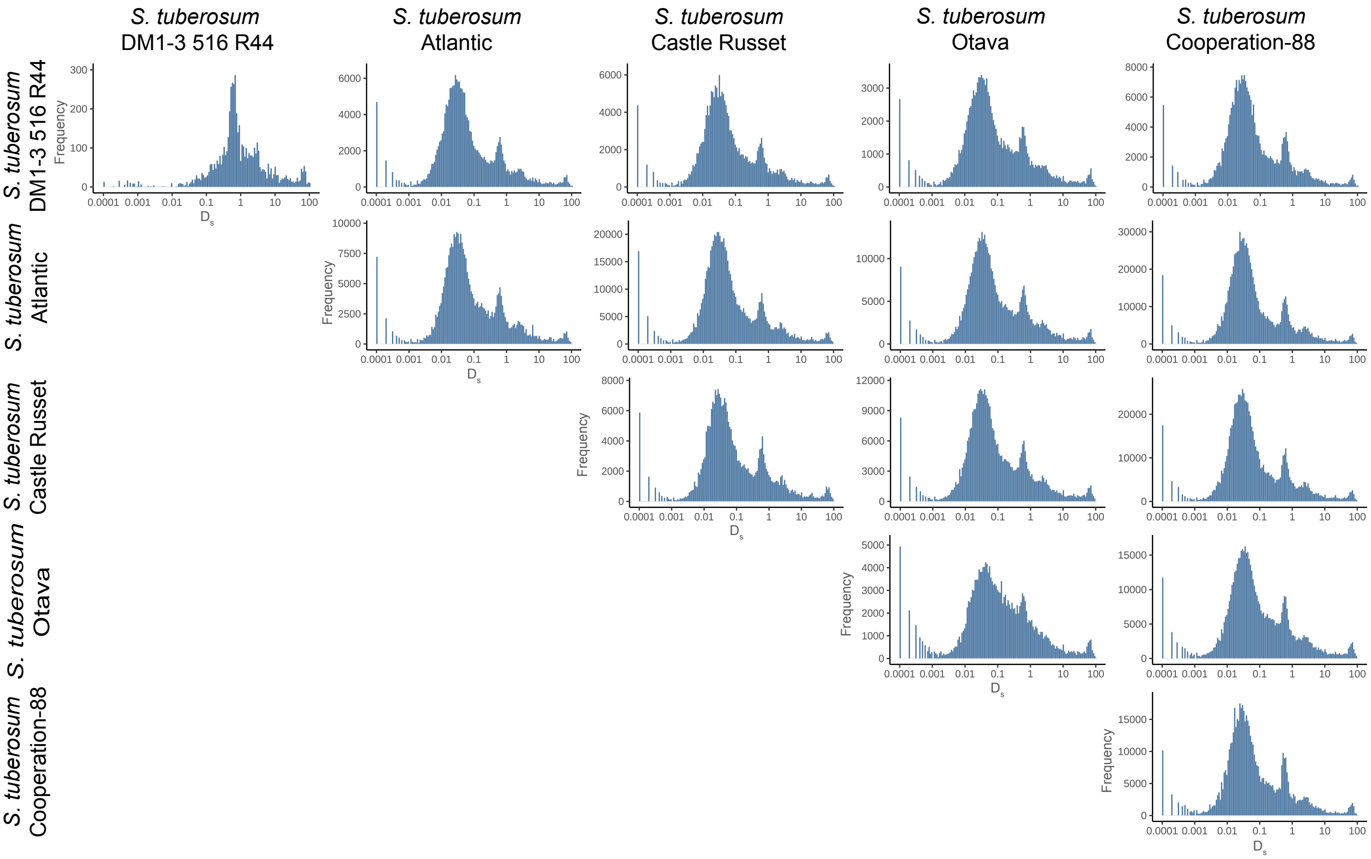


Supplementary Figure S15. Synonymous substitution rate ($D_s$) distributions of homologous sequences between *S. tuberosum* genomes. Values on the xaxis are on a logarithmic (base 10) scale.

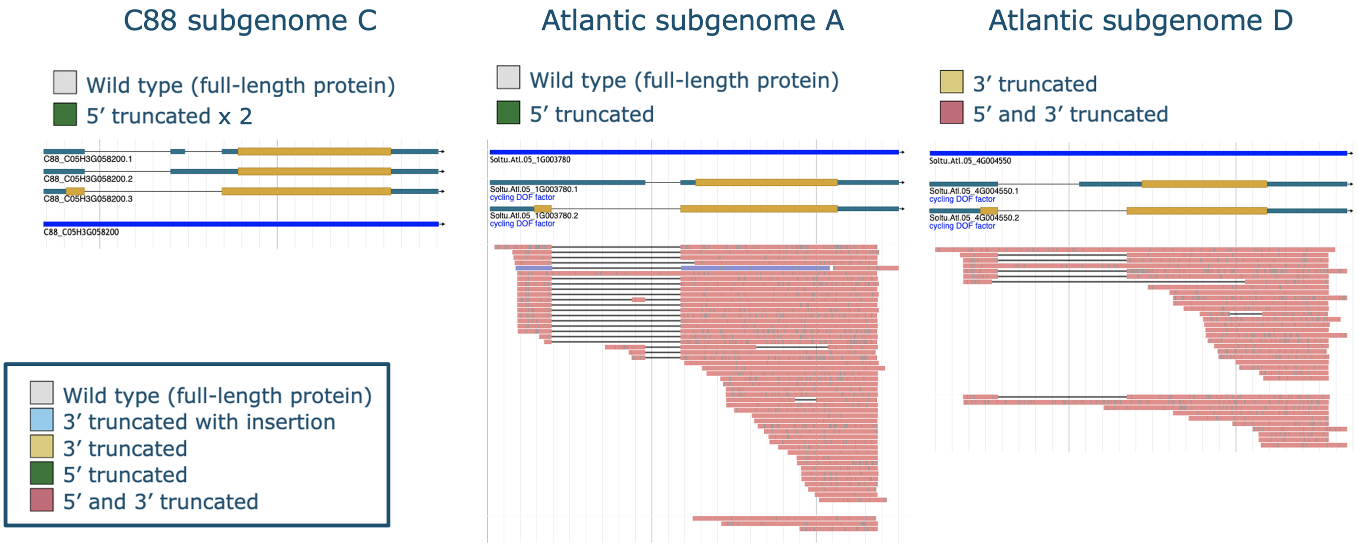


Supplementary Figure S16. *StCDF1* gene models and supporting read data in Spud DB Jbrowse instances of C88 and Atlantic. Screenshots were taken on January 9, 2023. The Jbrowse instances can be accessed via the following links: https://tinyurl.com/ytssy9be (C88), https://tinyurl.com/mv5rztd2 (Atlantic A), tinyurl.com/5n6dsvvx (Atlantic D).
