## Supplementary Analyses for "Exploring intra- and intergenomic variation in haplotype-resolved pangenomes"

### Exploring intra- and inter-genomic variation in haplotype-resolved pangenomes

Supplementary analyses

#### Supplementary Analysis 1. *Malus* pangenome construction

The *Malus* pangenome was constructed from five diploid apple assemblies: *M. domestica* cv. Gala, *M. domestica* 'Golden Delicious' GDDH13, *M. sieversii*, *M. sylvestris* and *M. baccata*. Haplotypes were resolved in the Gala, *M. sieversii* and *M. sylvestris* assemblies.

##### BUSCO completeness assessment

We performed the BUSCO analyses on proteome sets of the complete genomes as well as the separate subgenomes. The initial evaluation of the complete sets exhibited very high duplication rates, ranging between 27.4-34.8% for the unphased genomes and 80.9-86.2% in haplotype-resolved genomes (Fig. 1). For the second assessment, the proteomes of the haplotype-resolved genomes were split into two subgenome (and one unphased) subsets. Application of BUSCO on these sets resulted in a strongly reduced the number of gene duplications, to a similar level (26.7-35.2%) as in the unphased genomes (Fig. 2). Still, all assemblies remain with a notably high fraction of duplicated genes, which support the occurrence of the recent Maleae-specific whole-genome duplication (WGD) [1].

We further used the BUSCO analysis to assess the overlap and identify differences between subgenomes. The overall completeness of subgenomes (Fig. 2) shows that all three B subgenomes have more missing and fragmented genes than A subgenomes. This is most likely a result of the assembly and phasing methods that were used. Running BUSCO on the unphased sequences indicates a notable number of genes that were absent in the haplotypes, which can explain the higher number of missing genes in B subgenomes.

For the haplotype-resolved *Malus* genomes we estimated the intersection of BUSCO genes between their three proteome subsets (A, B, unphased) as shown in Fig. 3-5. There was a substantial overlap between subgenomes, most prominently in single-copy and duplicated genes. Still, each subgenome had its own unique single-copy genes (between 226 and 335). This was similar for duplicated genes, where the majority of gene duplications are present in both subgenomes. Notably, the A proteome subsets showed considerably more duplicated genes that the B subsets.

##### Optimal grouping analysis

*Malus* proteomes were clustered with 7 different PanTools clustering settings (so-called "relaxation modes") and benchmarked on BUSCO genes to determine the optimal homology grouping (Fig. 6). The highest $F_1$-score were obtained using relaxation mode 4 and 5 (Fig. 6B). In addition to the BUSCO statistics, we used the percentage of shared genes between subgenomes as a metric for selecting a suitable grouping (Fig. 6A). We specifically looked at the Gala genome because its two subgenomes appeared to share fewer genes compared to the remaining two genomes. In relaxation mode 4, 52% of Gala's homology groups had genes in both subgenomes, compared to 59% in mode 5. Ultimately, relaxation mode 5 was chosen based on having a higher $F_1$-score and a higher degree of gene content overlap between the subgenomes.

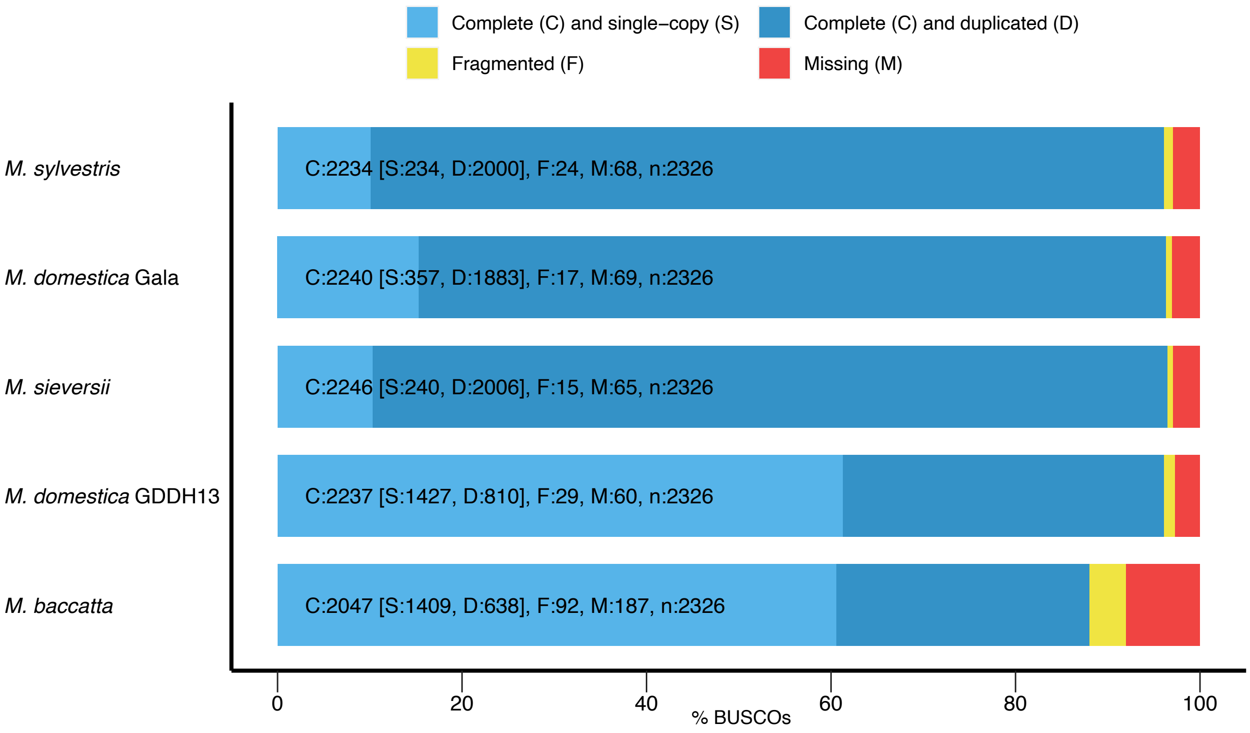


Fig. 1. BUSCO analysis of complete *Malus* proteomes.

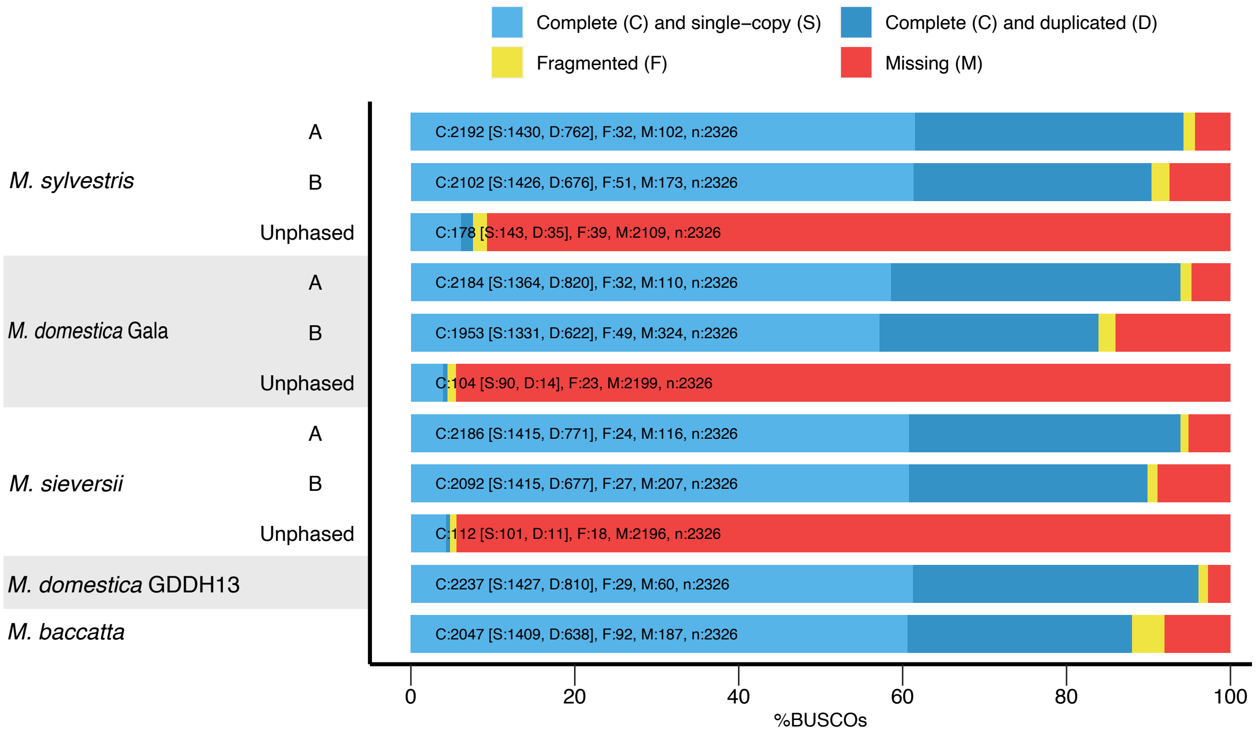


Fig. 2. BUSCO analysis of proteome subsets of *Malus* subgenomes.

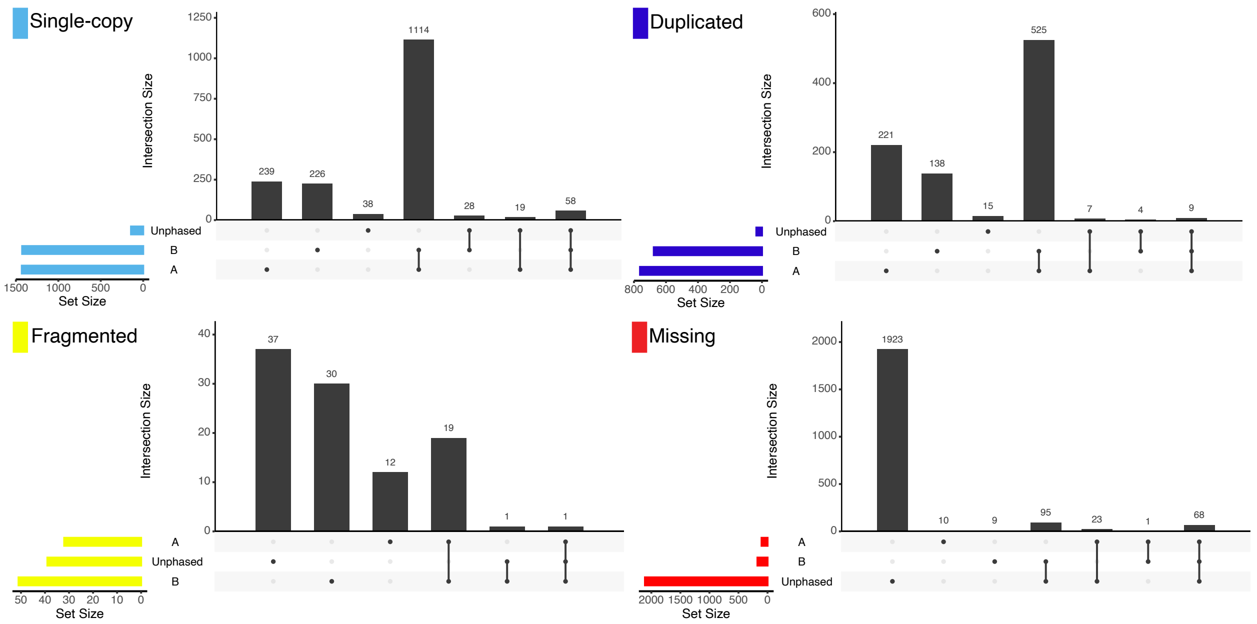


Fig. 3. BUSCO analysis of *M. sylvestris* with the eudicots odb10 dataset of 2,326 single-copy genes.

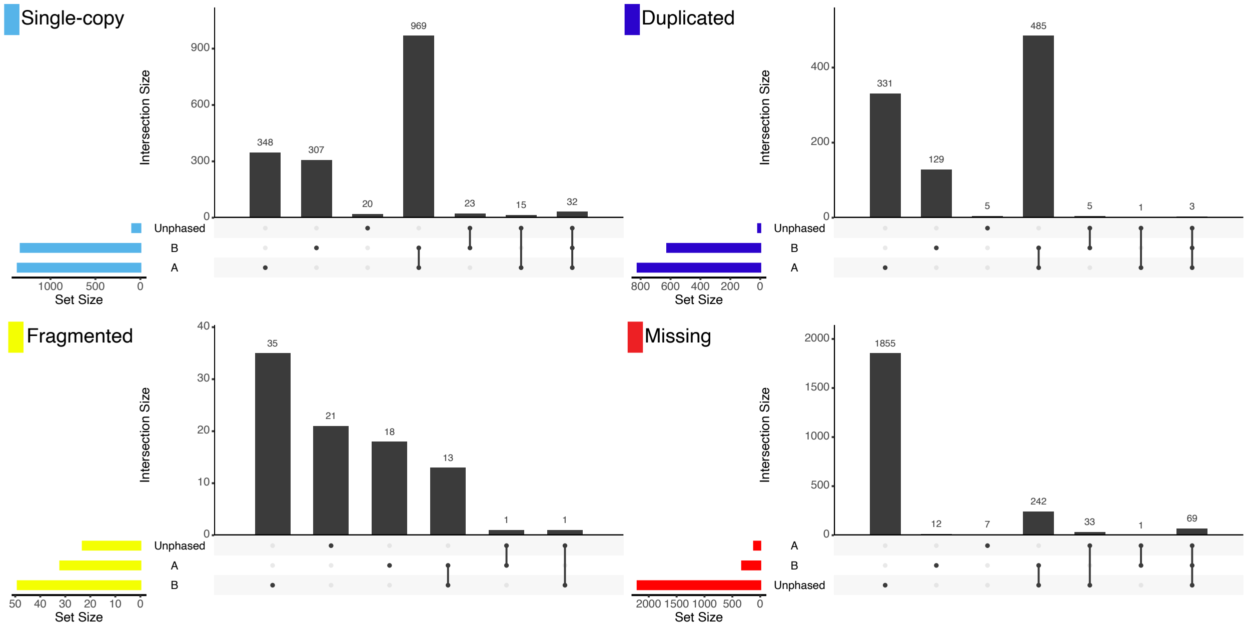


Fig. 4. BUSCO analysis of *M. domestica* cv. Gala with the eudicots odb10 dataset of 2,326 single-copy genes.

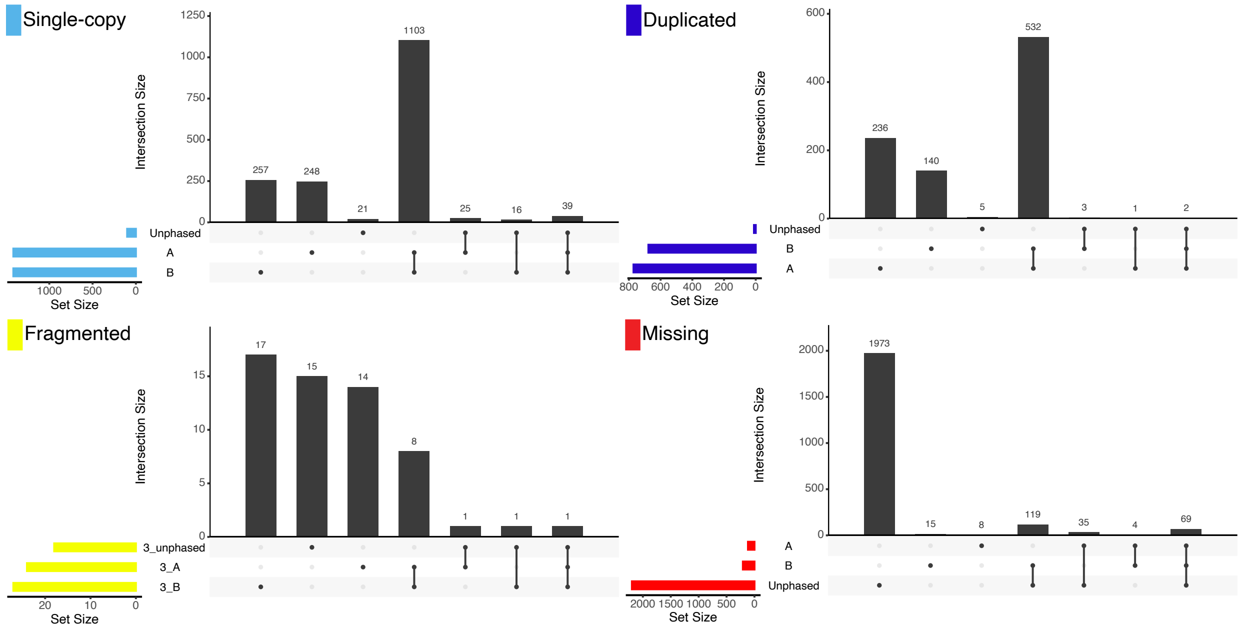


Fig. 5. BUSCO analysis of *M. sieversii* with the eudicots odb10 dataset of 2,326 single-copy genes.

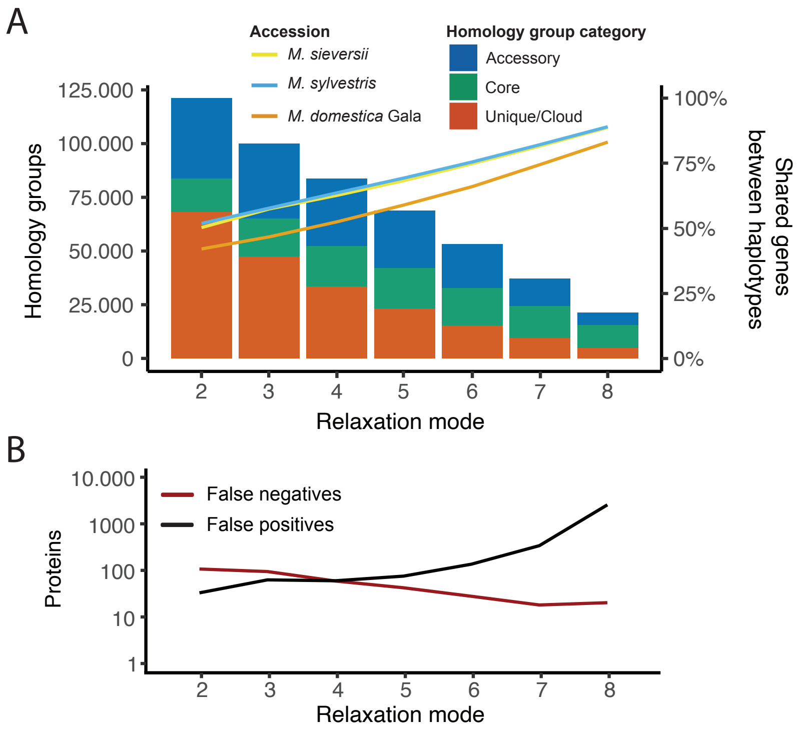


Fig. 6. (A) The effect of increasing relaxation modes (lowering clustering stringency) on the *Malus* pangenome composition, in terms of total number of homology groups (bar charts) and average percentage of genes shared between subgenomes (line graphs). (B) PanTools' BUSCO benchmark results of the seven homology grouping settings (relaxation modes).

### Supplementary Analysis 2. *Malus* gene content analyses

#### Characterizing intergenomic variation

Protein sequences of the *Malus* pangenome clustered into 53,261 homology groups, comprising 57.7% core, 33.1% accessory and 9.8% cloud groups. More than half of the cloud groups were singleton groups, each holding only a single protein without any similarity to any other protein in the pangenome. Individual genomes contain 54.5-60.9% core genes, 7.3-11.5% cloud genes, with the remainder being accessory (Fig. 7A). Gene regions of *M. sylvestris* were colored according to this classification (Fig. 7B). Within this genome the different gene categories appear to be randomly distributed, lacking a perceptible pattern or organization.

Overlap between genomes was measured as the union of two homology group sets and varied between 50.9-65.2%. *M. baccatta* shared least genes with any other genome, the three phased genomes showed most overlap in gene content. Interestingly, Gala shared less genes with the other *M. domestica* genome (GDDH13) than with the other two haplotype-resolved genomes, yet from GDDH13's viewpoint, Gala was the most similar genome in terms of gene content. The gene content of phased genomes was clustered in at least 5,000 additional homology groups compared to the unphased genomes, indicating significant variation that remains hidden in pseudo-haploid assemblies.

#### Characterizing intragenomic variation

We assessed the presence of genes in one or two subgenomes (Fig. 7C). Among the three phased apple genomes, *M. domestica* Gala had the lowest number of homology groups (57.2%) with mRNAs located on both subgenomes. Substantially more groups were identified in two subgenomes in the *M. sylvestris* (68.0%) and *M. sieversii* (67.4%) genomes. In Fig. 7D, gene regions of *M. sylvestris* are colored depending on whether the gene was found in the other haplotype. Overall, haplotype specific genes (black) are distributed randomly around the entire chromosome, but are also co-localized in specific regions.

We observed a large increase in distinct alleles as a direct result of resolving the haplotypes: phased genomes on average had 2.0 alleles, opposed to 1.3 in the unphased assemblies. Alleles were counted from CDS sequences in homology groups, where sequences were considered distinct alleles when having a SNP that can differentiate it. The majority of the phased genomes gene content is found in a mono-allelic state (39.6-43.6%). Still, high heterozygosity is implied by the large number of genes in a bi- (38.6-42.3%), tri- (8.5-8.8%) or tetra-allelic (5.7-6.2%) state.

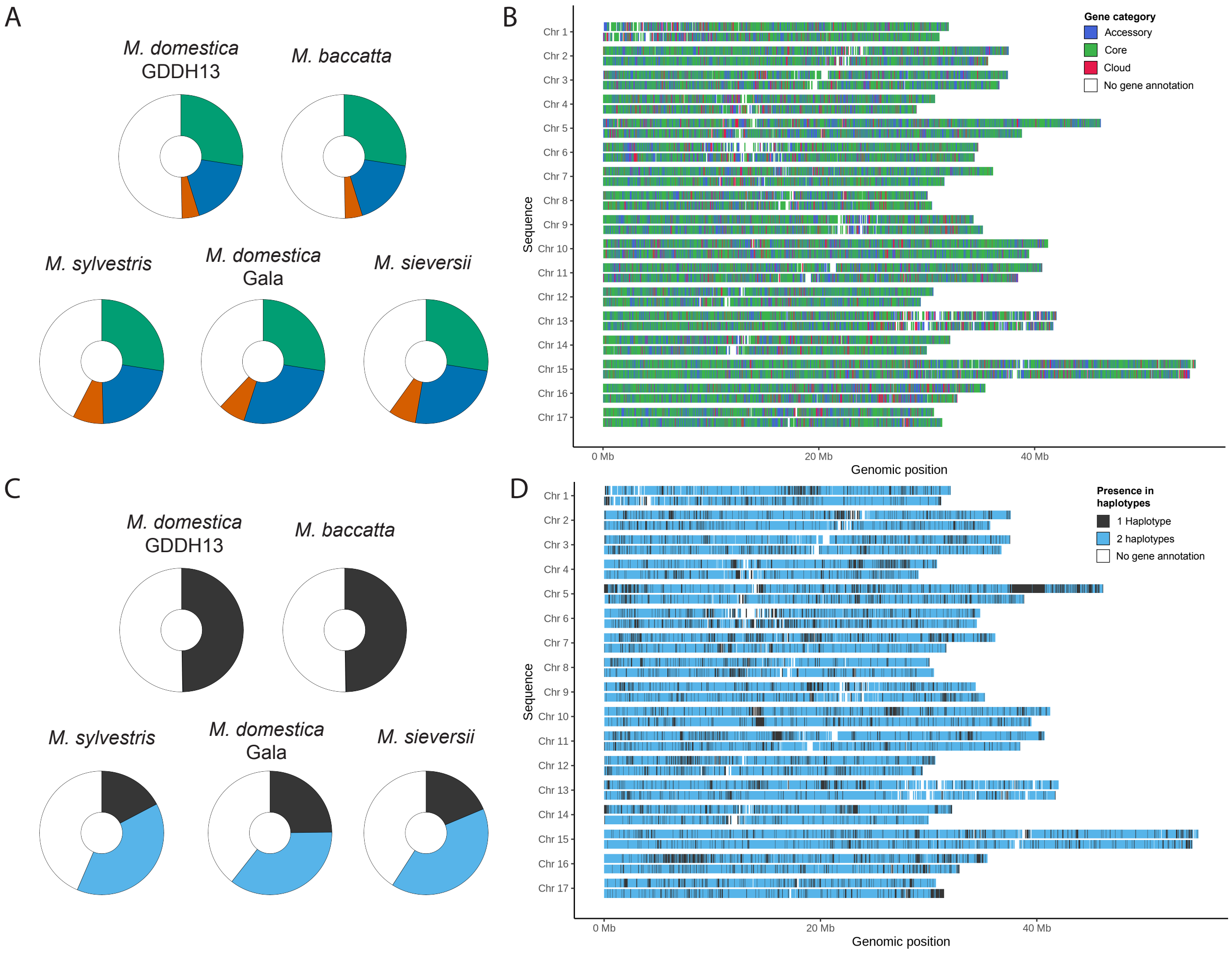


Fig. 7. Characterization of the *Malus* pangenome gene content. (A) Pie chart slices indicate the proportion of groups being core, accessory, or cloud. Each circle represents 68,751 homology groups of the pangenome. (B) Gene regions of *M. sylvestris* colored to represent shared homology with four other apple genomes. (C) Pie chart slices show the presence of genes in either one or two subgenomes. The circles represent an equal number of groups compared to plot A, although they have slightly a larger proportion of white due to genes located on unphased sequences. (D) *M. sylvestris* gene regions color-based by their presence in the two haplotypes of chromosomes.

### Supplementary Analysis 3. *Malus* evolutionary analyses

#### Assessing multiple phylogenetic methods

Phylogenetic relationships were established in the *Malus* pangenome at the genome, subgenome and sequence level. Inconsistent haplotype assignment in the genome assemblies results in conflicting signals when inferring phylogeny of subgenomes. Here we provide subgenome trees as a proof of principle to showcase the potential of the updated PanTools functionalities, despite the issues posed by the arbitrary haplotype assignment. The four phylogenetic methods that we review are explained in the main results (*'Establishing the evolutionary history in the pangenome'*) and methods section (*'Phylogenetic analyses'*). In the following section we discuss their application to the *Malus* pangenome.

No genome core phylogeny was inferred as only 46 single-copy groups were identified, which would result in poor resolution and low statistical support. A subgenome-level core phylogeny was inferred on 2,902 single-copy groups and 33,980 SNPs. Subgenomes of the three wild apples cluster together as expected. The domesticated apple subgenomes also cluster together; however, the Gala subgenomes were not adjacent to each other as GDDH13 was positioned between them.

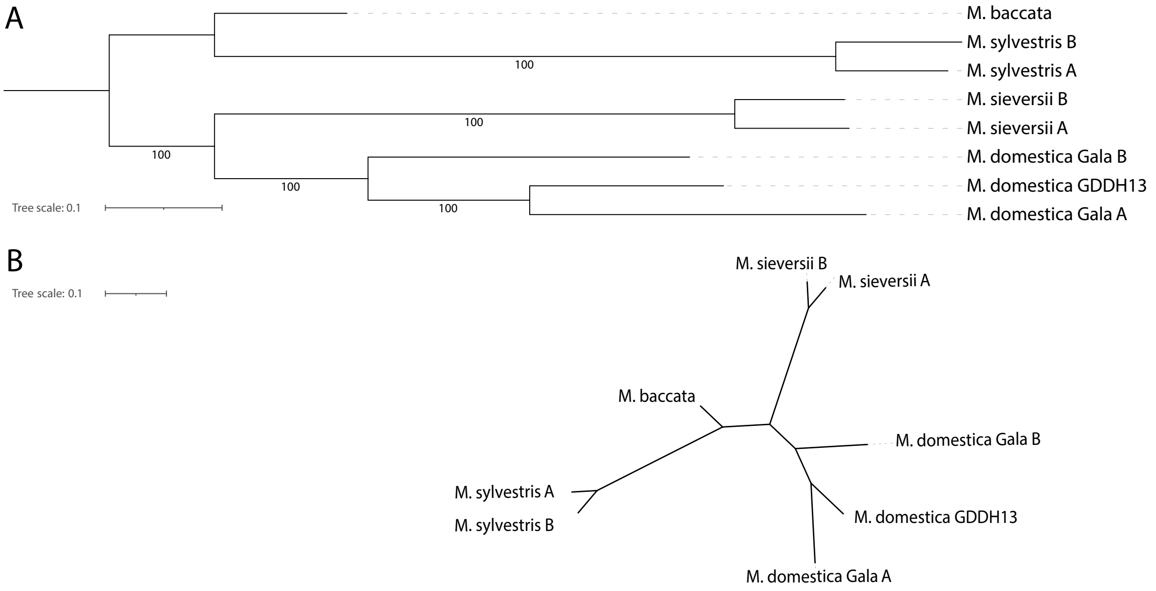


Fig. 8. Core SNP phylogeny at subgenome-level of five *Malus* genomes. The topology is represented in rectangular form (A) and split graph (B). The root was placed at the midpoint node of the rectangular tree.

The genome $k$-mer tree (Fig. 9A) showed the divergence of *M. baccatta* in the first split and *M. domestica* nested within the wild apples. This topology is in agreement with earlier high resolution phylogeny estimations [2], [3], [4], [5]. The subgenome $k$-mer tree (Fig. 9B) also displays the separation of the Gala A and B subgenomes, similar to the topology of the subgenome core phylogeny.

The sequence-level $k$-mer distance tree (Fig. 9C) shows 17 clades corresponding to the number of chromosomes of the *Malus* genus. No conflicts were observed in the assigned chromosome numbers, supporting a correct topology. Except for Chr 2, every chromosomal clade shares an early branching point to another chromosomal clade, suggesting common ancestry of chromosomes. Based on the earlier synteny analysis, we could confirm that the co-placed chromosomes were duplicated earlier through the recent polyploidy event. Within the chromosomal clades, all *M. sylvestris* sequences and most *M. sieversii* sequences were positioned in pairs, thus clustering the two homologous chromosomes. This adjacent placement occurred for only two Gala chromosome pairs; in most other clades, the Gala A subgenome was co-located with GDDH13, with the B subgenome randomly placed in the clade.

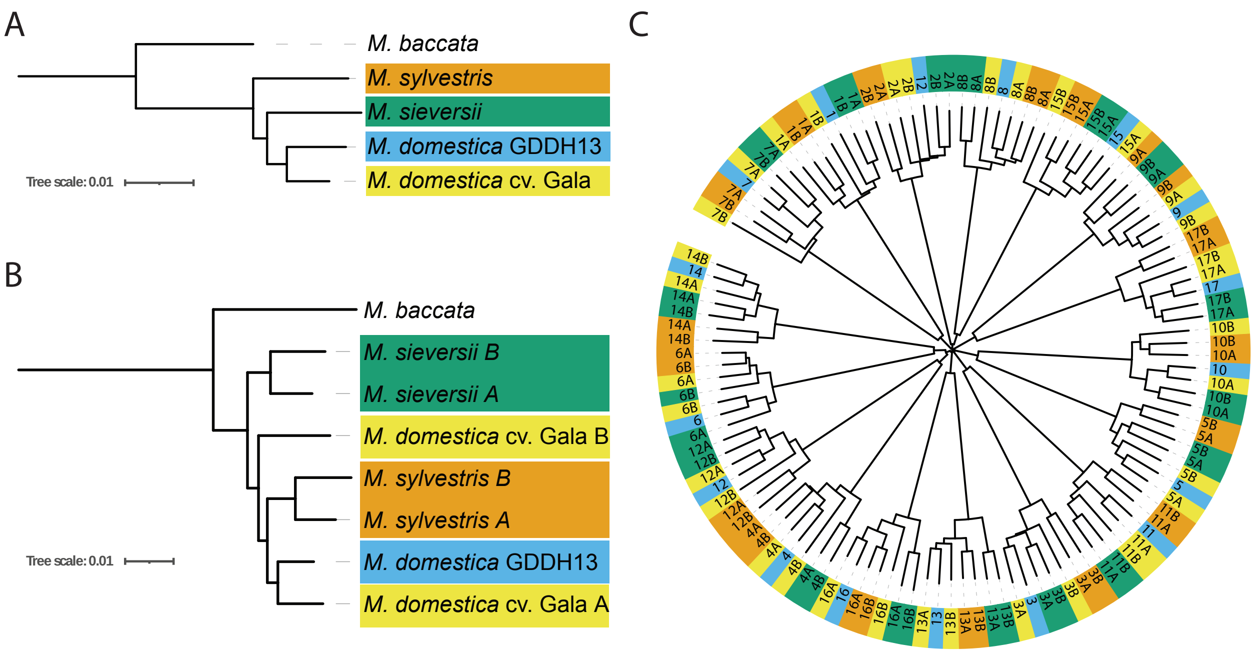


Fig. 9. Genome (A), subgenome (B) and sequence (C) $k$-mer distance phylogenetic trees from the *Malus* pangenome. The coloring of the (sub)genome tree serves as legend for the sequence tree.

#### Synonymous mutation analysis

To gain insights into the WGD history of the *Malus* genus we calculated synonymous ($D_s$; silent mutation) substitution rates of homologous genes, between but also within genomes. The distribution of the mutation rates is shown in Fig. 10, where intra-genomic comparisons are presented on the diagonal. The youngest peak ($D_s$ 0.01-0.02), derived from the low mutation rates of orthologous sequences, represents the species divergence event. Considering the divergence of *M. sieversii* and *M. sylvestris* $\sim$1.8 million years ago [5] and an obtained $D_s$ 0.0138 from the first peak, the estimated substitution rate is $3.83 \times 10^{-9}$ sites per year. This rate is nearly identical to the original estimate based only on a small set of single-copy genes [4].

Apart from speciation peaks observed in the intergenomic analysis, we also observed peaks with similar mutation rates in the intragenomic comparison plots. In Gala this peak was noticeably higher than in the other two wild *Malus* genomes, suggesting a higher level of divergence between subgenomes. Given the hybrid ancestry of *M. domestica* Gala, high heterozygosity indicates preserved genetic information from its progenitors.

The plots exhibit two additional peaks. The second peak is most prominent in every plot, around $D_s$ 0.16-0.19. This peak represents distances between paralogous sequences derived from the Pyrinae tetraploidization event [6]. Although the third peak centered around $D_s$ 1.2-1.9 provides a weaker signal, the distribution is nearly identical to an earlier study that hypothesized this to signal the ancient eudicot paleohexaploidy ($\gamma$) event [7].

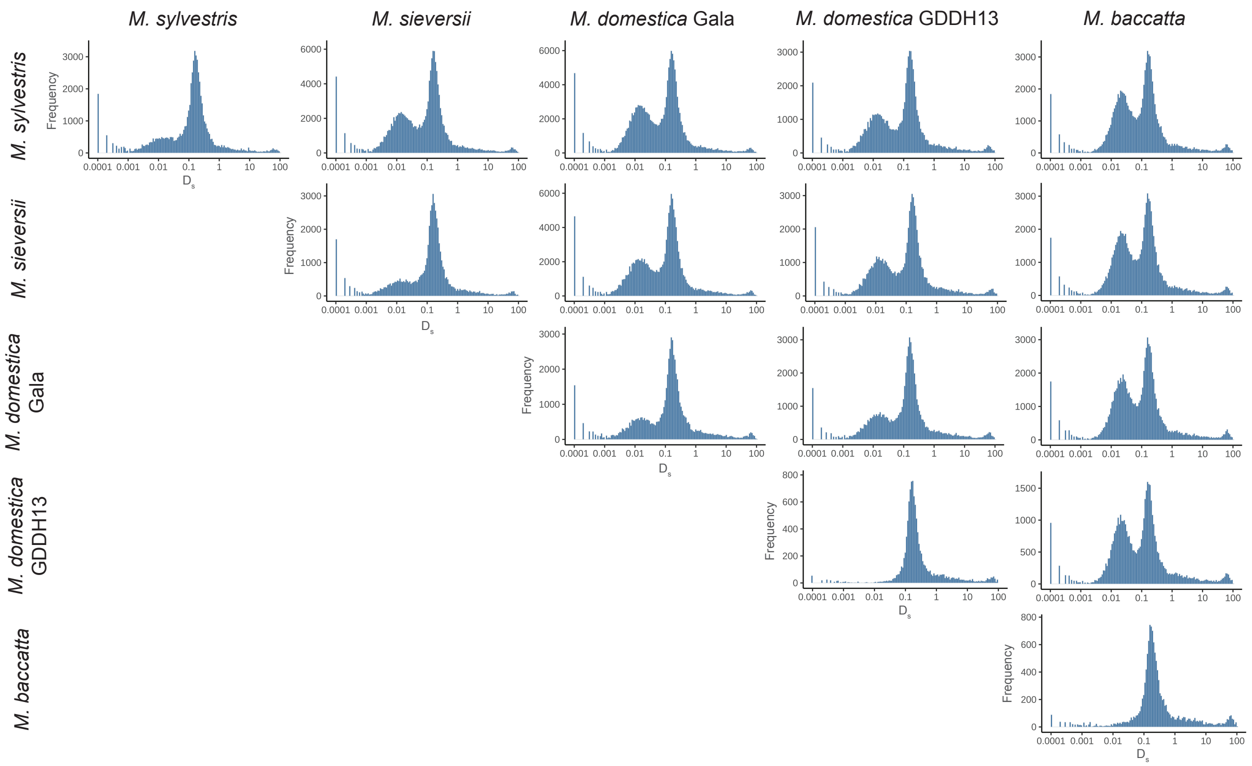


Fig. 10. Synonymous substitution rate ($D_s$) distributions of homologous sequences between *Malus* genomes. Values on the x-axis are on a logarithmic (10) scale.

### Supplementary Analysis 4. Visualizing genomic organization and variation of *S. tuberosum* genomes

#### Gene retention analysis

Using the PanTools 'gene_retention' function we calculated syntenic gene retention against *S. tuberosum* DM 1-3 516 R44 Chr 2 and 11. The retention pattern against DM Chr 11 (Fig. 11A) was representative for the majority of visualizations, showing strong conservation in distal parts of the chromosome but high fractionation in the pericentromeric regions. The four chromosome haplotypes of C88 and Otava generally show the same level of retention, except for the central region of the chromosome where each of Otava's haplotypes show a distinct retention pattern, and C88 shows two disparate haplotype pairs. Moreover, the plot illustrates that the four Atlantic Chr 11 haplotypes can be very disparate, especially surrounding the centromeric region, where two haplotypes were completely missing.

Retention patterns of the first half of DM Chr 2 (Fig. 11B) were similar between genomes, in contrast to the extremely fluctuating second half. Half of the haplotypes were disparate in the first 8 Mb of the chromosome, showing two highly retained sequences. Only the C88 haplotypes display conserved collinearity across the entire chromosome. The Atlantic and Otava genome show multiple regions with complete loss of synteny, along with translocations. These translocated regions were recognized by the loss of retention in Chr 11 and sudden increase of Chr 10 in Atlantic (red line) and Chr 1 in Otava (blue lines). All four Otava haplotypes display a translocation whereas in Atlantic this occurred for just a single haplotype.

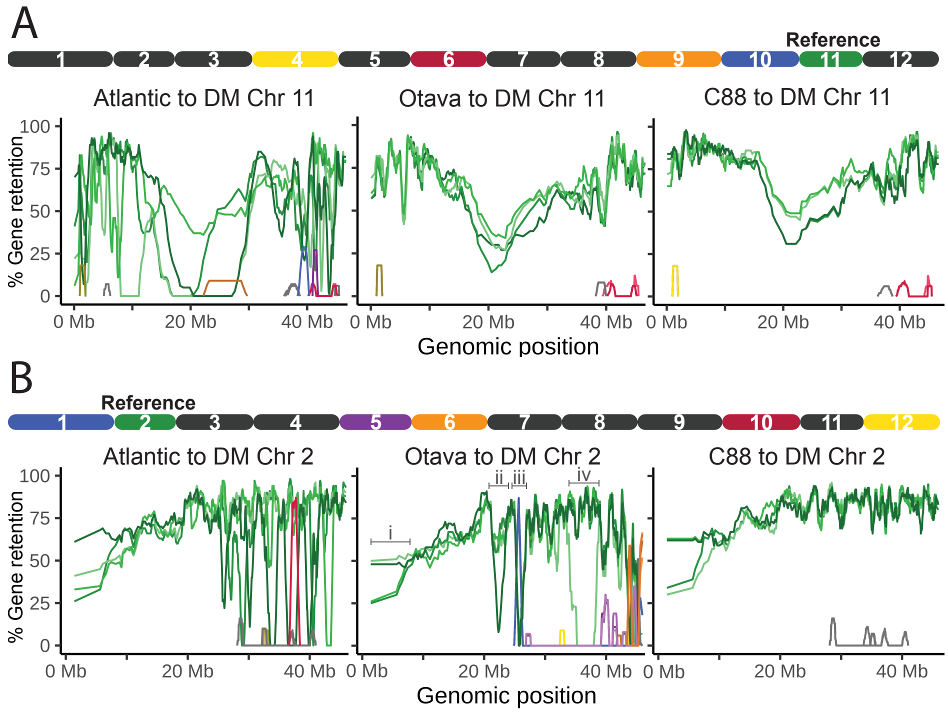


Fig. 11. Syntenic gene retention of *S. tuberuosum* DM 1-3 516 R44 Chr 11 (A) and Chr 2 (B). The different shades of colors in the line graph indicate different haplotypes of a chromosome. A schematic representation of the DM genome above the retention line graphs illustrates which chromosomes show retention to the reference query, indicated by their color.

#### Chromosome visualizations

In this analysis, we revisit the two chromosomes from the previous gene retention analyses, but now look at synteny relationships and gene absence/presence variation as a way to explore the underlying relationship of intriguing retention patterns. We demonstrate two examples with chromosomes from Otava and C88. Because the earlier retention plots were plotted against DM, the coordinates cannot be directly adopted.

C88's Chr 11 sequences were visualized in Fig 12A to examine the variation in the pericentromeric regions. The majority of genes in the regions flanking the centromere (20-30Mb) colored blue, illustrating they occur in a 2:2 gene distribution. Furthermore, two large synteny breakpoints together with an inversion were visible between A-B and C-D. Both findings support the earlier observation where two haplotypes display lower gene retention in the pericentromeric regions. Just to the right, in a subsequent region (30-35Mb) of haplotype C, we observed that most of the genes were haplotype specific. This genomic region colored predominantly orange in the other three haplotypes, indicating a 1:3 gene distribution.

Otava's Chr 2 haplotypes are visualized in Fig 12B. The visualization shows many small-scale (inverted) translocations, especially between the A and B haplotypes. These variants were specific to A, as none of the translocation appeared in C or D. We discuss four regions that showed a strong loss of synteny in the previous retention analysis (Fig 11B. Consistent with the retention plot, the first 8 Mb (i) of the four haplotypes were highly dissimilar. The leftmost dip (ii) in retention specific to haplotype 2D appeared as synteny break point to 2C (around $\sim$25Mb). The observed loss caused by the translocation (iii) was not visible because the event was shared among all haplotypes. The final loss of retention in haplotype B (iv, region 36-40Mb) was supported by a clearly visible synteny breakpoint, further revealing most lost genes still have three copies located on the other homeologous chromosomes.

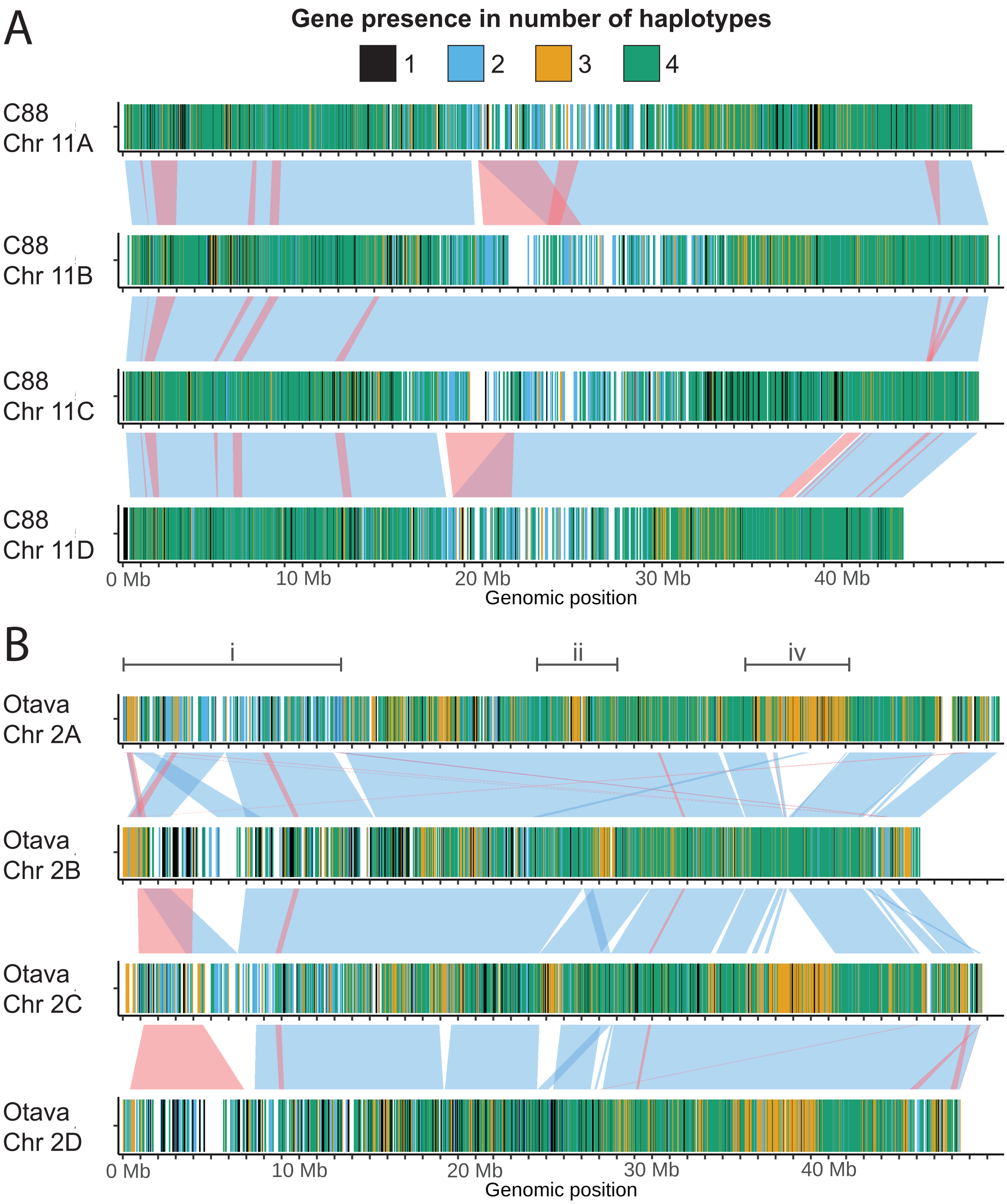


Fig. 12. Genetic and structural variation within (A) *S. tuberosum* C88 Chr 11 and (B) Otava Chr 2. Gene regions are colored by presence in number haplotypes. Syntenic blocks are drawn between two sequences, inverted blocks are red.

#### Whole-chromosome alignments

We performed 12 whole-chromosome alignments using the minimap2 [8] enhanced PanTools functionality (explained in Methods section I). A general observation in many dot plots was that the Atlantic and Castle Russet haplotypes appeared as a curve rather than a diagonal as a consequence of short chromosome sequences. Most pairwise alignments against these two genomes were highly fragmented and very noisy. Still, the dot plots revealed rearrangements in all four haplotype-resolved assemblies. The identified structural rearrangements were not found as quadruplets, but instead were specific for one to three haplotypes.

We showcase the alignments of *S. tuberosum* Chr 12 as these most clearly displayed large structural variations (Fig 13A). These full-length chromosome alignments showed an overall high collinearity among themselves. Contrarily, the pericentromeric regions appear as noisy blocks, conveying to be highly divergent and repetitive. This was especially evident for the Otava haplotypes (marked with red block), where substantial intra-genomic variation located in centromeric regions completely fragmented the alignments. Apart from fragmentation, Chr 12 alignments were characterized by a large ∼20Mb inversion. Relative to the DM genome, all four Otava haplotypes display this large inversion, whereas C88 only shows two inversions. When zooming into the inverted regions of C88 (Fig 13B), we observed high fragmentation in the centromere and several additional smaller rearrangements.

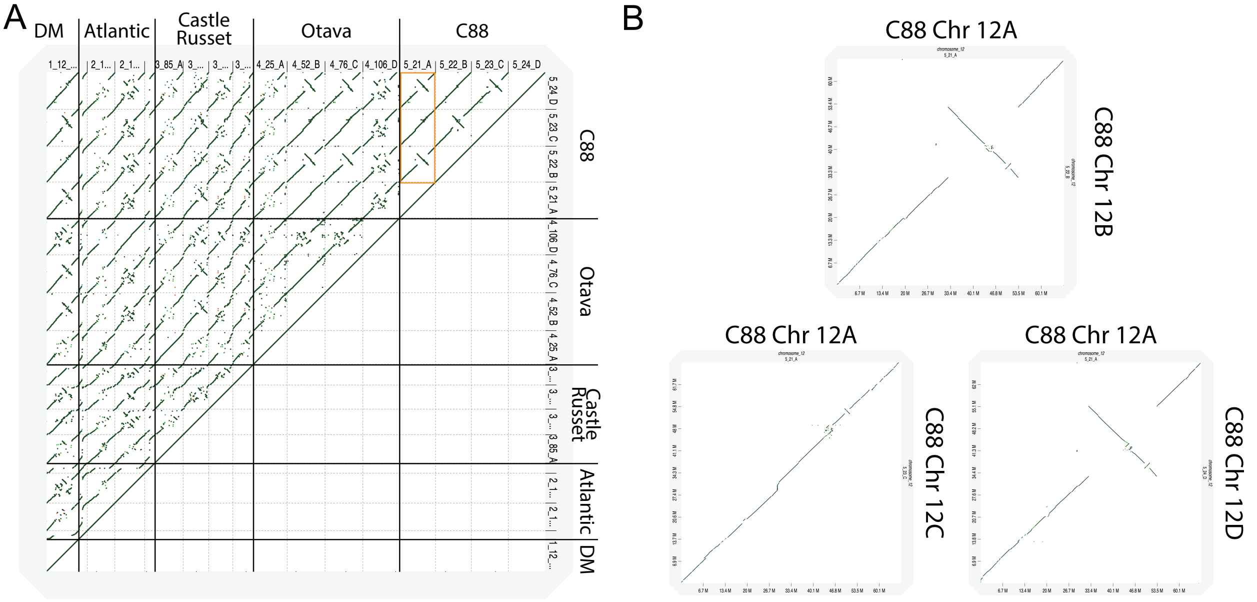


Fig. 13. Dot plot visualization of *S. tuberosum* Chr 12 alignments. (B) Higher resolution visualization of the intra-genomic alignments of C88 Chr 12 (marked in plot A). Small matches in the alignment were removed.

#### Supplementary Analysis 5. *Malus*: Integrative chromosome overview

##### *M. domestica* Gala Chr 3 visualization

Here we re-explore the PanTools 'sequence_visualization' function. In previous examples, we demonstrated the ability to combine intra- and intergenomic gene presence variation with synteny. Now, we incorporate additional features to demonstrate the tool's ability to generate an integrative overview by combining different (pan)genomic attributes.

We showcase *M. domestica* Gala Chr 3 (Fig. 14) as one of many chromosomes with a substantially larger A haplotype compared to B. The plot clearly captures the haplotype length differences, showing that haplotype 3A is nearly 5Mb longer thanhaplotype 3B. Both haplotypes exhibit a large proportion of genes which seem to be present in only a single haplotype (annotation bar 4). Using the homology groups, we assessed how many genes were shared between the two haplotypes. Around 27% of the 3A genes did not have any homolog in 3B, whereas roughly 17% of the 3B gene content was not present in 3A. Regardless of high gene absence/presence variation, the genomic collinearity of the haplotypes was highly conserved (bar 5) and was only disrupted by larger blocks of co-localized haplotype-specific genes. In addition to these synteny breakpoints, a single haplotype-specific small-scale inversion (red blocks) was found, but this did not break the larger syntenic block spanning half of the chromosome.

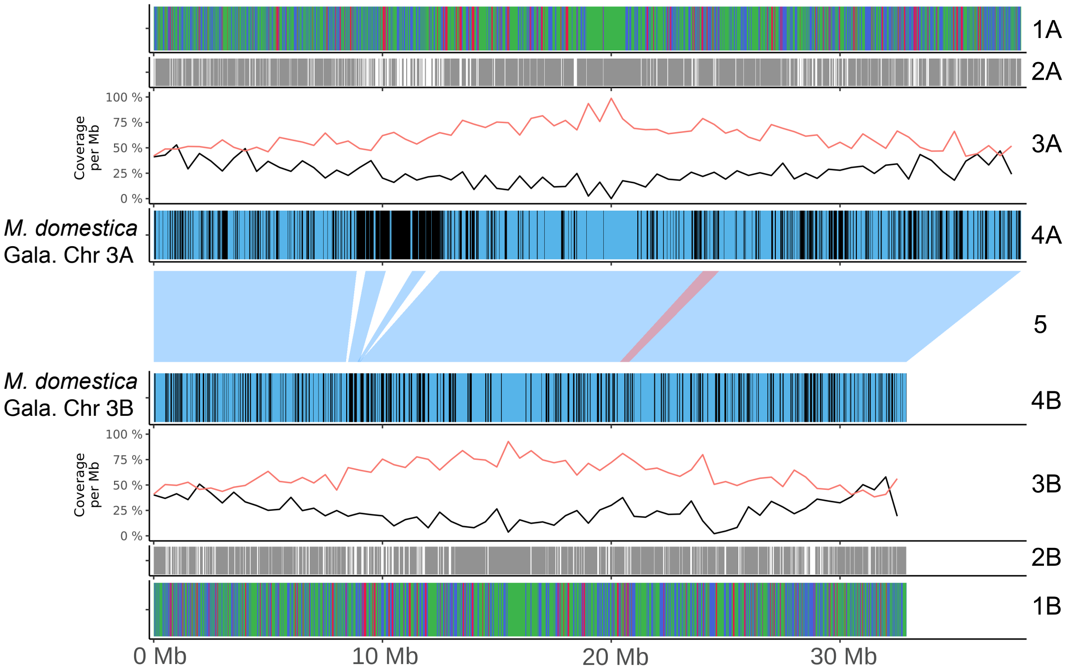


Fig. 14. *M. domestica* Gala Chr 3 represented by various visualizations of (pan)genomic features. Annotation bars from top to middle (and bottom to middle): intergenomic gene presence: core (green), accessory (blue) or cloud (red); genes with another homolog on another Gala chromosome (grey); repeat (red) and gene (black) coverage of 500 Kb windows; intragenomic gene presence: single (black) or both (blue) haplotypes; syntenic blocks between two haplotypes, with inverted blocks in red.

The observable pattern of gene and repeat coverage (bar 3) follows a typical pattern in plants; high gene and low repeat coverage in the subtelomeric regions, and *vice versa* in the pericentromeric regions. The strongest anti-correlation of these two genomic features is found near the center of the chromosome (19-20 Mb), indicating the centromere locations. Although centromeres generally consist of tandemly repeated noncoding sequences, the gene coverage indicates several protein-coding genes are located within these regions.

The outer bars (bar 1) visualize core, accessory and cloud genes. Where in potato this demonstrated clear a localization of accessory and cloud genes in the middle of the chromosome, here genes of different categories appear to be randomly distributed. Bar 2 illustrates if a gene occurs in any other Gala chromosome.

By combining the information of bar 1A, 2A, and 4A for haplotype A genomic region 7.5-12.5MB, we can examine that most genes in this region are specific to this genomic segment within Gala. This observation was made from the patterns in bar 3A (mostly filled black) and 4A (mostly unfilled). Since this region appears as a mosaic of different gene categories in bar 1A, the majority of genes are not necessarily specific to the Gala genome.
