## Supplementary material for "Exploring intra- and intergenomic variation in haplotype-resolved pangenomes": README.html

The following PanTools commands were used to perform the analyses and create all figures of the manuscript. The commands were not necessarily executed in this order. When the same command was applicable to both potato and apple, only one command is shown, with the database refered as **pangenome\_DB**.

### Instructions for PanTools v4.3.0

The analysis was originally performed using PanTools on the **phased\_pangenomics** development branch, which was merged with the **pantools\_v4** branch at commit d5eca936 (https://git.wur.nl/bioinformatics/pantools/-/commits/) and released in PanTools v4.3.0 (https://git.wur.nl/bioinformatics/pantools/-/releases/v4.3.0). The original command-line arguments were slightly revised in the release versions. Additionally, the `genome_alignment` functionality has been removed from the current release. The following set of instructions allows to perform the analyses with PanTools v4.3.0 (and later):

```
pantools build_pangenome pangenome_DB 5_genomes.txt
pantools add_annotations pangenome_DB 5_annotations.txt
```

Repeats were only identifed in the three haplotype-resolved apple genomes with EDTA https://github.com/oushujun/EDTA.

```
perl EDTA.pl --genome Msylvestris_diploid_v2.chr.fa --cds Msylvestris.pep.fas --overwrite 1 --sensitive 1 --anno 1 --evaluate 1 --threads 12
perl EDTA.pl --genome Gala_diploid_v2.chr.fa.gz --cds Gala.pep.fas --overwrite 1 --sensitive 1 --anno 1 --evaluate 1 --threads 12
perl EDTA.pl --genome Msieversii_diploid_v2.chr.fa.gz --cds Msieversii.pep.fas --overwrite 1 --sensitive 1 --anno 1 --evaluate 1 --threads 12

pantools add_repeats apple_DB 3_apple_repeats.txt
pantools repeat_overview apple_DB
```

Different BUSCO sets were used to assess completeness

```
pantools busco_protein apple_DB --odb10=eudicots_odb10 -t=12 --phasing 
pantools optimal_grouping apple_DB apple_DB/busco/eudicots_odb10/protein_phased -t=24 --relaxation=2,3,4,5,6,7,8 --phasing

pantools busco_protein potato_DB --odb10=solanales_odb10 -t=12 --phasing 
pantools optimal_grouping potato_DB potato_DB/busco/solanales_odb10/protein_phased/ -t=12 --relaxation=2,3,4,5,6,7,8 --phasing

pantools grouping_overview pangenome_DB
pantools change_grouping apple_DB -v=4 # version is not the relaxation but identifier of clustering actually the version. It is relaxation  
pantools change_grouping potato_DB -v=5
```

##### Adding phasing information to the pangenome

The chromosome numbering was based on a the clustering in the 'kmer\_classification' distance tree to DM1-3 (potato) and GDDH13 (apple), haplotype letters were assigned randomly. The phasing identifier files that were used (**potato\_identifiers.txt** & **potato\_identifiers.txt**) were added to this README folder.

```
pantools add_phasing pangenome_DB phasing_identifier.txt
```

##### Phylogenies

```
pantools core_phylogeny pangenome_DB --sequence --phasing
pantools kmer_classification pangenome_DB --sequence --phasing
Rscript pangenome_DB/gene_classification/gene_distance_tree.R
```

```
pantools metrics pangenome_DB --sequence
pantools msa -t=24 pangenome_DB
pantools calculate_dn_ds pangenome_DB -t=24 --homologs
```

In the potato pangenome, 'gene\_classification' was run on all chromosome sequences via the
min\_100\_genes.txt rule file, which only included sequences with at least 100 gene annotations. In apple, every sequence of the fragmented *M. baccata* assembly were included in adition to all chromosomes.

```
pantools gene_classification potato_DB --sequence --phasing --selection-file=min_100_genes.txt
pantools gene_classification apple_DB --sequence --phasing --selection-file=min_100_genes_all_baccata.txt
```

##### Synteny analysis

```
pantools calculate_synteny pangenome_DB --selection-file=min_100_genes.txt -t=24 --sequence --run
pantools add_synteny pangenome_DB pangenome_DB/synteny/mcscanx.collinearity
pantools synteny_statistics pangenome_DB -t=12
pantools gene_retention pangenome_DB --phasing --selection-file=min_100_genes.txt
```

For the potato sequence visualizations, a rule file was included to only show annotation bars where regions are colored by gene presence in number of haplotypes.

```
pantools sequence_visualization apple_DB --selection-file=min_100_genes.txt
pantools sequence_visualization potato_DB --selection-file=min_100_genes.txt --rules=haplo_presence.txt
```

#### Analysis on *StCDF1* in potato pangenome

The included node is the *StCDF1* homology group. The regions file input for the MSA was created from the 'group\_info' output.

```
pantools group_info potato_DB --node=232273529 
pantools msa potato_DB --method=regions --regions-file=StCDF1_regions
```

### Original command-line arguments

The following commands were used for the original analyses conducted with the **phased\_pangenomics** branch prior to commit d5eca936:

```
pantools build_pangenome pangenome_DB -gf 5_genomes.txt
pantools add_annotations pangenome_DB -af 5_annotations.txt
```

```
pantools add_repeats apple_DB -if 3_apple_repeats.txt
pantools repeat_overview apple_DB
```

```
pantools busco_protein apple_DB --busco10 eudicots_odb10 -t 12 --phasing 
pantools optimal_grouping apple_DB apple_DB/busco/eudicots_odb10/protein_phased -t 24 -relaxation 2,3,4,5,6,7,8 --phasing

pantools busco_protein potato_DB --busco10 solanales_odb10 -t 12 --phasing 
pantools optimal_grouping potato_DB potato_DB/busco/solanales_odb10/protein_phased/ -t 12 --relaxation 2,3,4,5,6,7,8 --phasing

pantools grouping_overview pangenome_DB
pantools change_grouping apple_DB -v 4 # version is not the relaxation but identifier of clustering actually the version. It is relaxation  
pantools change_grouping potato_DB -v 5
```

```
pantools add_phasing pangenome_DB -if phasing_identifier.txt
```

```
pantools core_phylogeny pangenome_DB --sequence --phasing
pantools kmer_classification pangenome_DB --sequence --phasing
Rscript pangenome_DB/gene_classification/gene_distance_tree.R
```

```
pantools metrics pangenome_DB --sequence
pantools calculate_dn_ds pangenome_DB -t 24 --msa --homologs
pantools genome_alignment pangenome_DB --sequence -tn 24
```

```
pantools gene_classification -dp potato_DB --sequence --phasing -sf min_100_genes.txt
pantools gene_classification -dp apple_DB --sequence --phasing -sf min_100_genes_all_baccata.txt
```

```
pantools calculate_synteny pangenome_DB -sf min_100_genes.txt -tn 24 --sequence --run
pantools add_synteny pangenome_DB -if pangenome_DB/synteny/mcscanx.collinearity
pantools synteny_statistics pangenome_DB -t 12
pantools gene_retention pangenome_DB --phasing -sf min_100_genes.txt
```

```
pantools sequence_visualization apple_DB -sf min_100_genes.txt
pantools sequence_visualization potato_DB -sf min_100_genes.txt --rules haplo_presence.txt
```

```
pantools group_info potato_DB --node 232273529 
pantools msa potato_DB --method regions -R StCDF1_regions
```
