## Supplementary material for "Exploring intra- and intergenomic variation in haplotype-resolved pangenomes": README.pdf

### Exploring intra- and inter-genomic variation in haplotype-resolved pangenomes

#### PanTools ‘sequence_visualization’ output

This supplementary file holds two folders containing plots generated by PanTools ‘sequence_visualization’ function for all haplotype-resolved genome assemblies. Apple visualizations were generated by including the following rule settings: ‘gene_classification, ‘haplotype_presence’, ‘other_chromosomes’, ‘repeat_coverage’, and ‘gene_coverage’. Potato plots were created with only ‘haplotype_presence’ bars. In both apple and potato, synteny relationships are drawn between haplotypes. These are included regardless of number of rules set. Figure 1 displays legends for the different type of annotation bars, which are universal and applicable to both apple and potato visualizations.

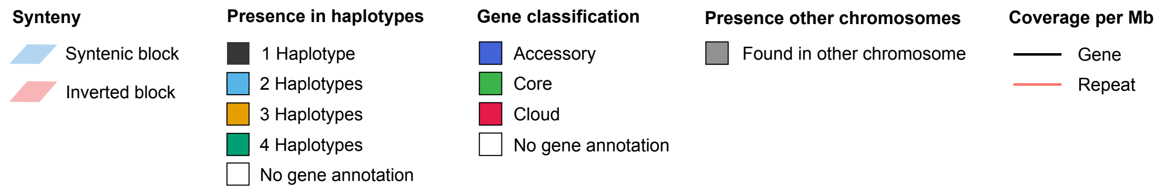


Fig. 1. Legends for different annotation bars
