## Supplementary figures and images for "Exploring intra- and intergenomic variation in haplotype-resolved pangenomes"

### Atlantic_chr_1.png

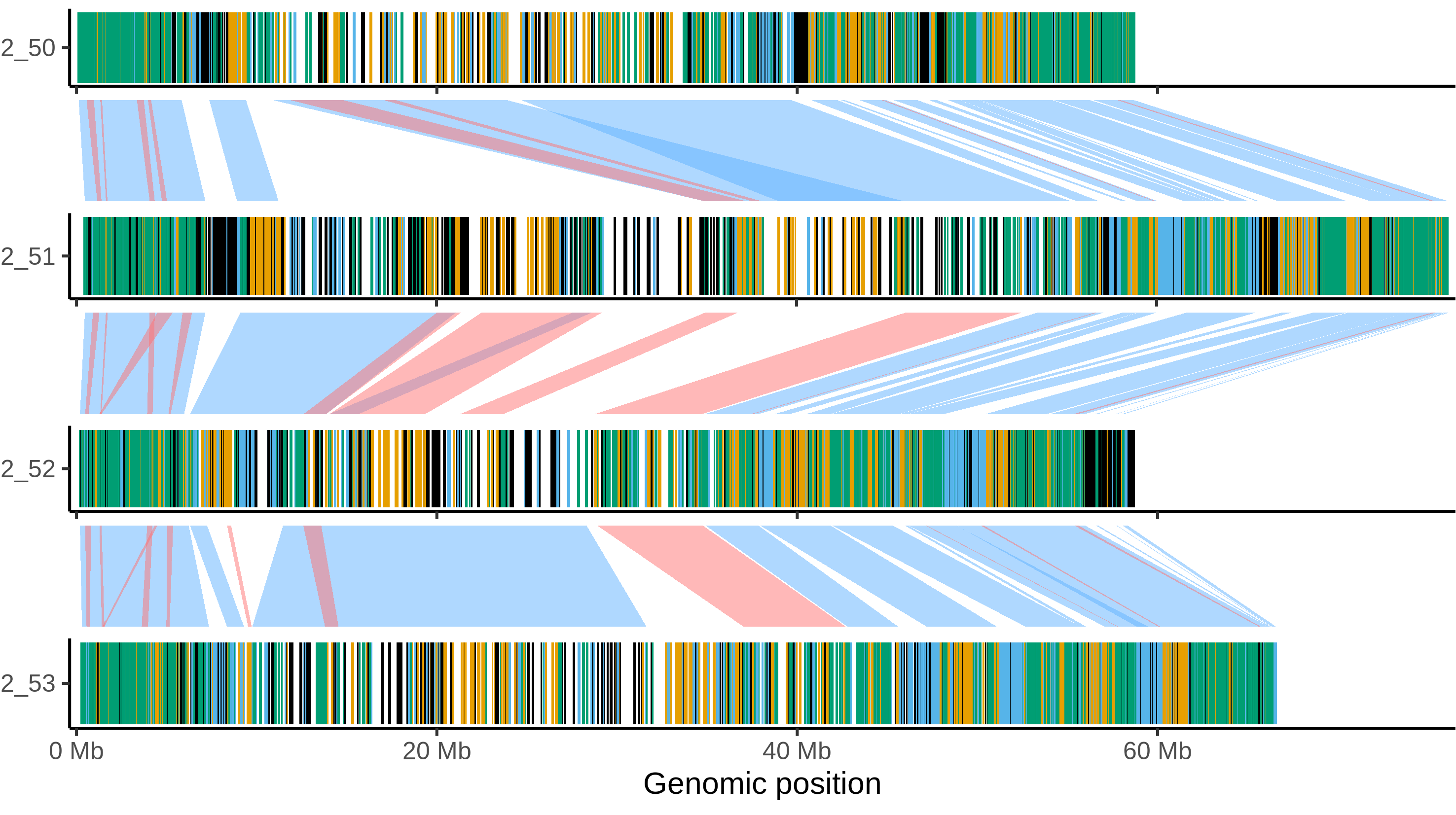

### Atlantic_chr_2.png

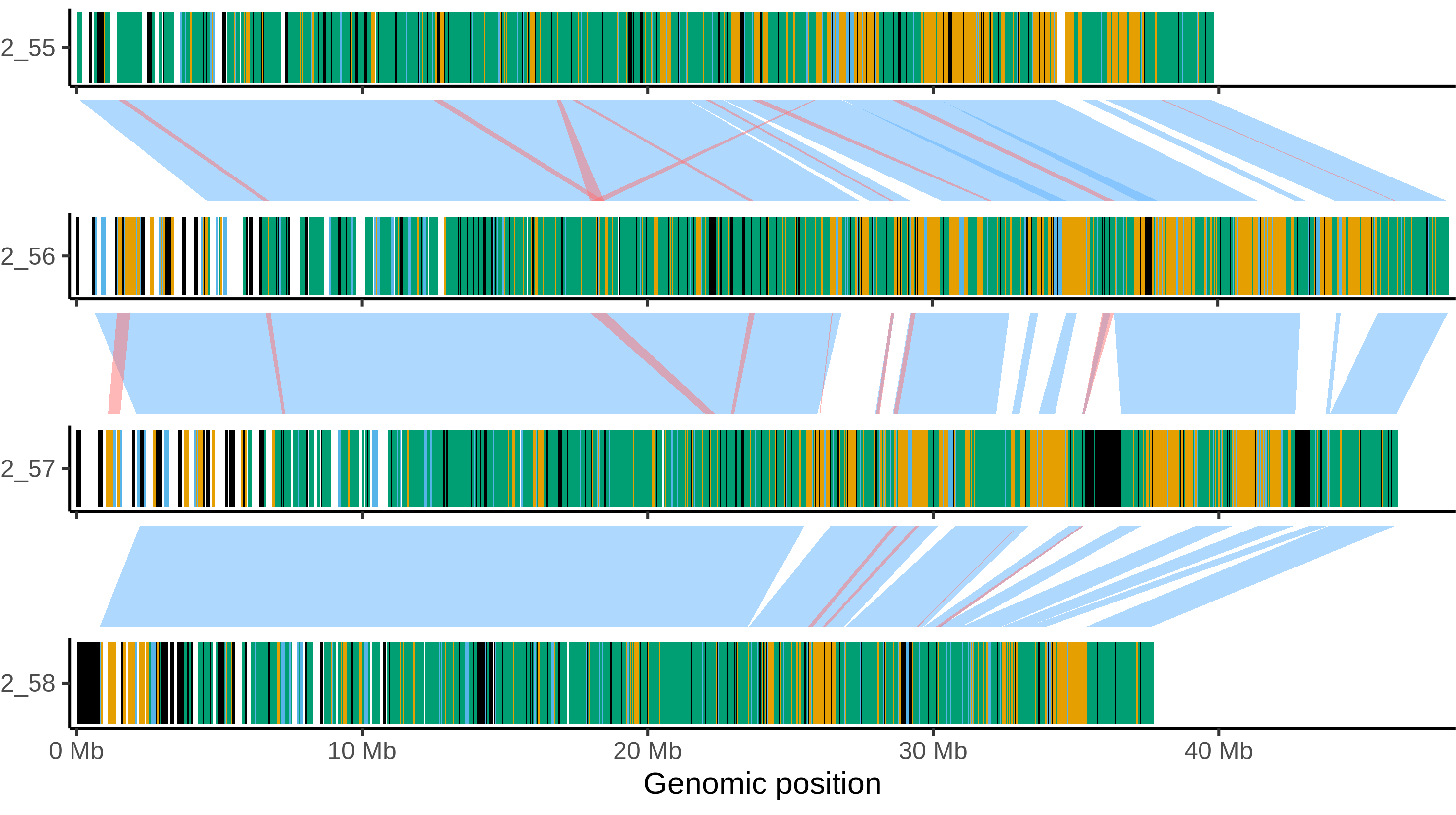

### Atlantic_chr_3.png

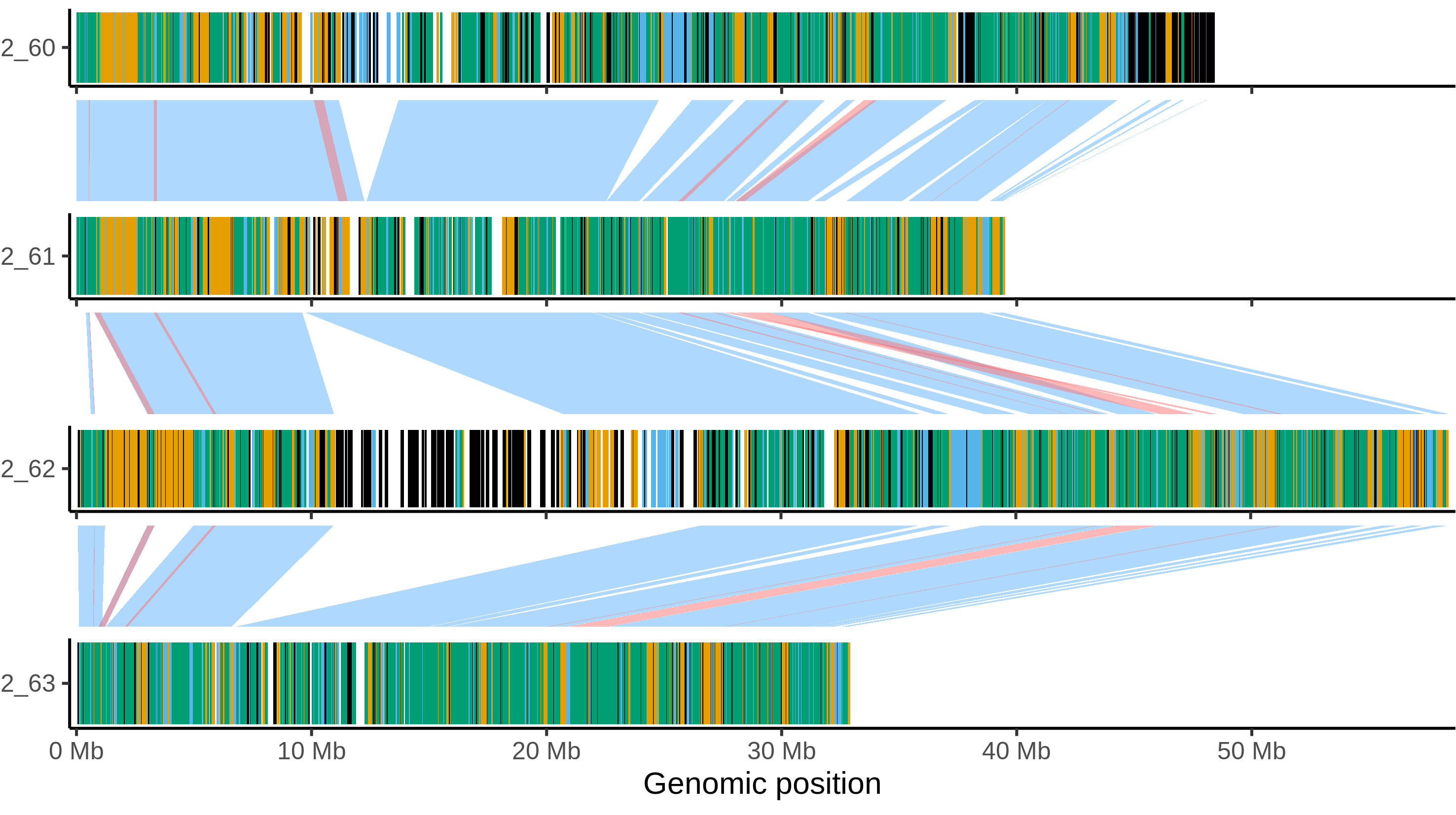

### Atlantic_chr_4.png

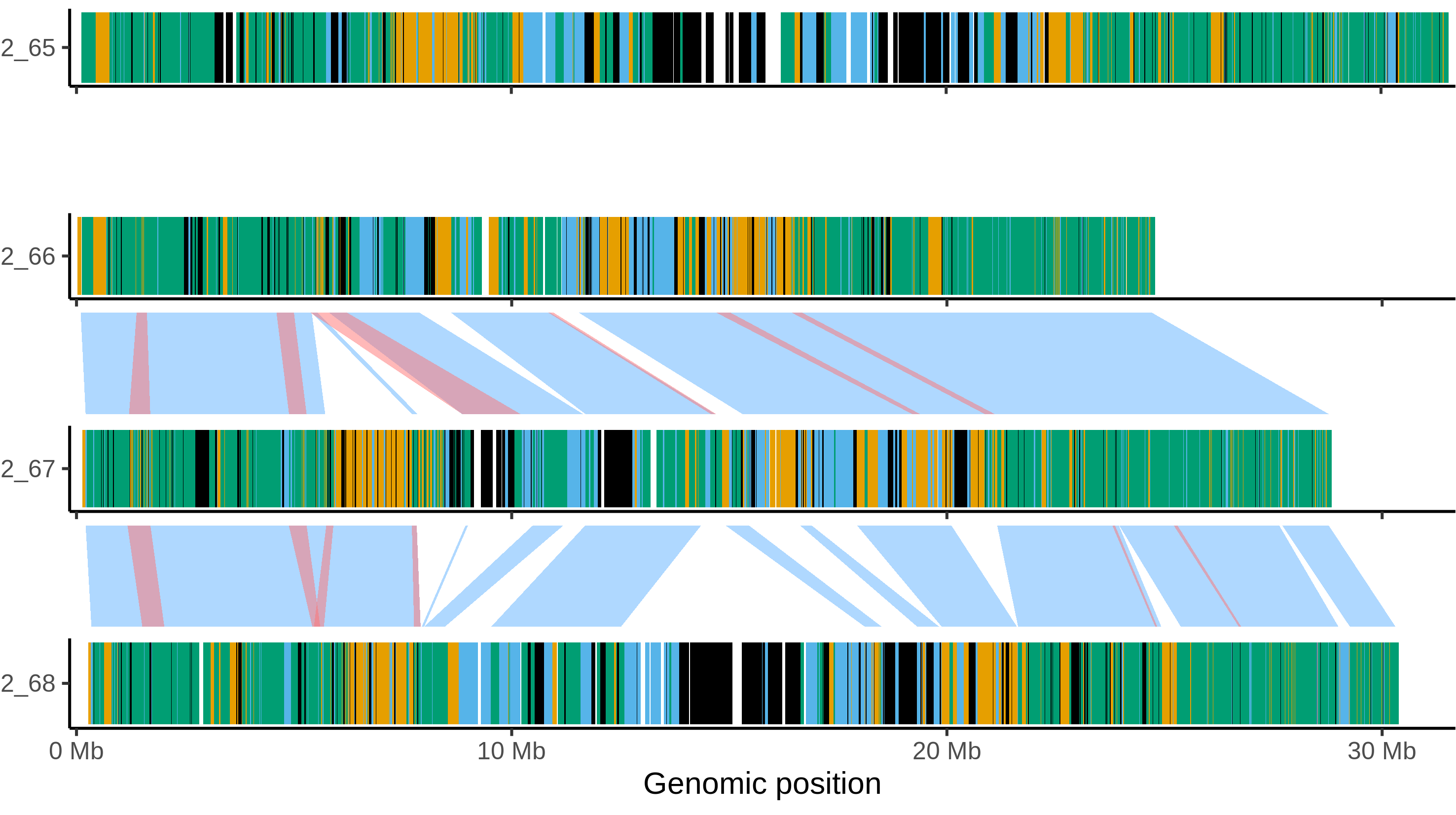

### Atlantic_chr_5.png

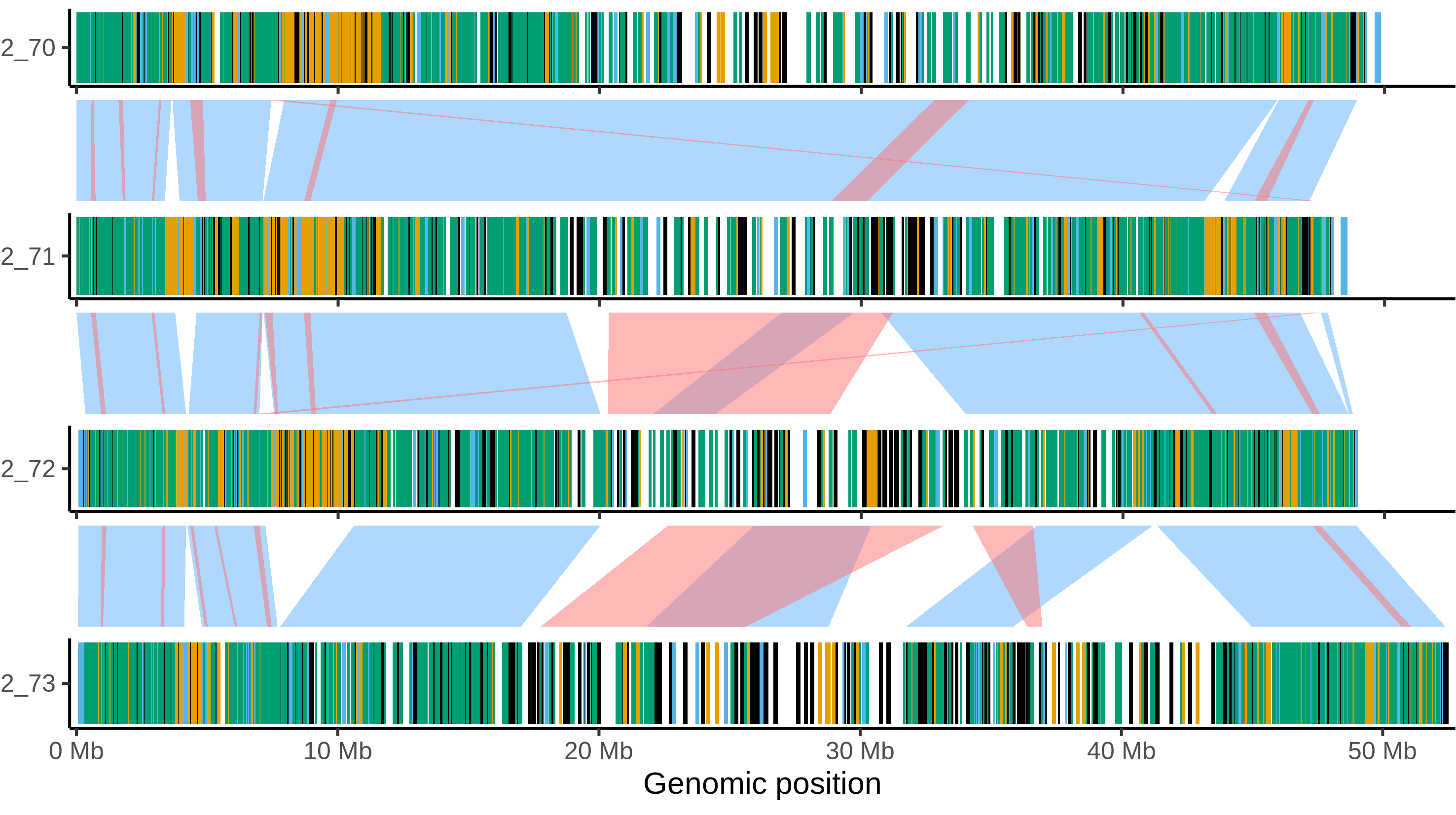

### Atlantic_chr_6.png

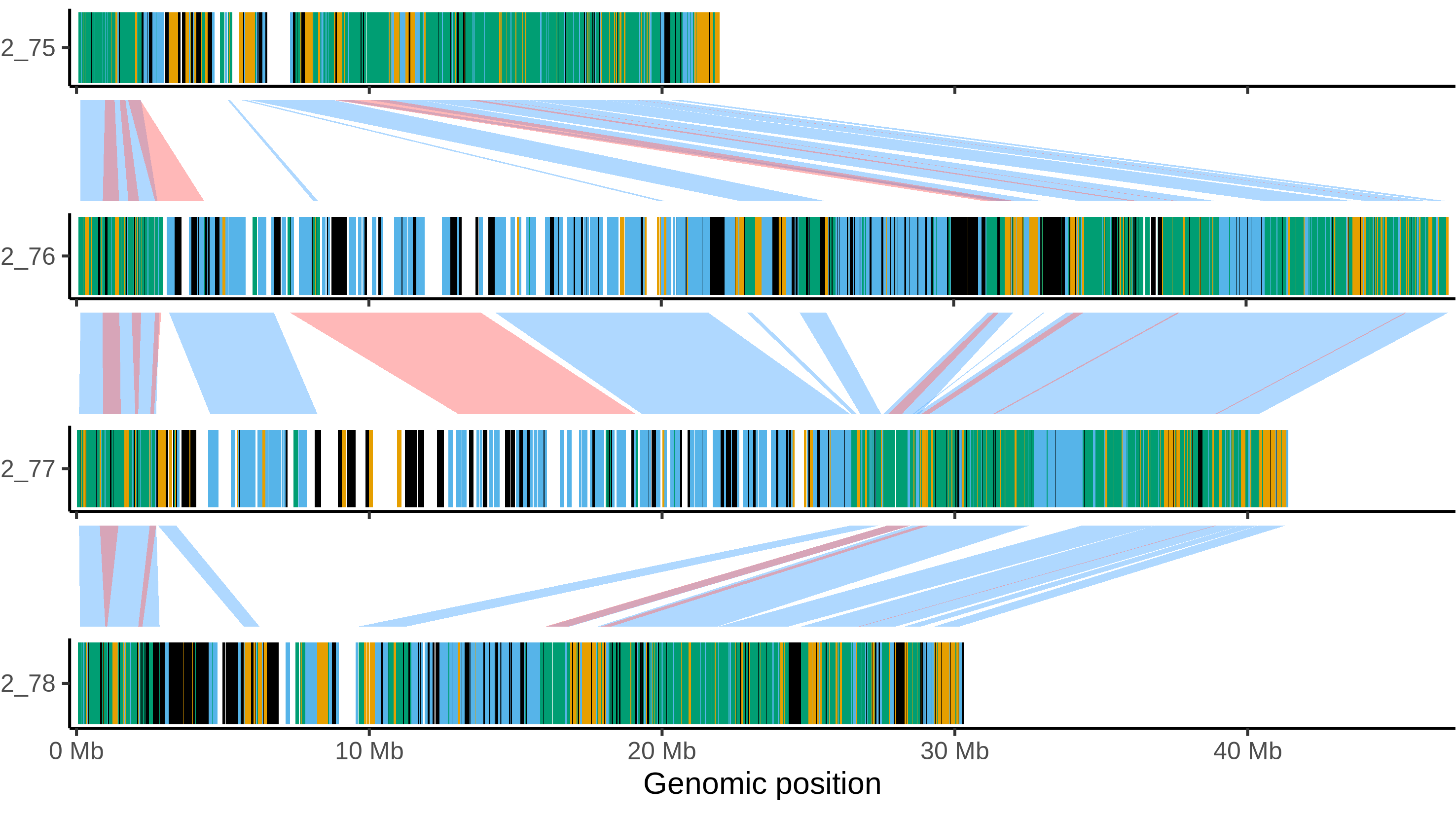

### Atlantic_chr_7.png

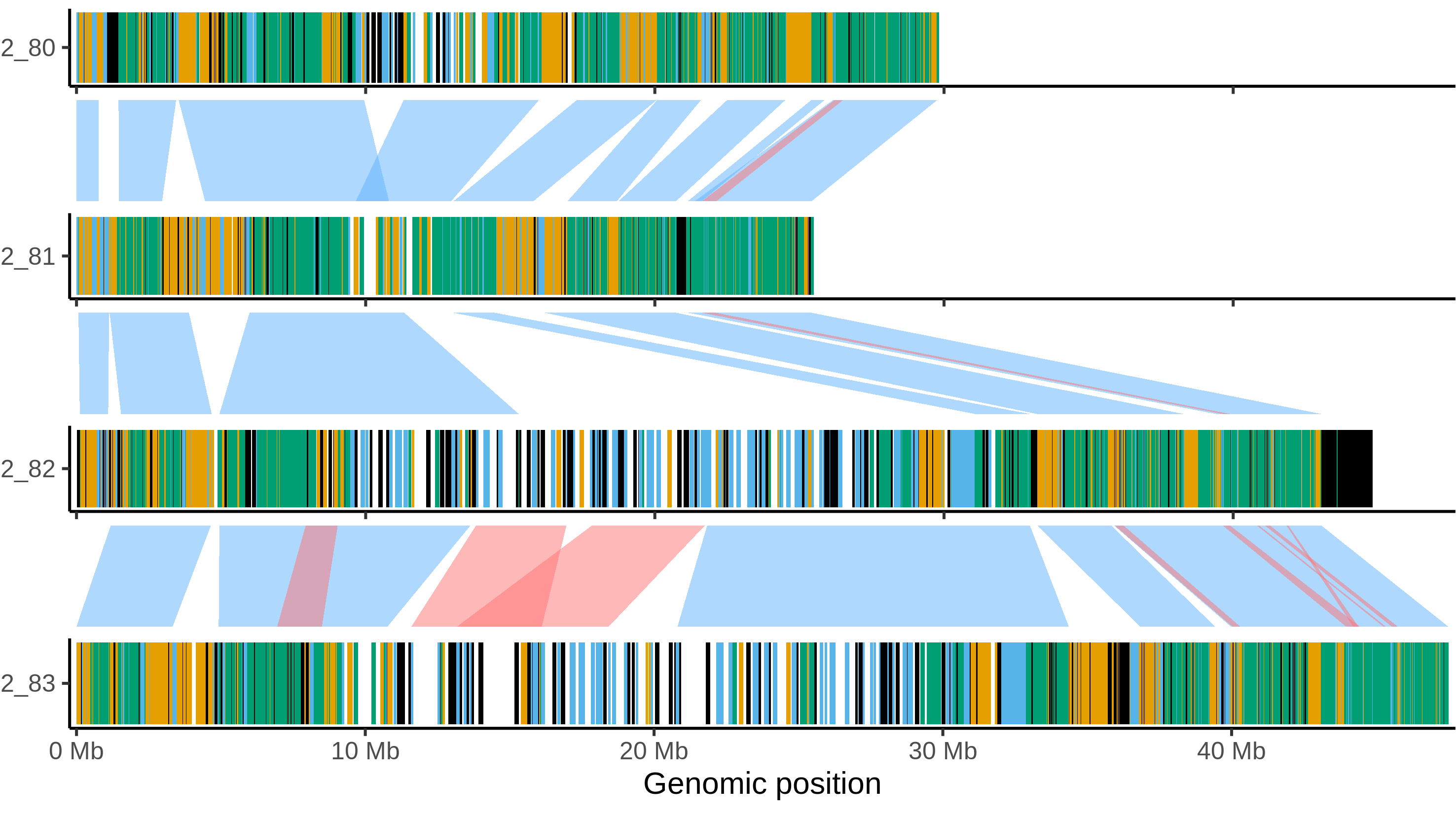

### Atlantic_chr_8.png

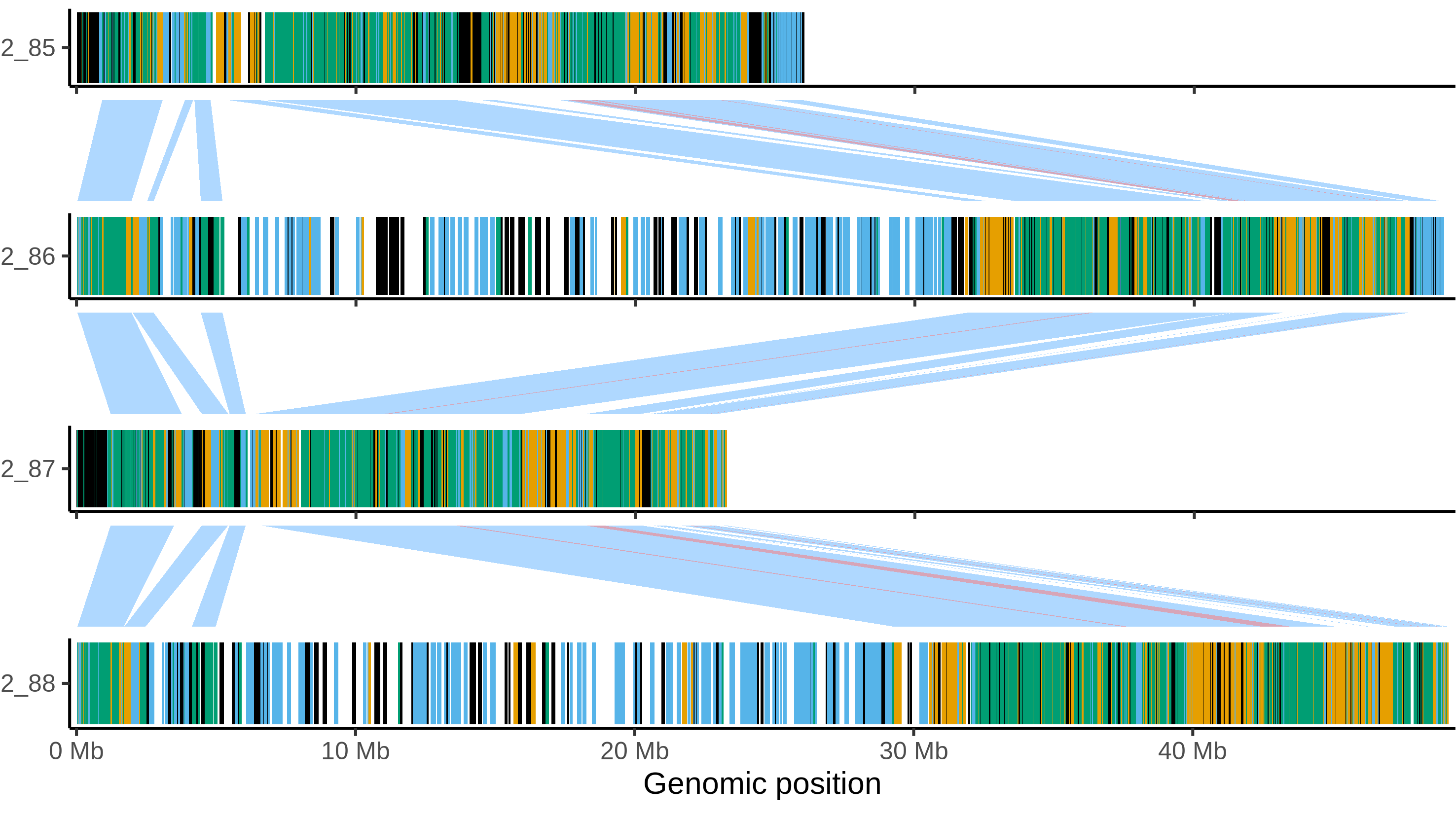

### Atlantic_chr_9.png

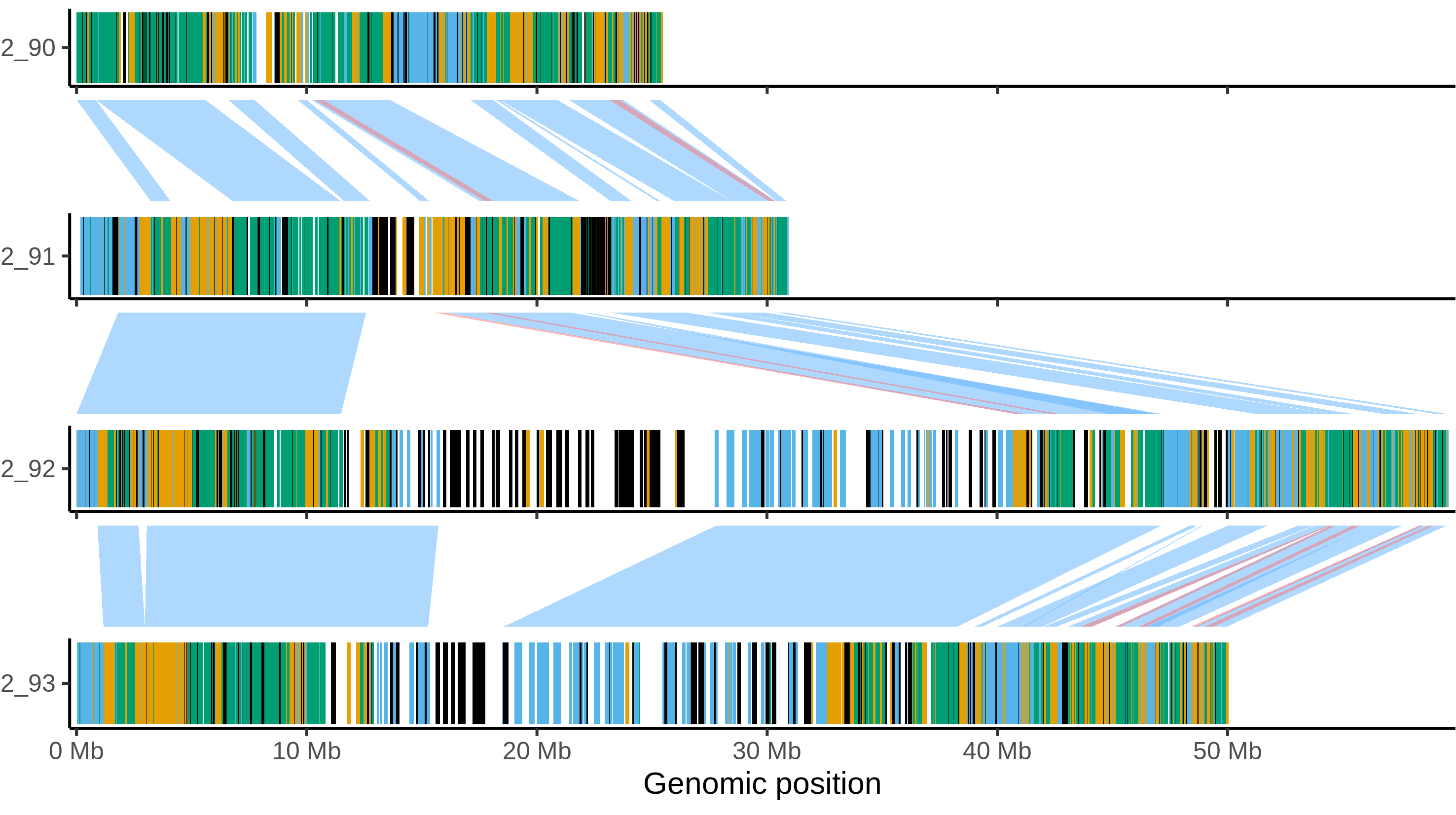

### Atlantic_chr_10.png

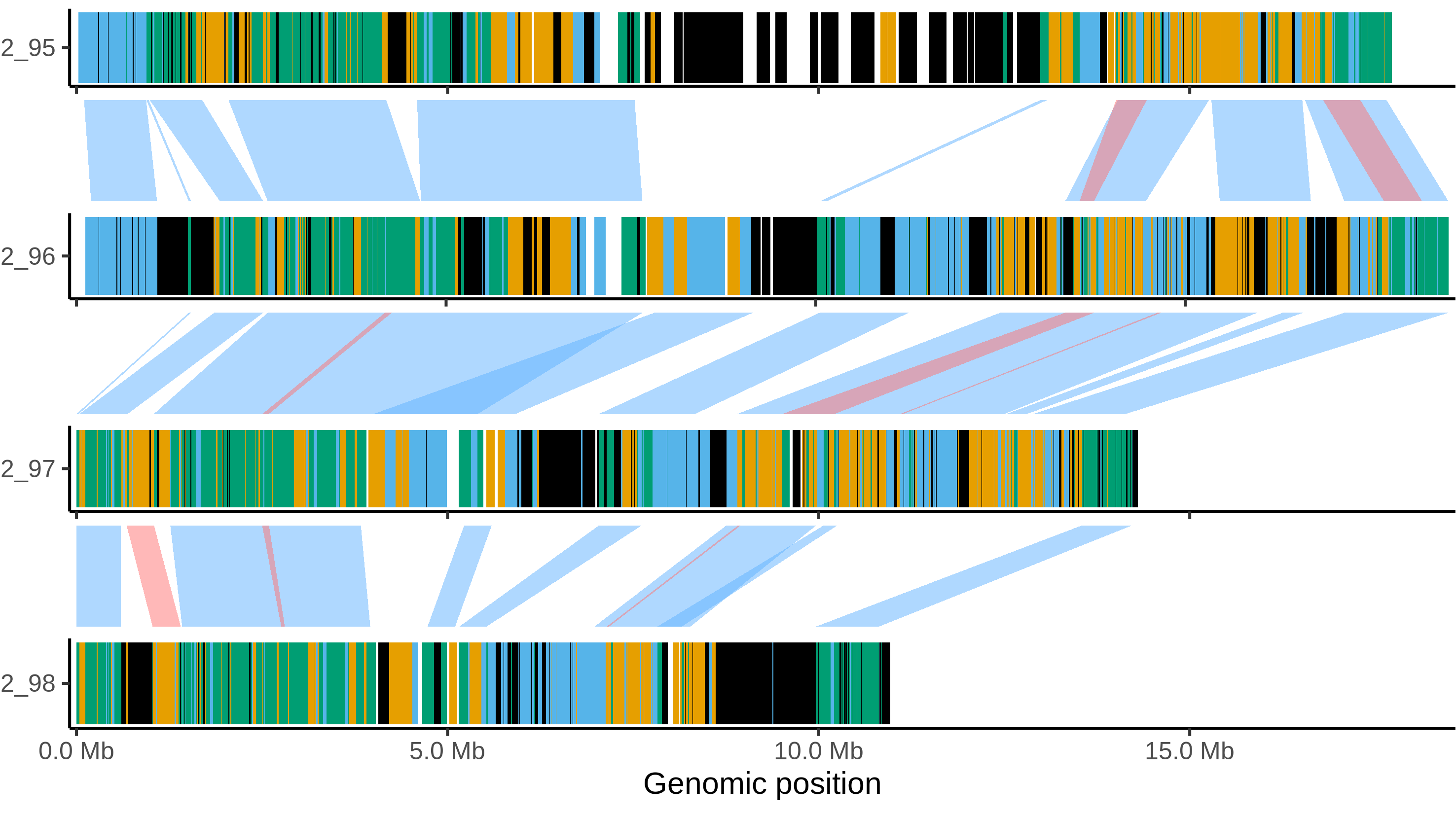

### Atlantic_chr_11.png

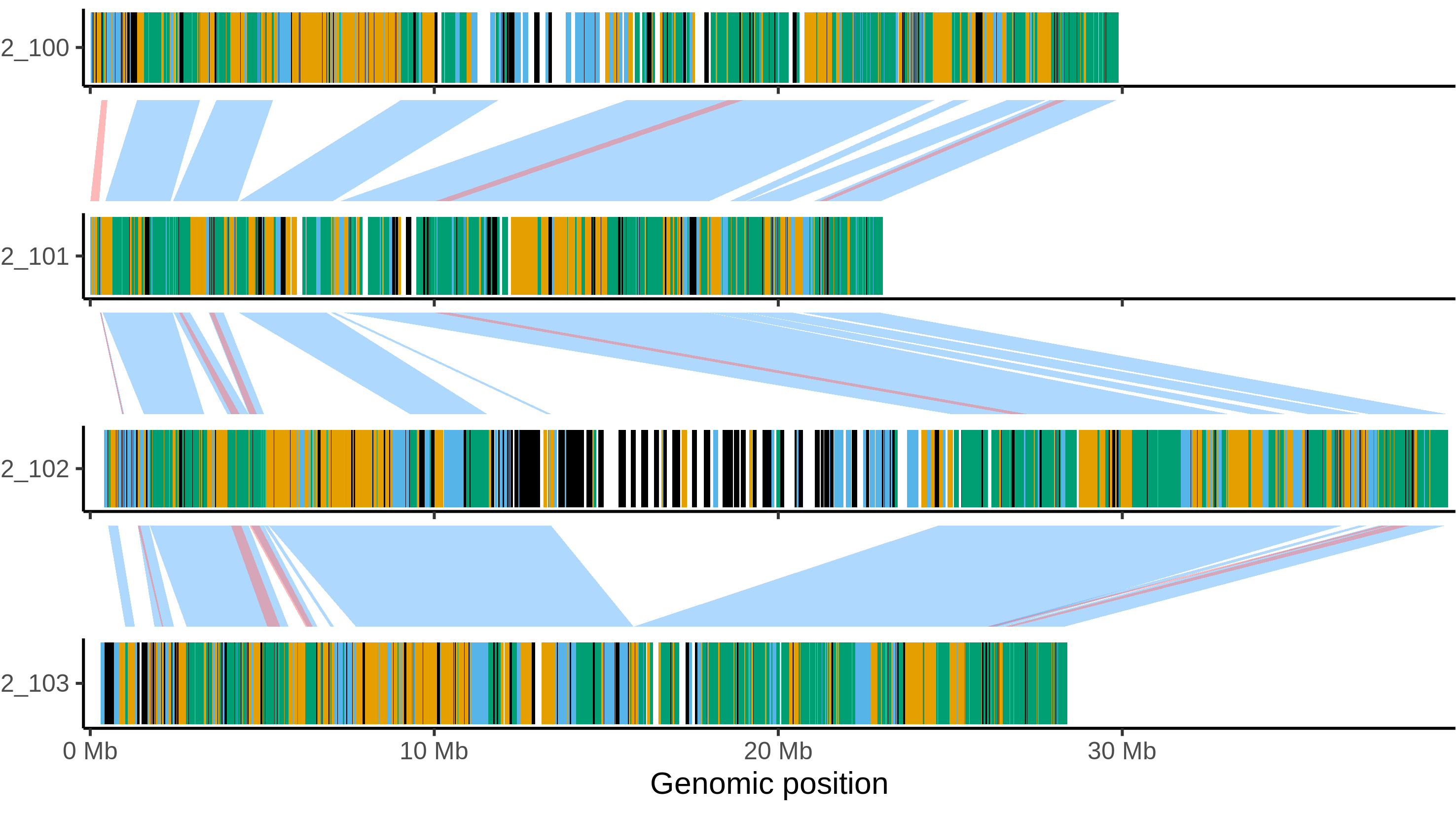

### Atlantic_chr_12.png

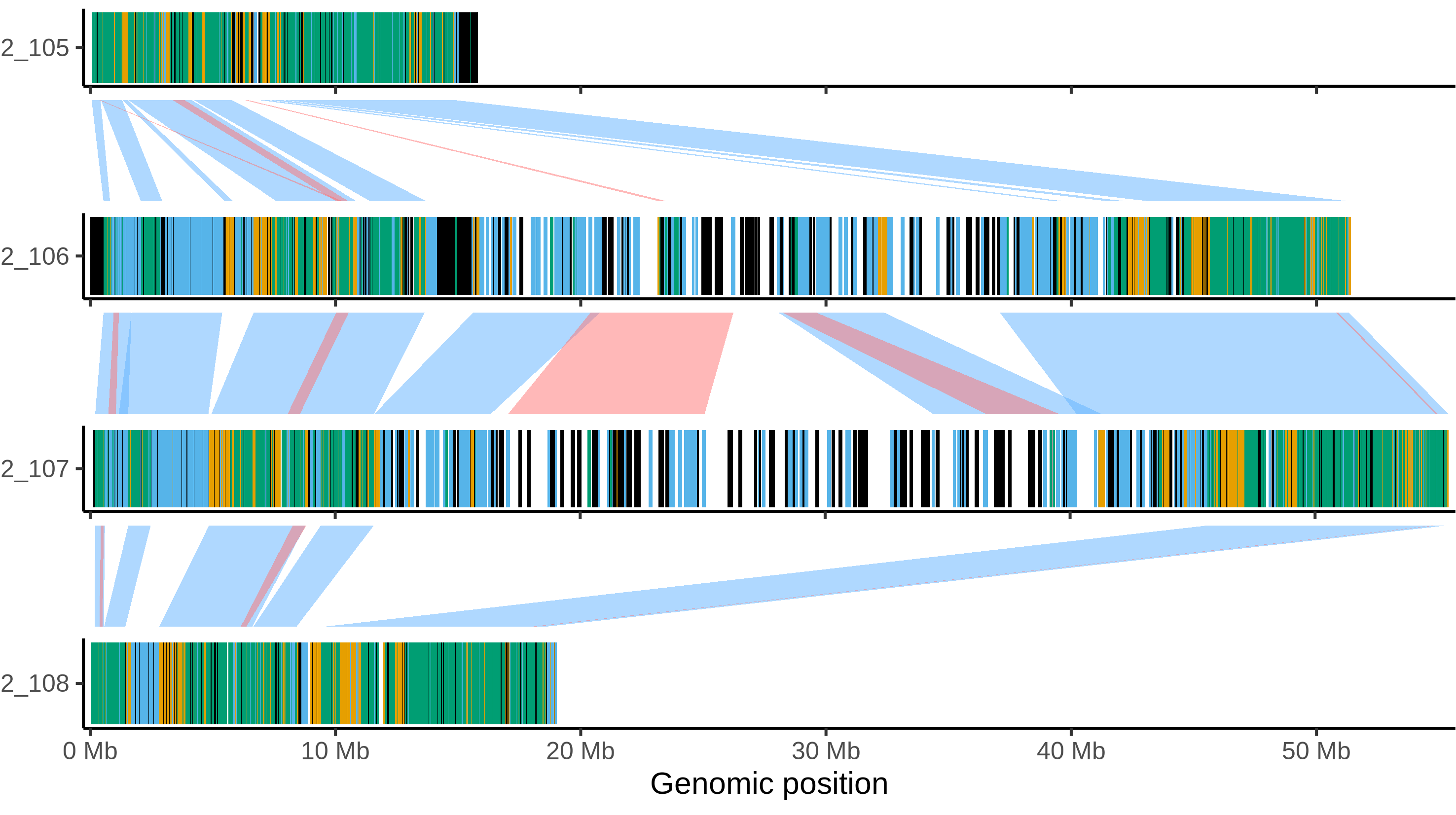

### C88_chr_1.png

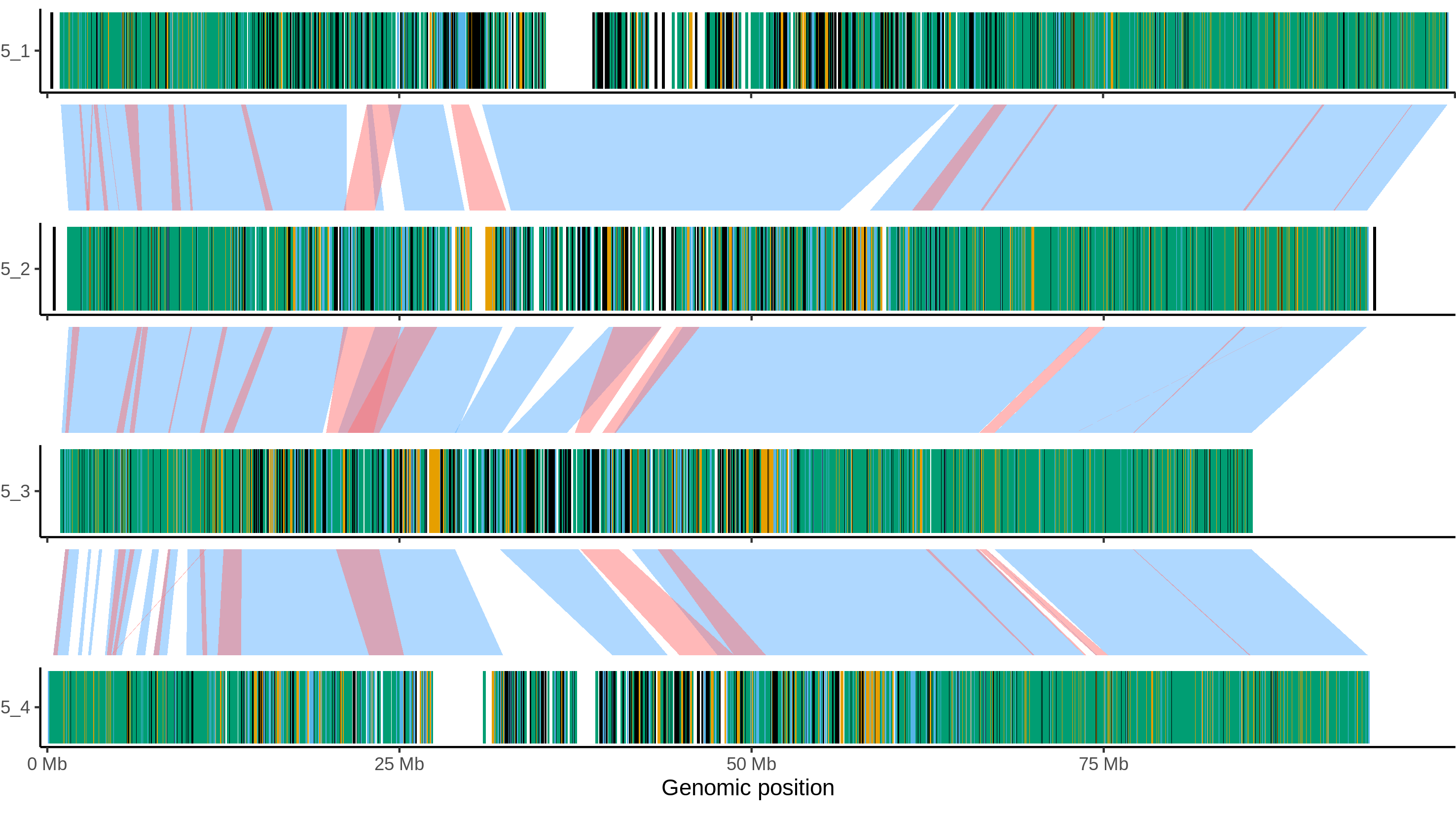

### C88_chr_2.png

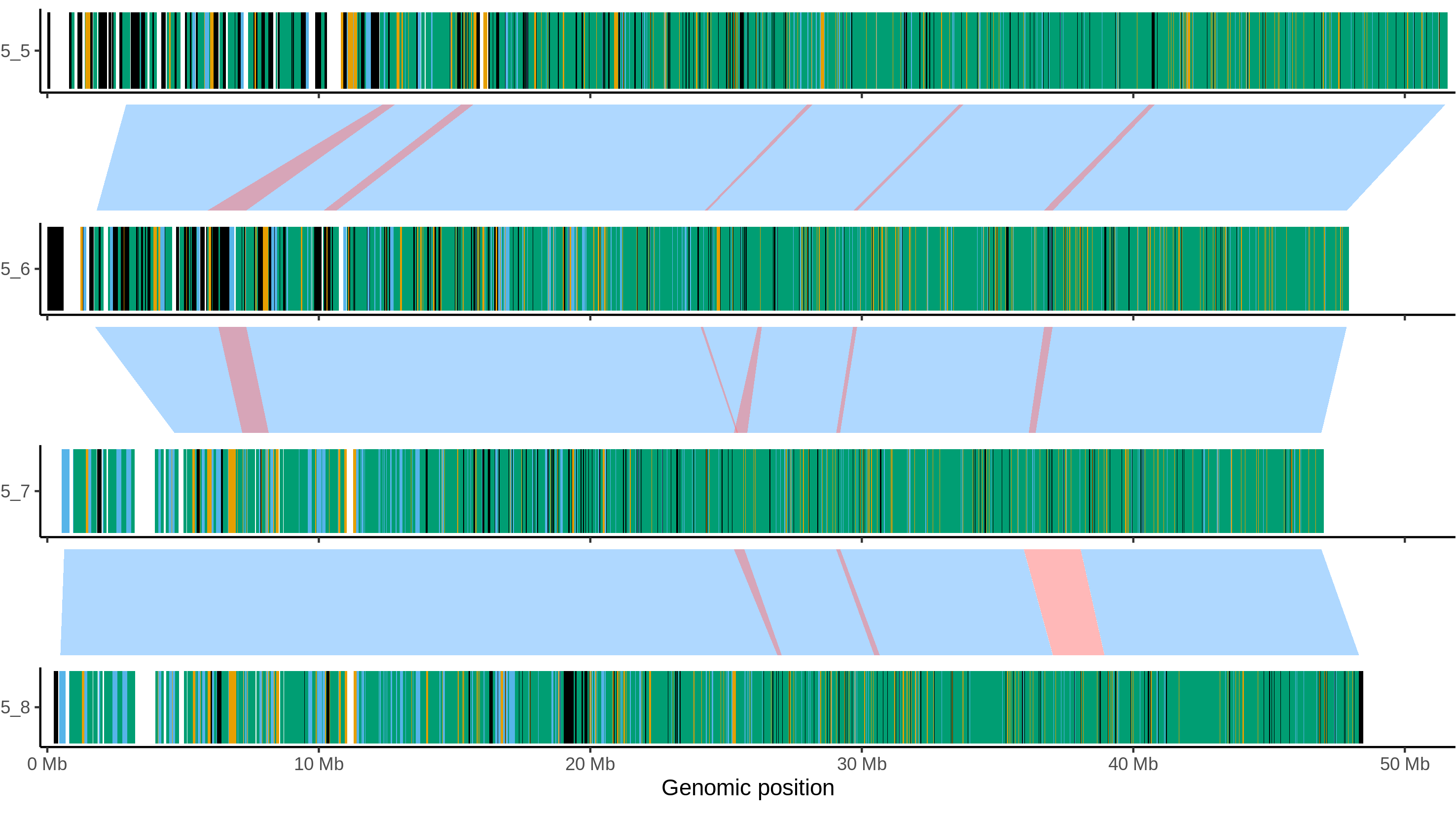

### C88_chr_3.png

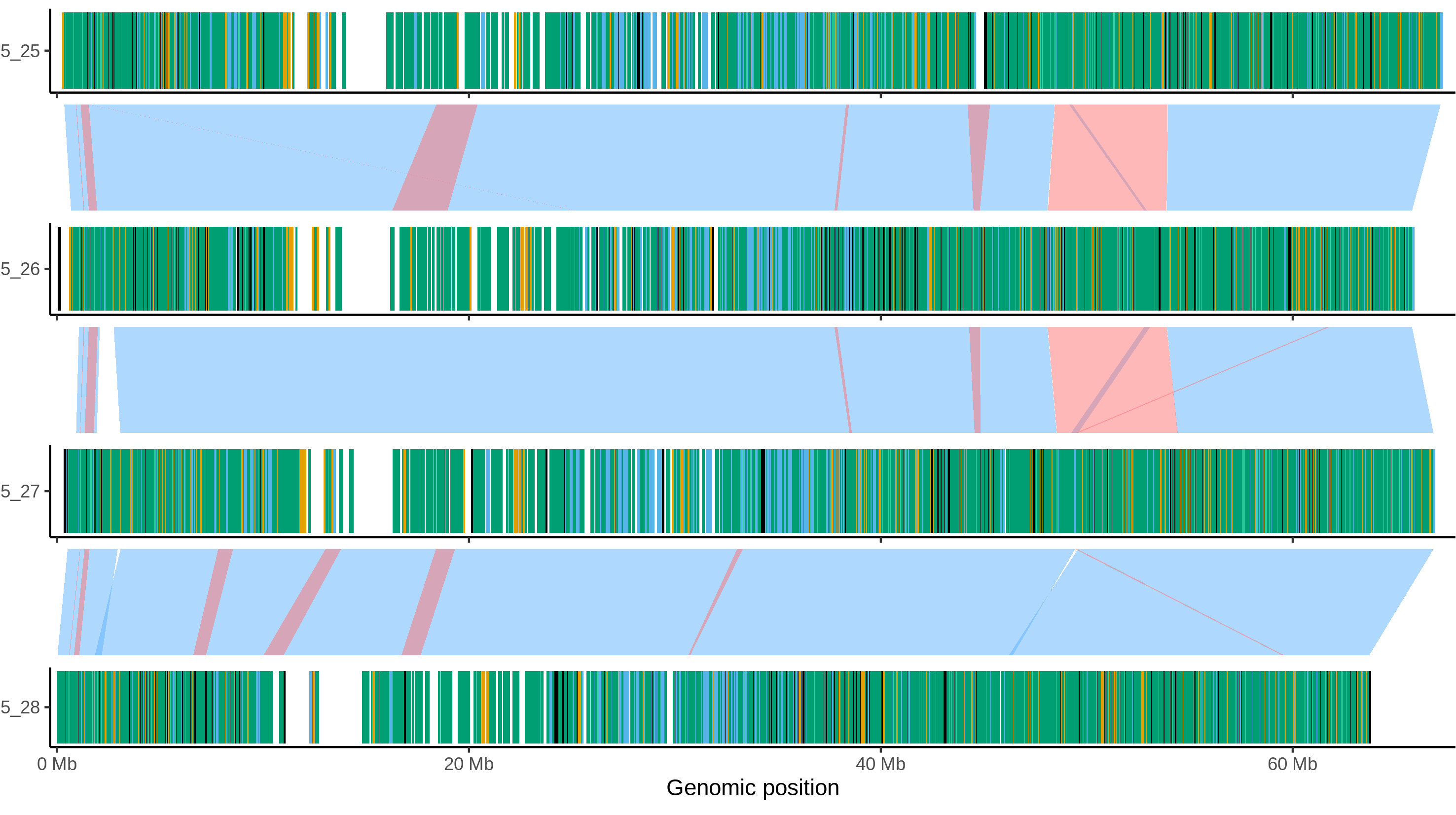

### C88_chr_4.png

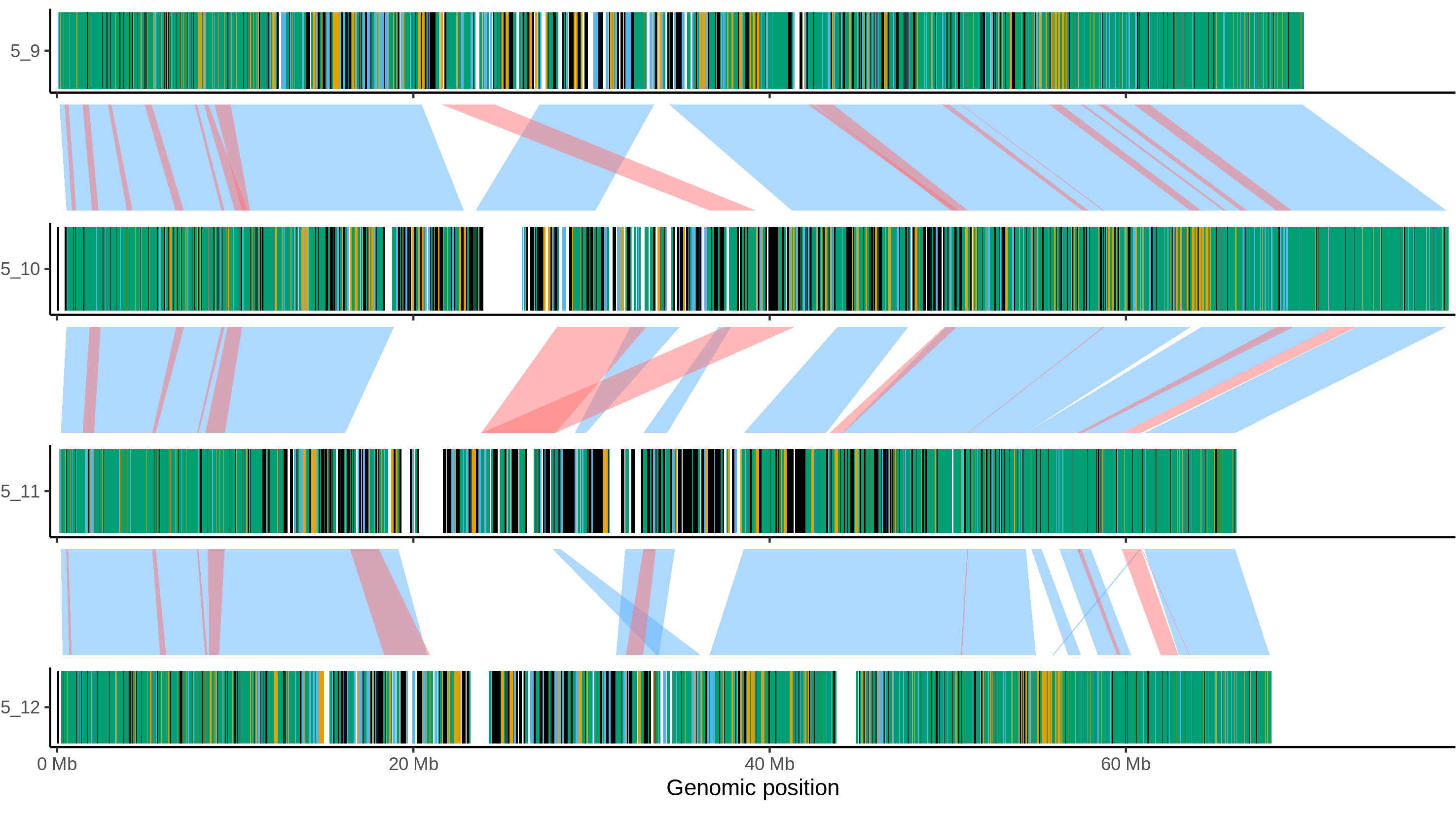

### C88_chr_5.png

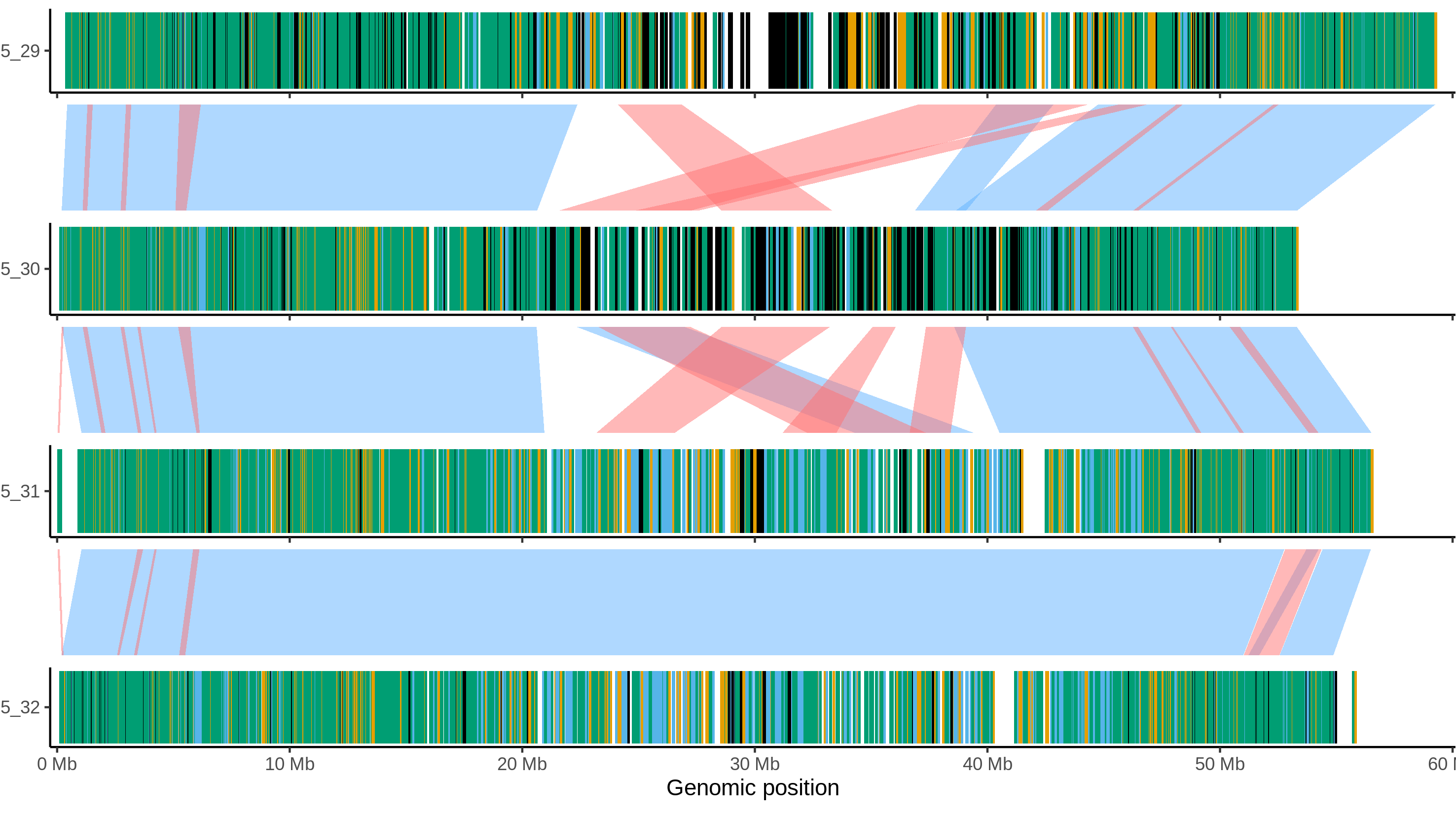

### C88_chr_6.png

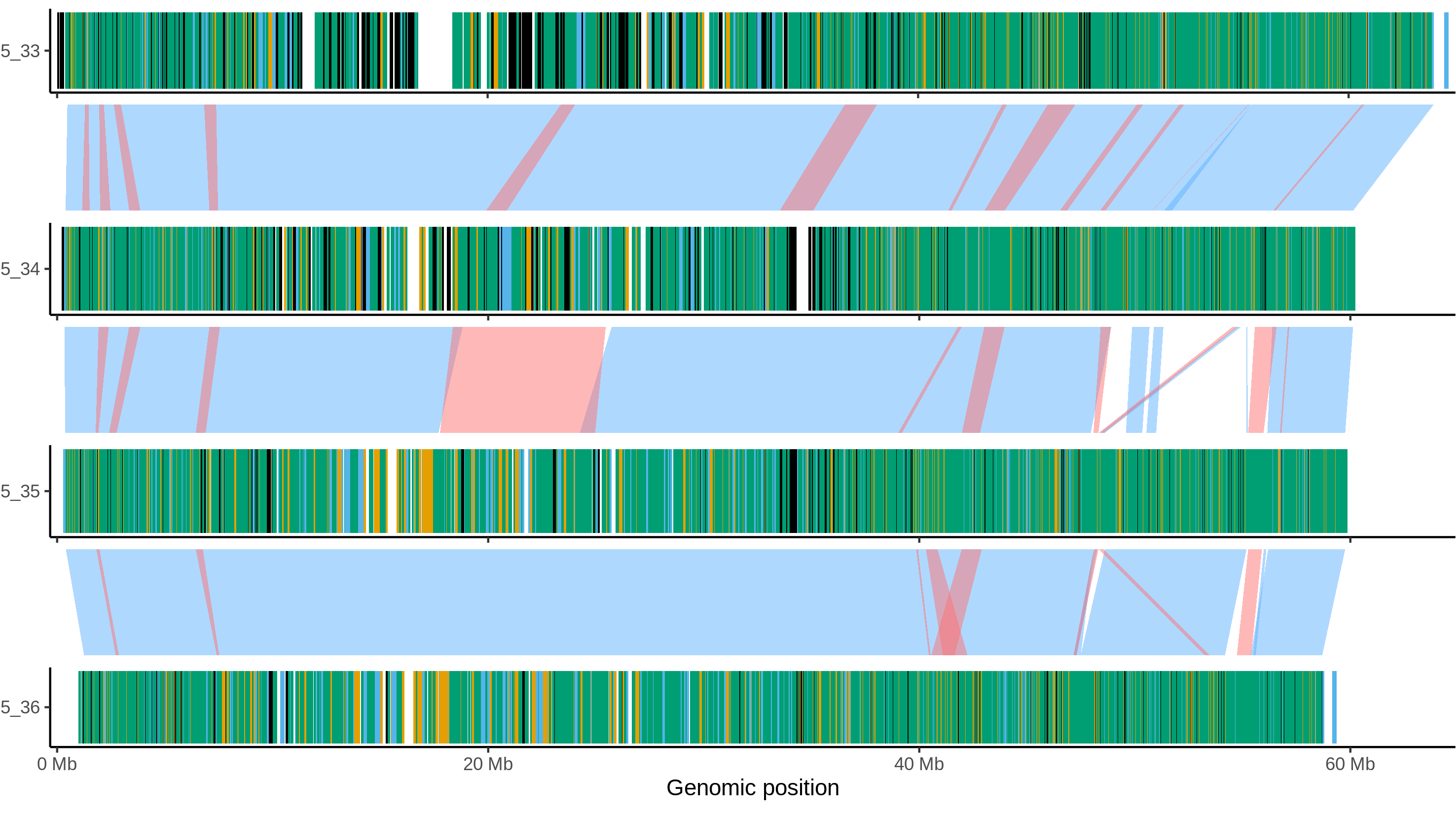

### C88_chr_7.png

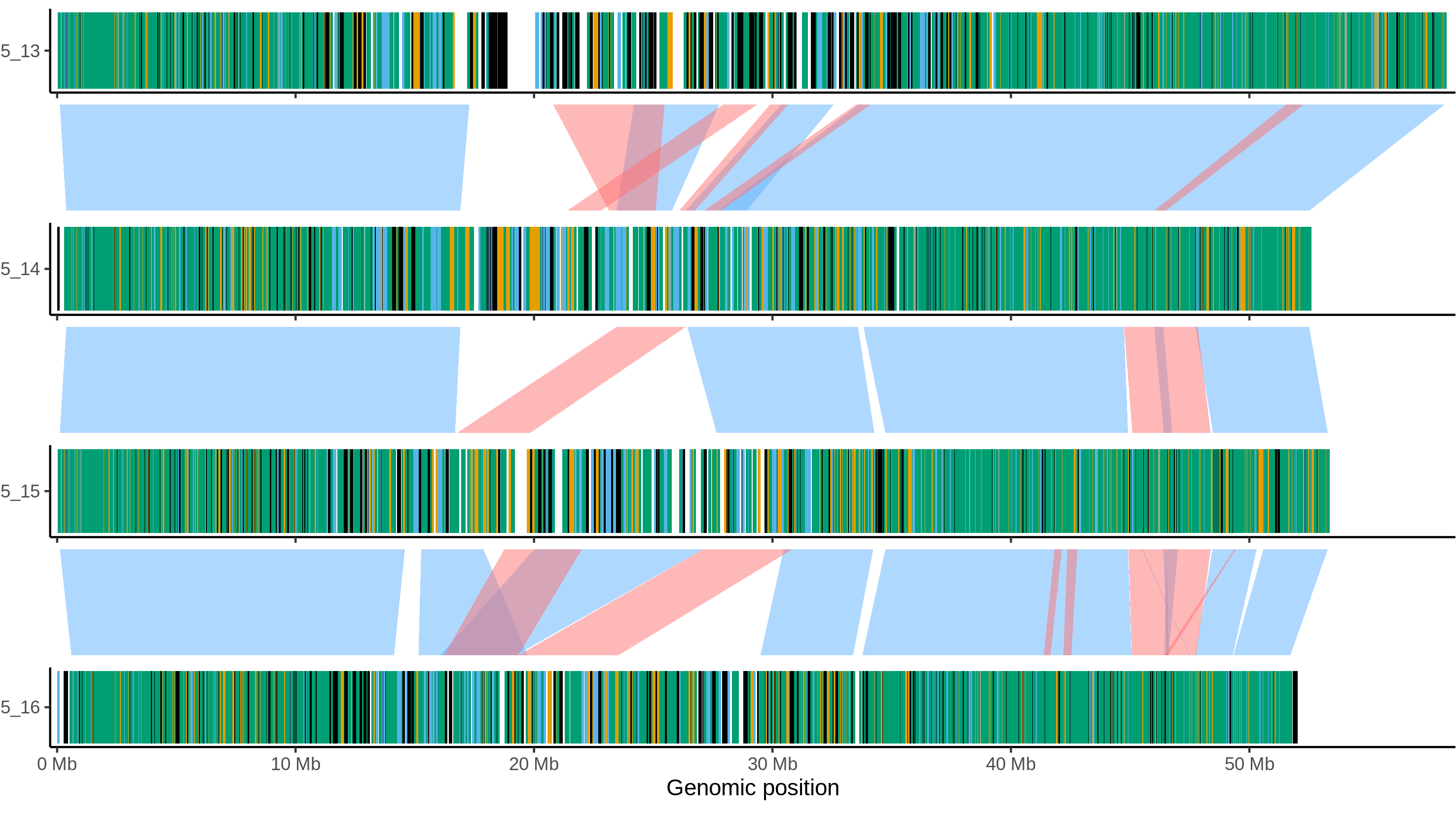

### C88_chr_8.png

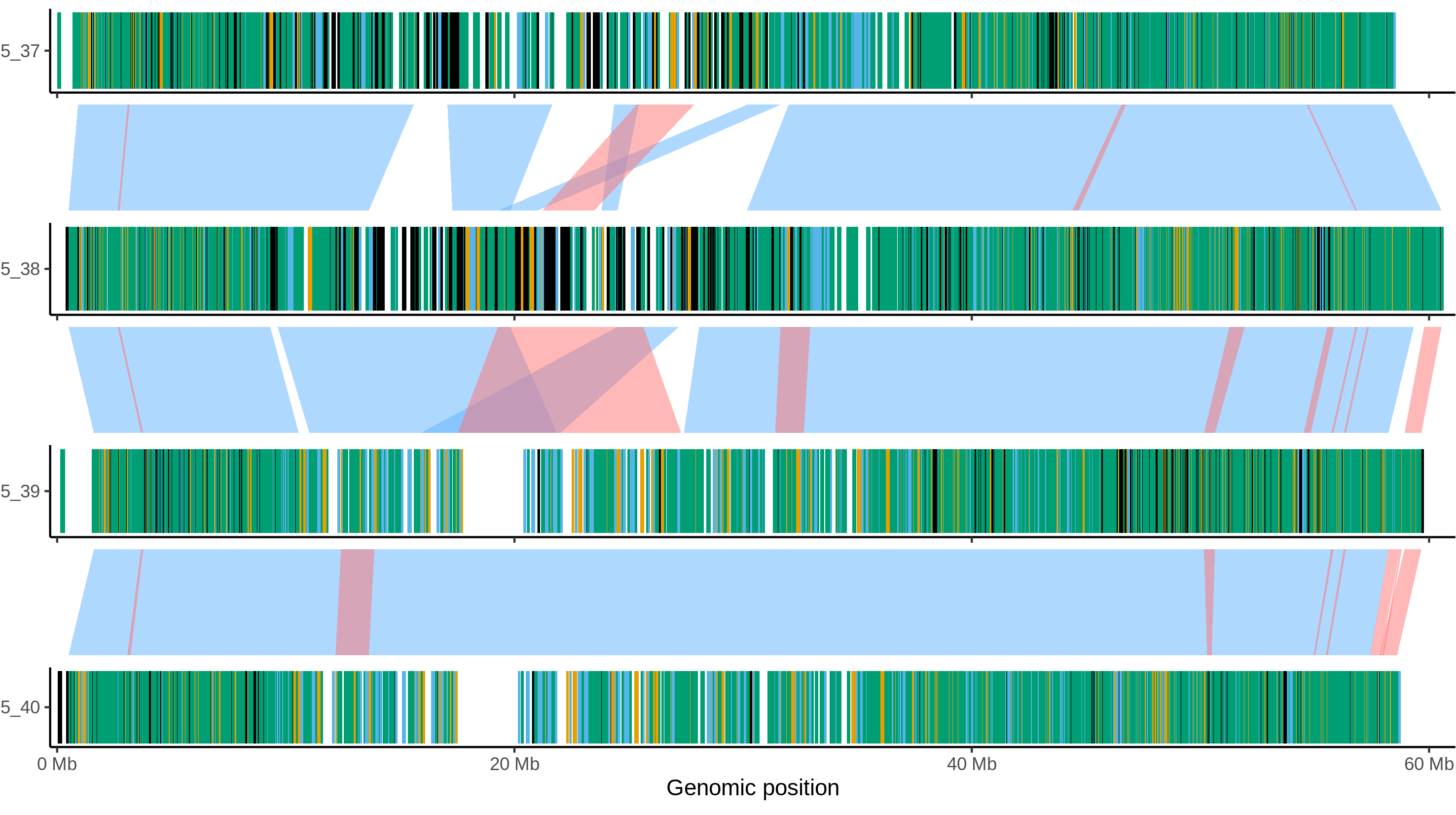

### C88_chr_9.png

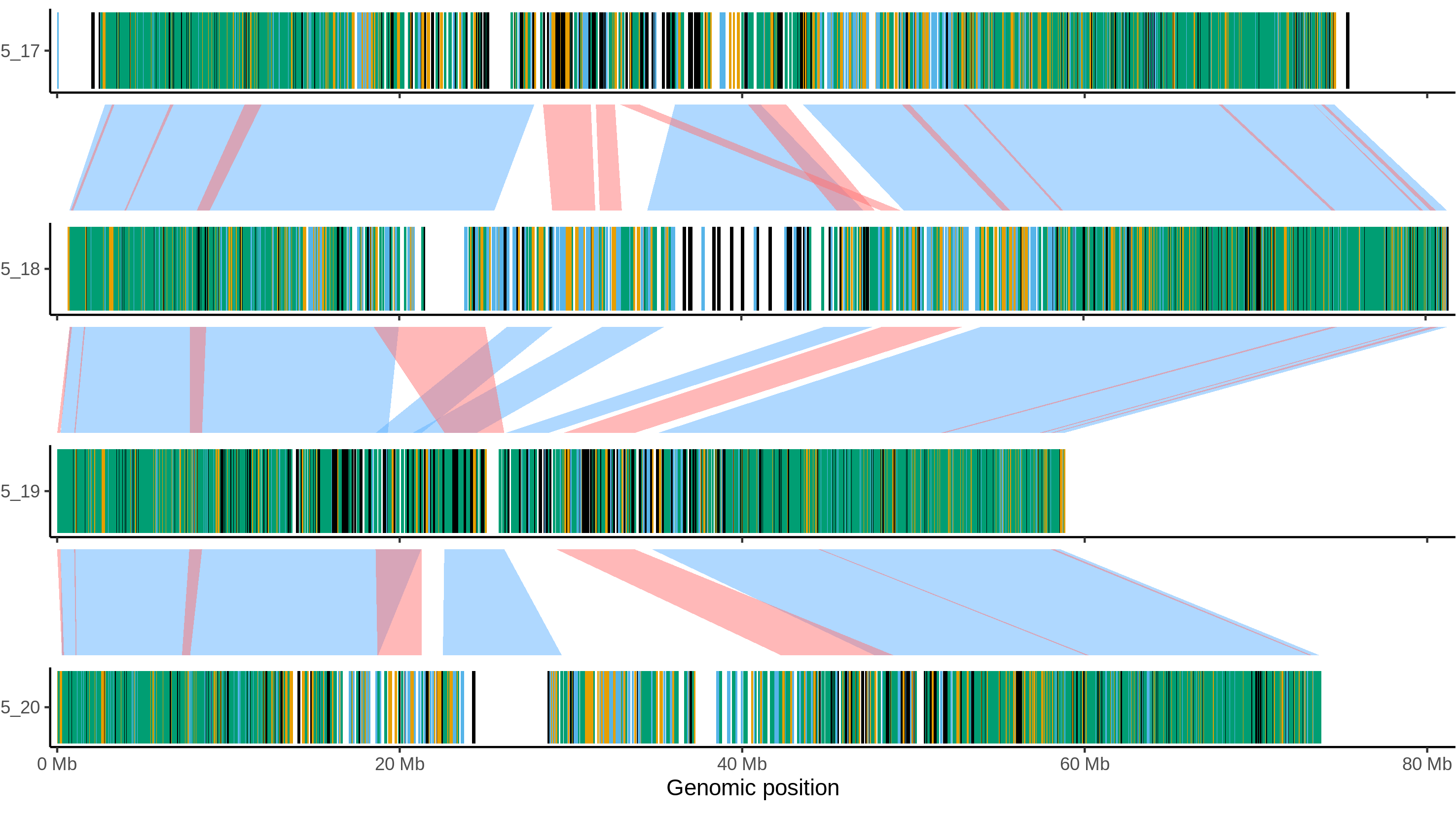

### C88_chr_10.png

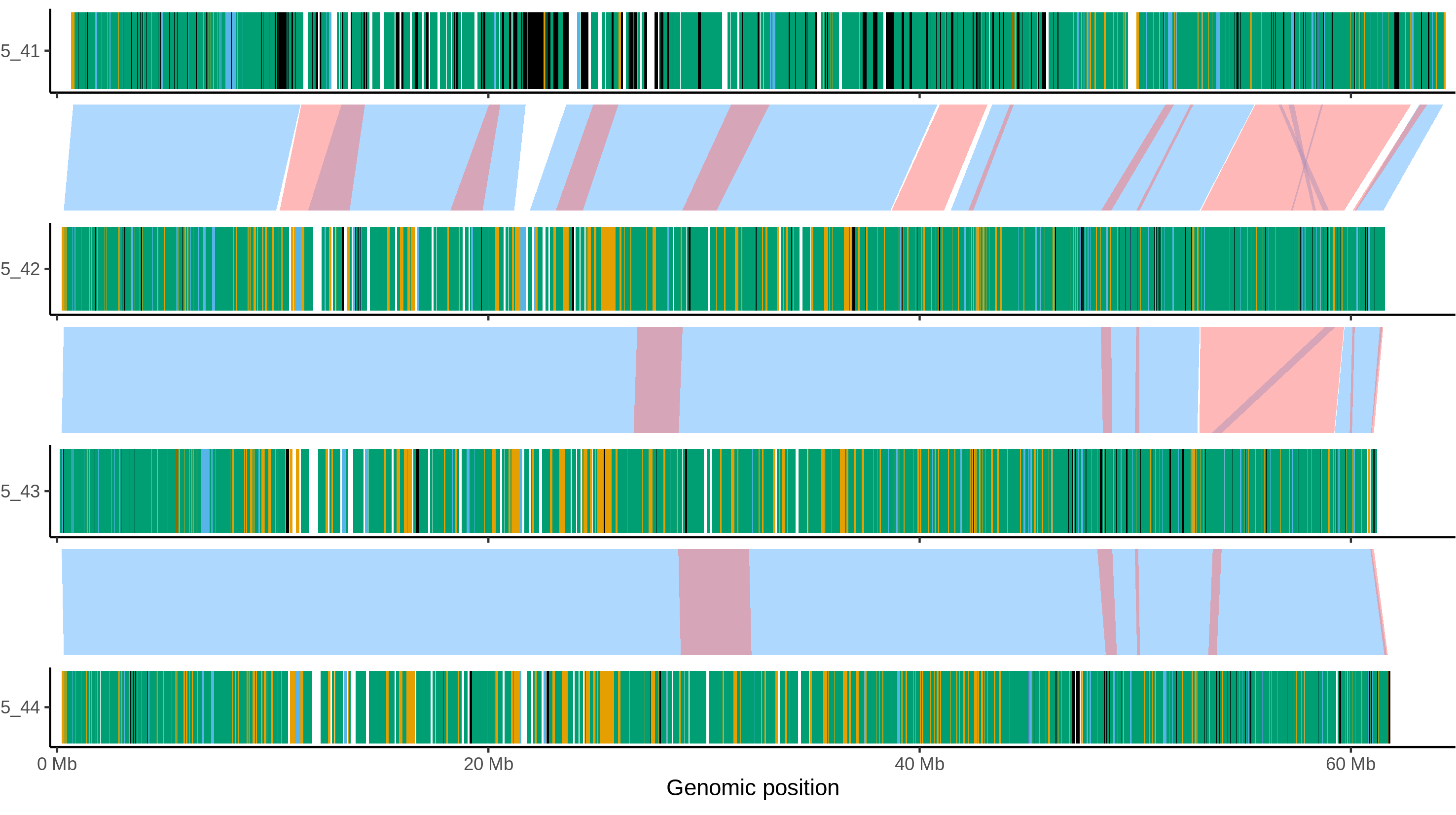

### C88_chr_11.png

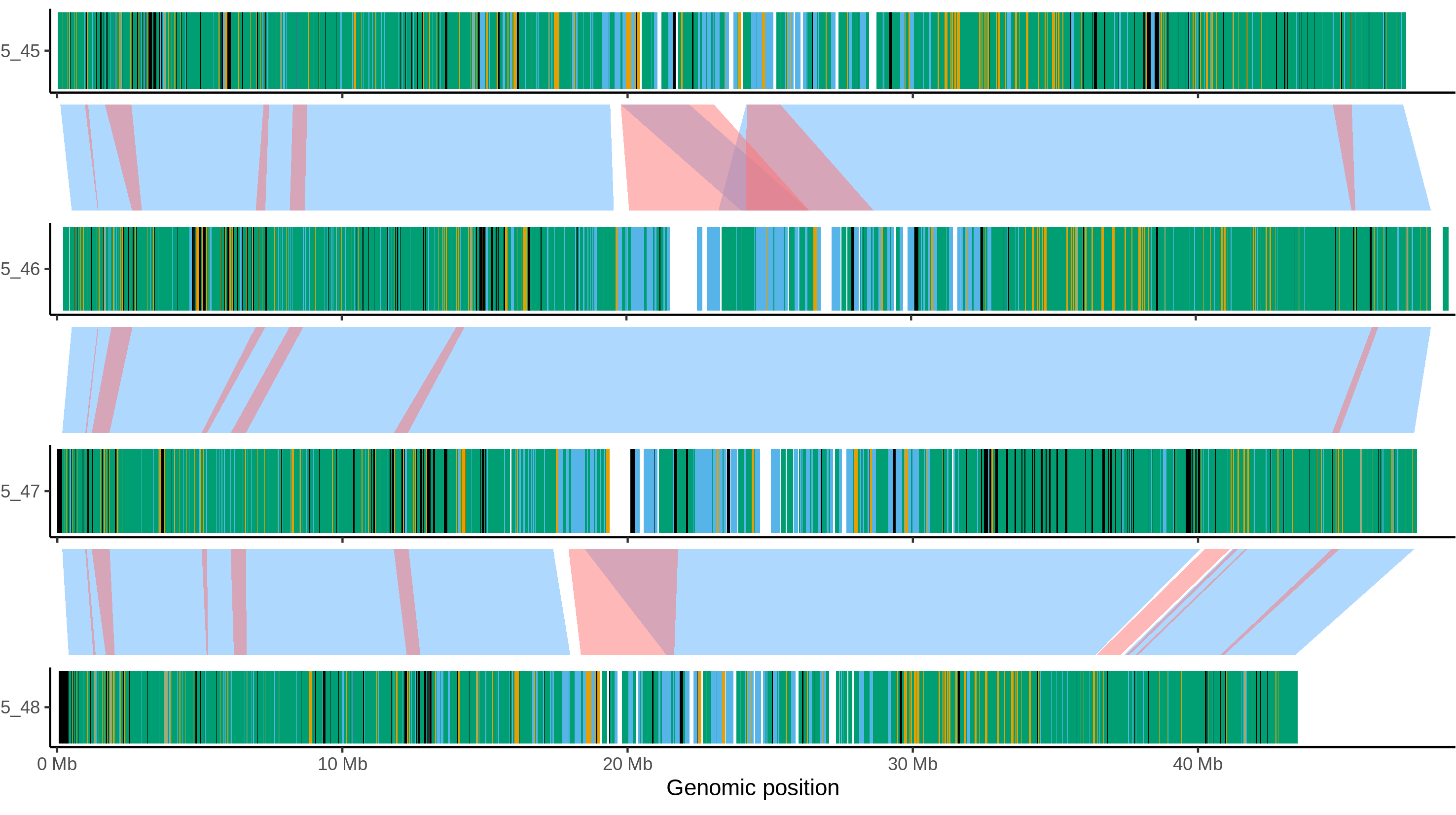

### C88_chr_12.png

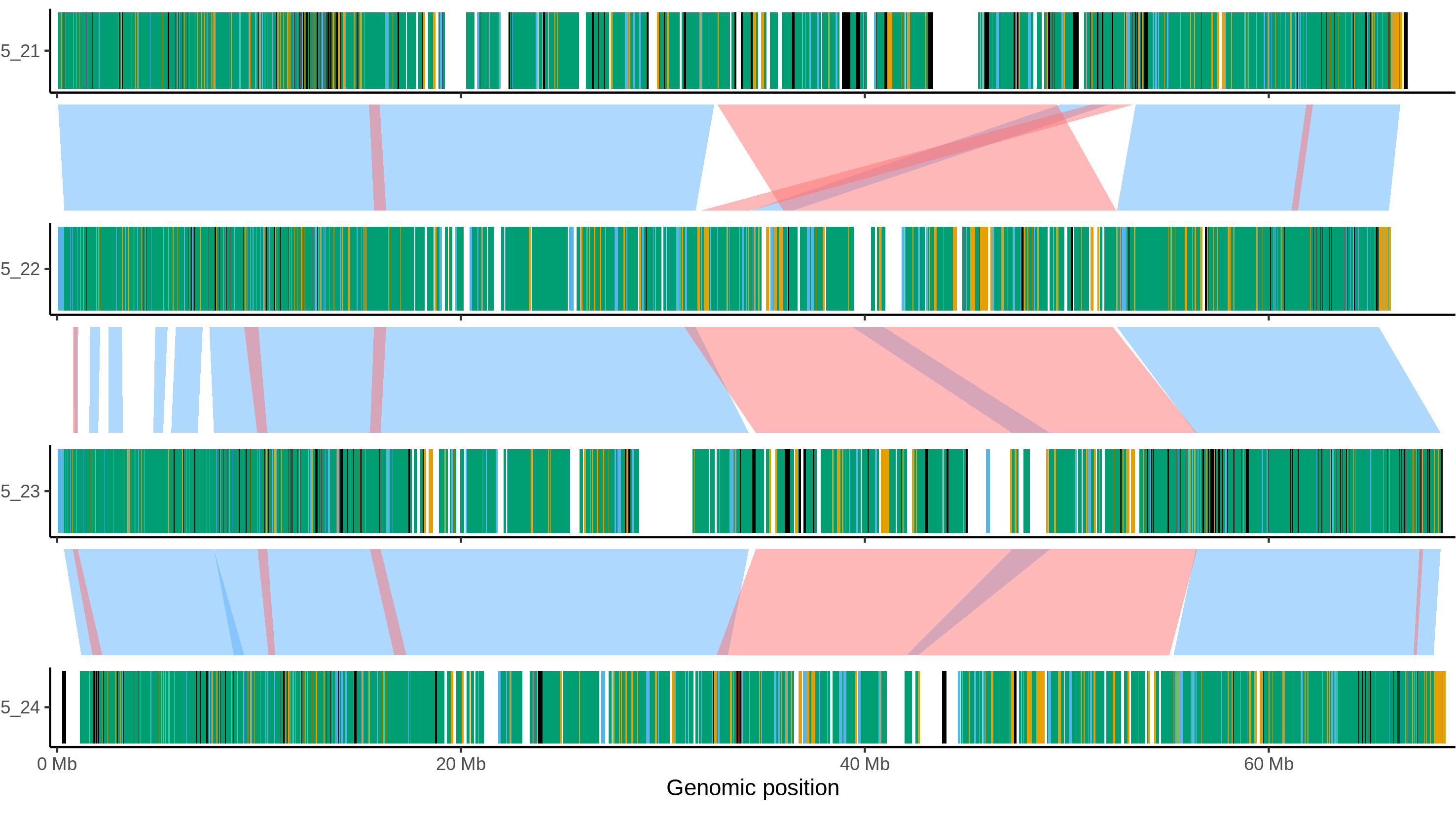

### Castle_russet_chr_1.png

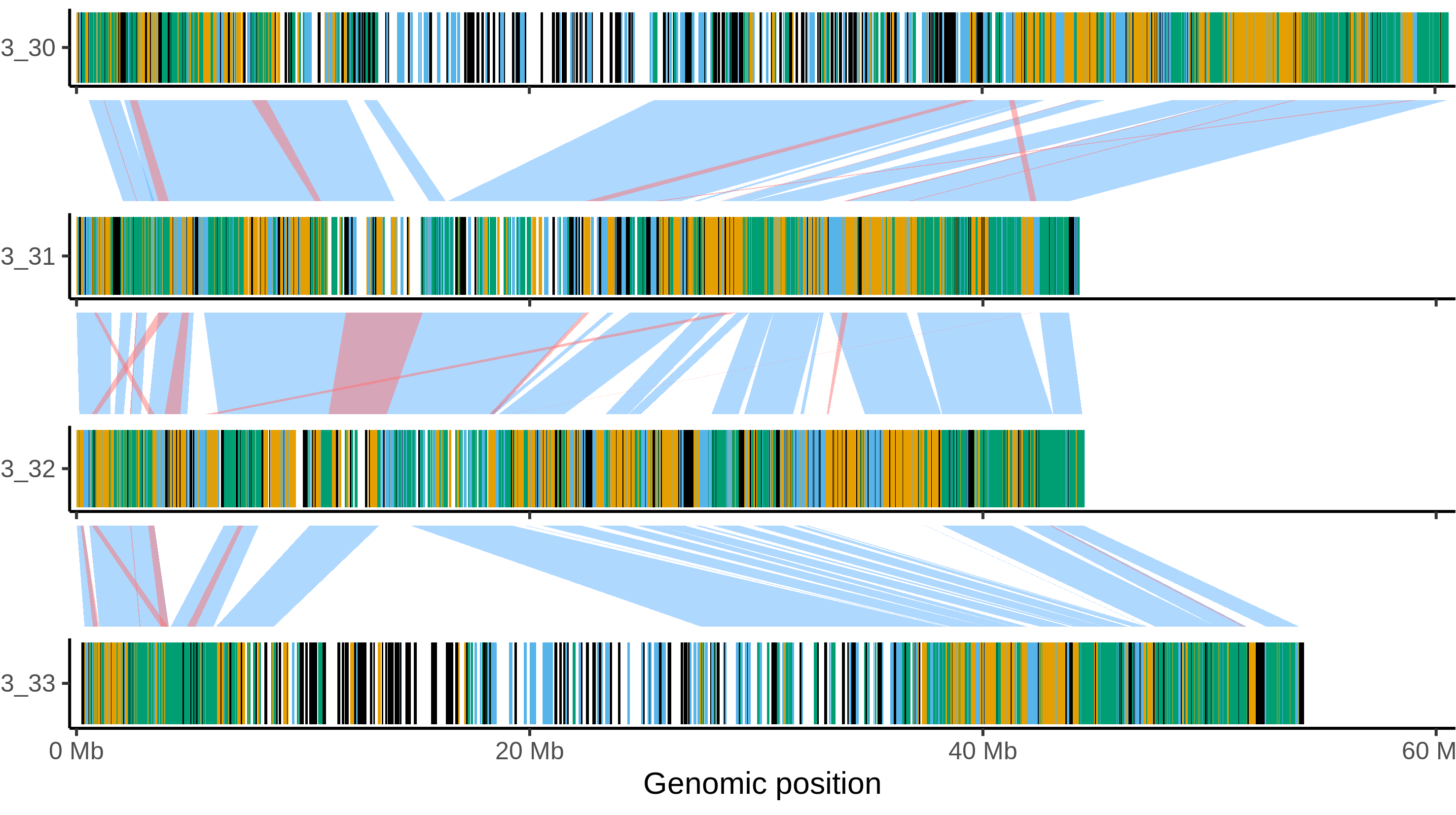

### Castle_russet_chr_2.png

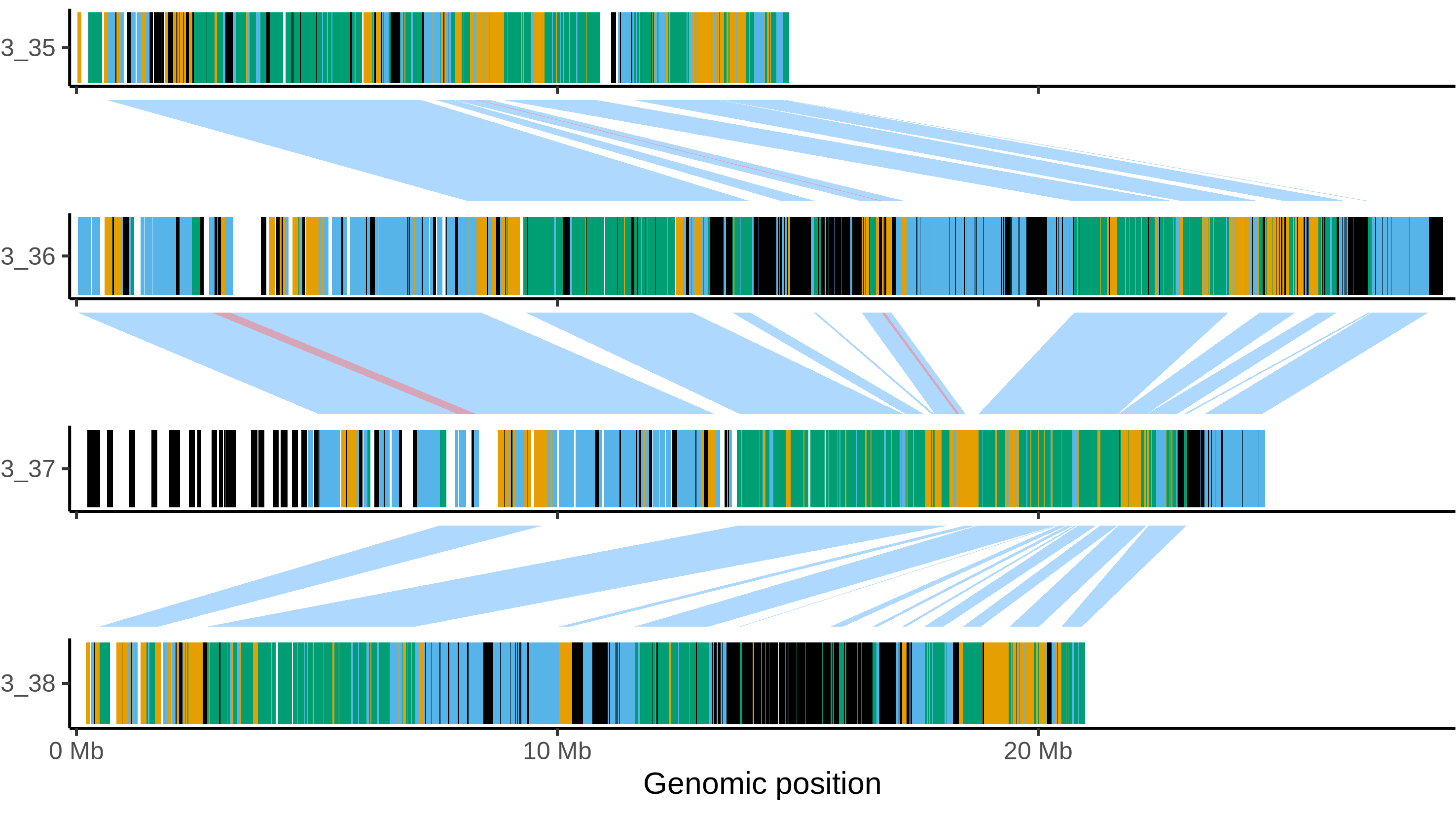

### Castle_russet_chr_3.png

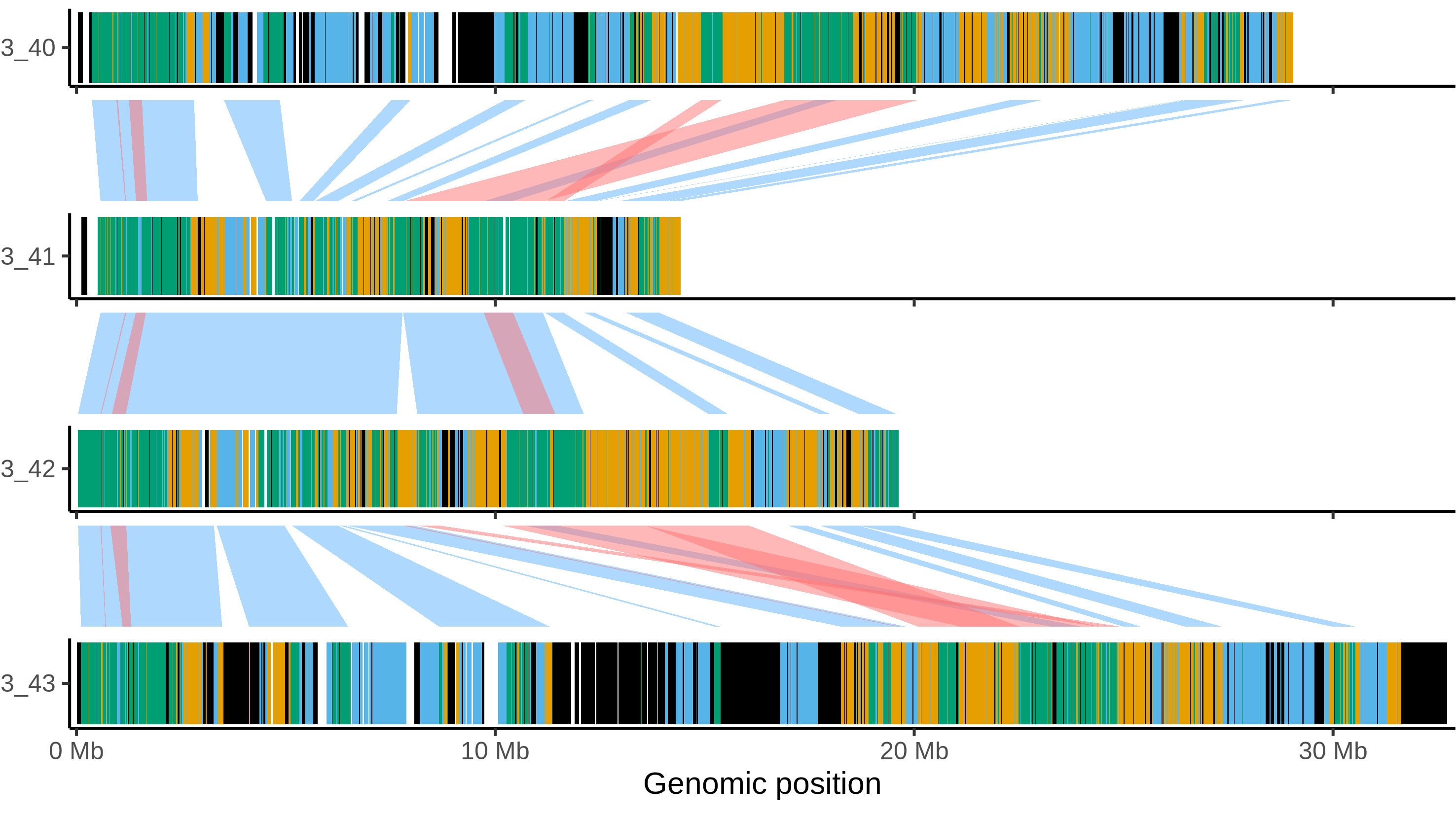

### Castle_russet_chr_4.png

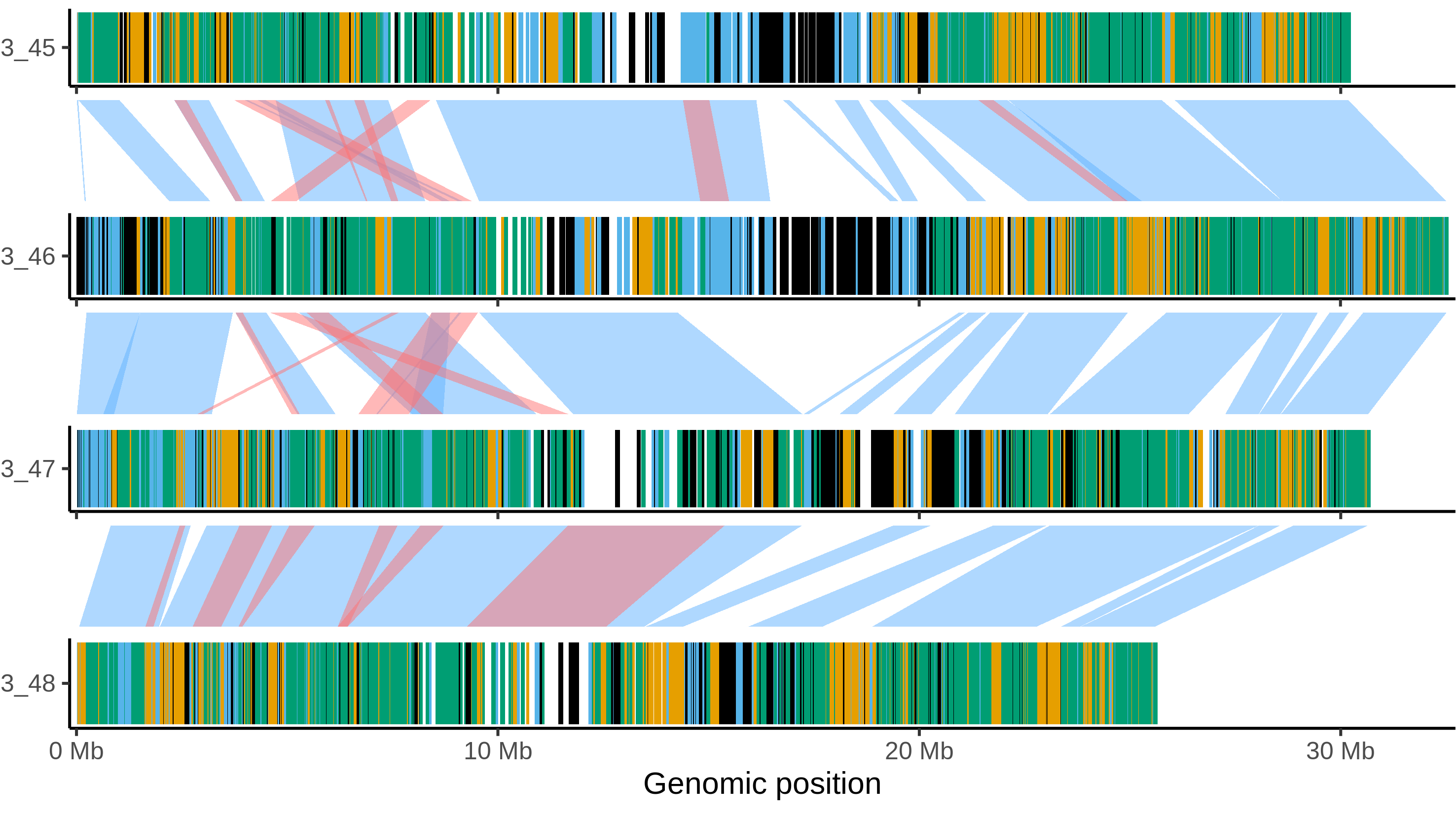

### Castle_russet_chr_5.png

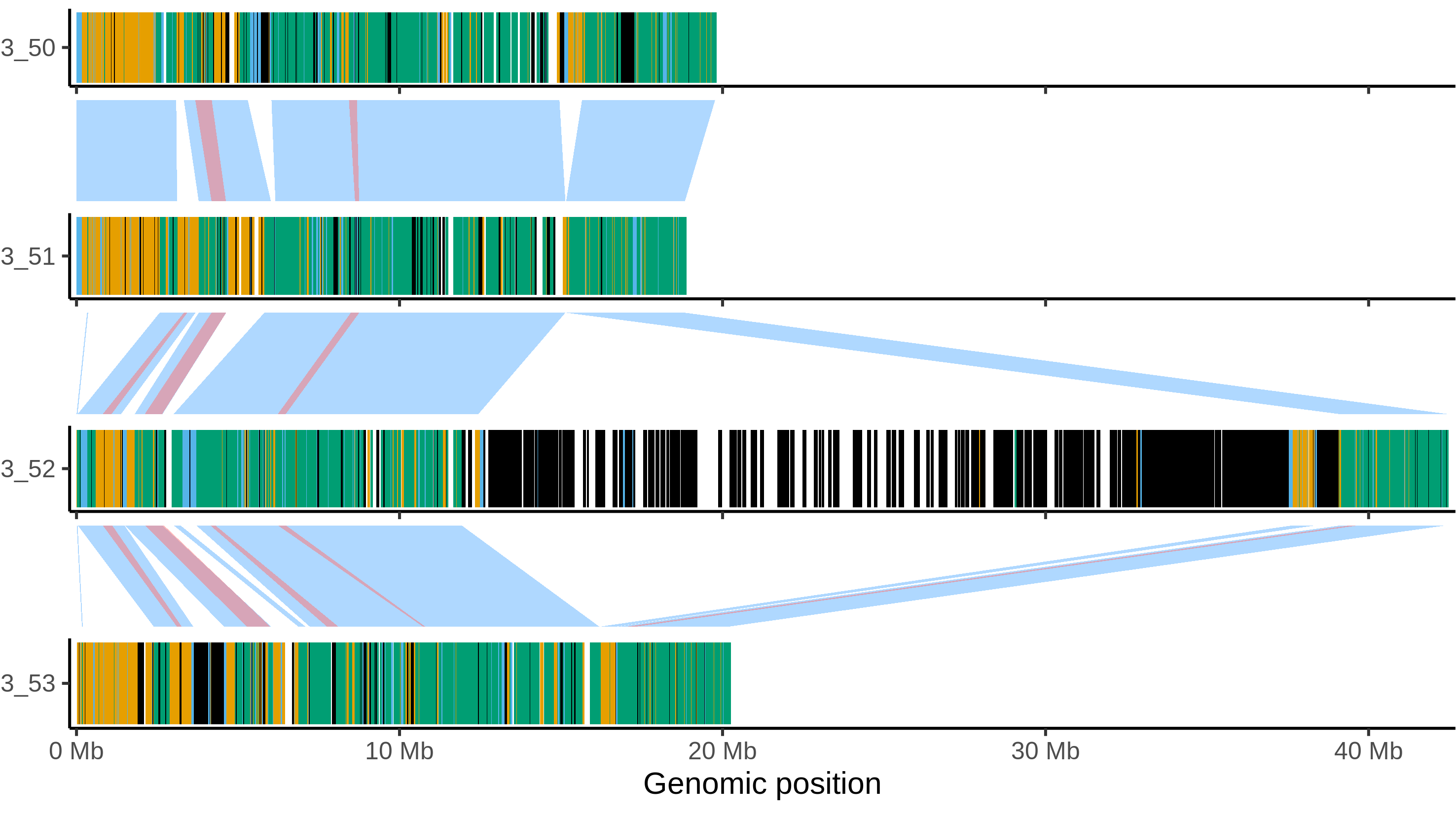

### Castle_russet_chr_6.png

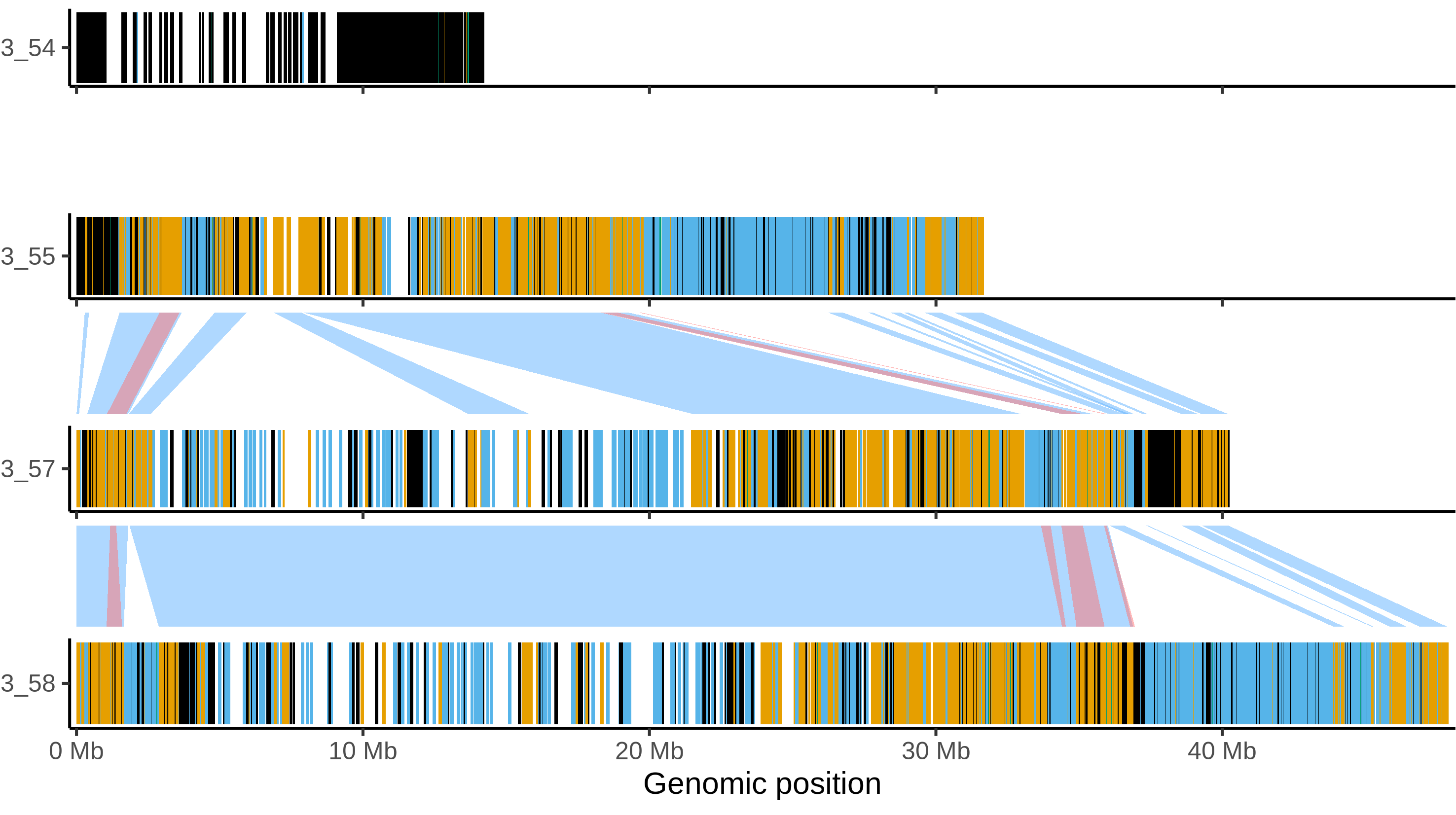

### Castle_russet_chr_7.png

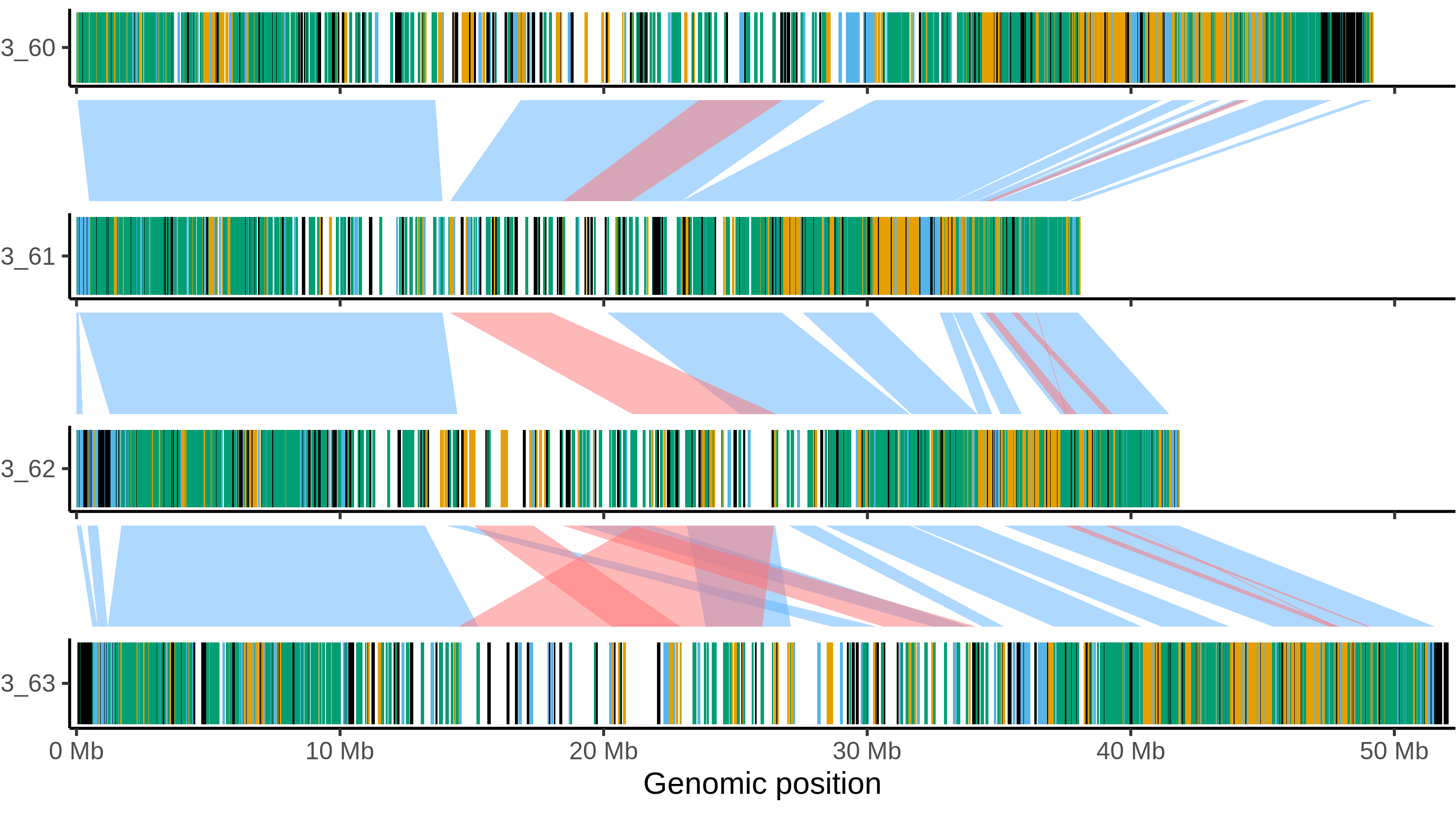

### Castle_russet_chr_8.png

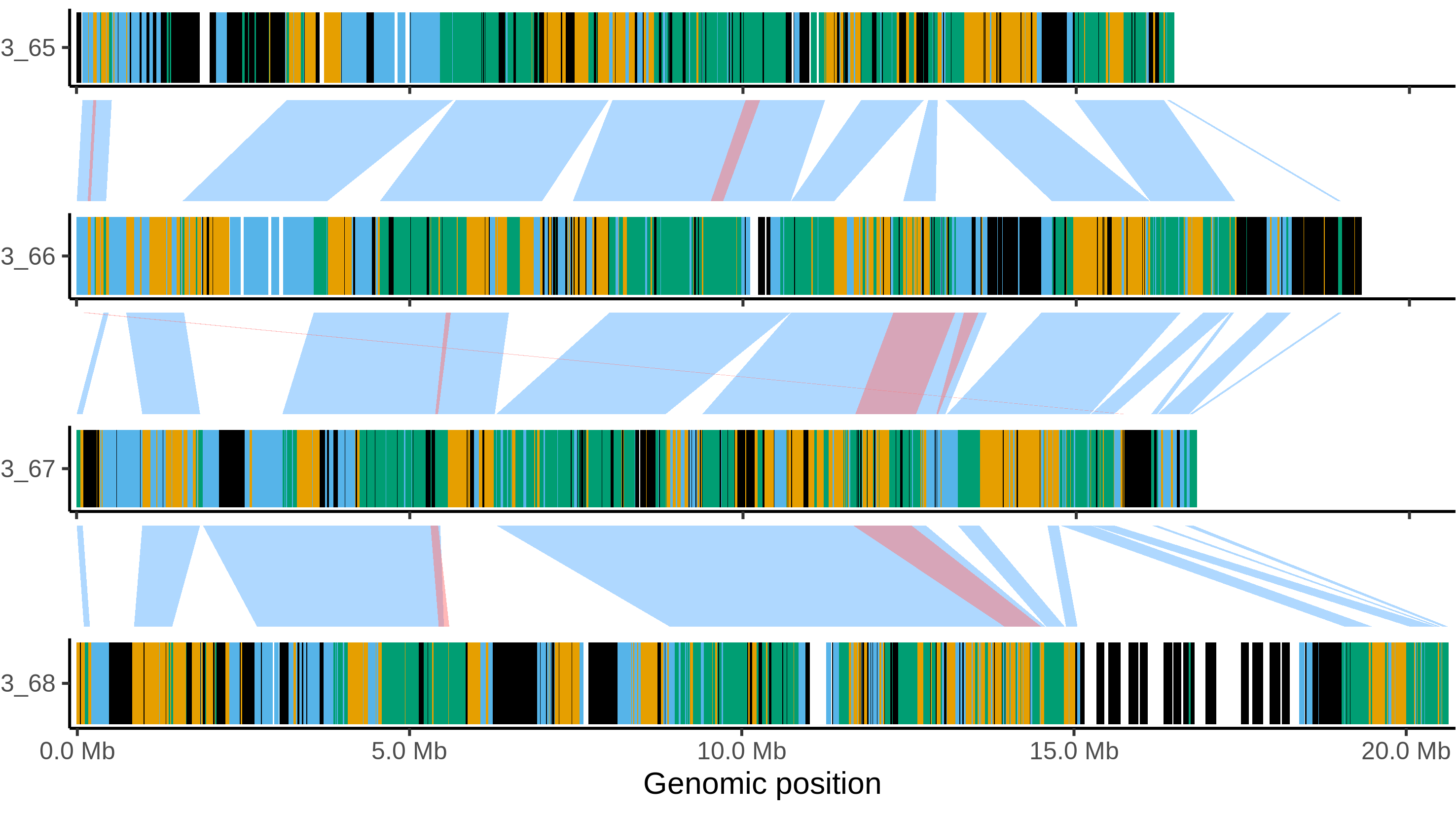

### Castle_russet_chr_9.png

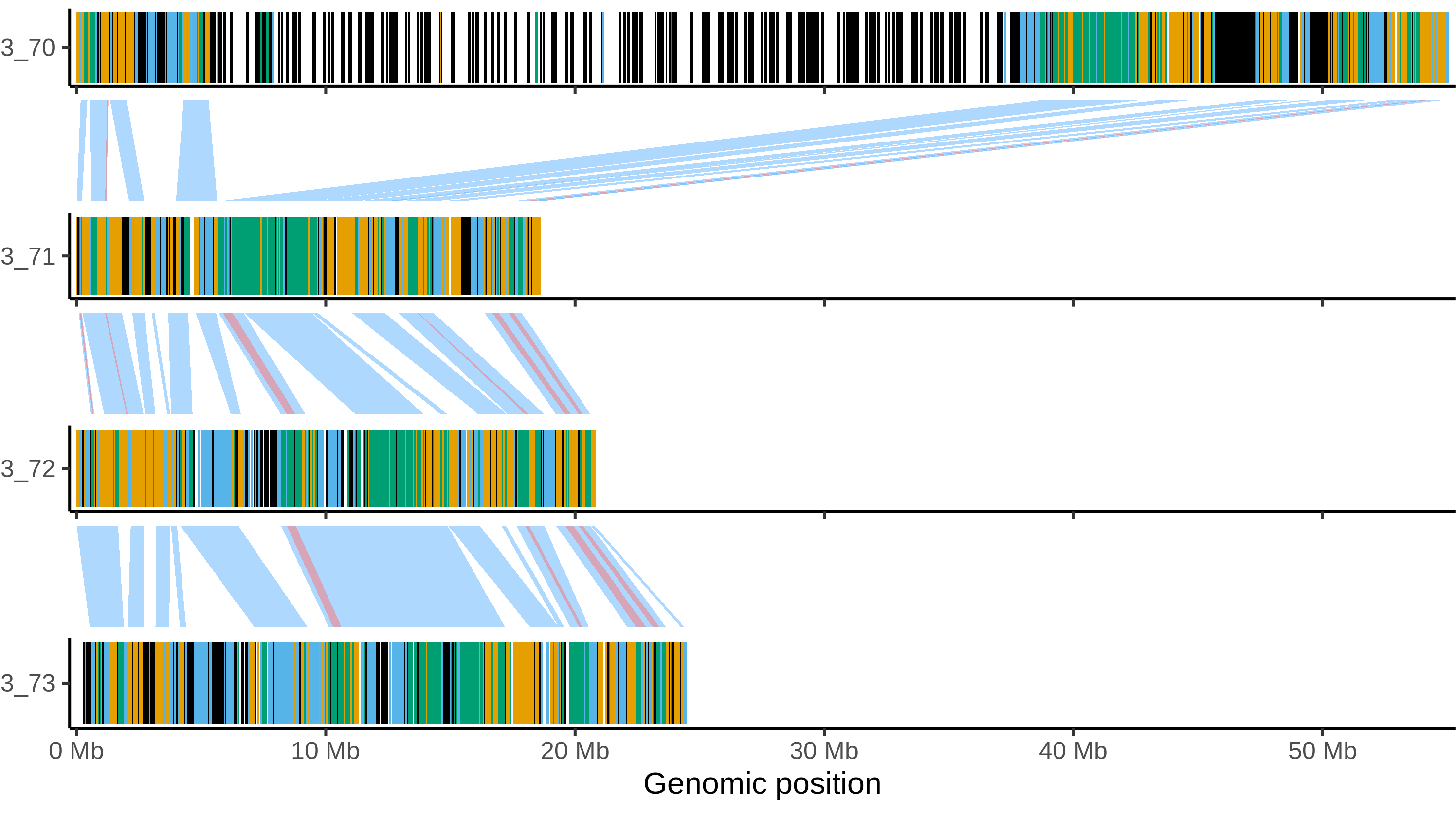

### Castle_russet_chr_10.png

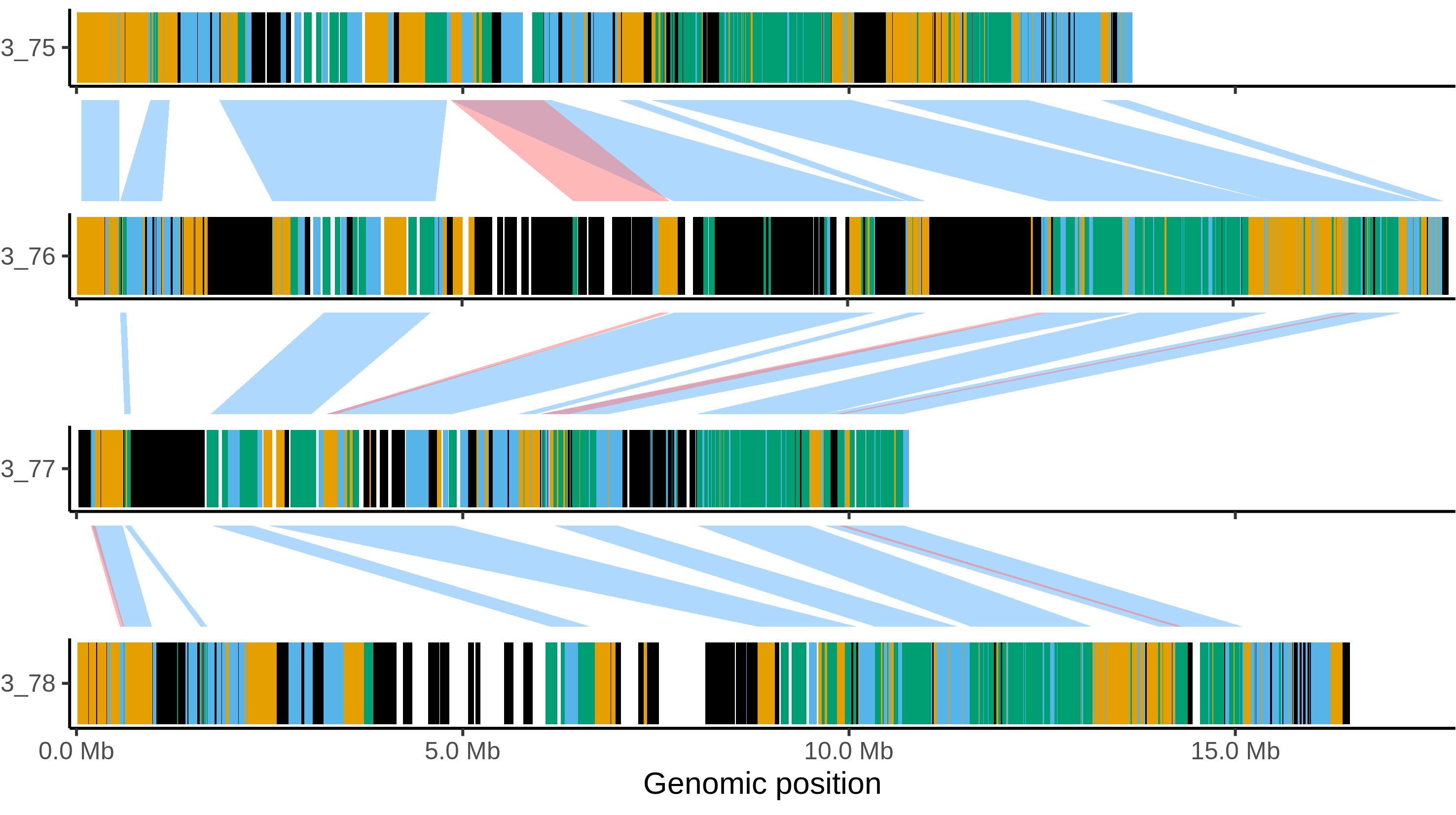

### Castle_russet_chr_11.png

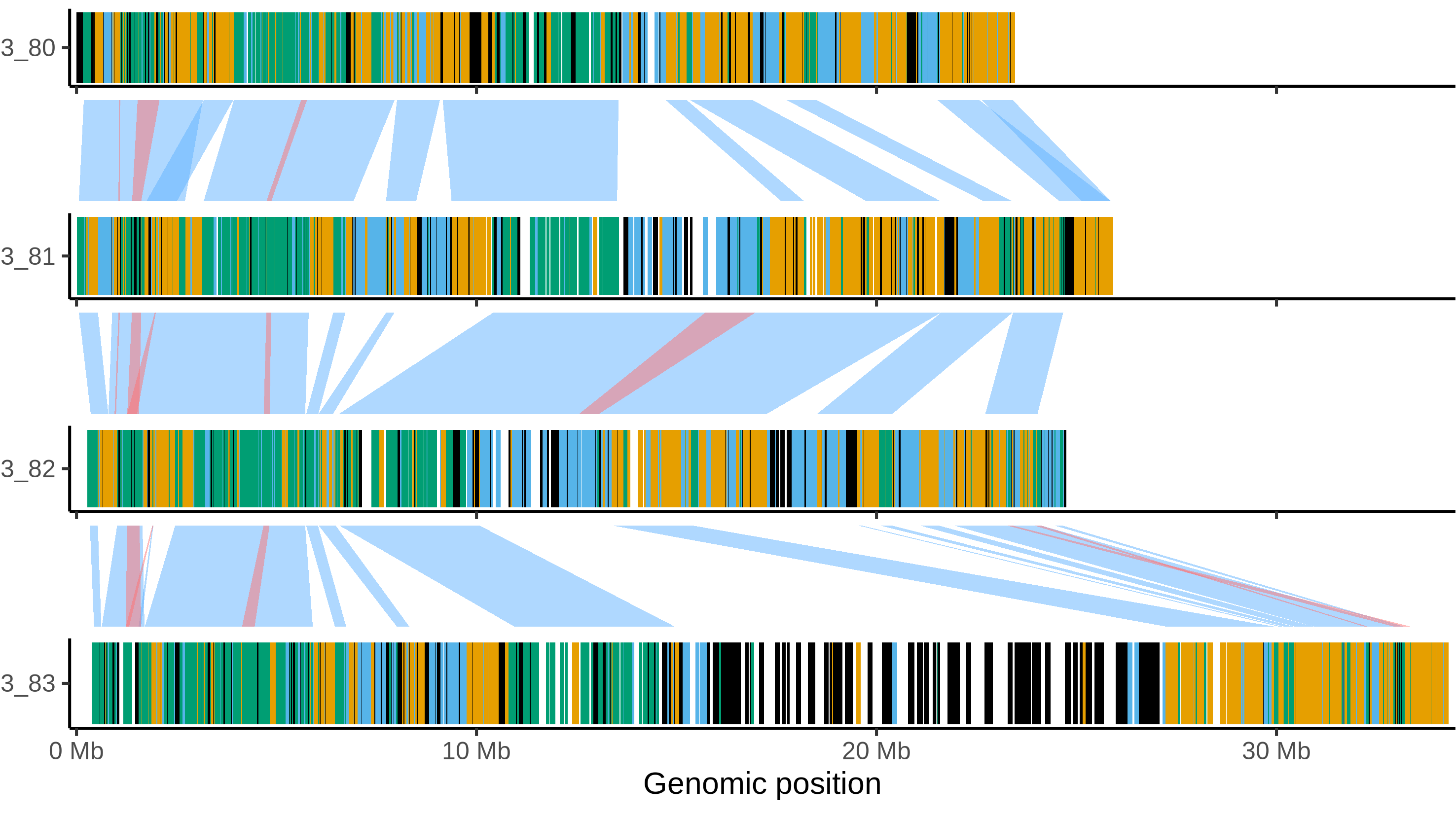

### Castle_russet_chr_12.png

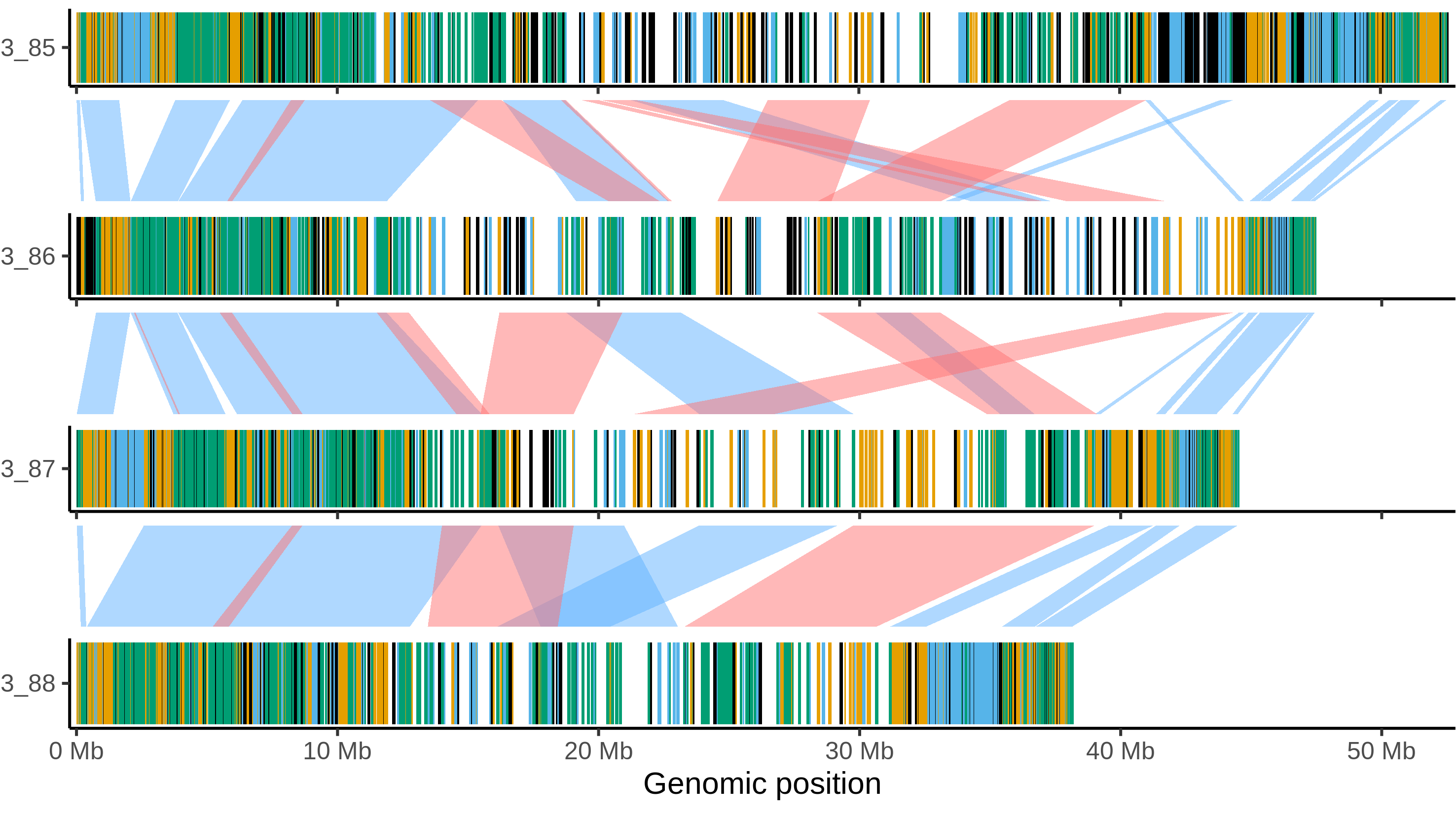

### mdomestica_gala_chr_1.png

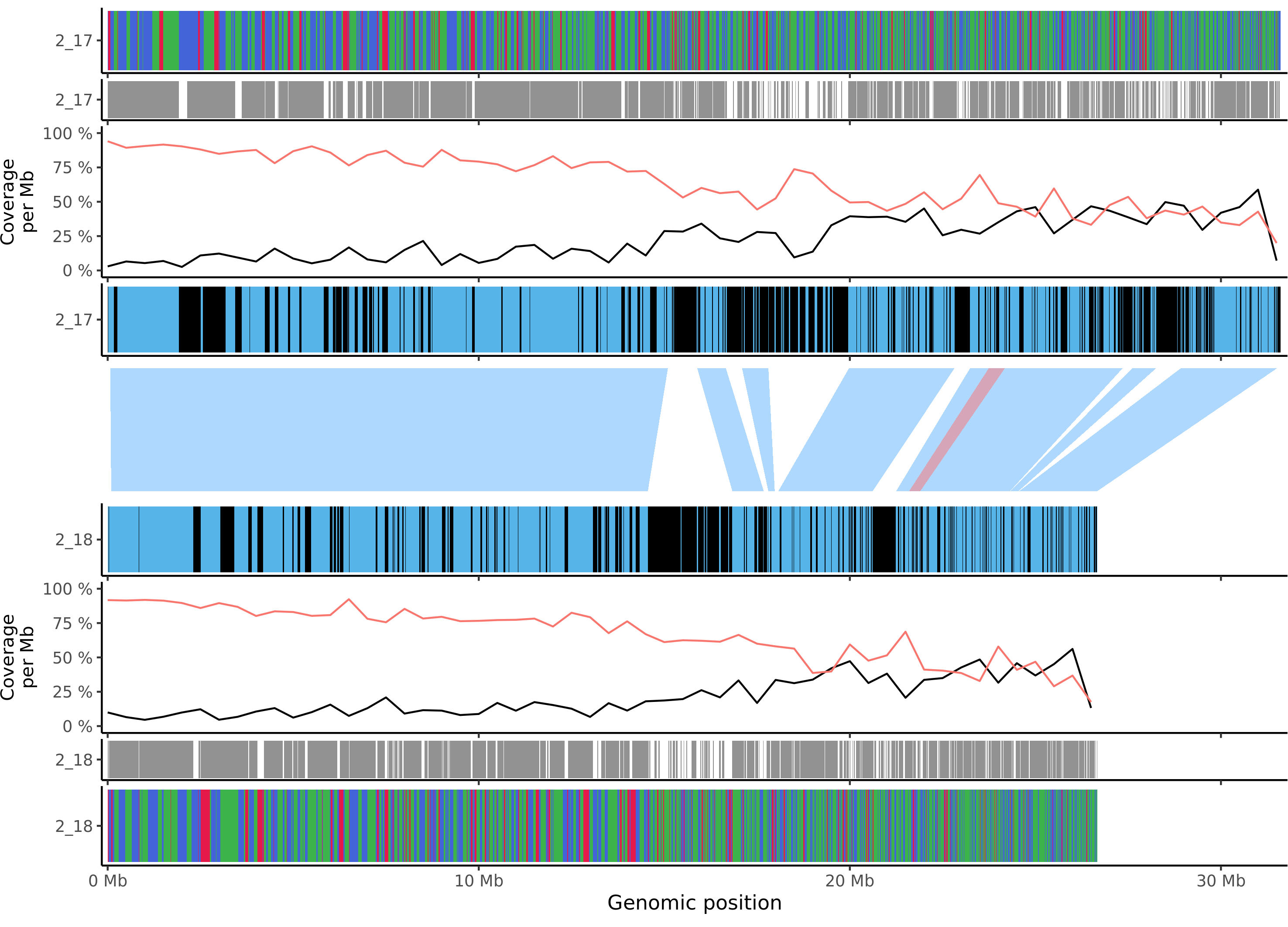

### mdomestica_gala_chr_2.png

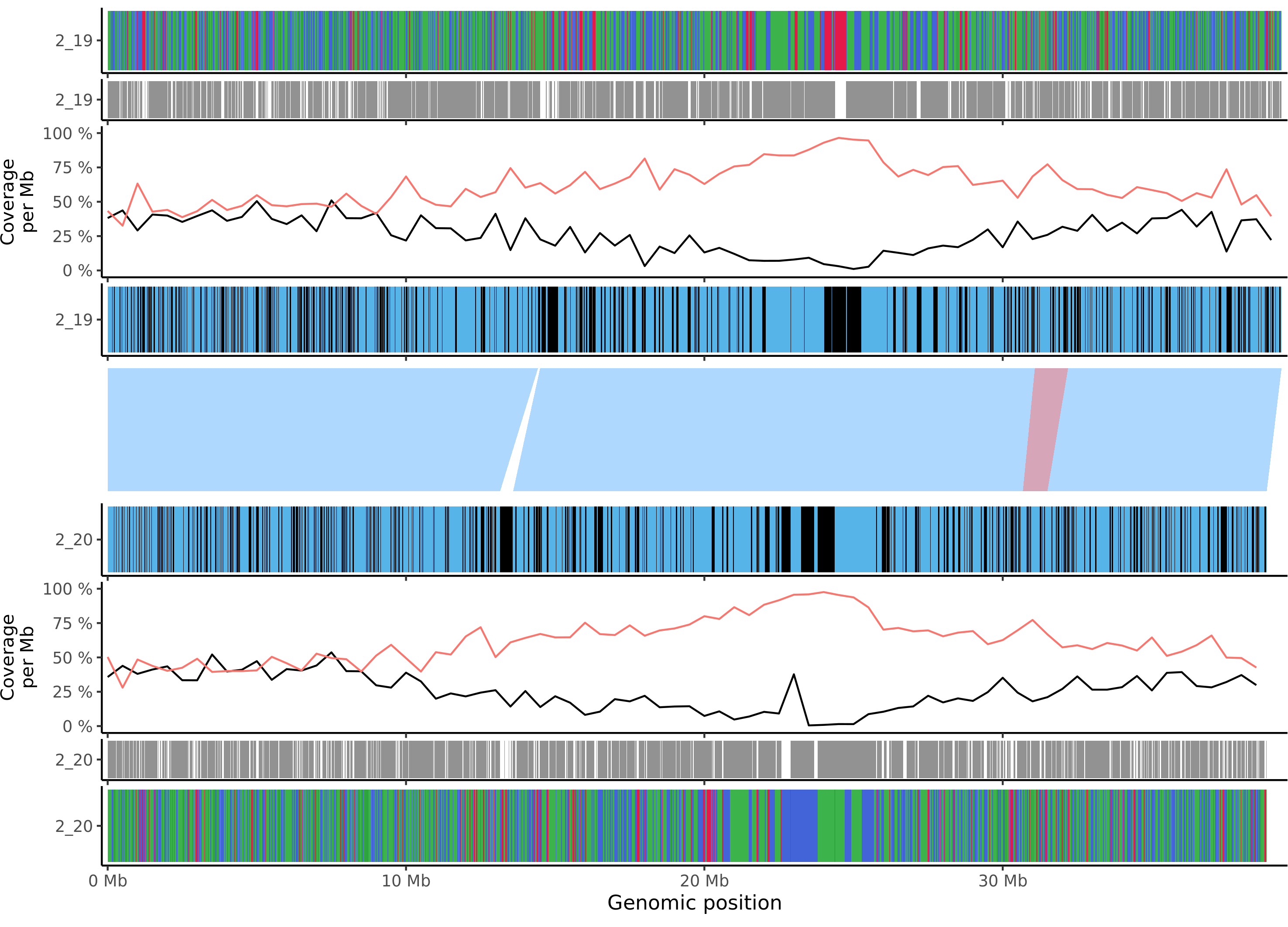

### mdomestica_gala_chr_3.png

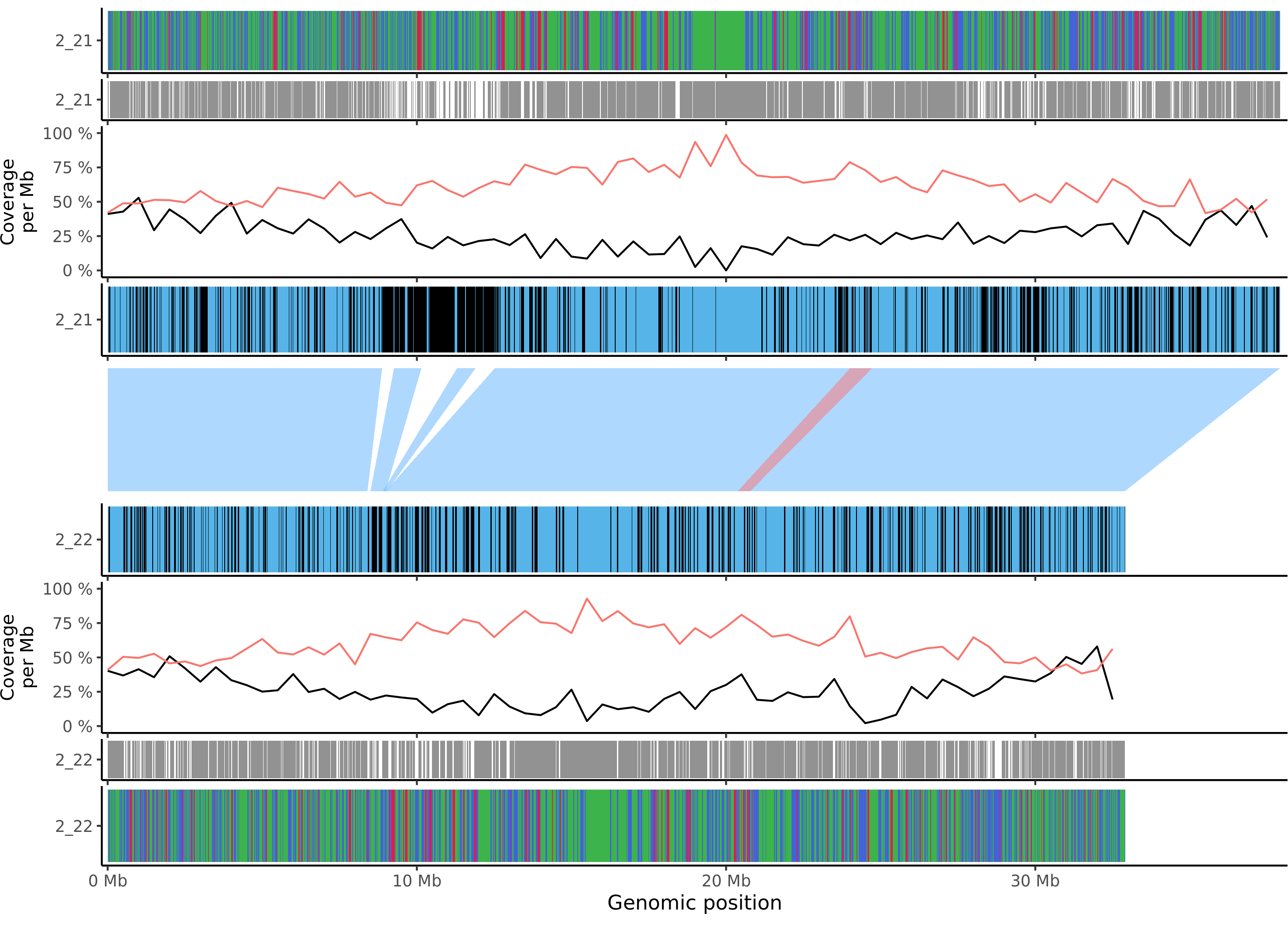

### mdomestica_gala_chr_4.png

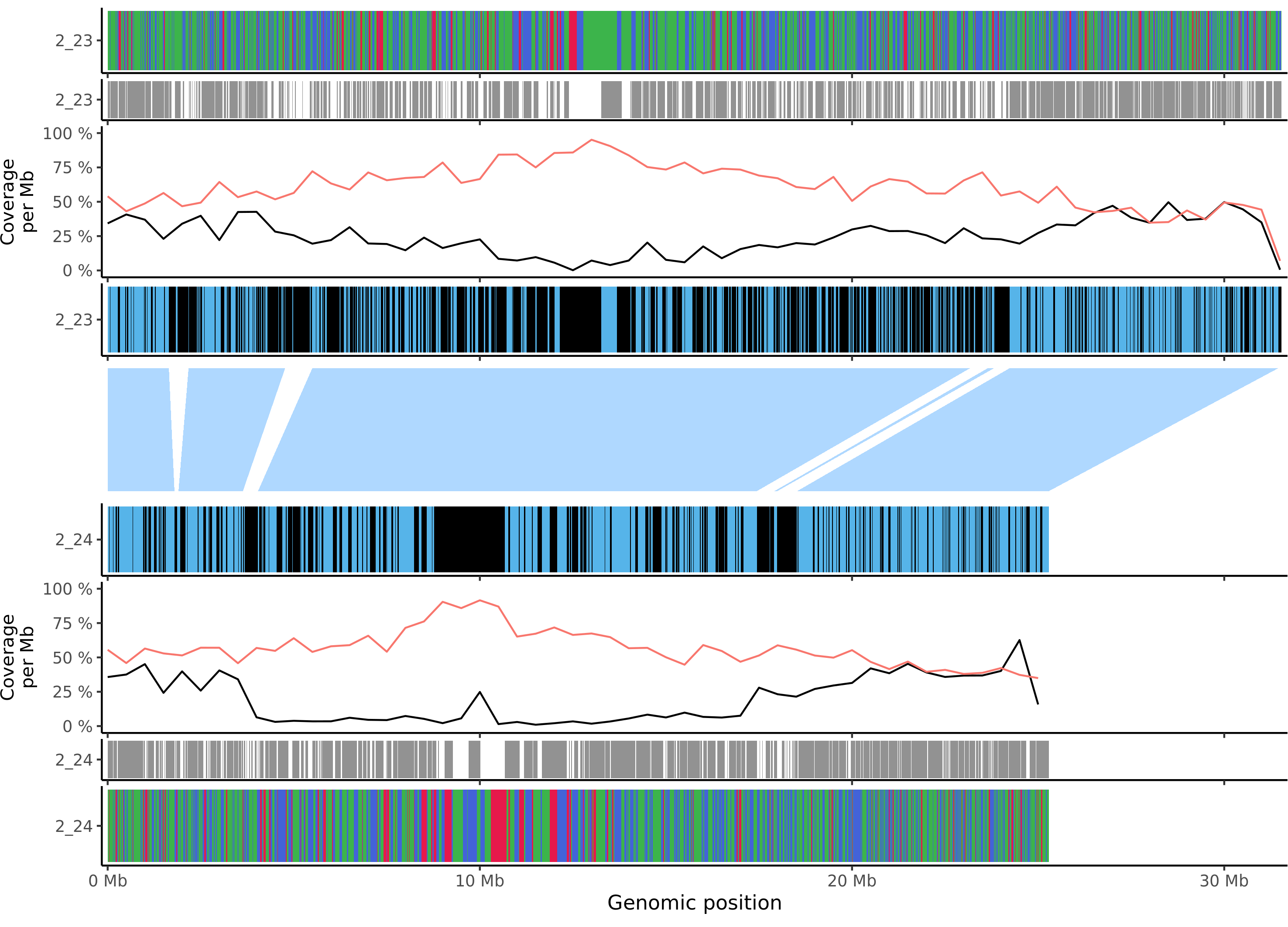

### mdomestica_gala_chr_5.png

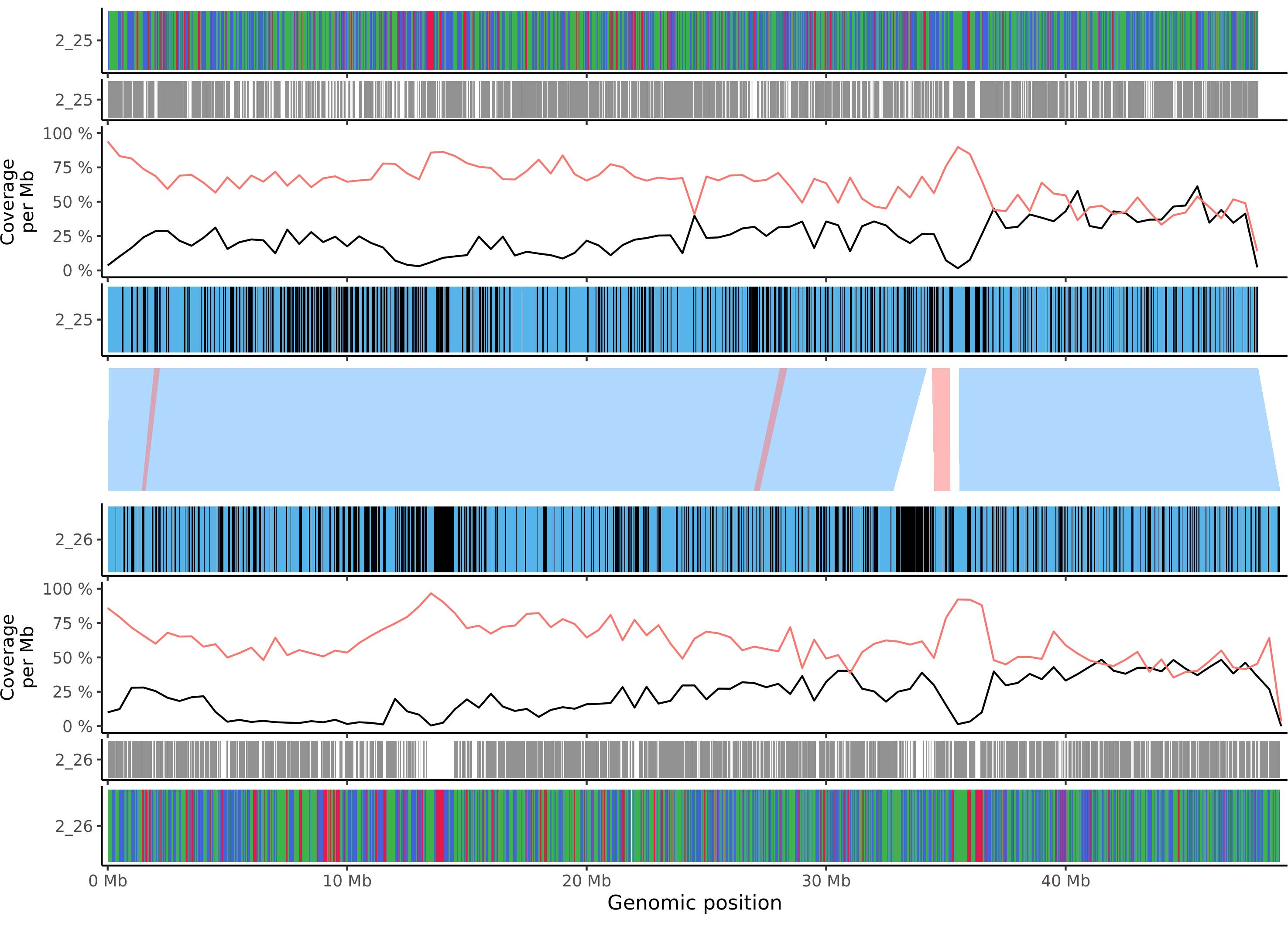

### mdomestica_gala_chr_6.png

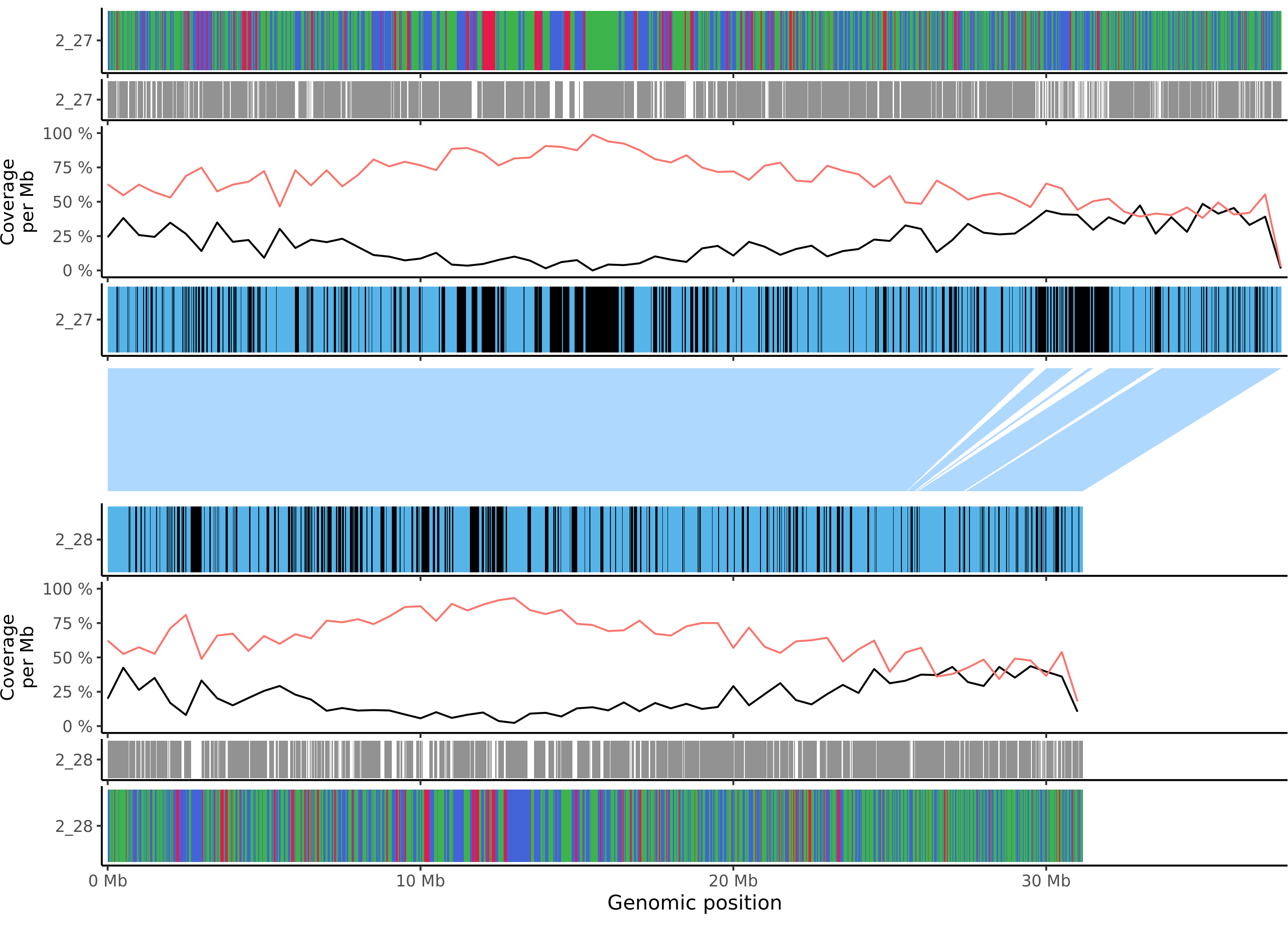

### mdomestica_gala_chr_7.png

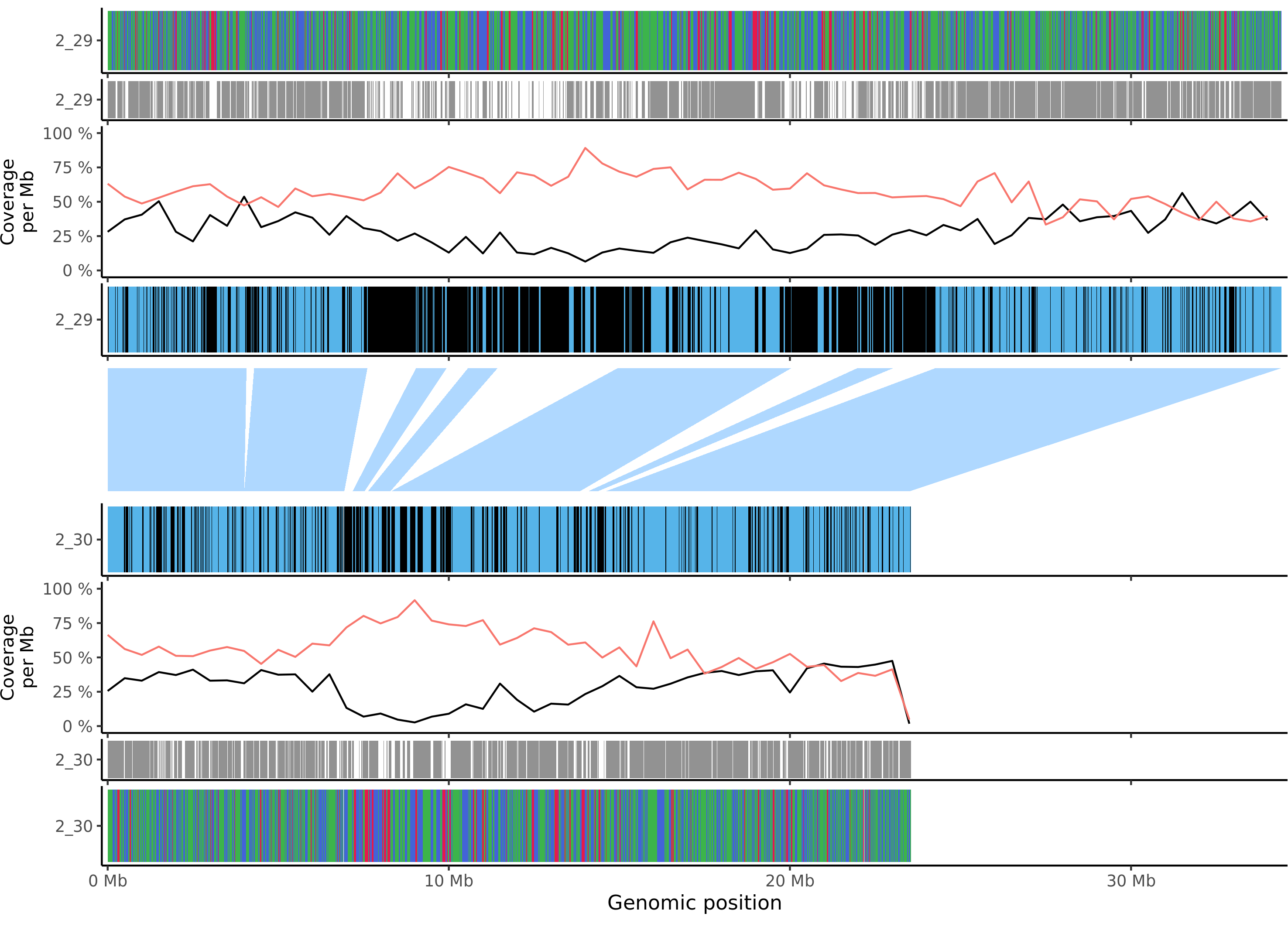

### mdomestica_gala_chr_8.png

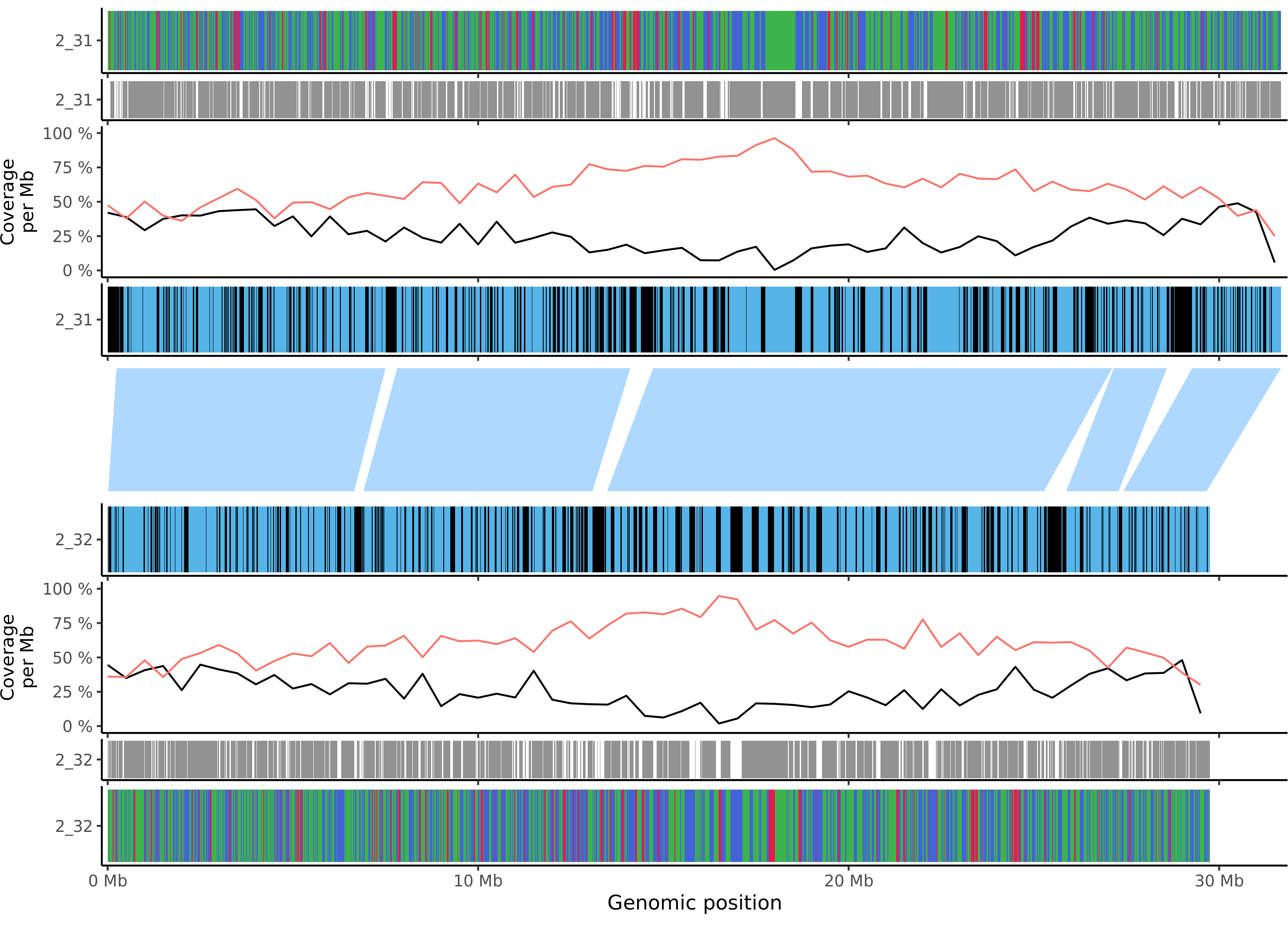

### mdomestica_gala_chr_9.png

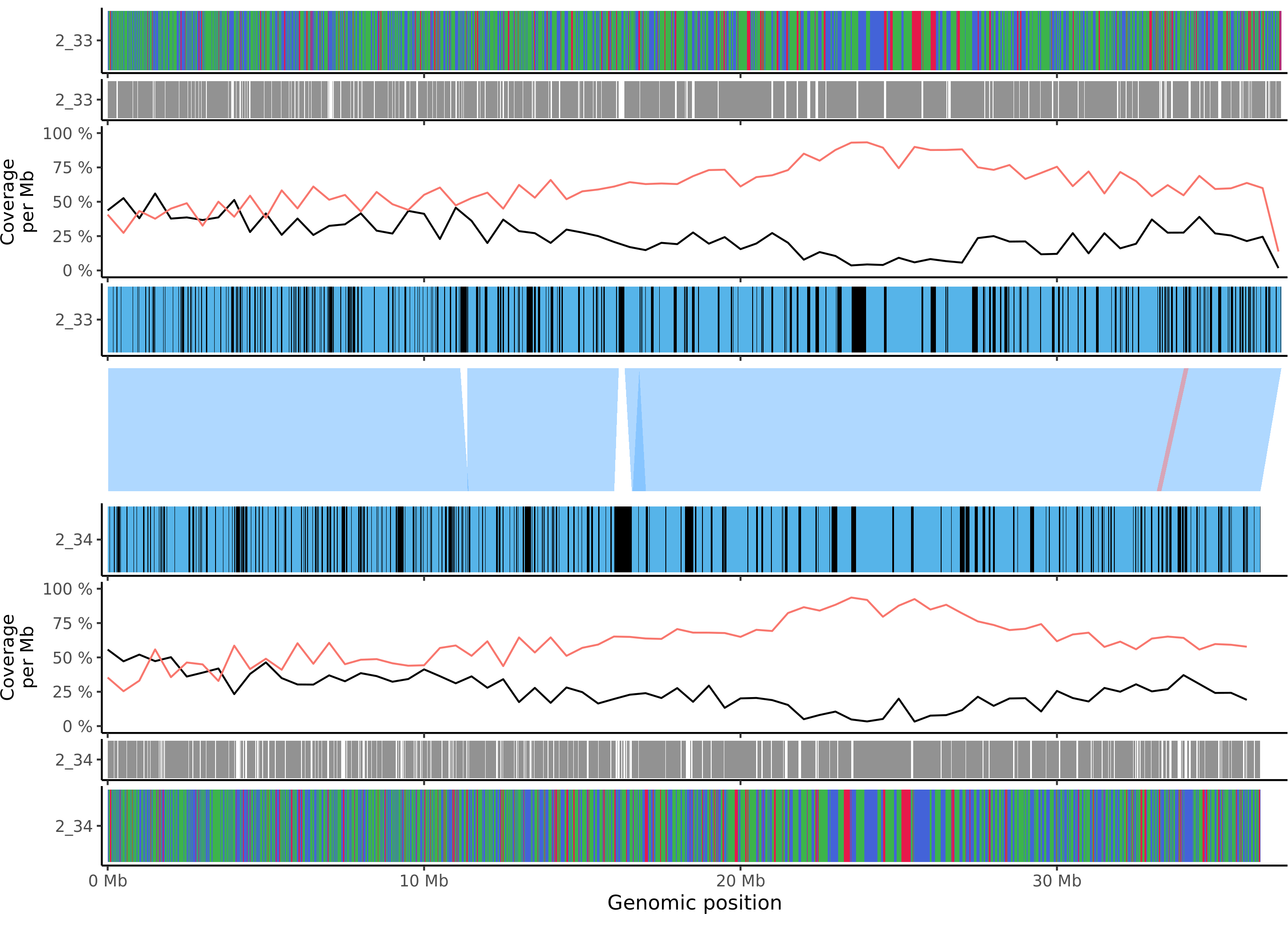

### mdomestica_gala_chr_10.png

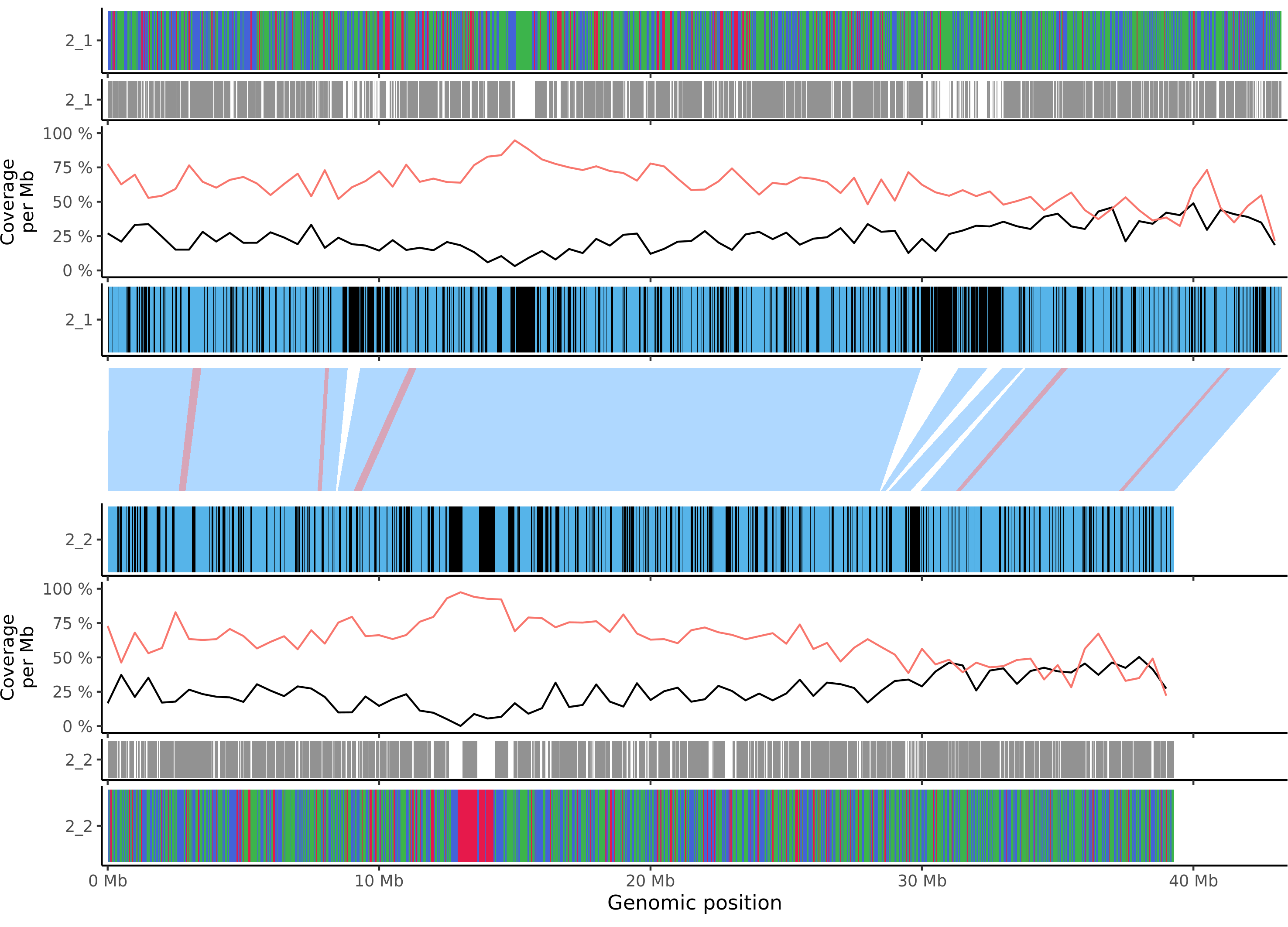

### mdomestica_gala_chr_11.png

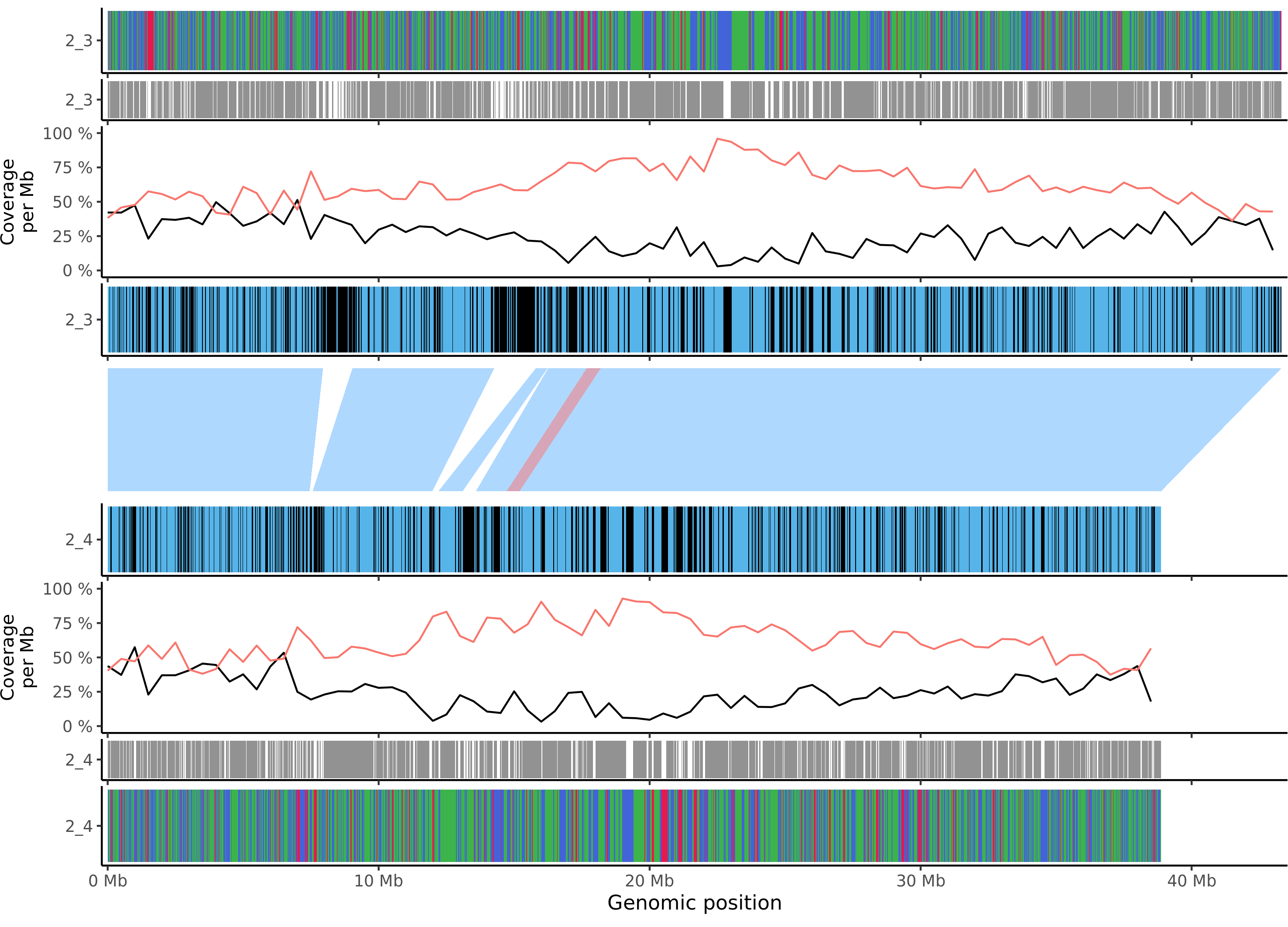

### mdomestica_gala_chr_12.png

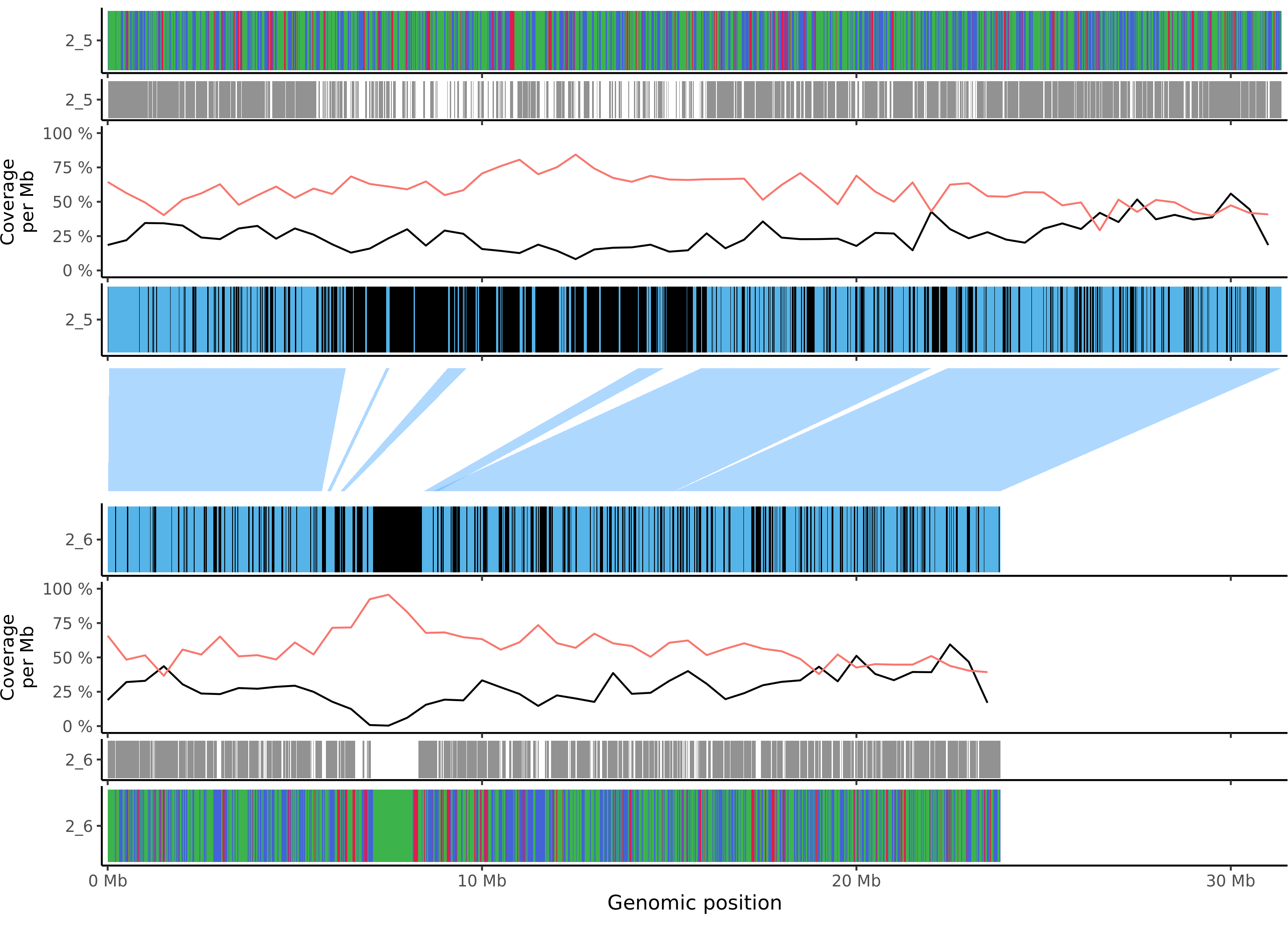

### mdomestica_gala_chr_13.png

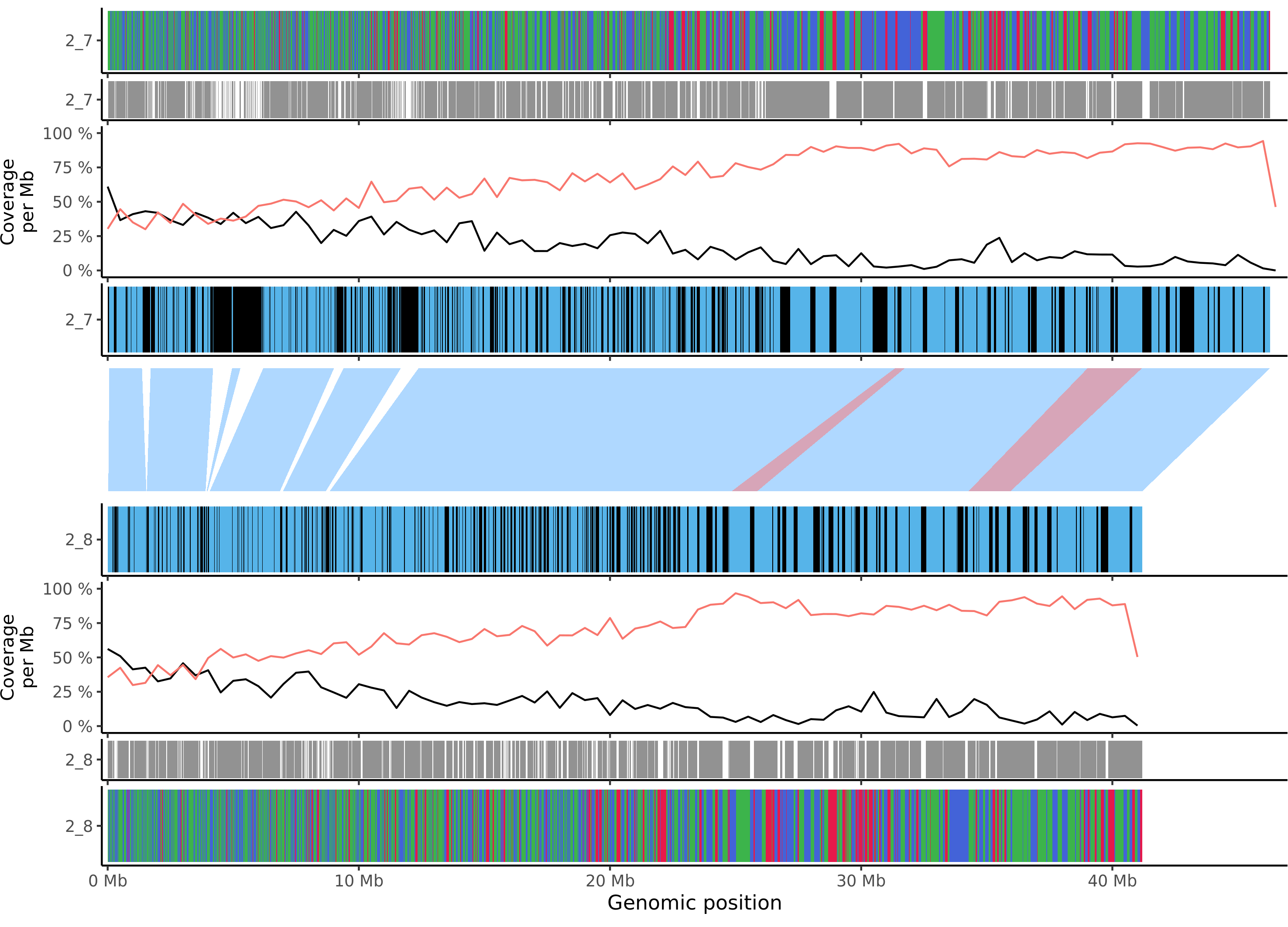

### mdomestica_gala_chr_14.png

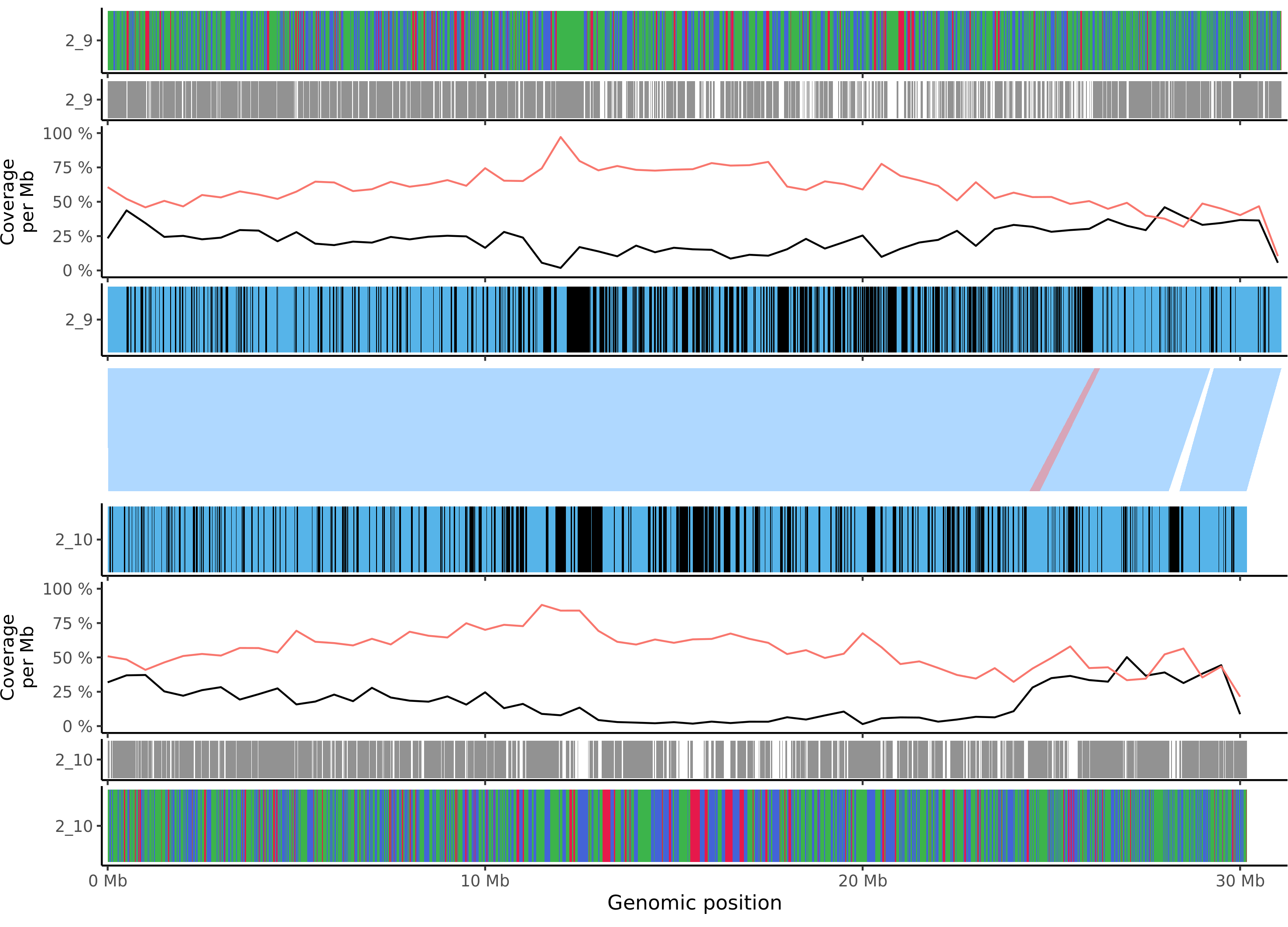

### mdomestica_gala_chr_15.png

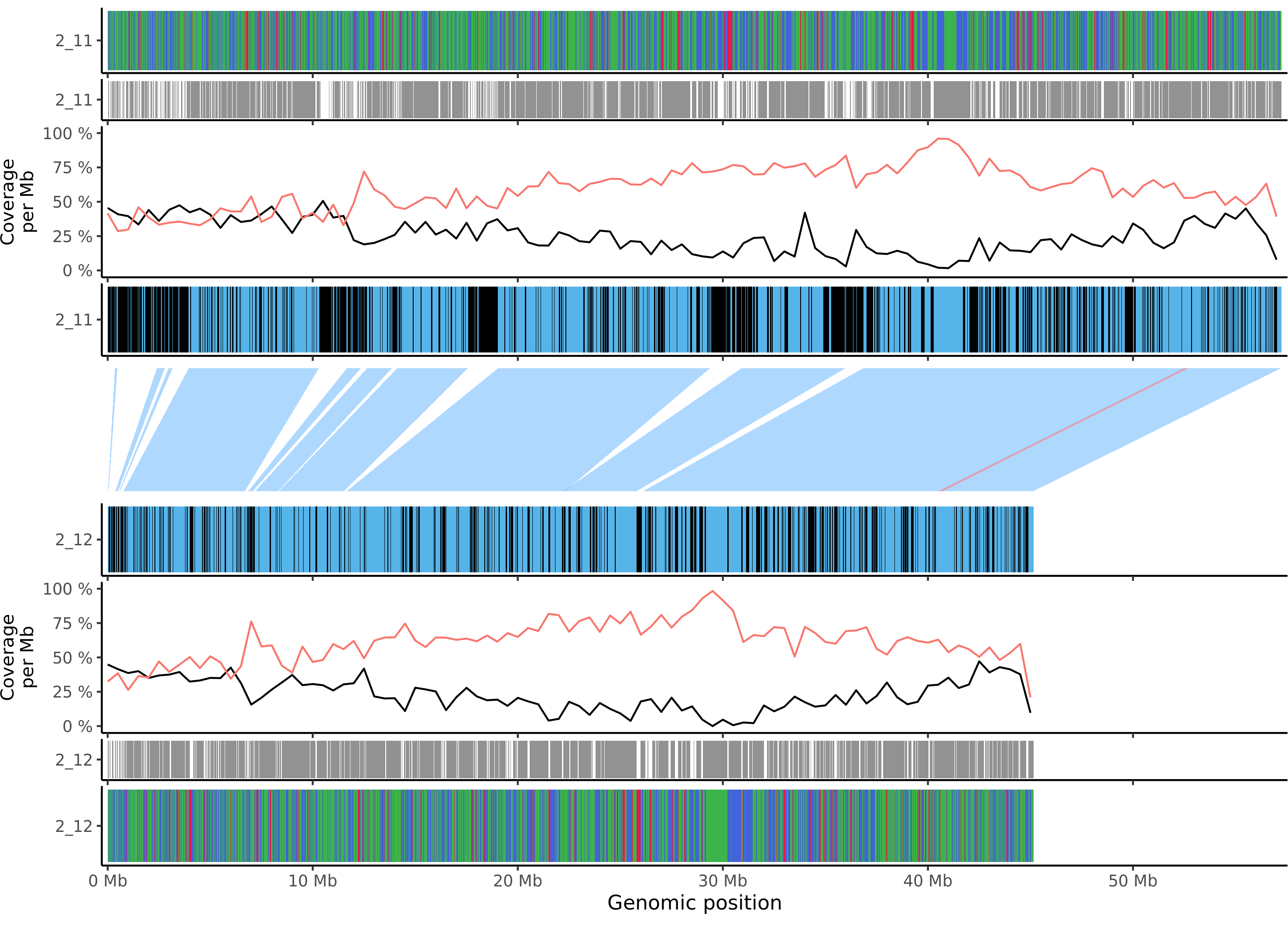

### mdomestica_gala_chr_16.png

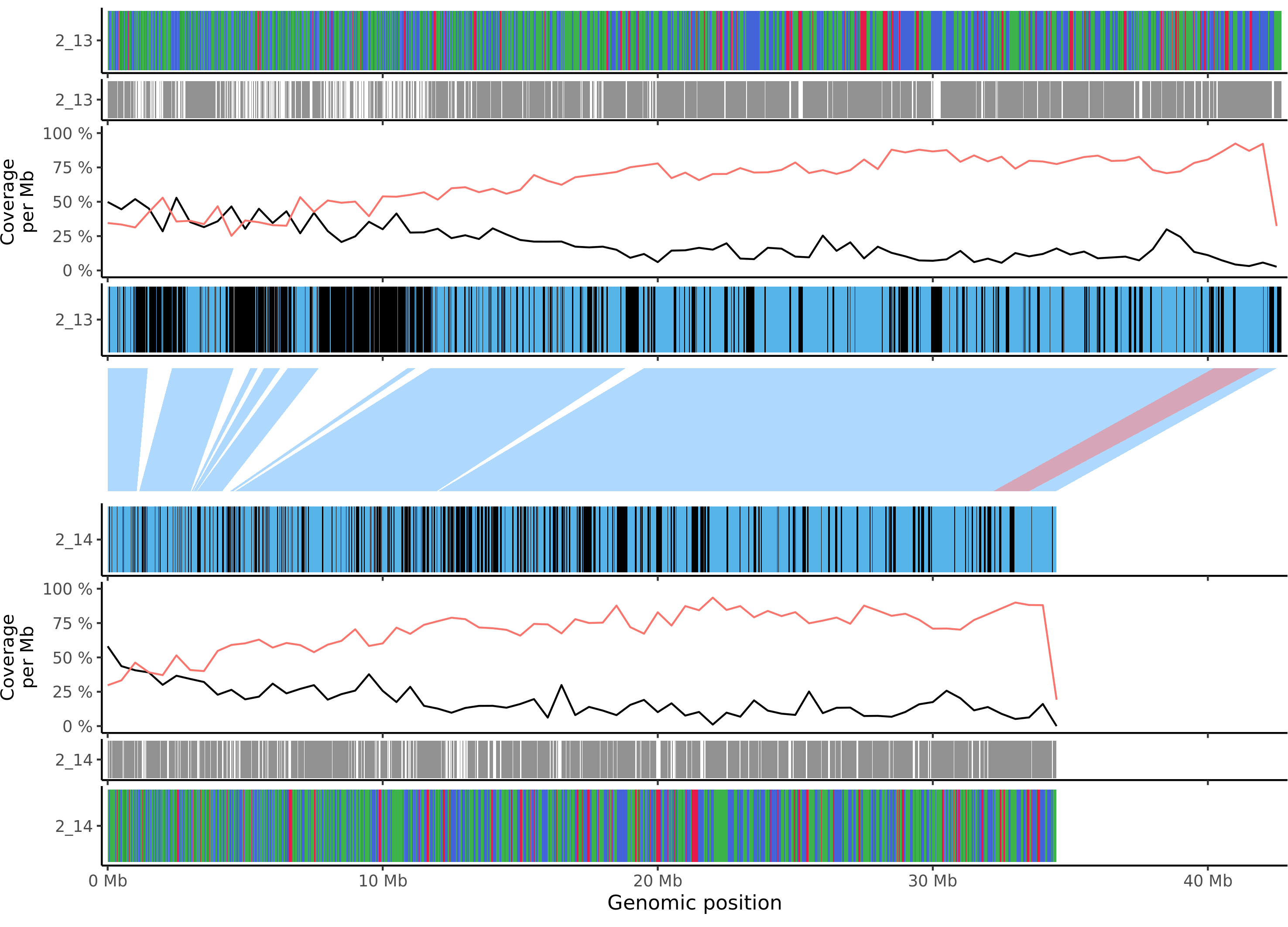

### mdomestica_gala_chr_17.png

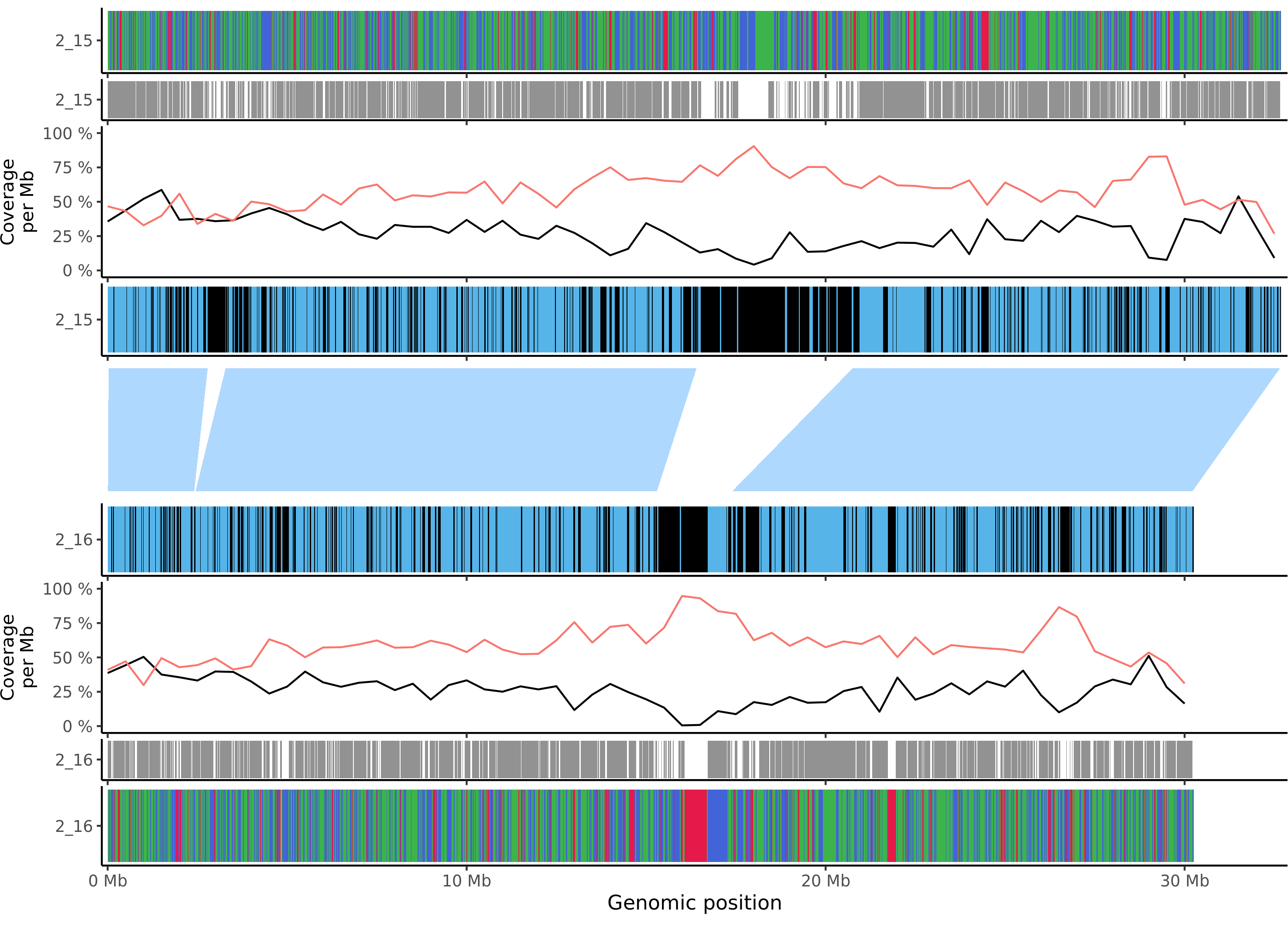

### msieversii_chr_1.png

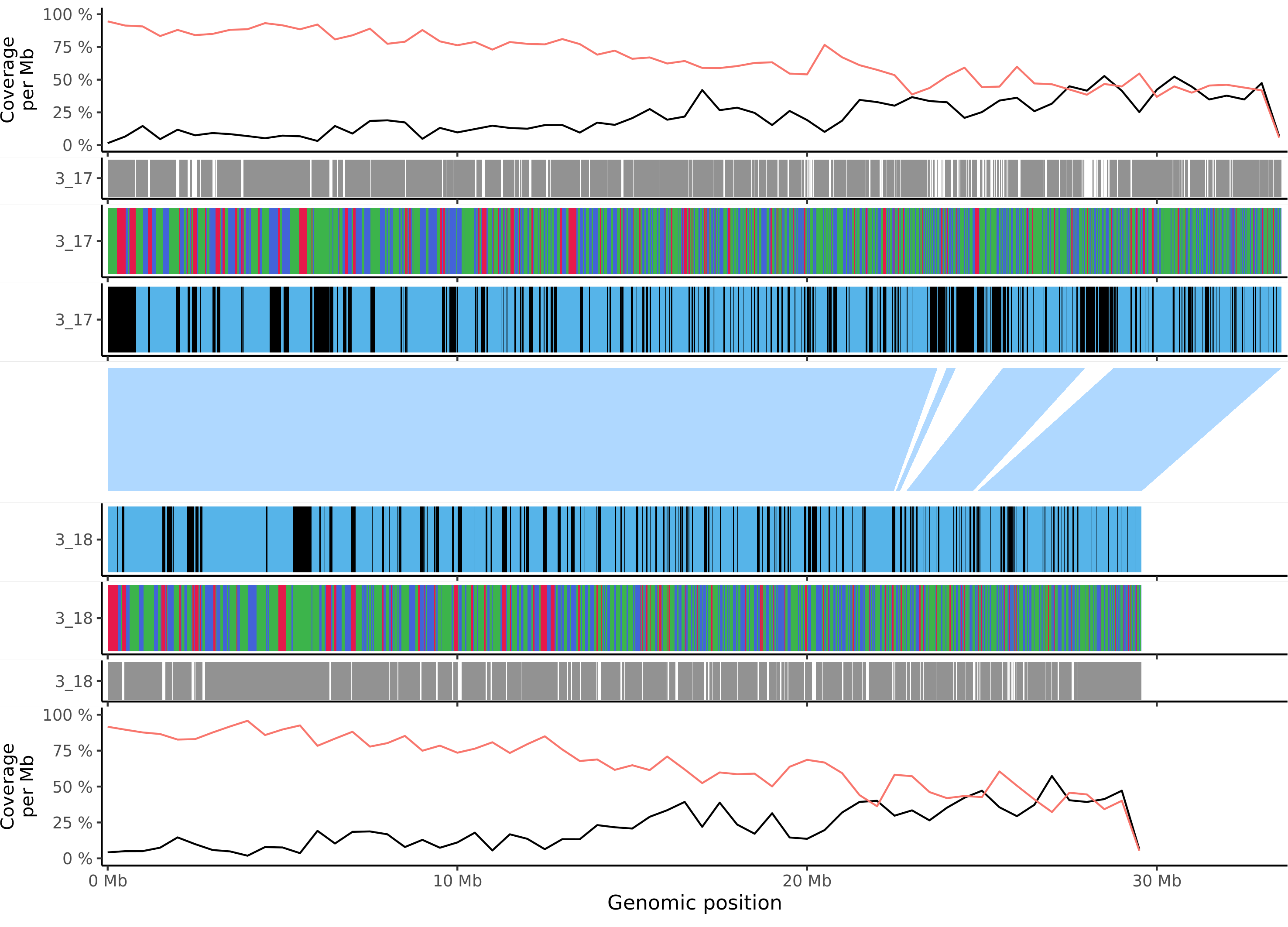

### msieversii_chr_2.png

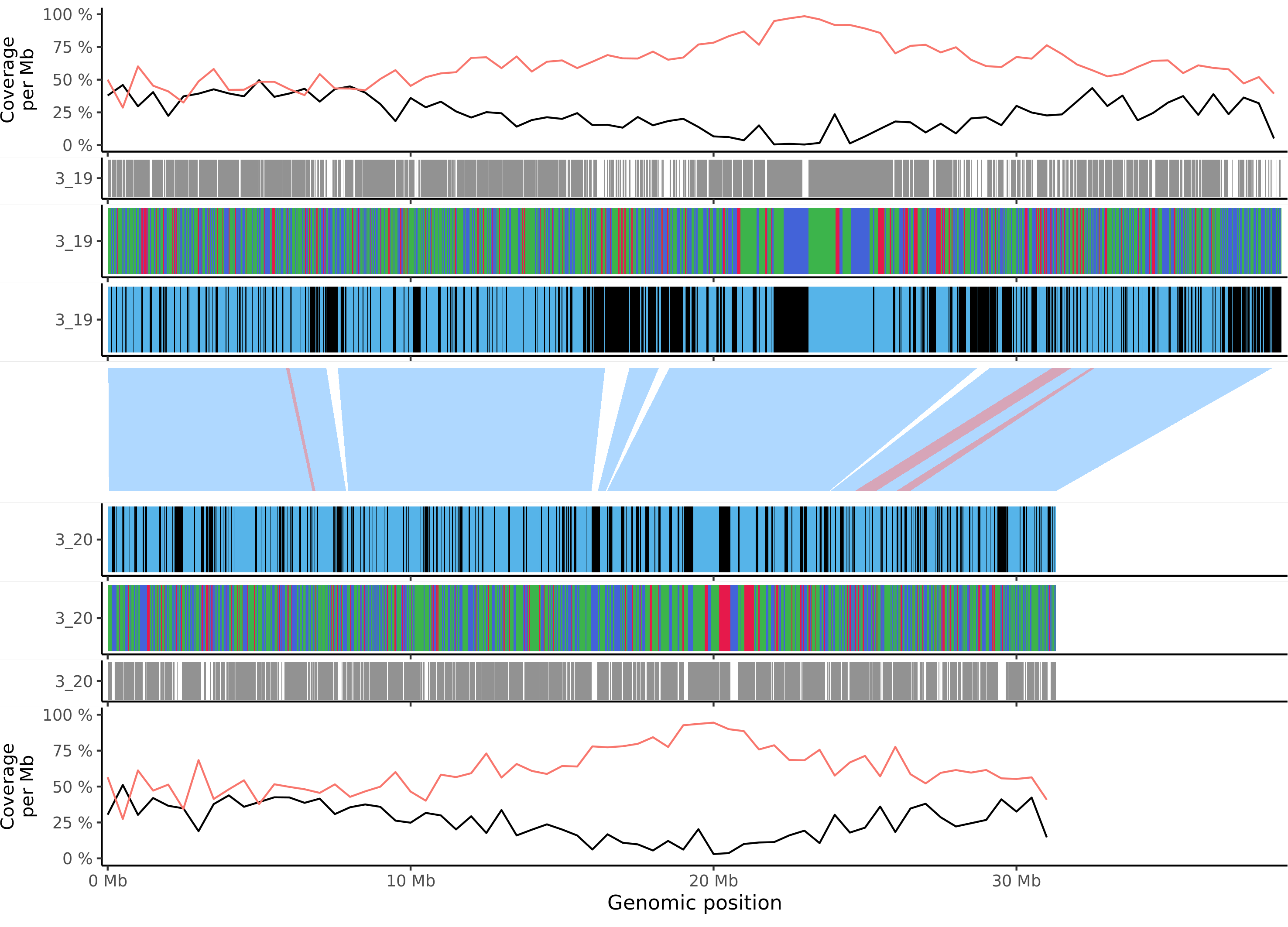

### msieversii_chr_3.png

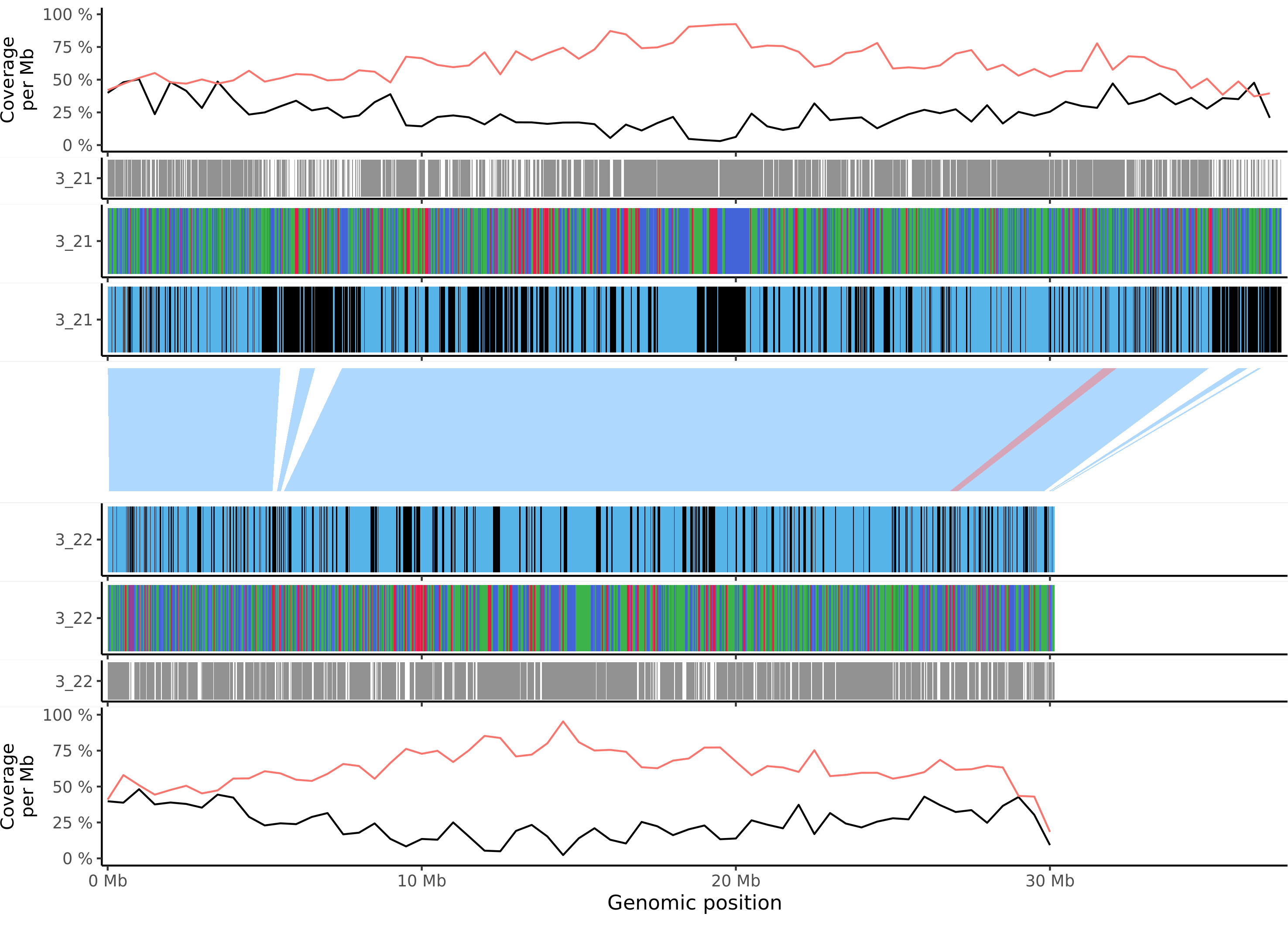

### msieversii_chr_4.png

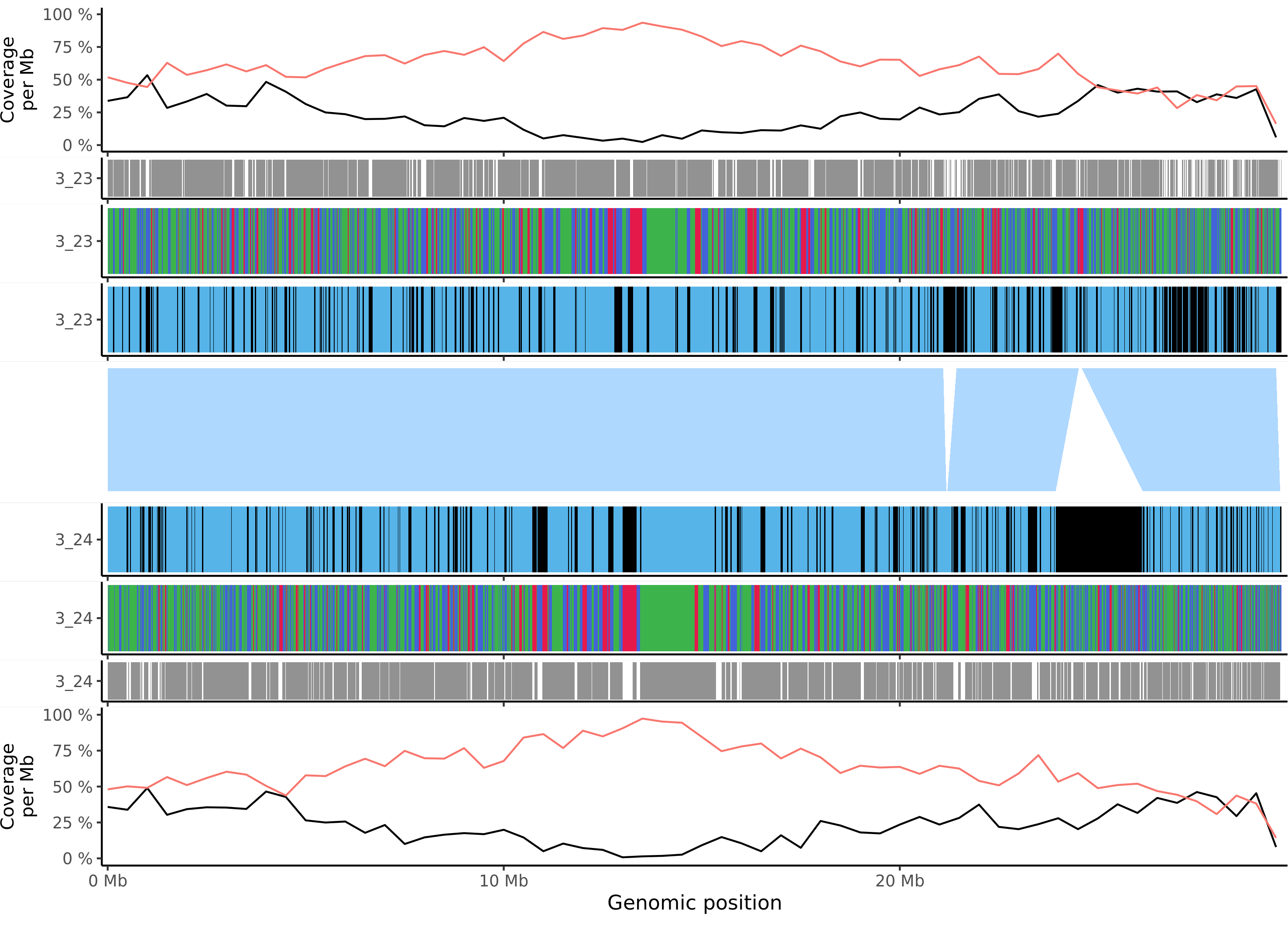

### msieversii_chr_5.png

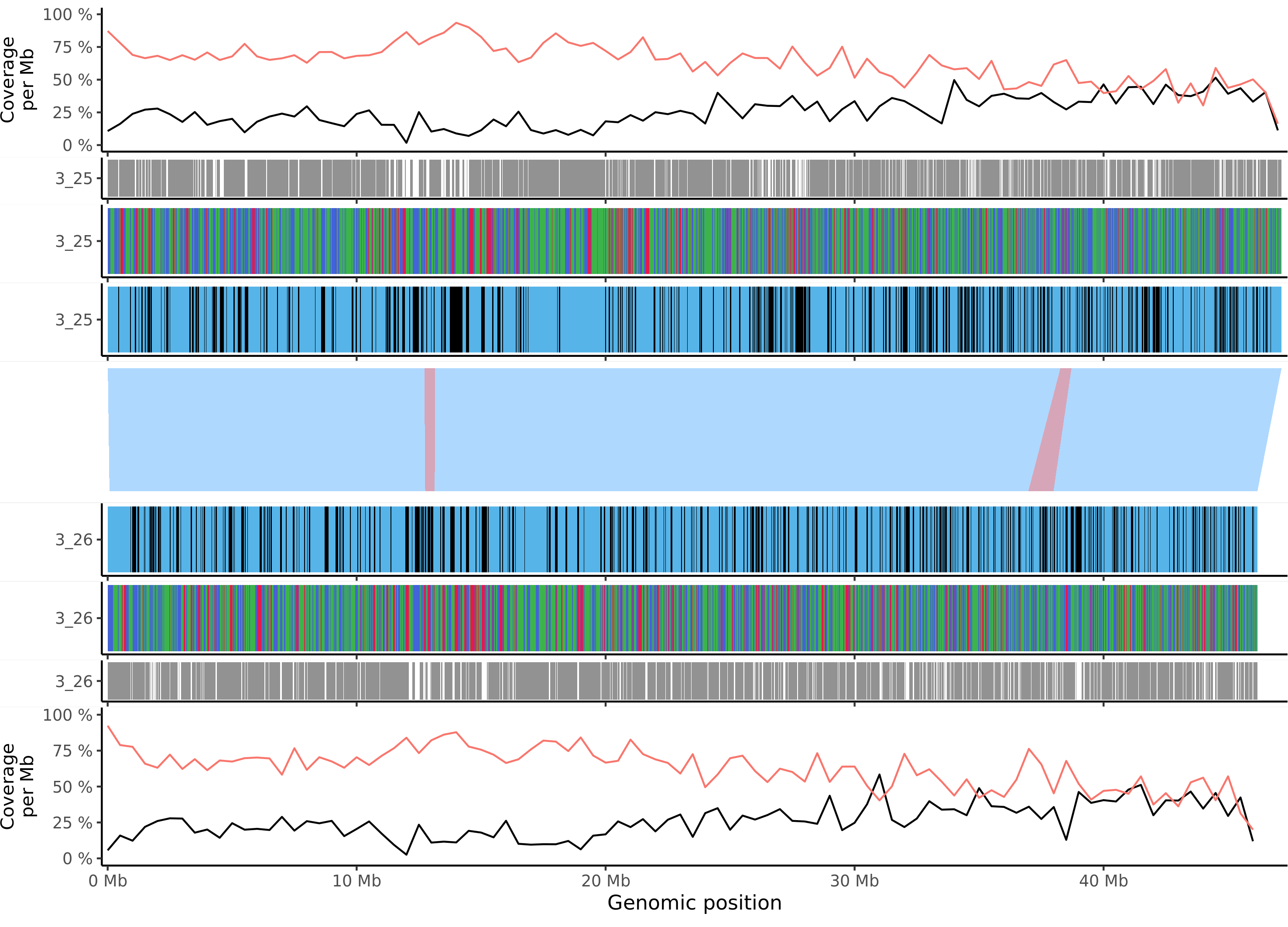

### msieversii_chr_6.png

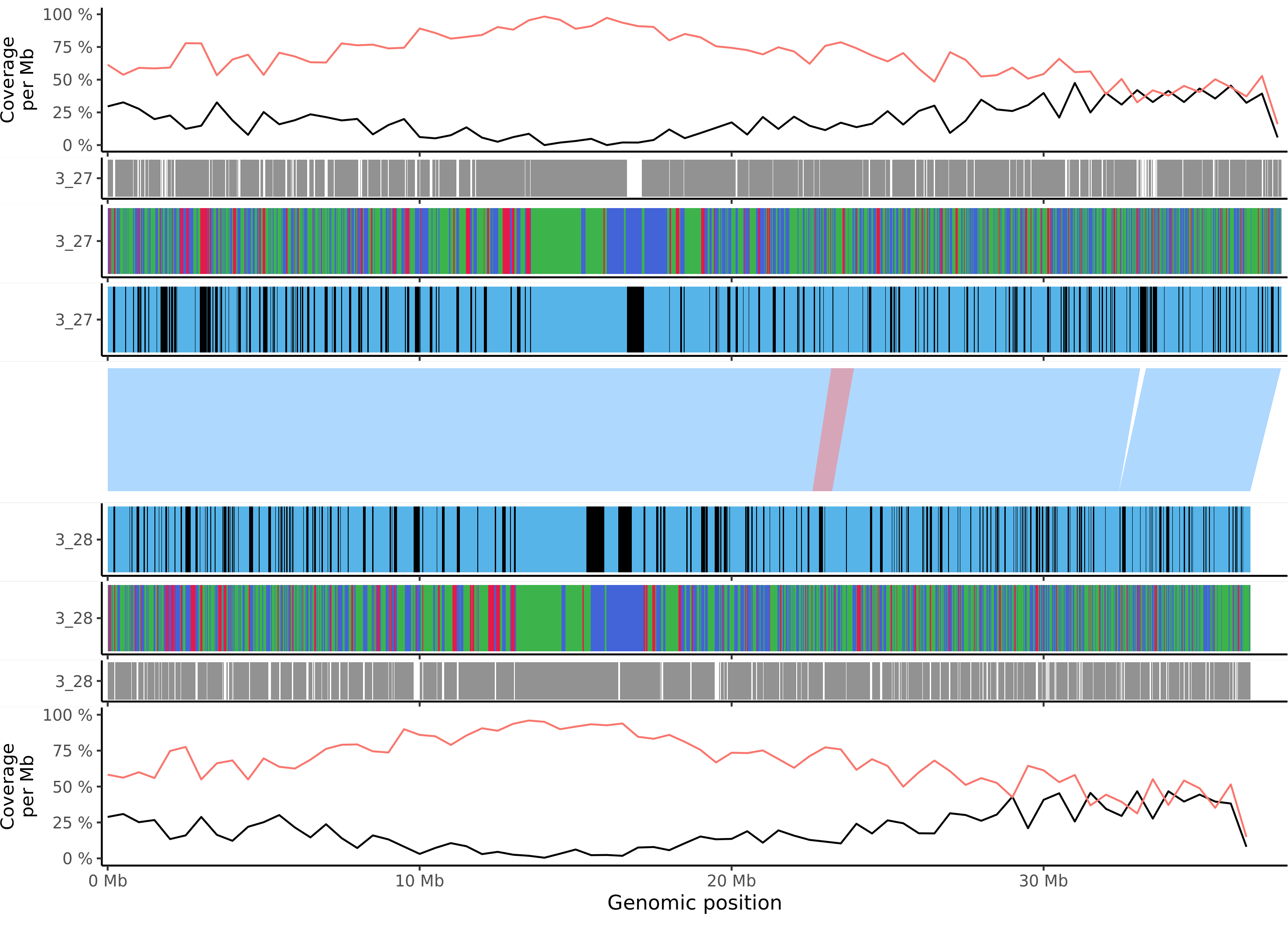

### msieversii_chr_7.png

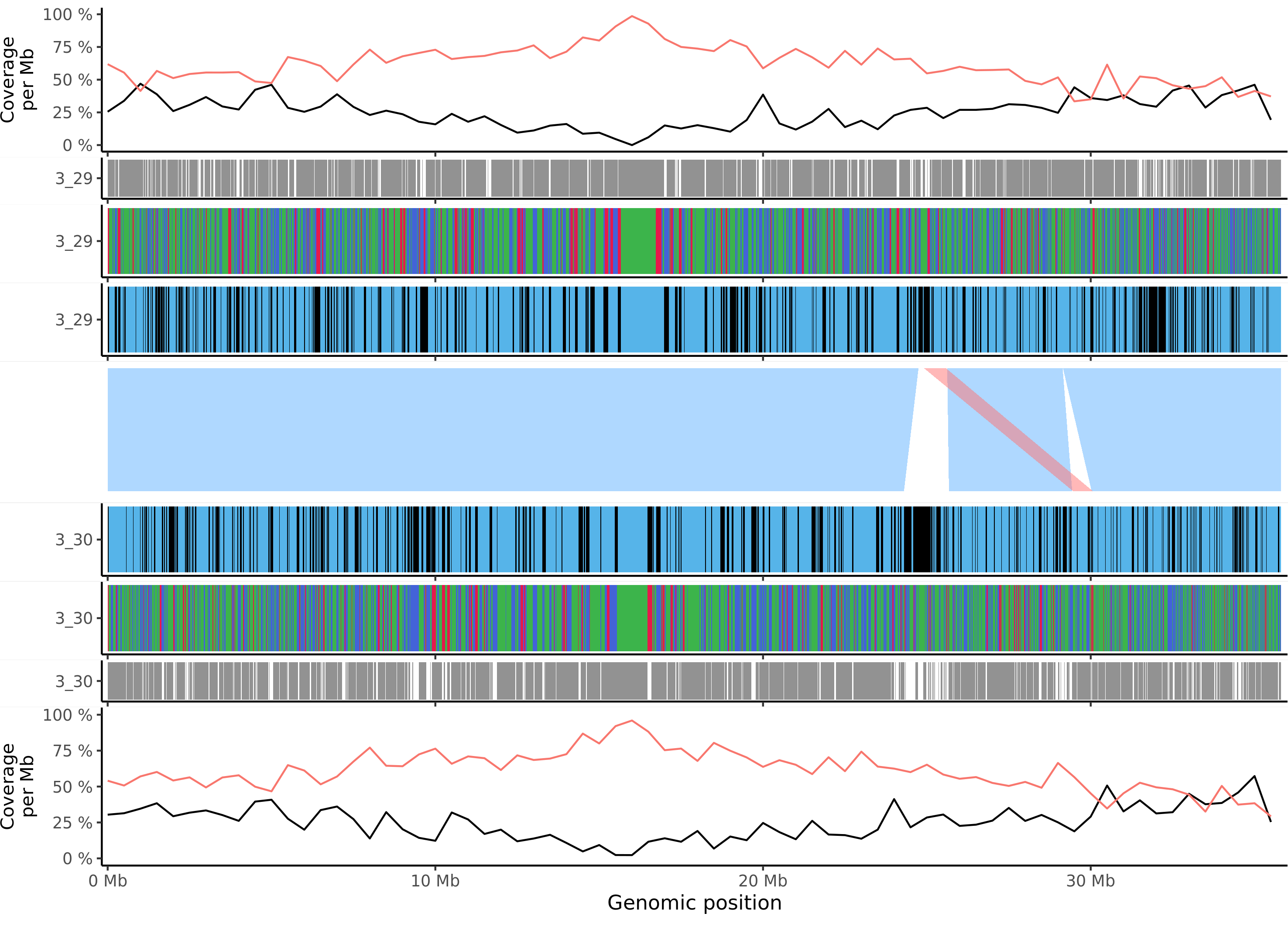

### msieversii_chr_8.png

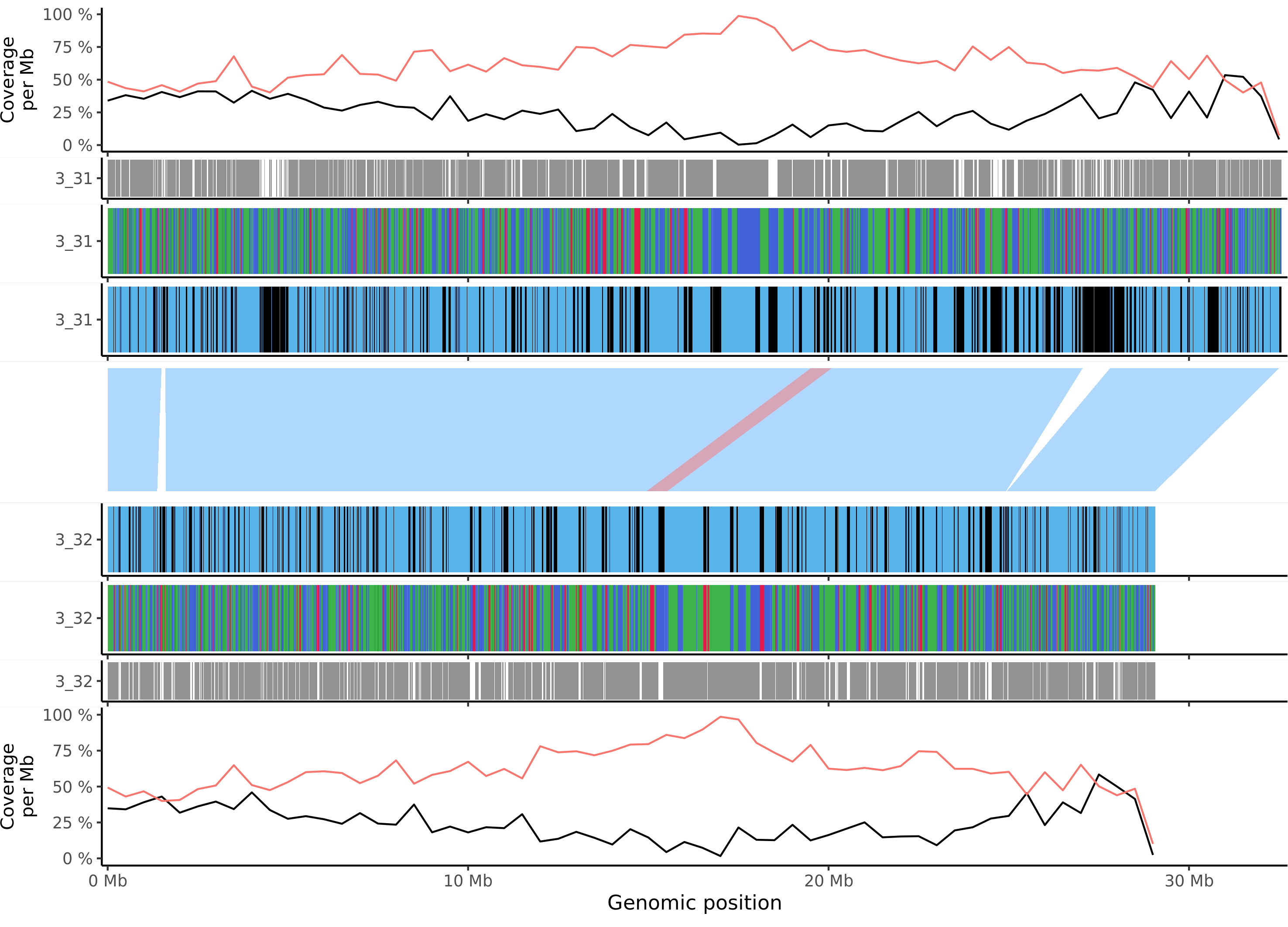

### msieversii_chr_9.png

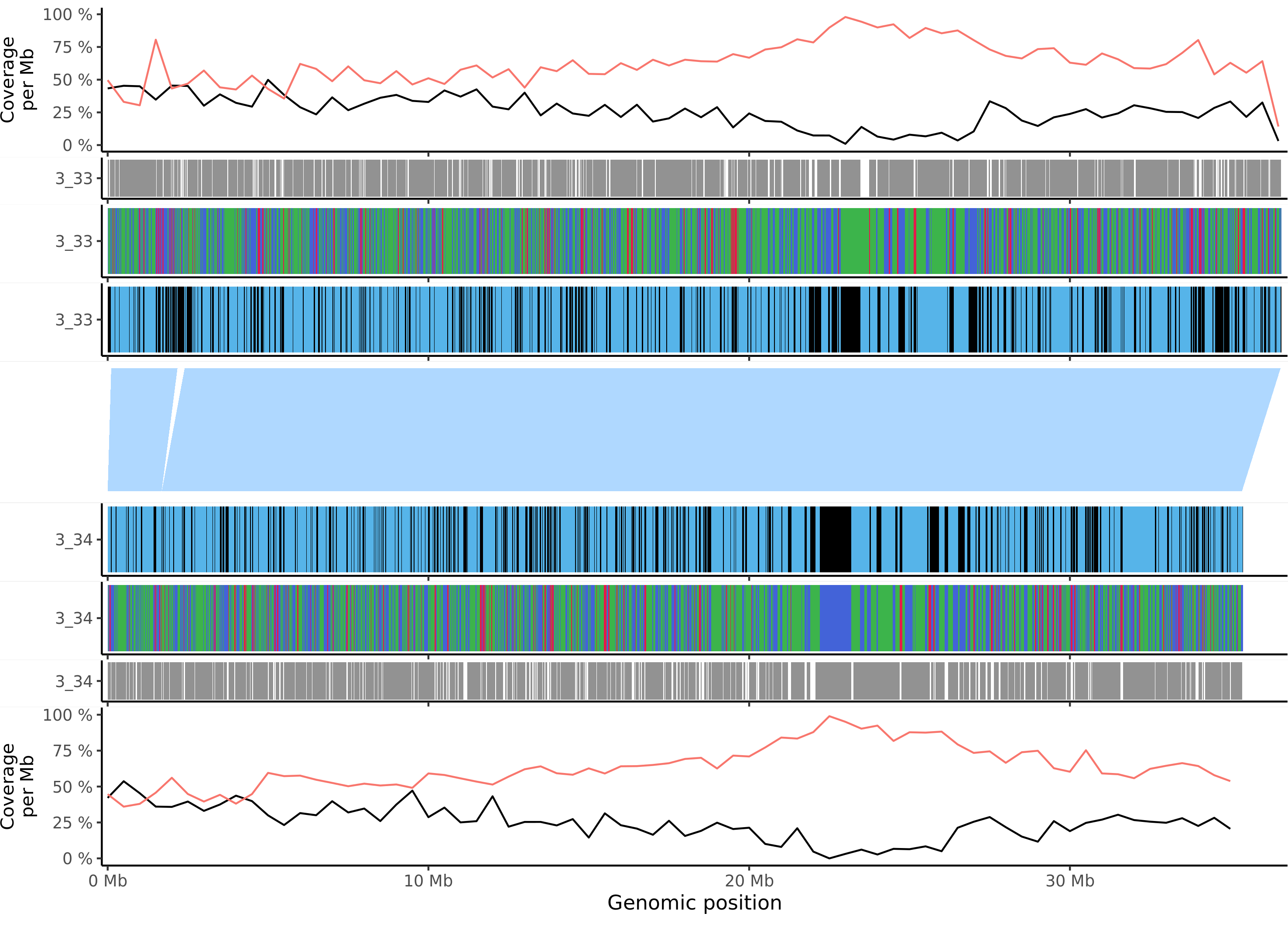

### msieversii_chr_10.png

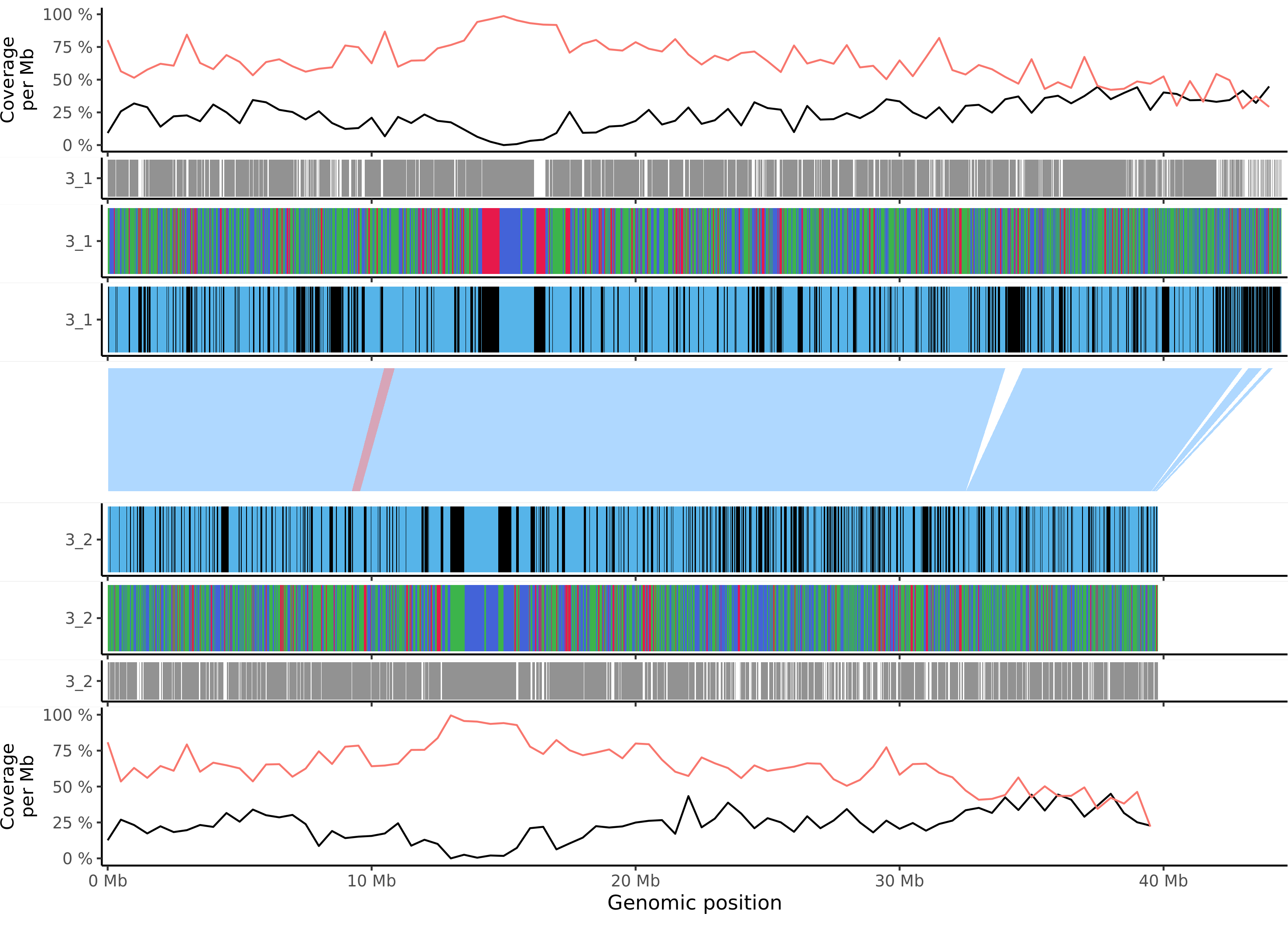

### msieversii_chr_11.png

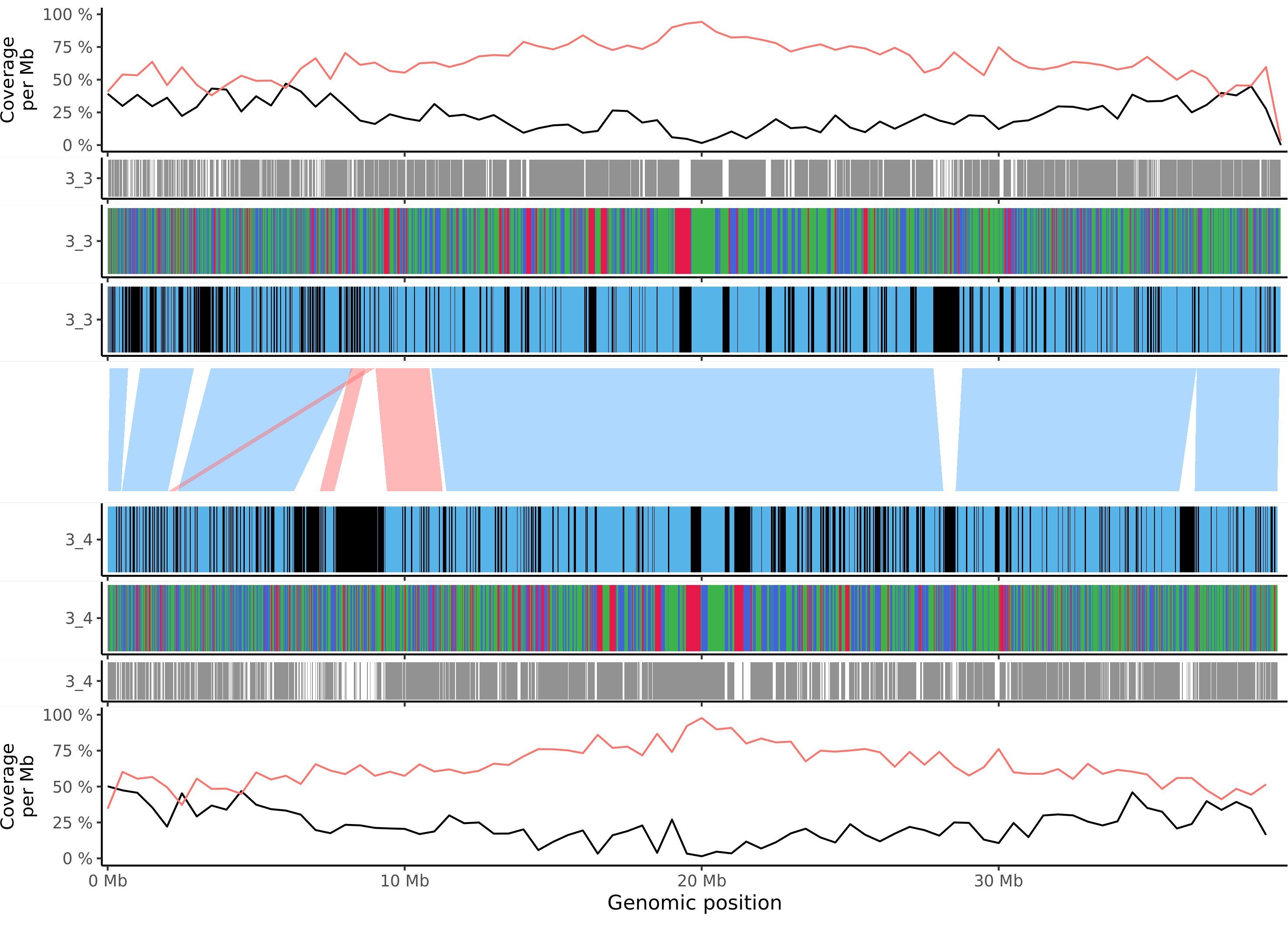

### msieversii_chr_12.png

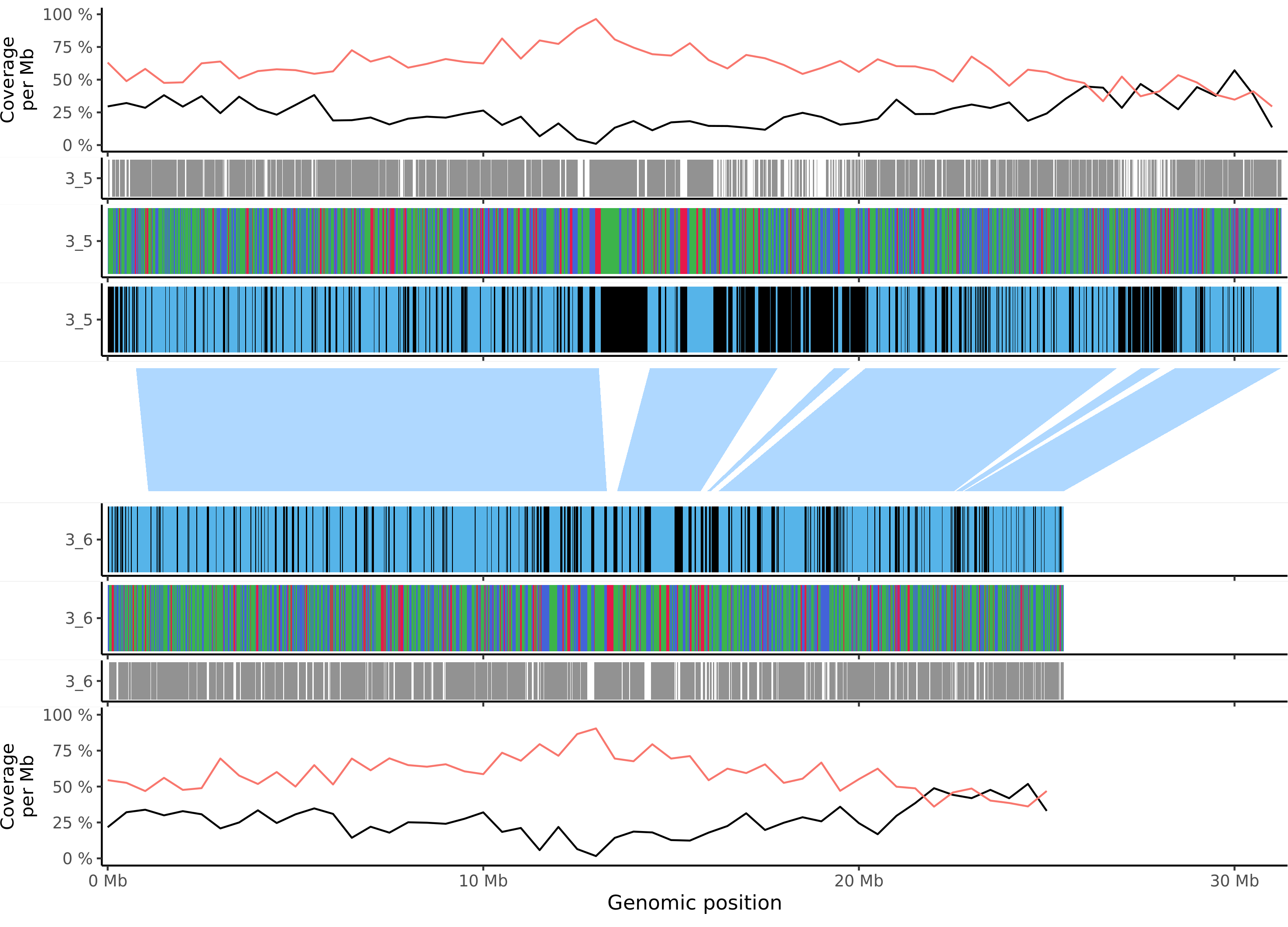

### msieversii_chr_13.png

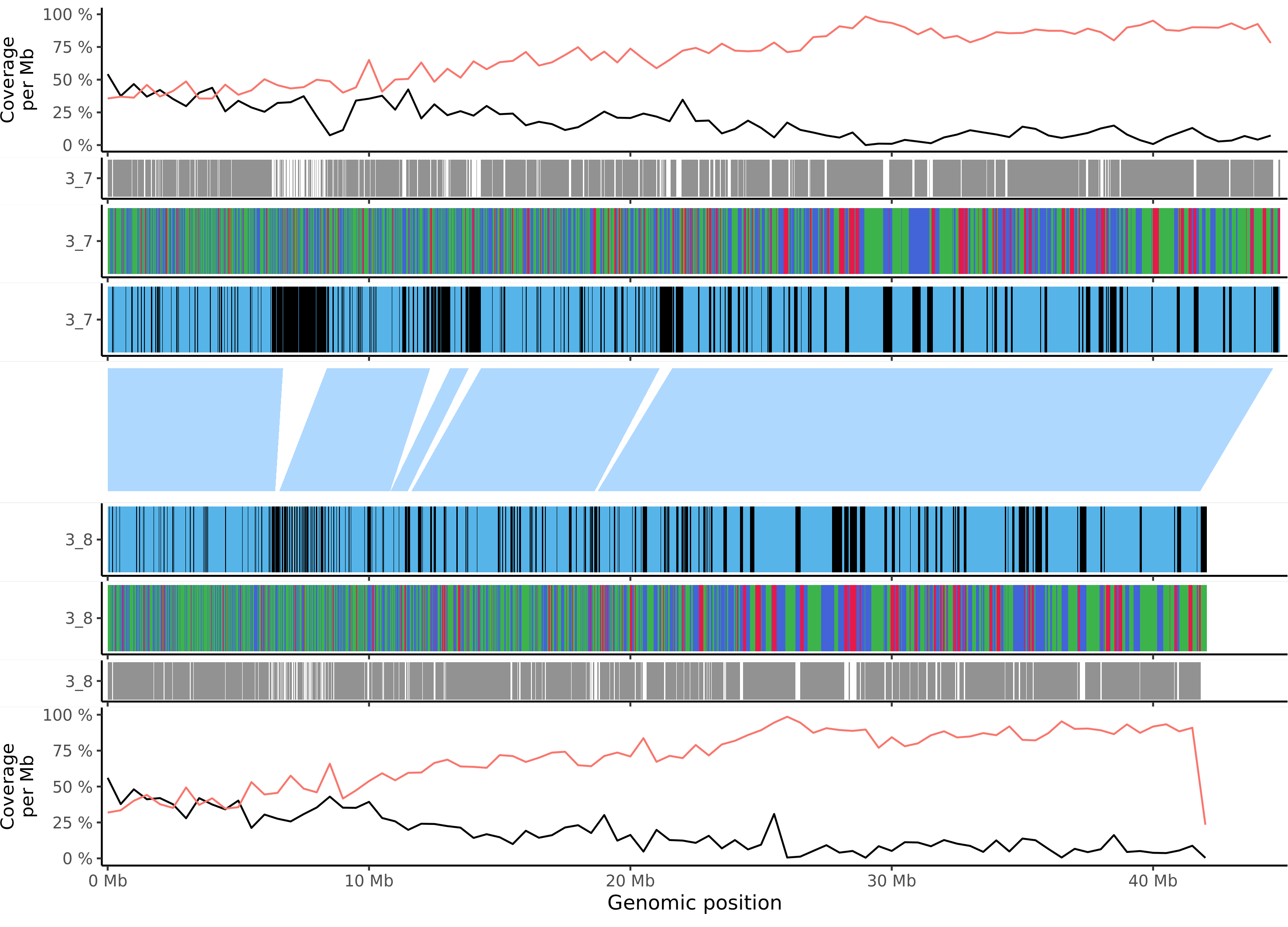

### msieversii_chr_14.png

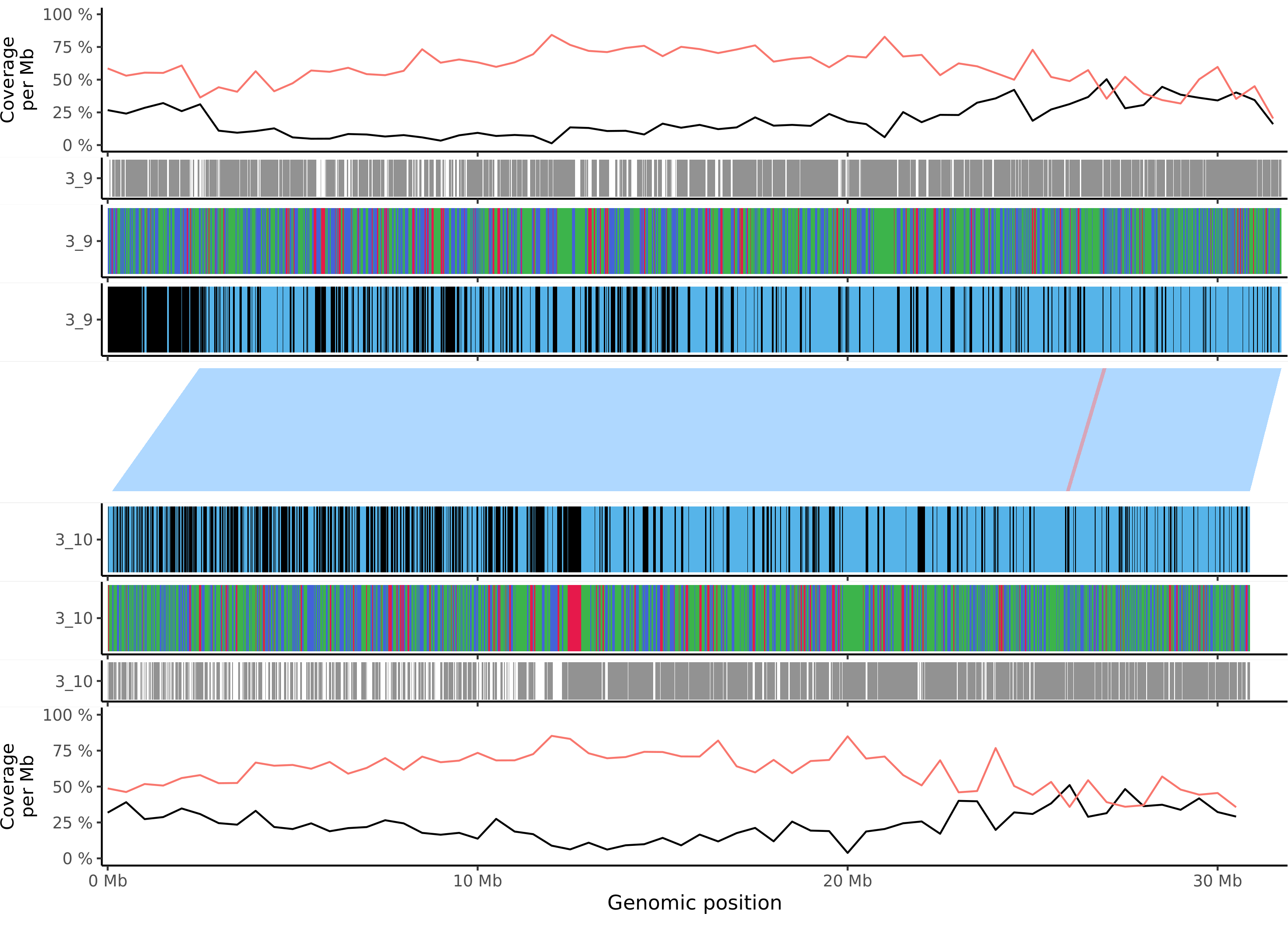

### msieversii_chr_15.png

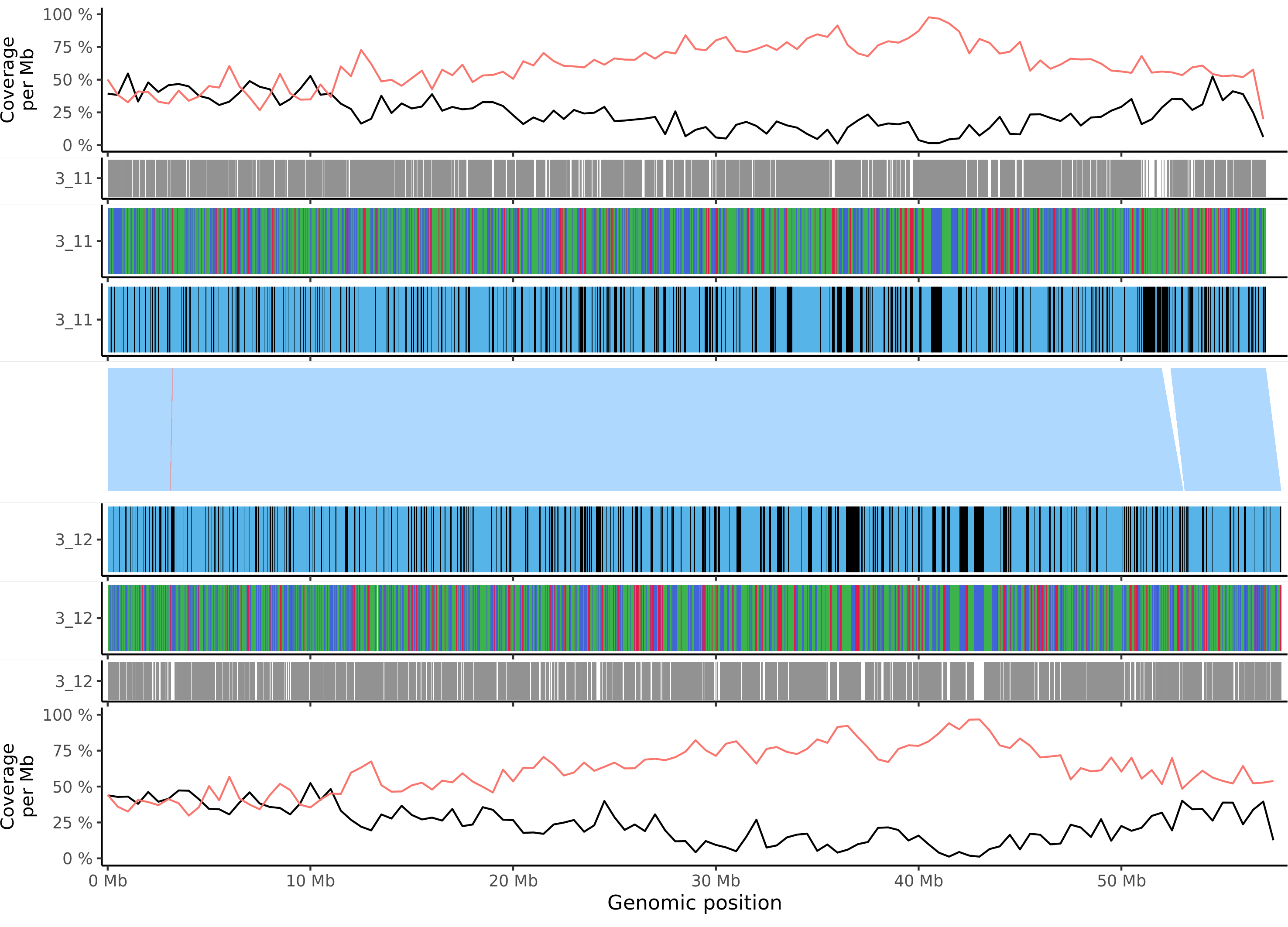

### msieversii_chr_16.png

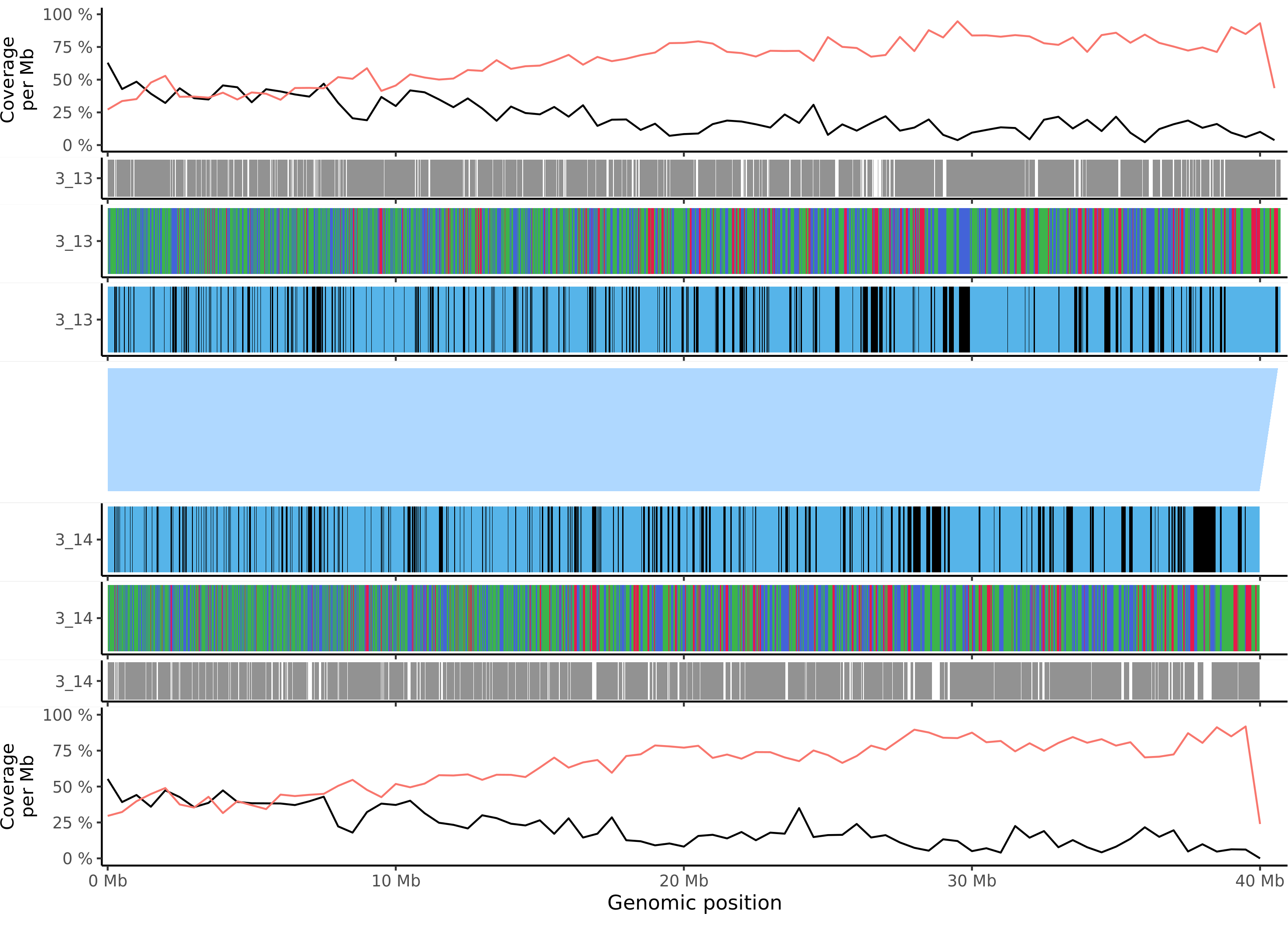

### msieversii_chr_17.png

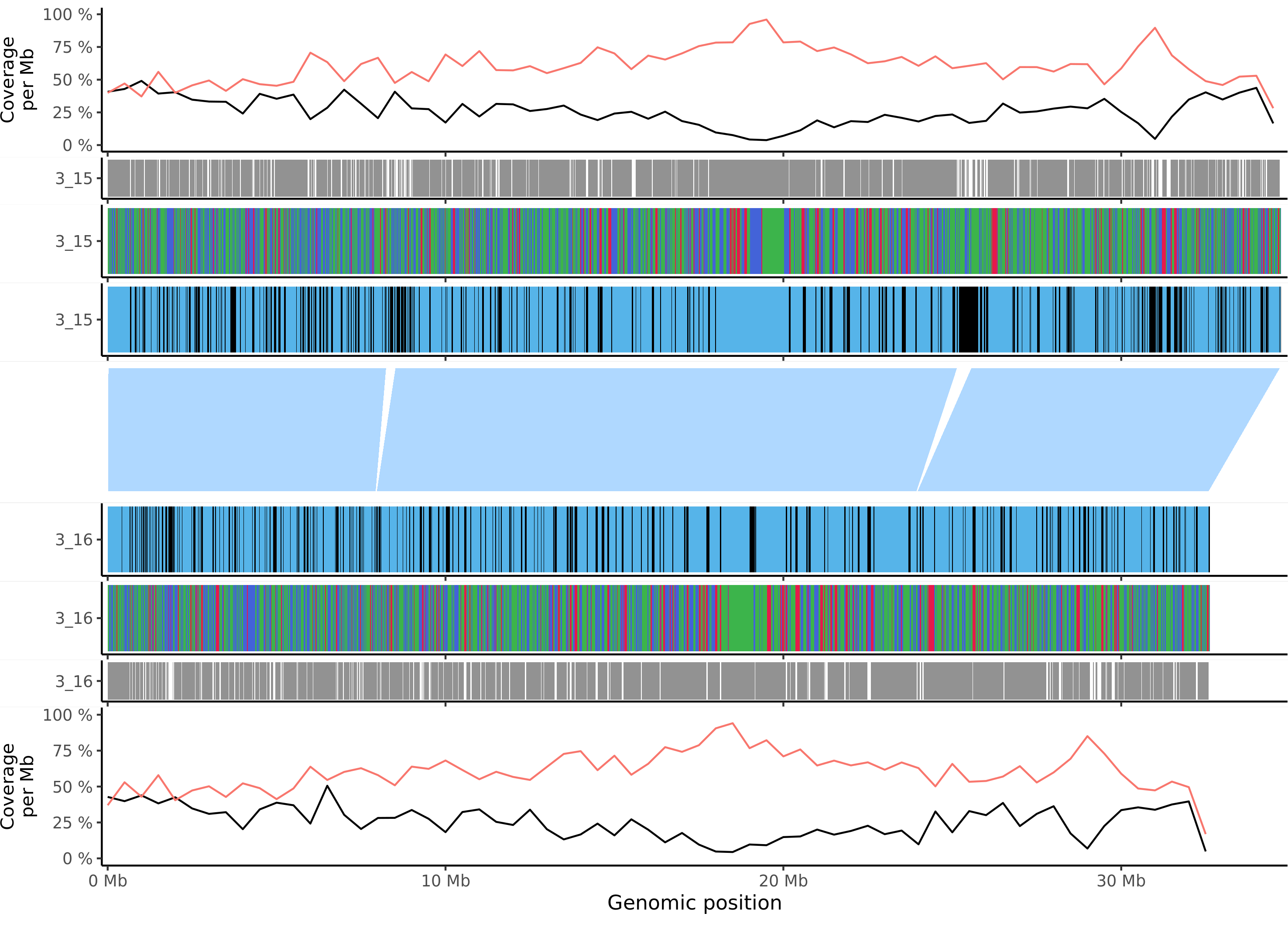

### msylvestris_chr_1.png

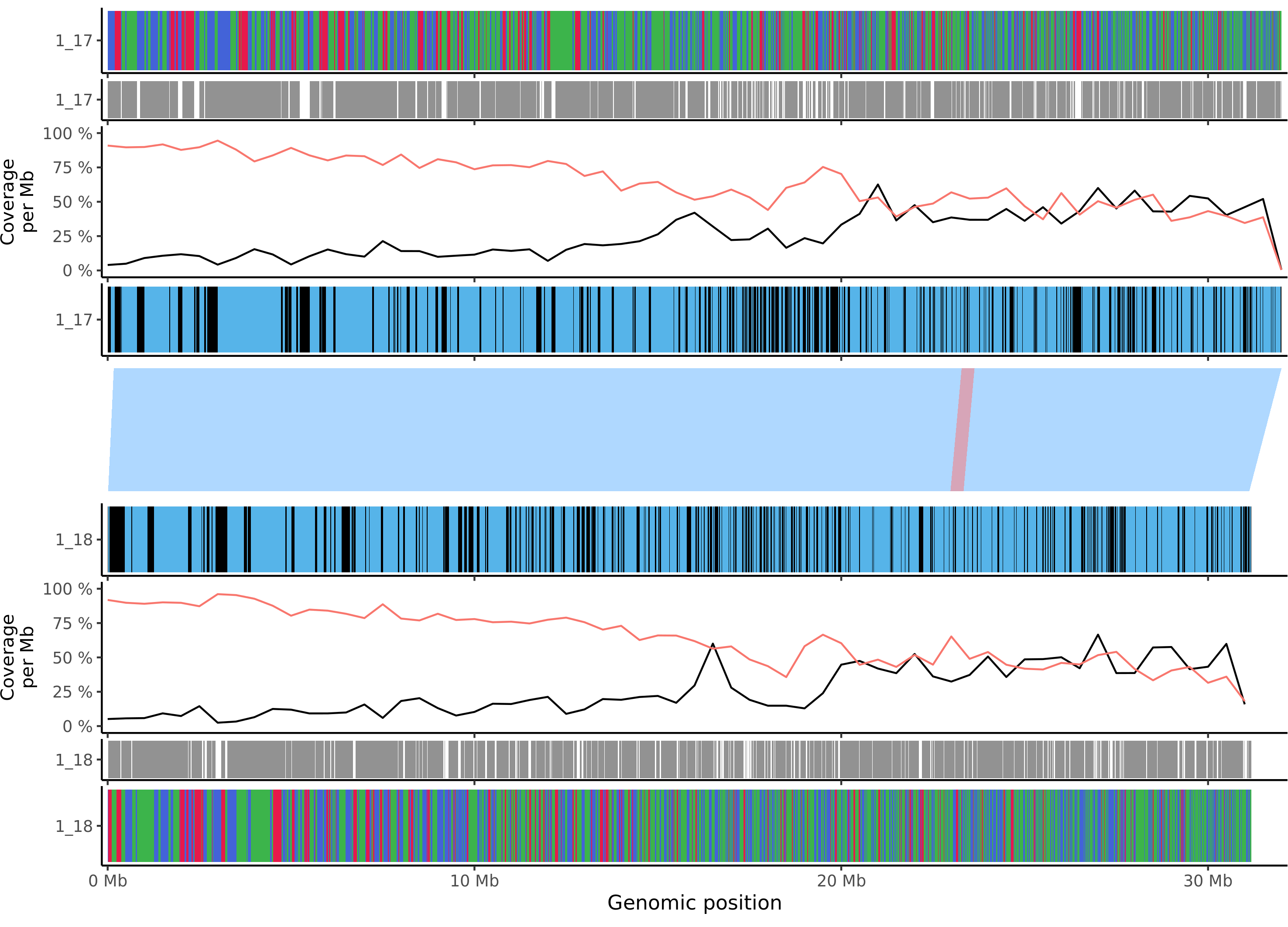

### msylvestris_chr_2.png

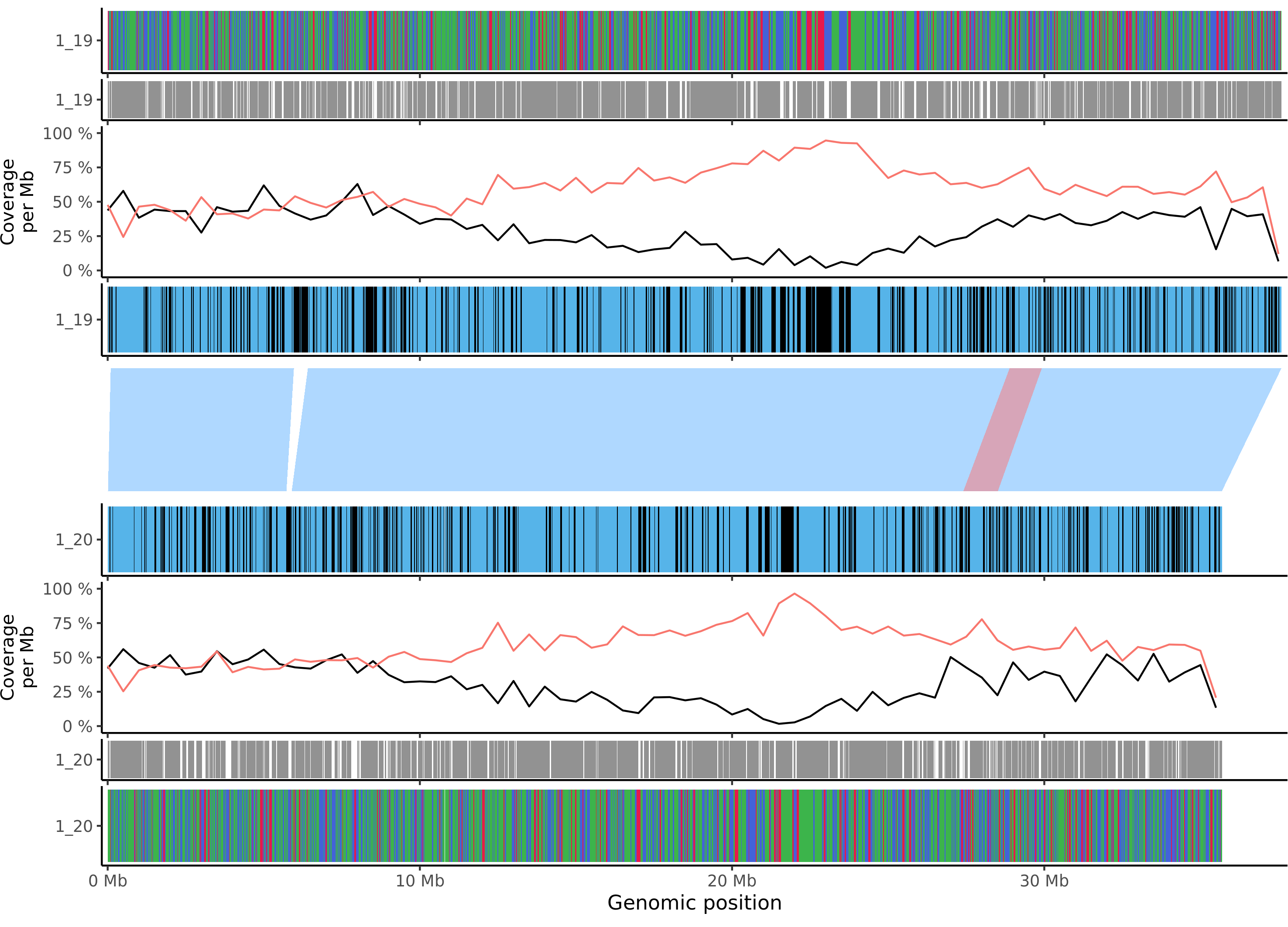

### msylvestris_chr_3.png

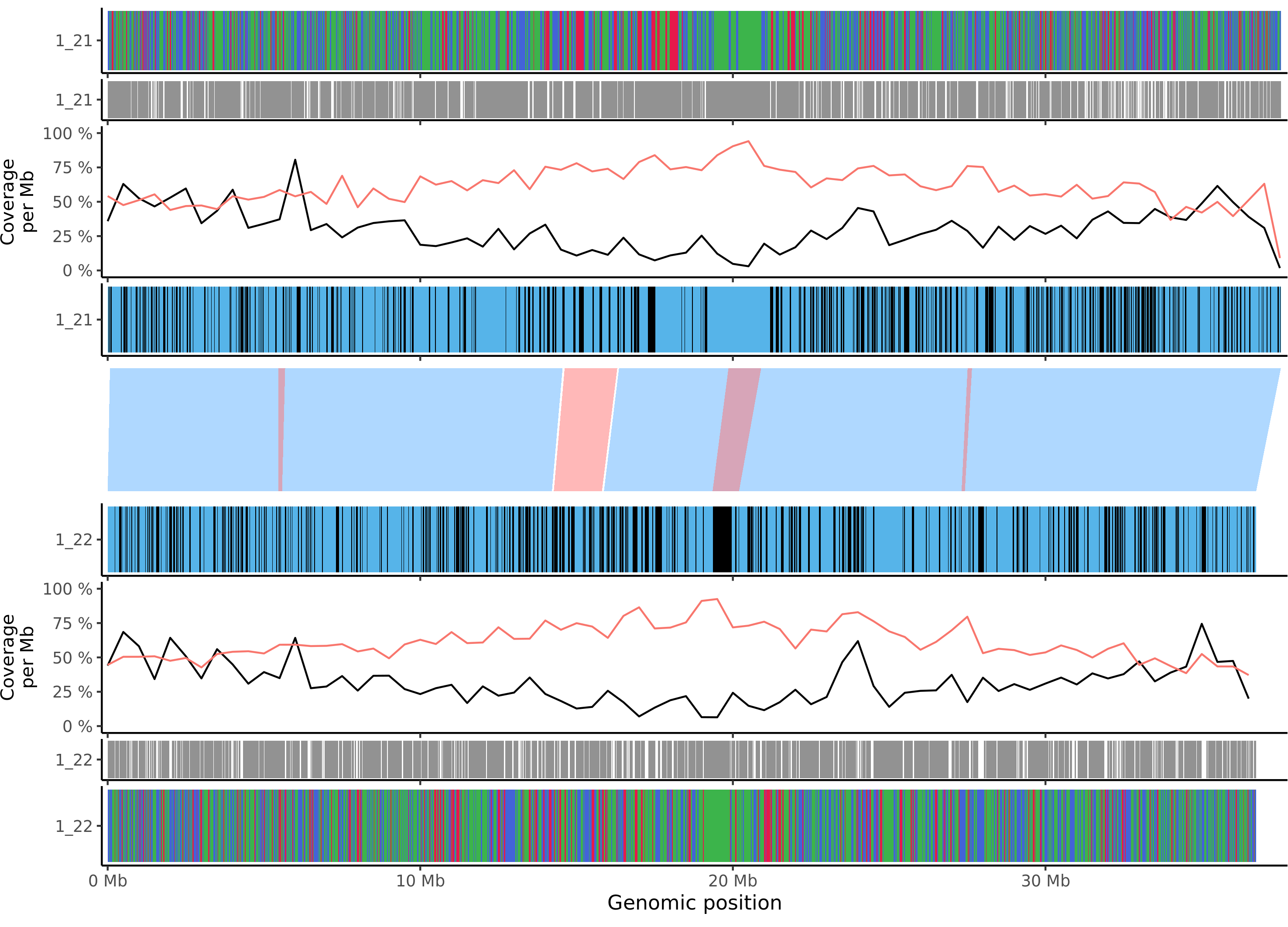

### msylvestris_chr_4.png

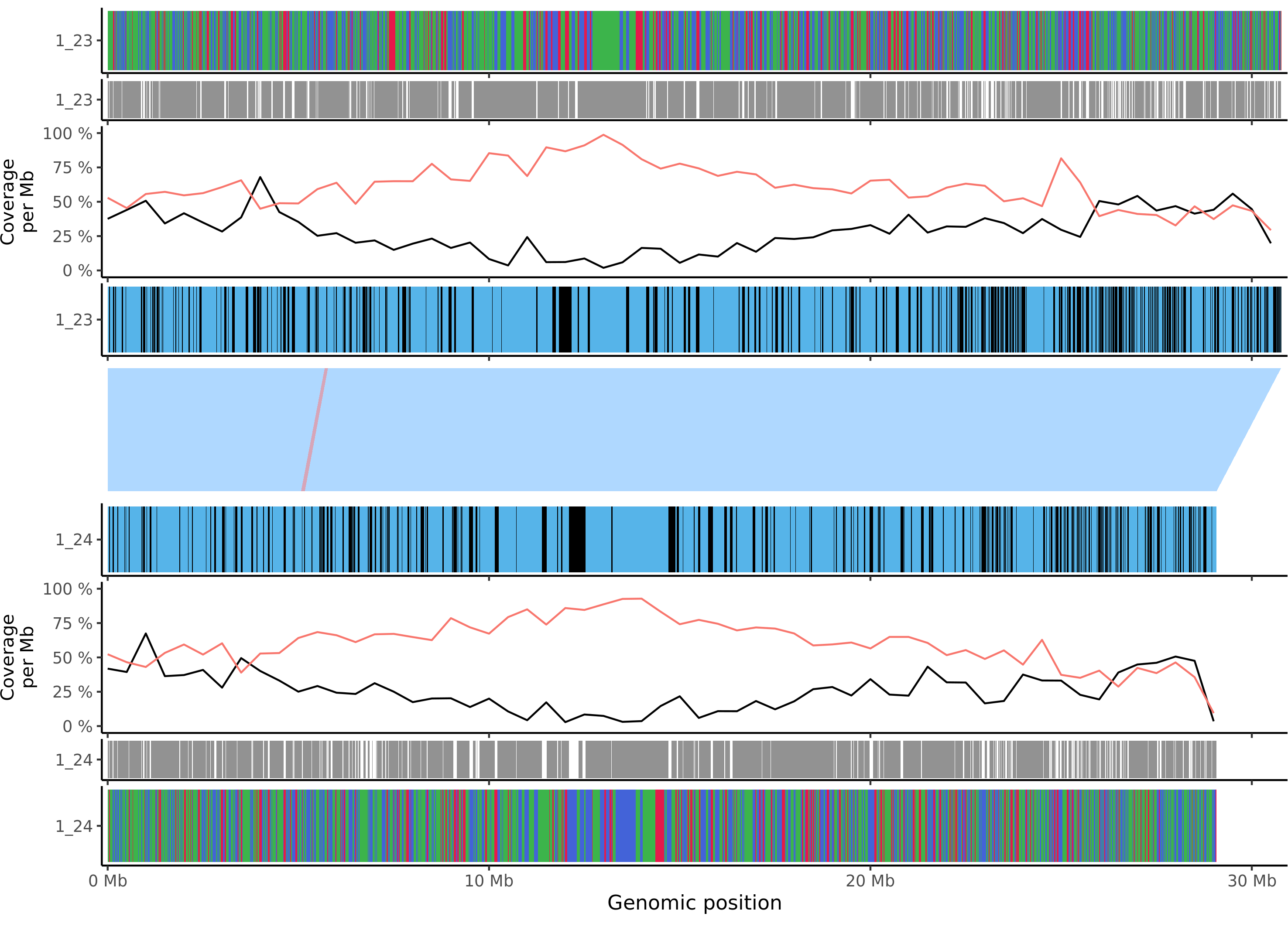

### msylvestris_chr_5.png

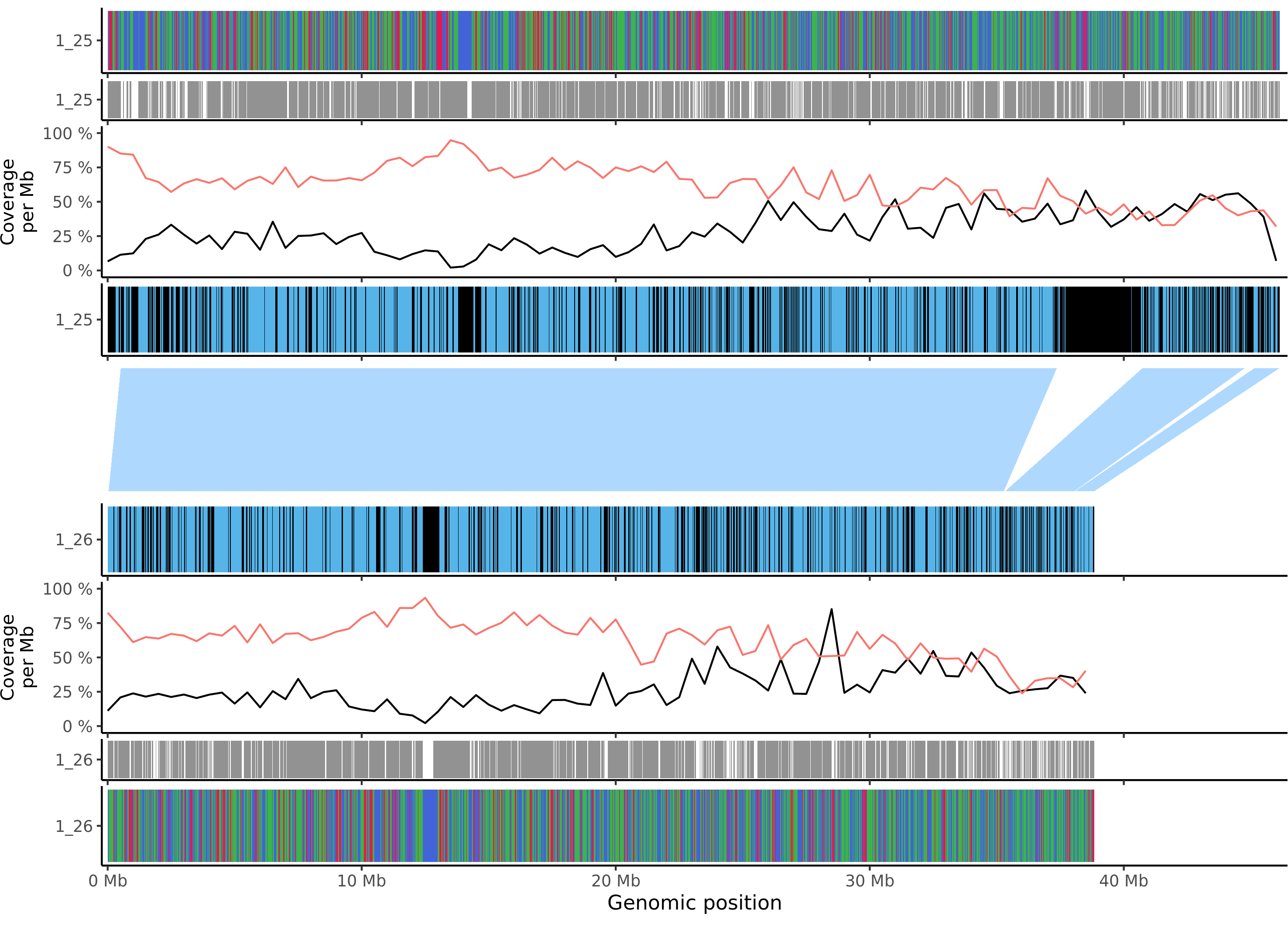

### msylvestris_chr_6.png

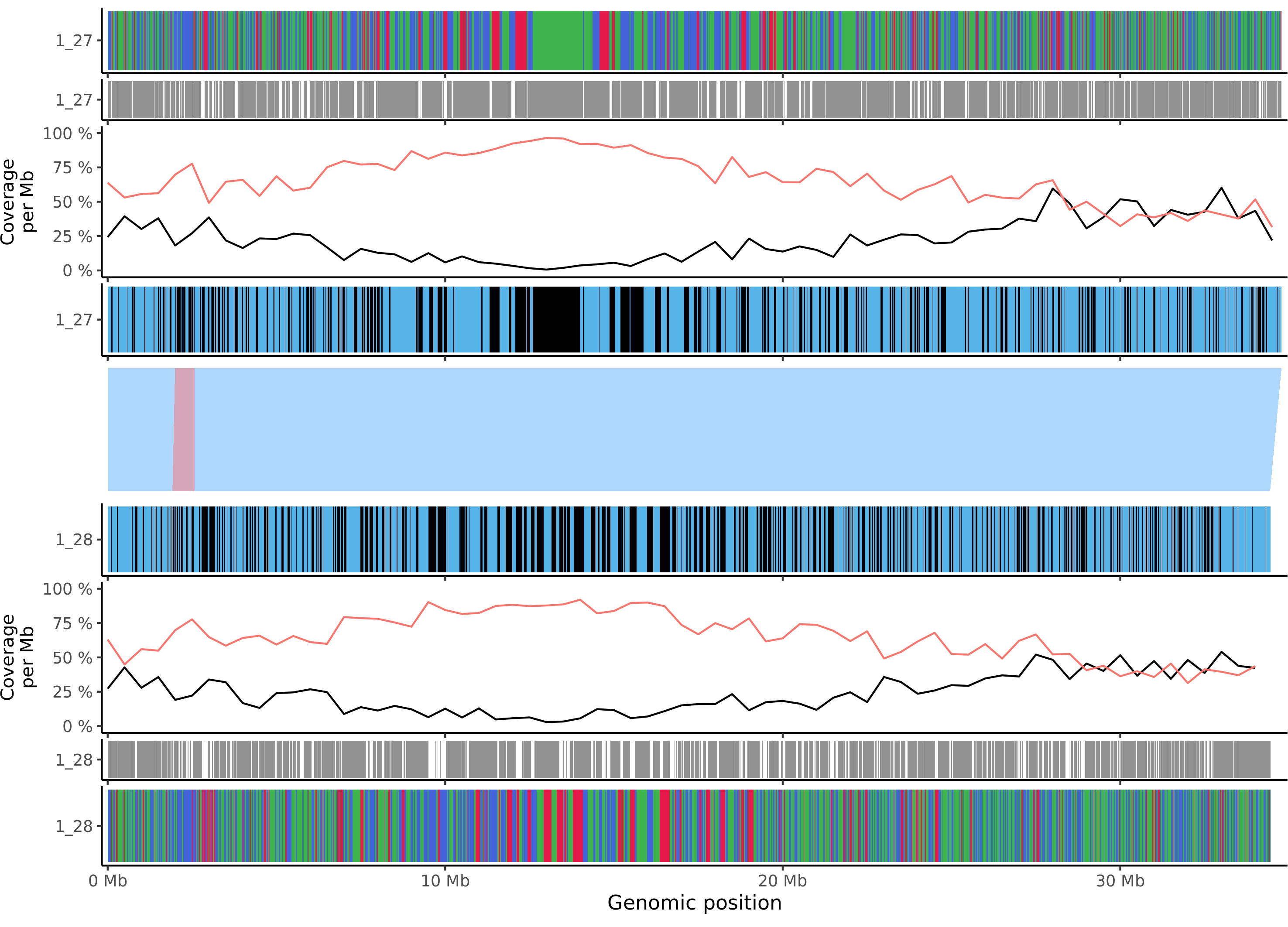

### msylvestris_chr_7.png

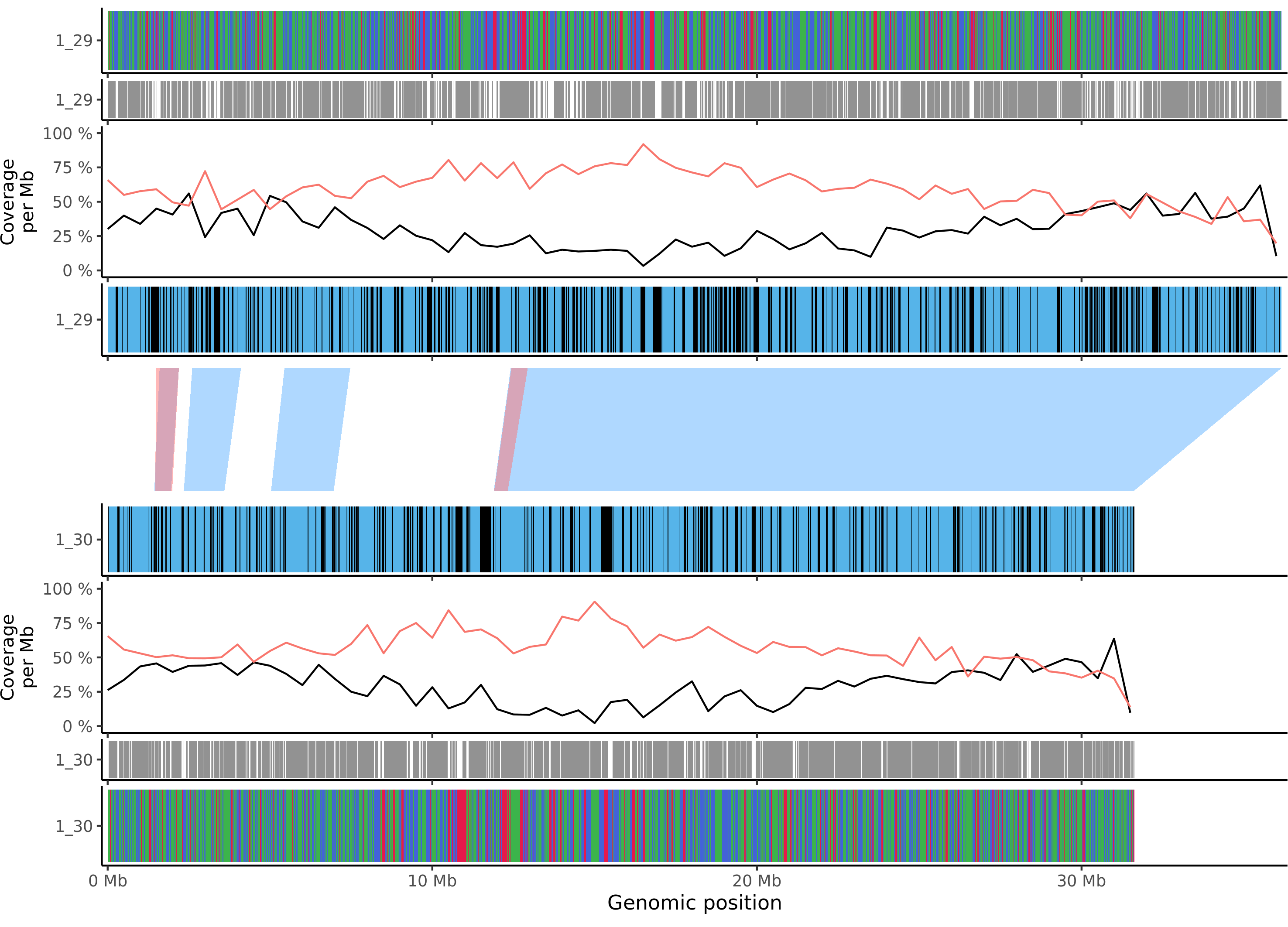

### msylvestris_chr_8.png

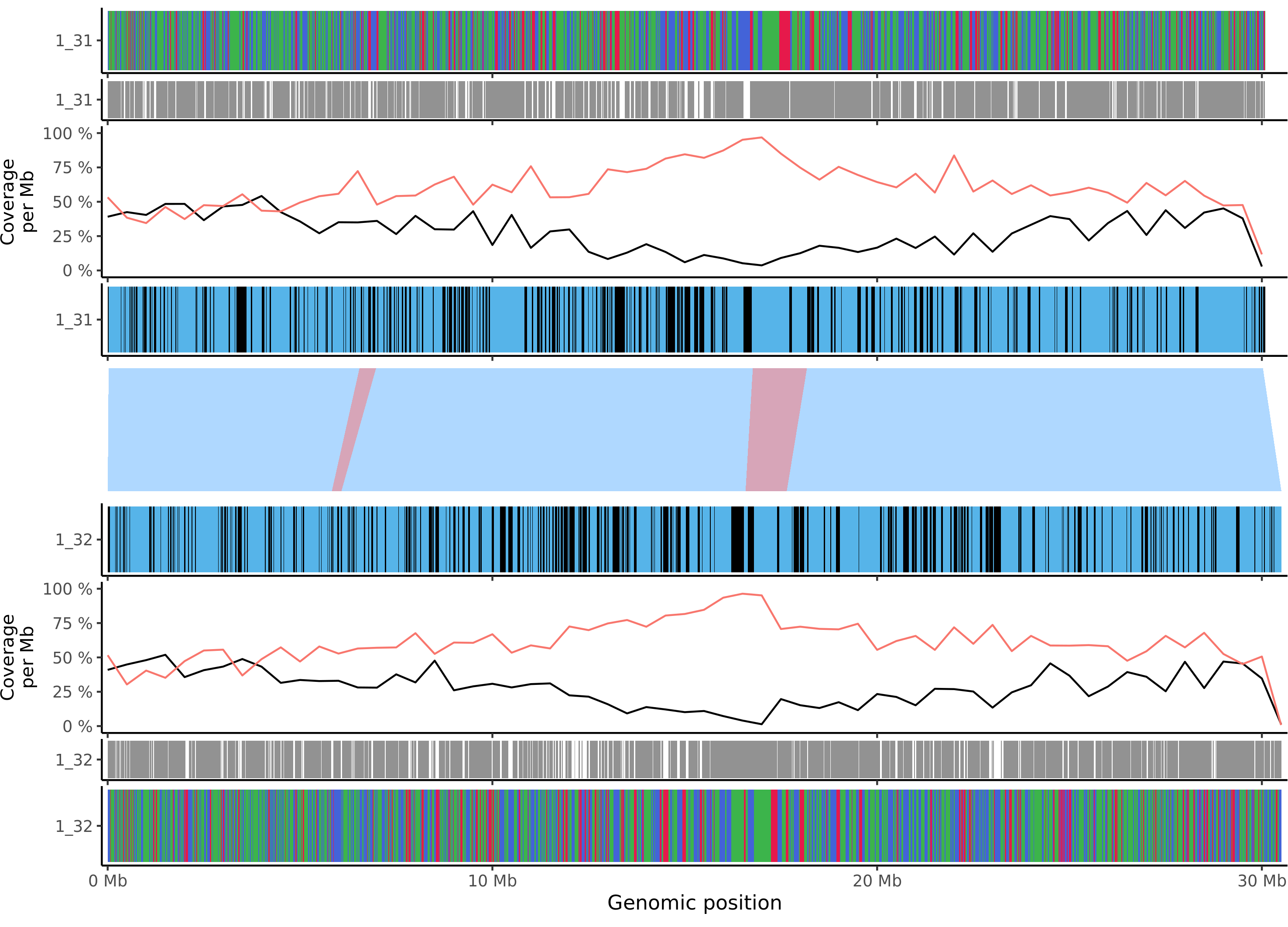

### msylvestris_chr_9.png

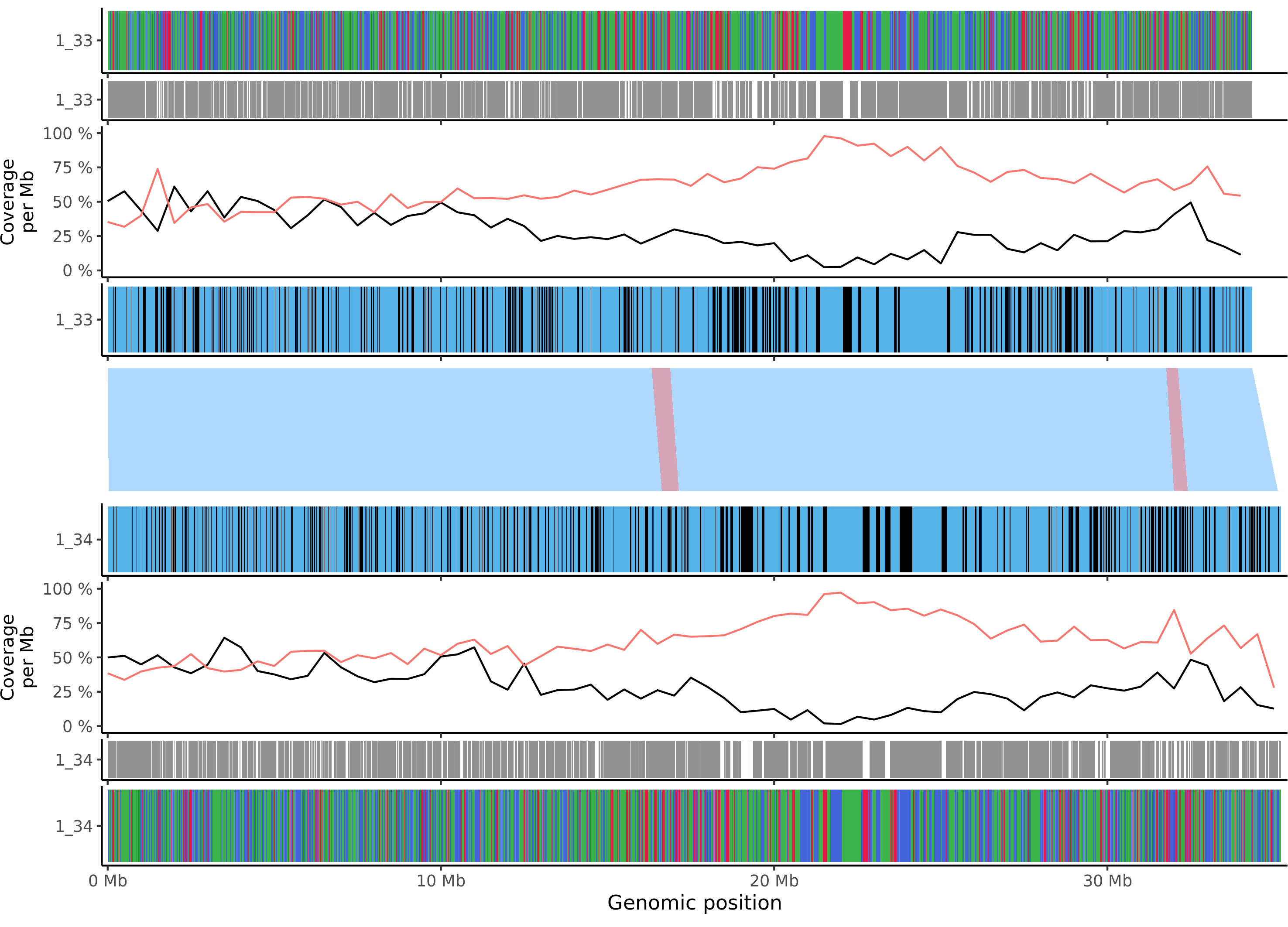

### msylvestris_chr_10.png

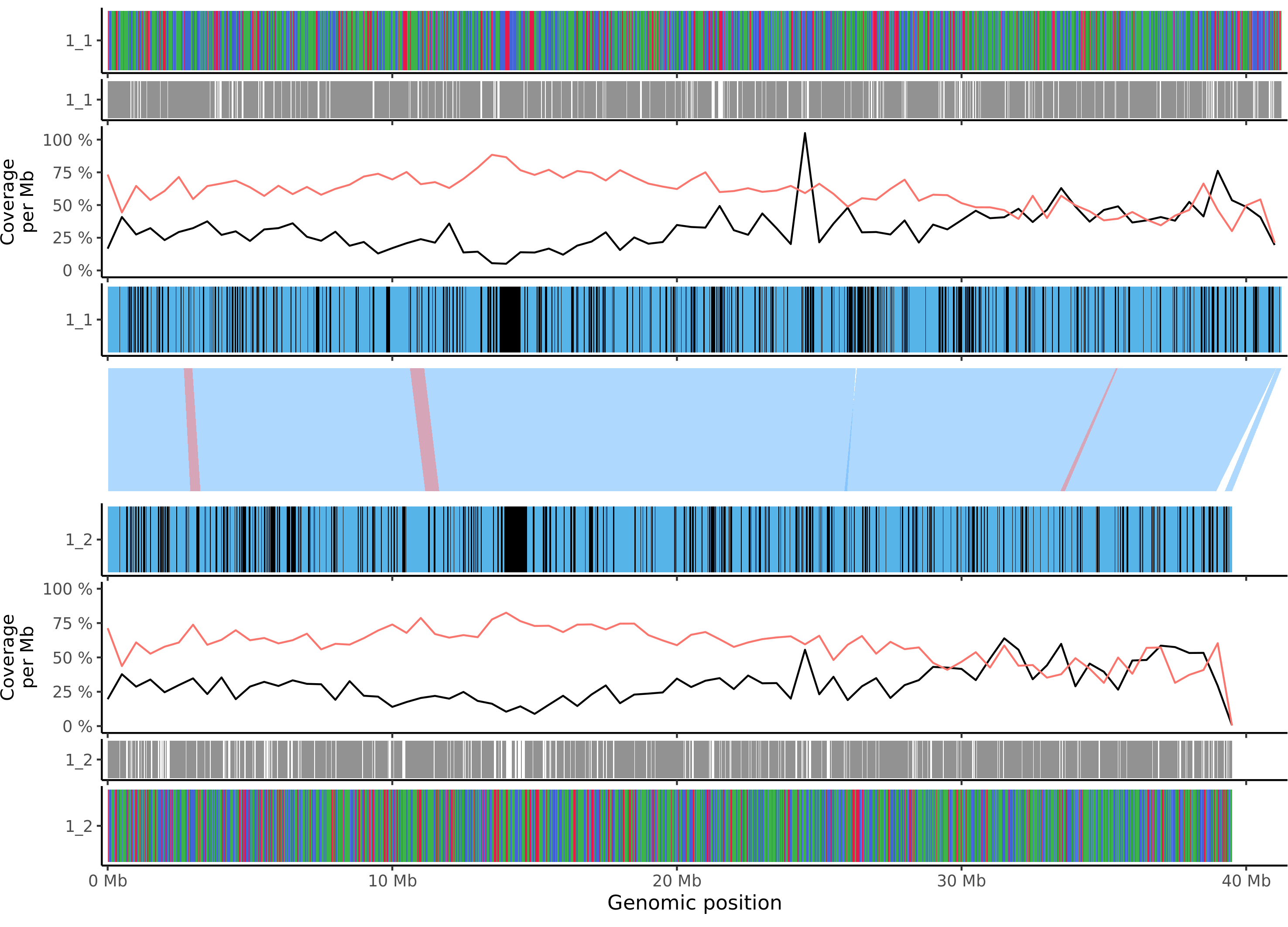

### msylvestris_chr_11.png

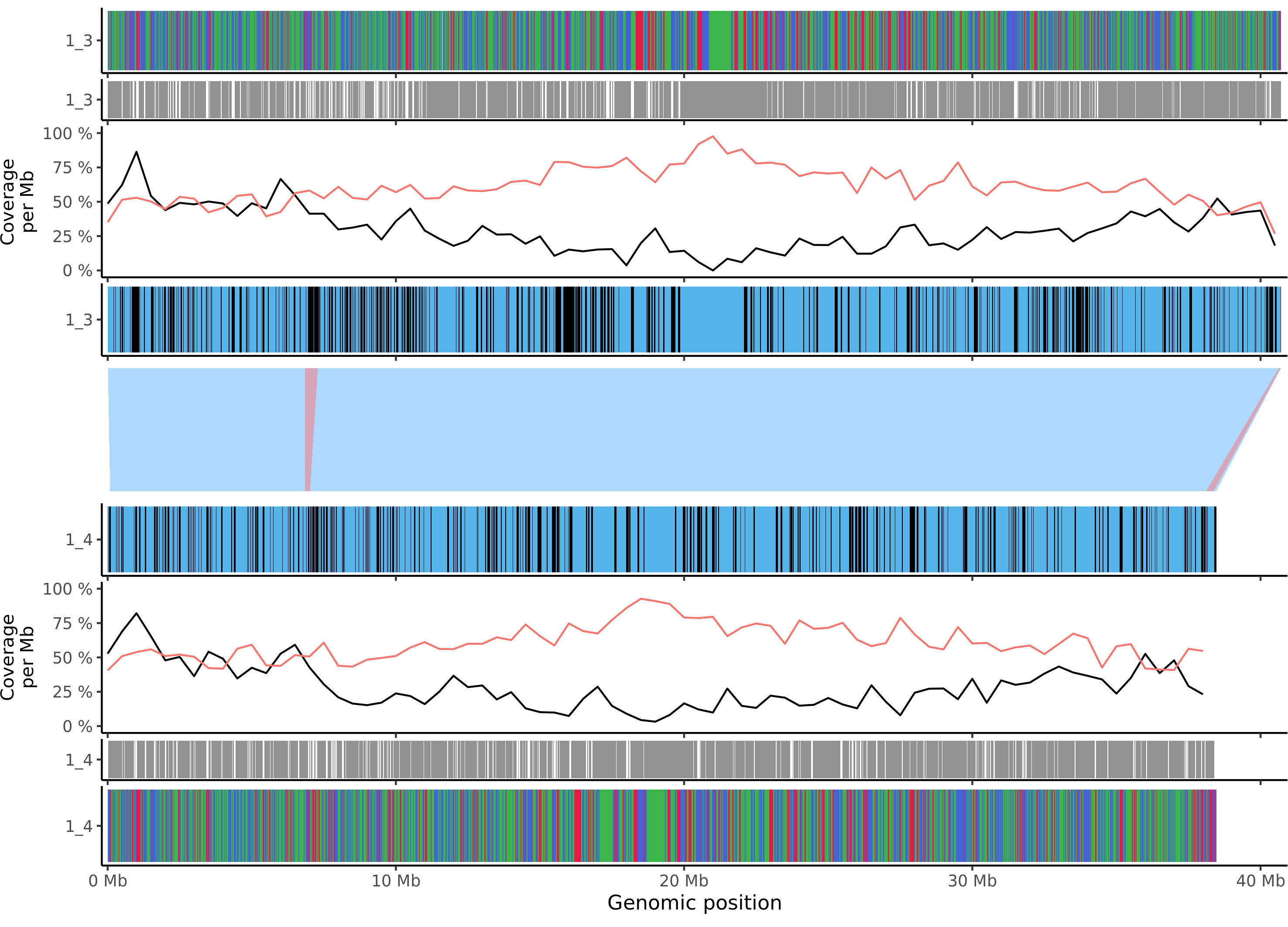

### msylvestris_chr_12.png

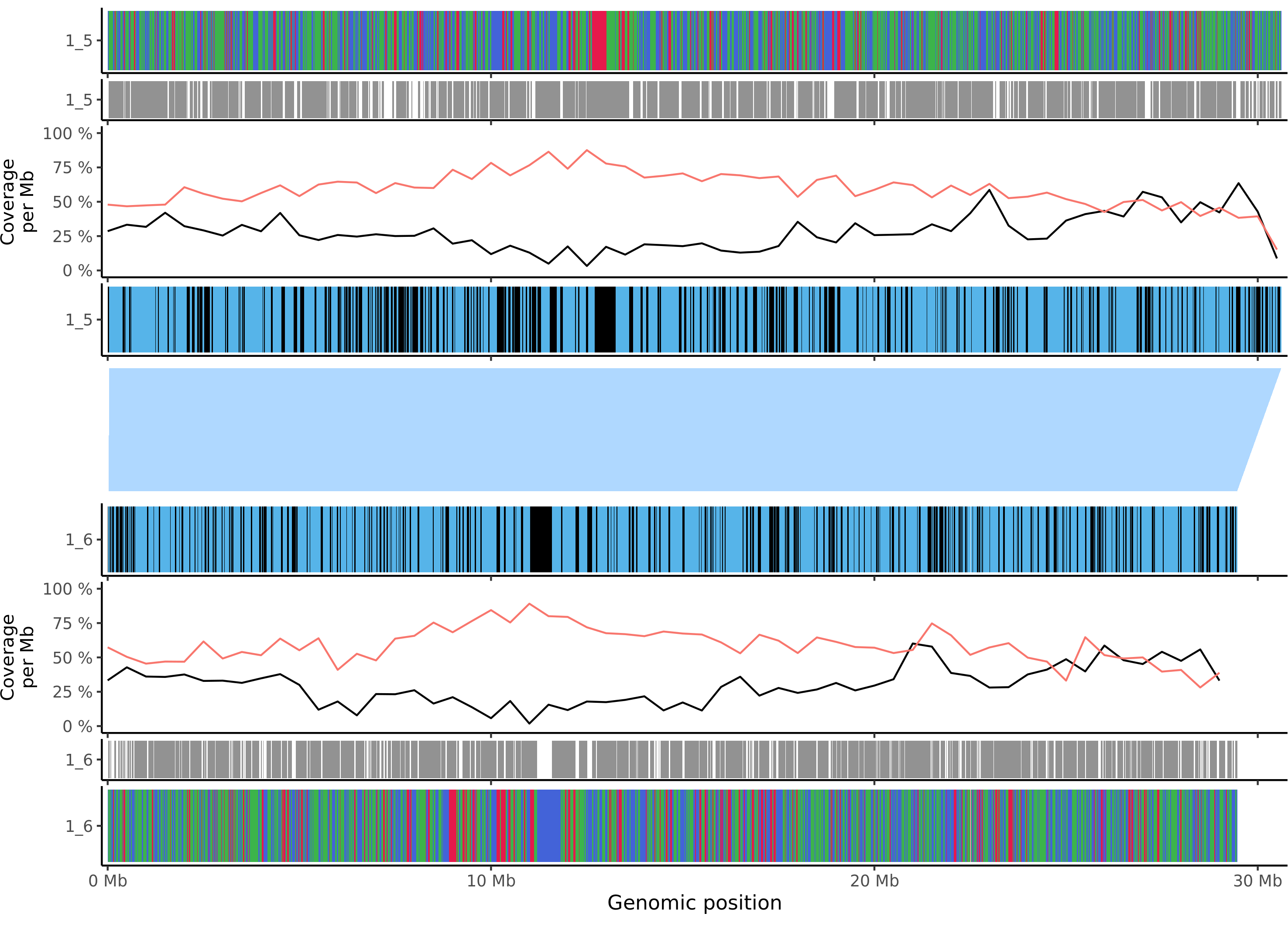

### msylvestris_chr_13.png

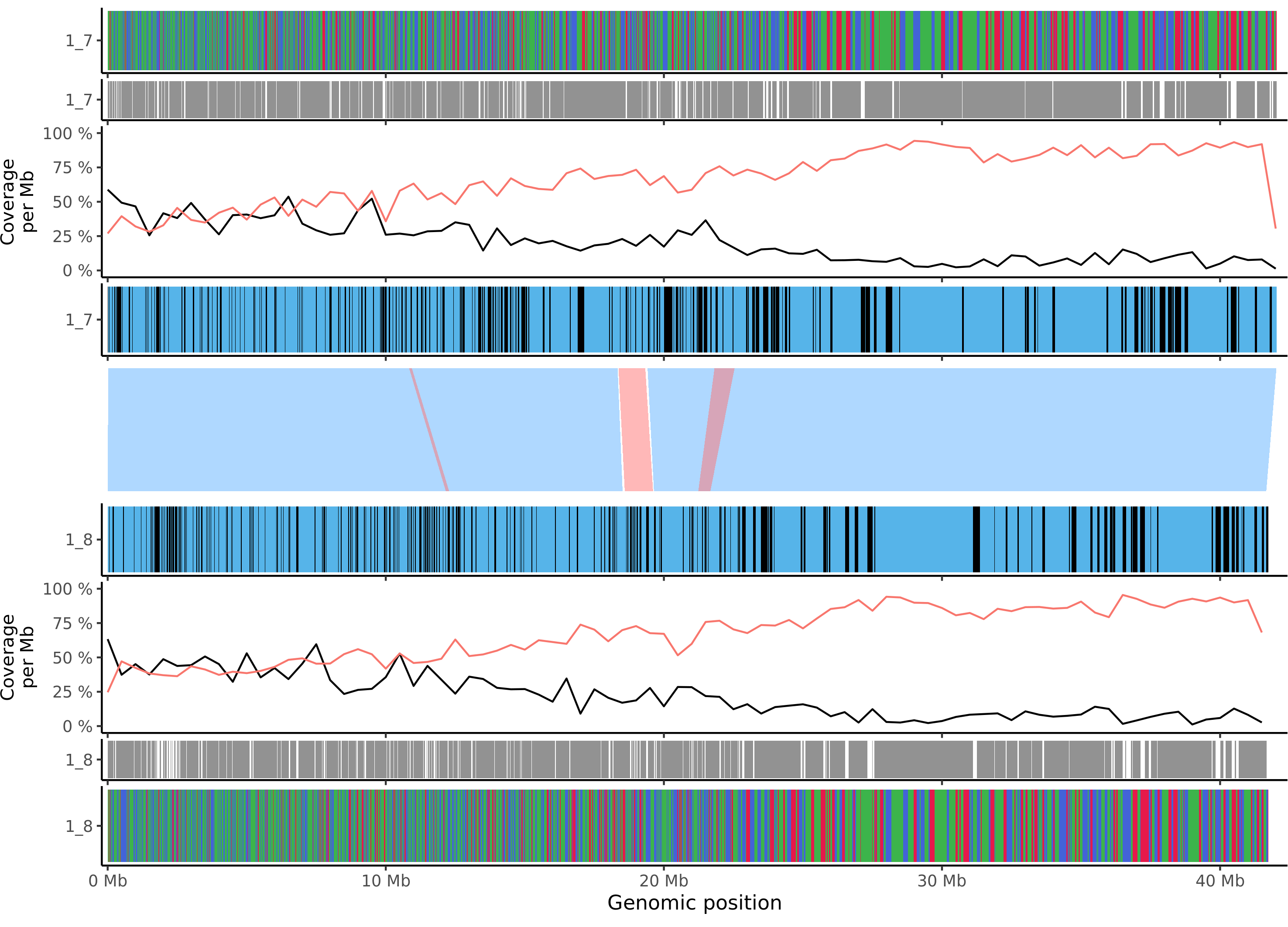

### msylvestris_chr_14.png

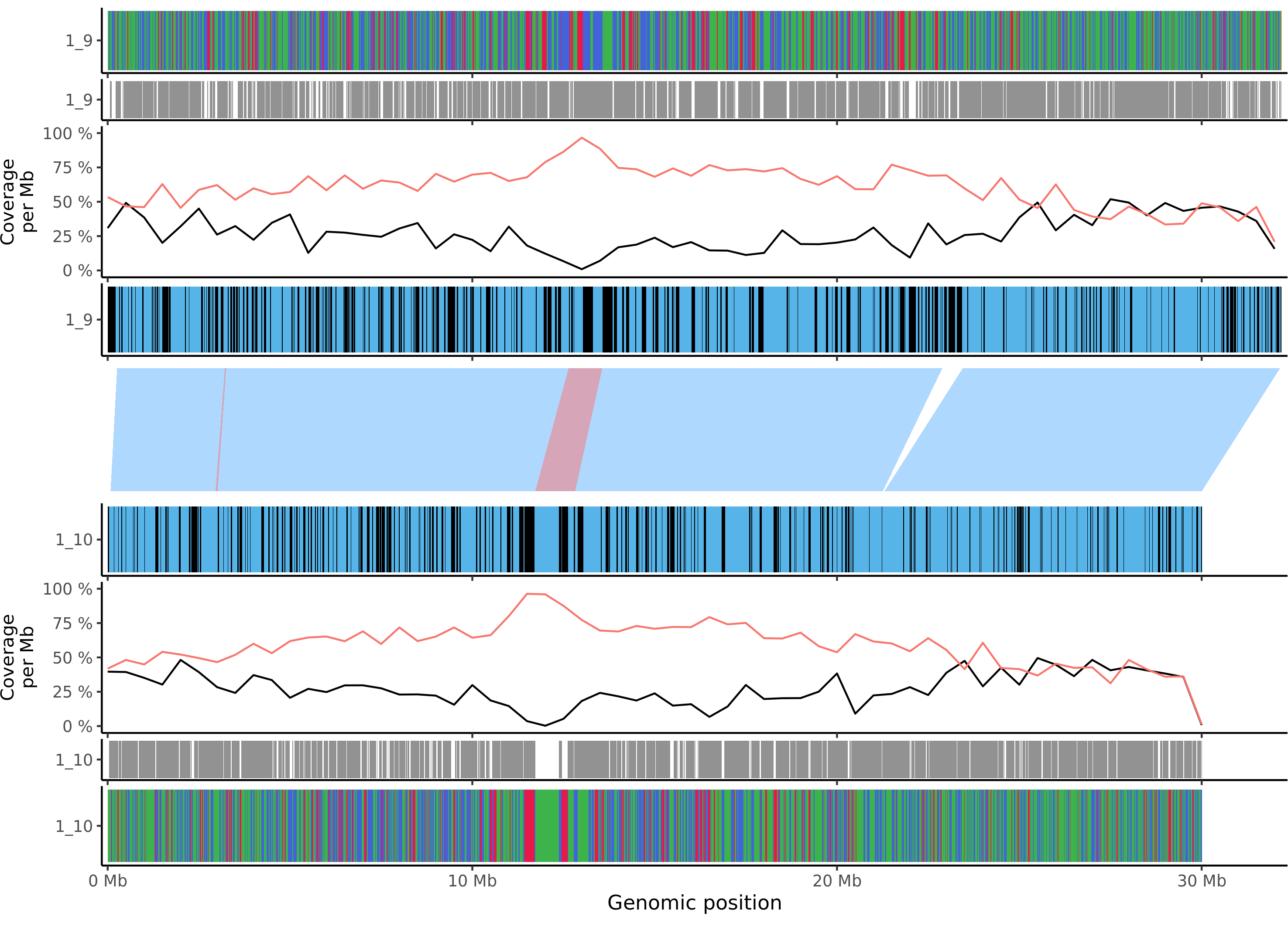

### msylvestris_chr_15.png

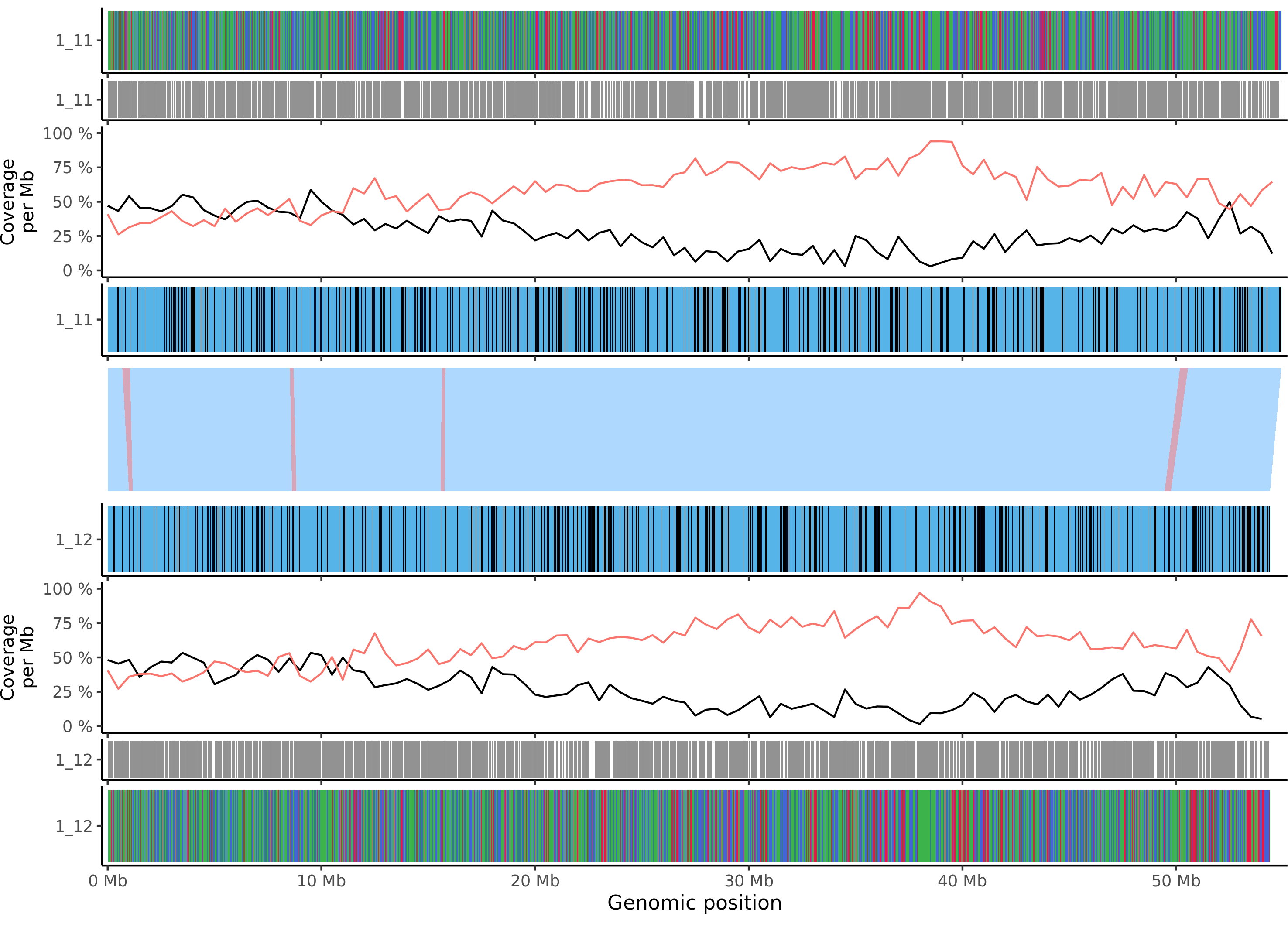

### msylvestris_chr_16.png

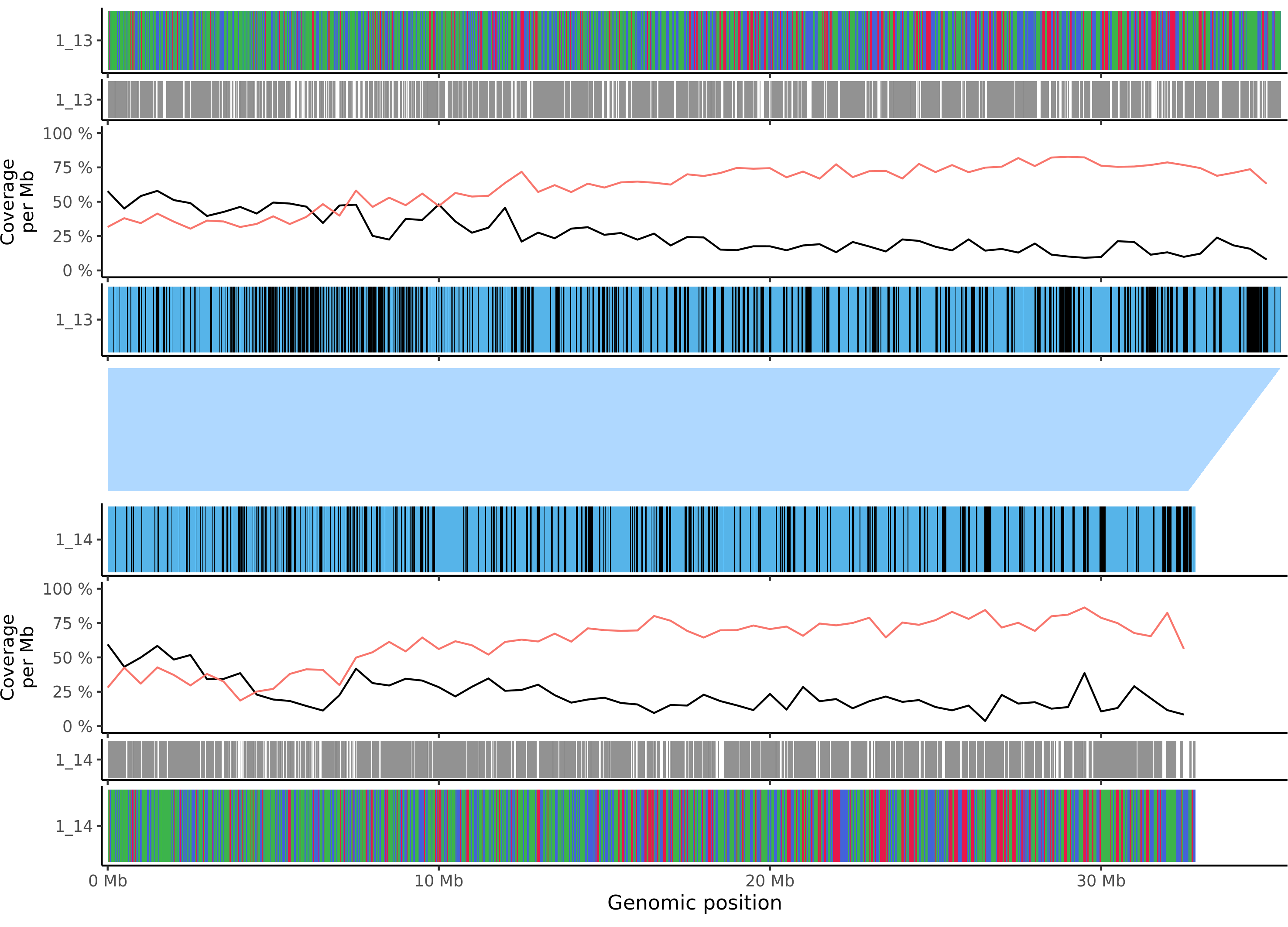

### msylvestris_chr_17.png

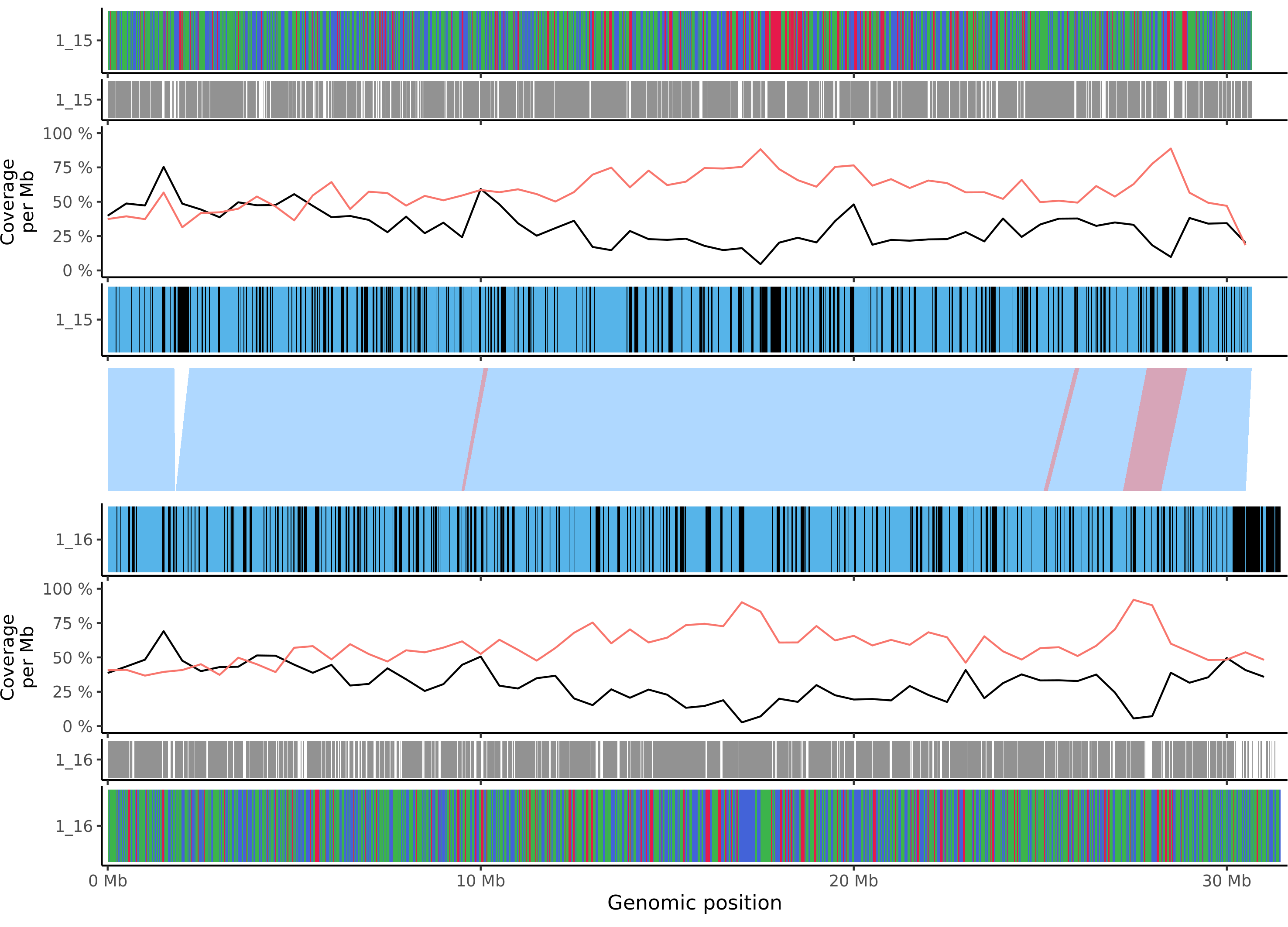

### Otava_chr_1.png

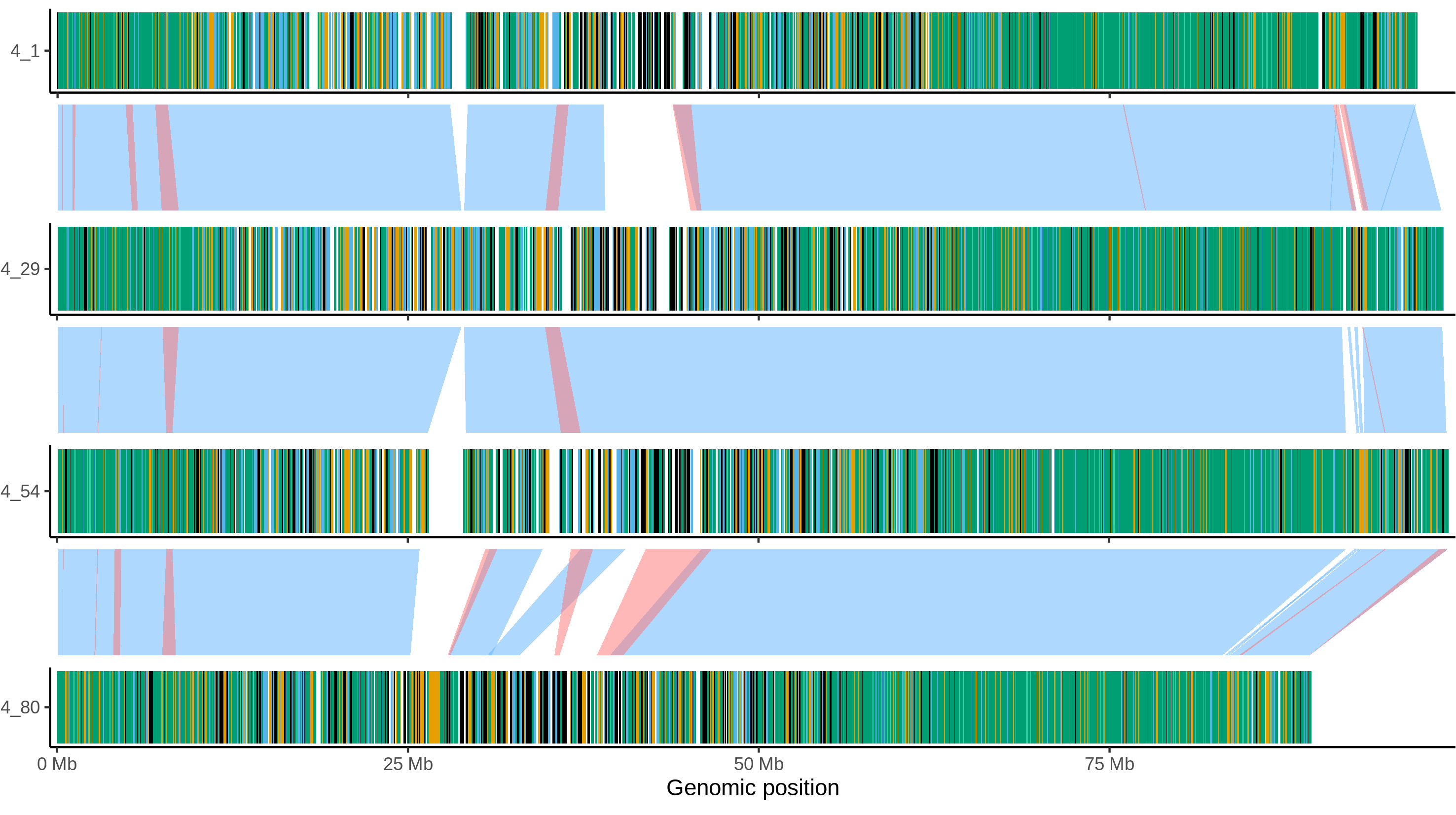

### Otava_chr_2.png

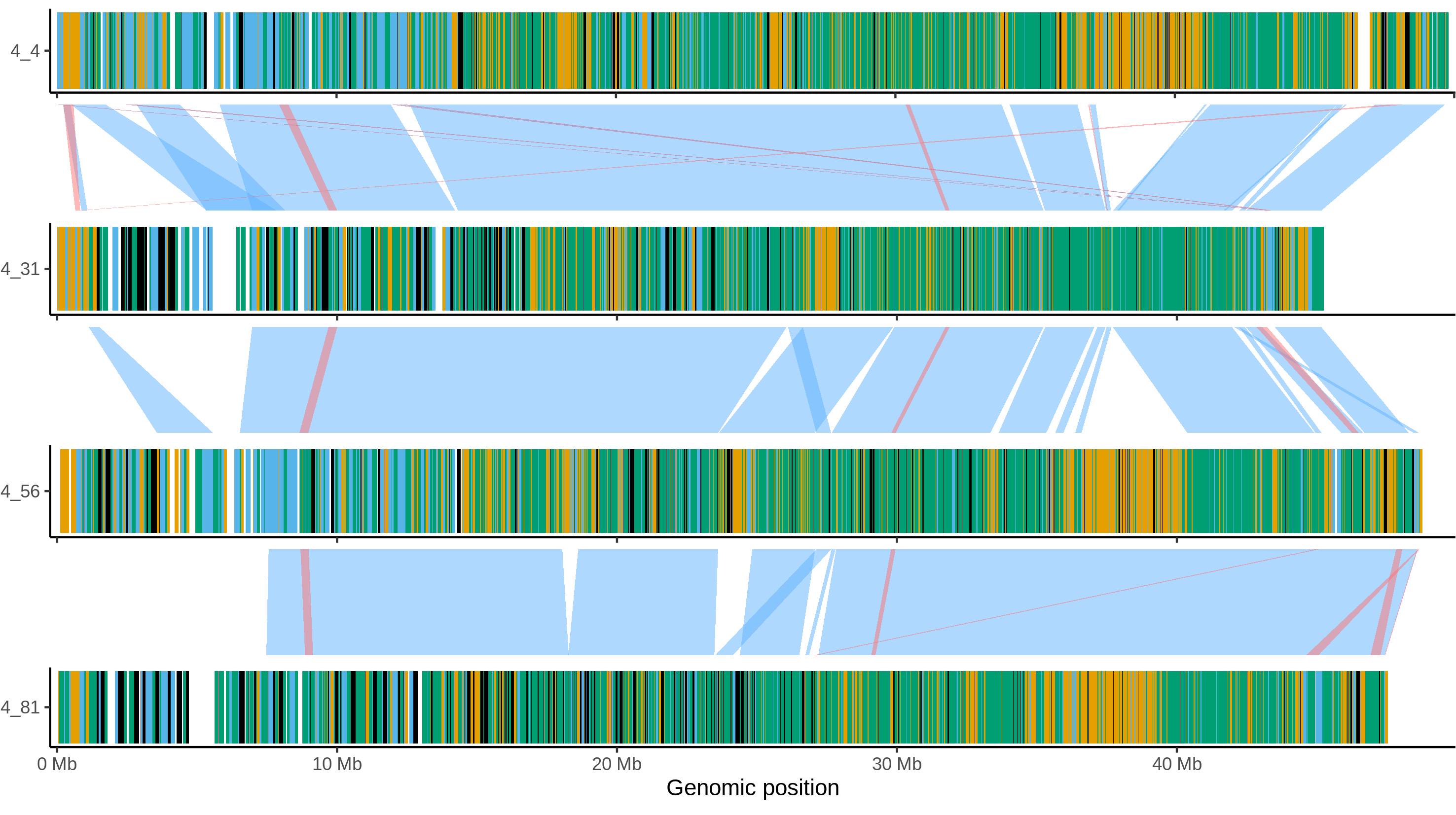

### Otava_chr_3.png

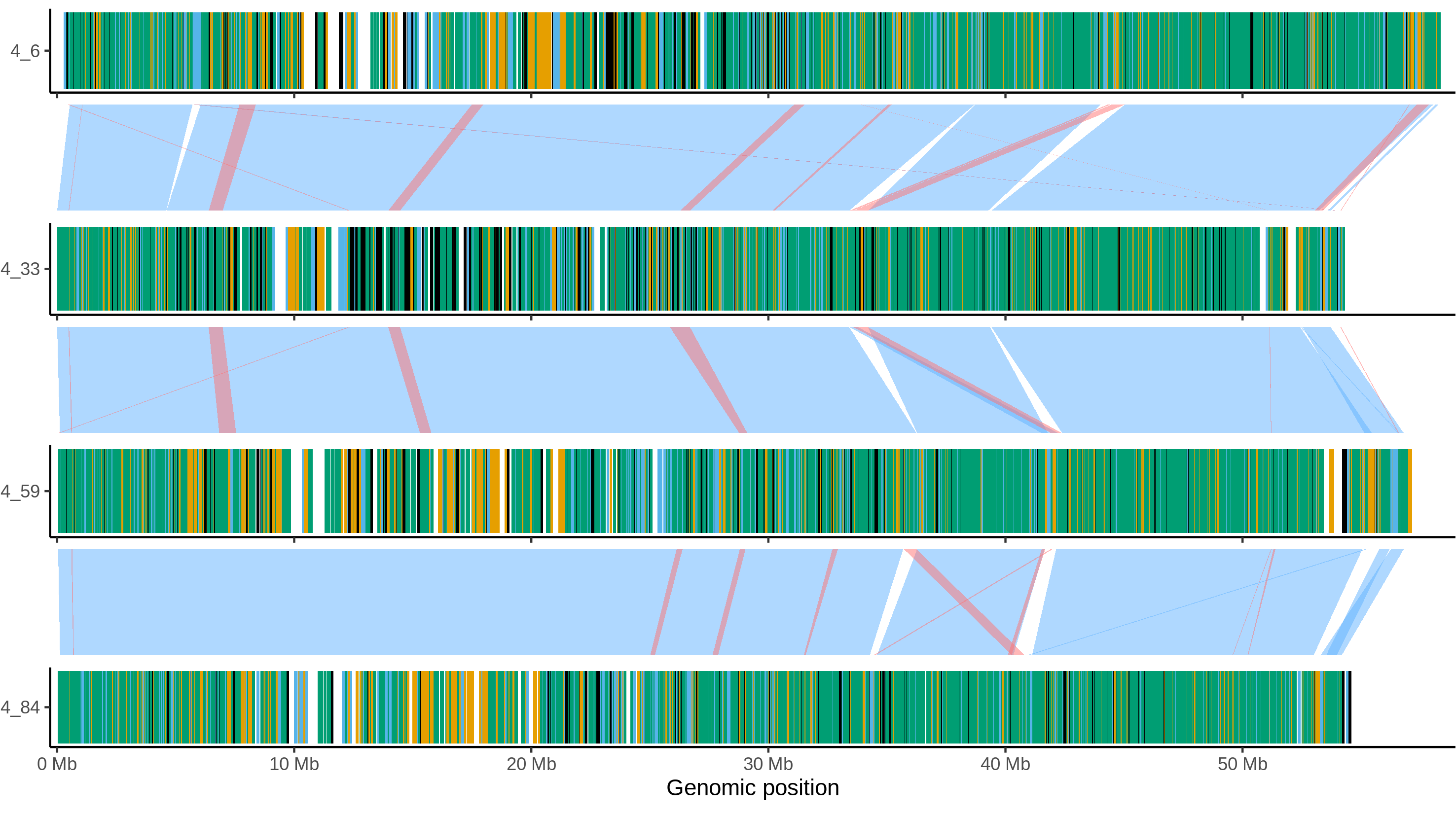

### Otava_chr_4.png

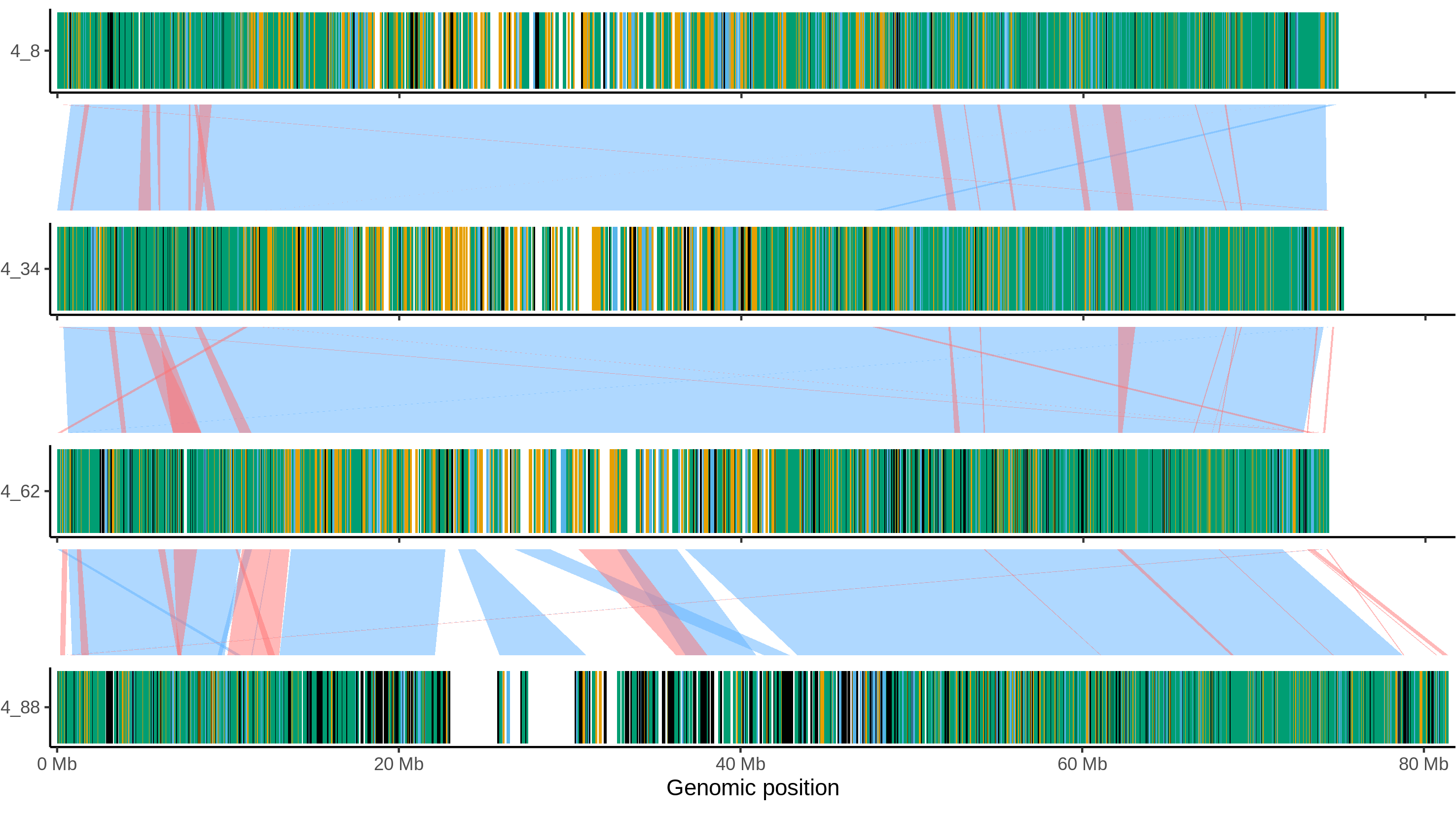

### Otava_chr_5.png

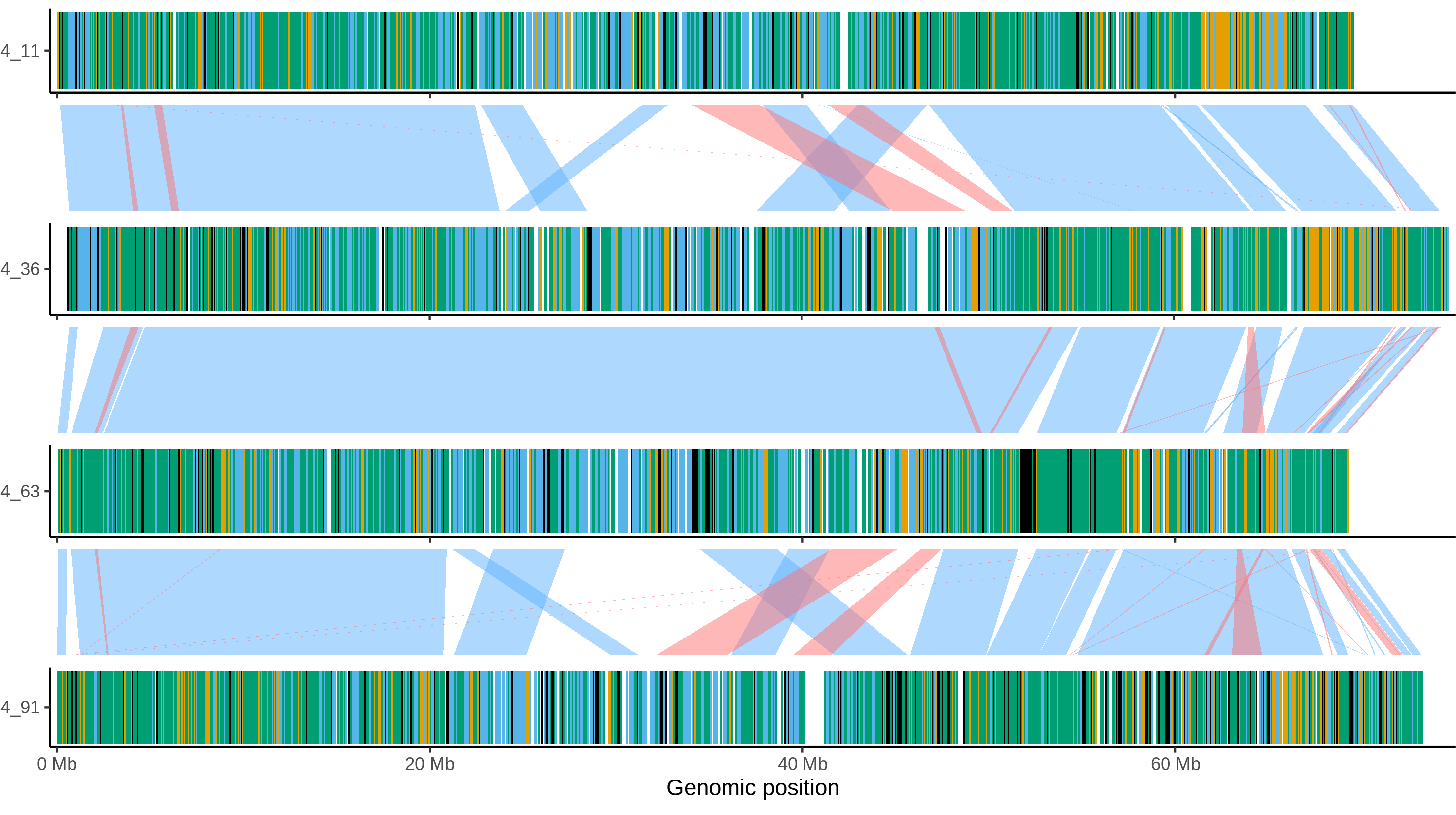

### Otava_chr_6.png

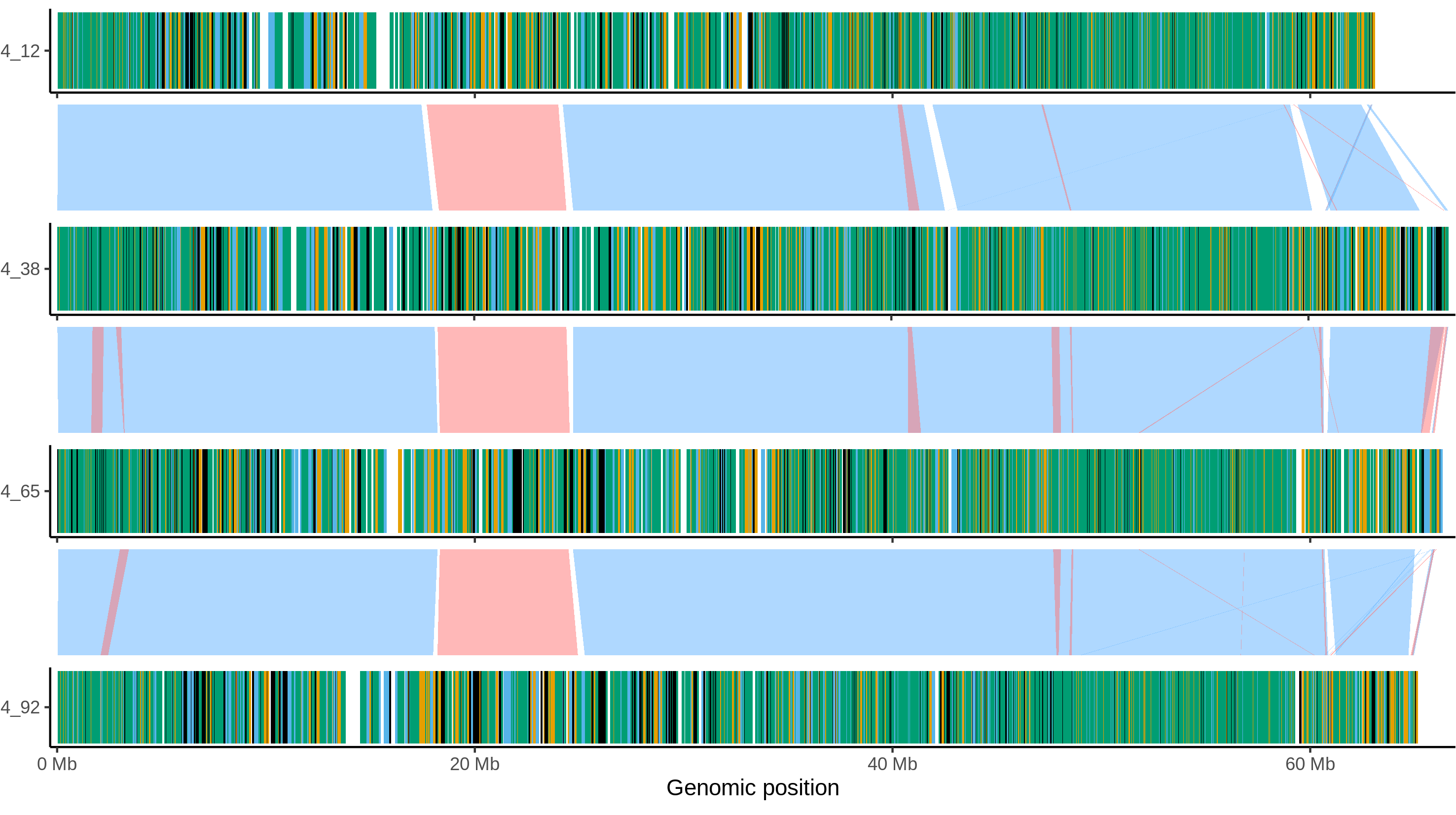

### Otava_chr_7.png

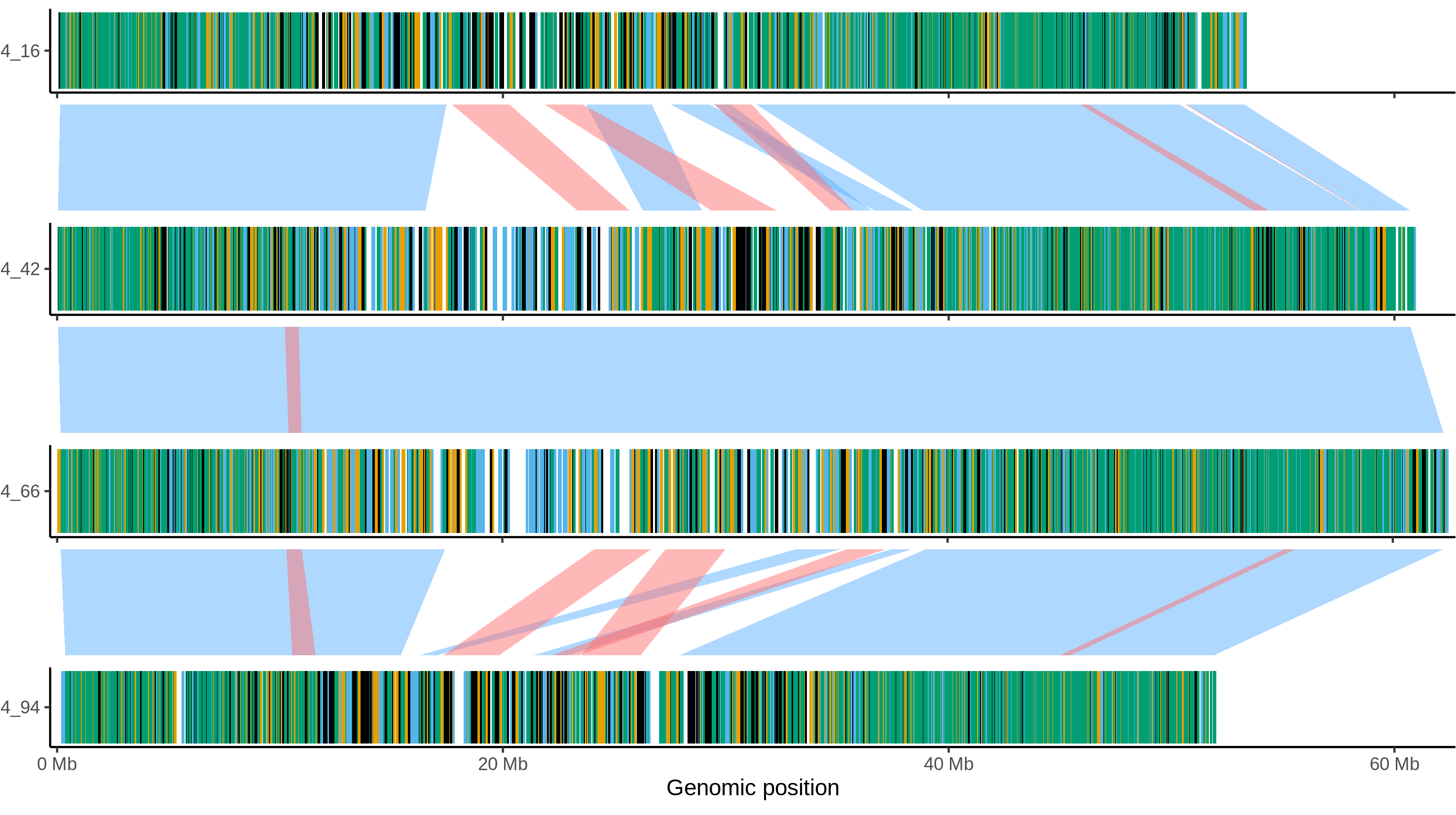

### Otava_chr_8.png

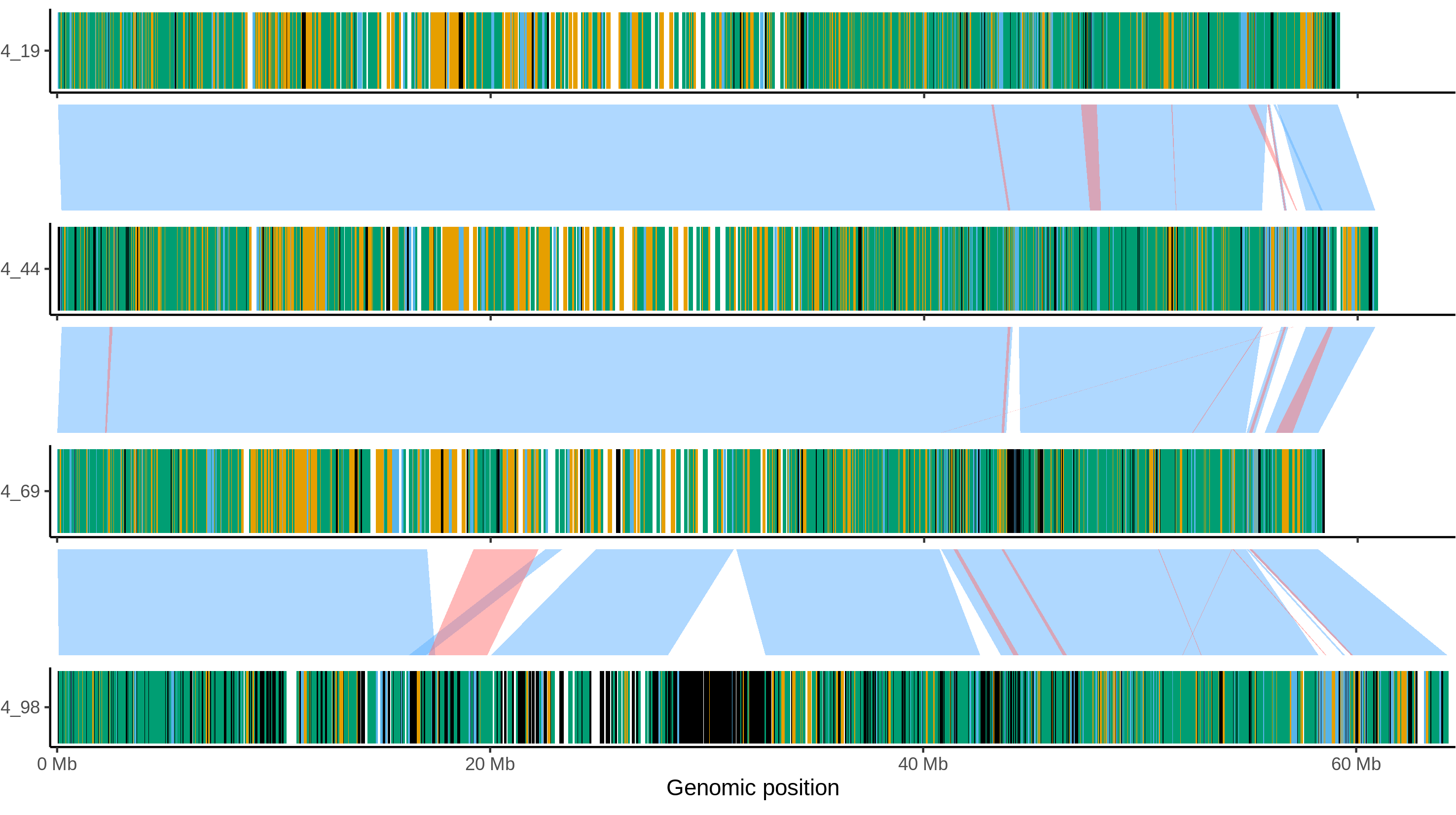

### Otava_chr_9.png

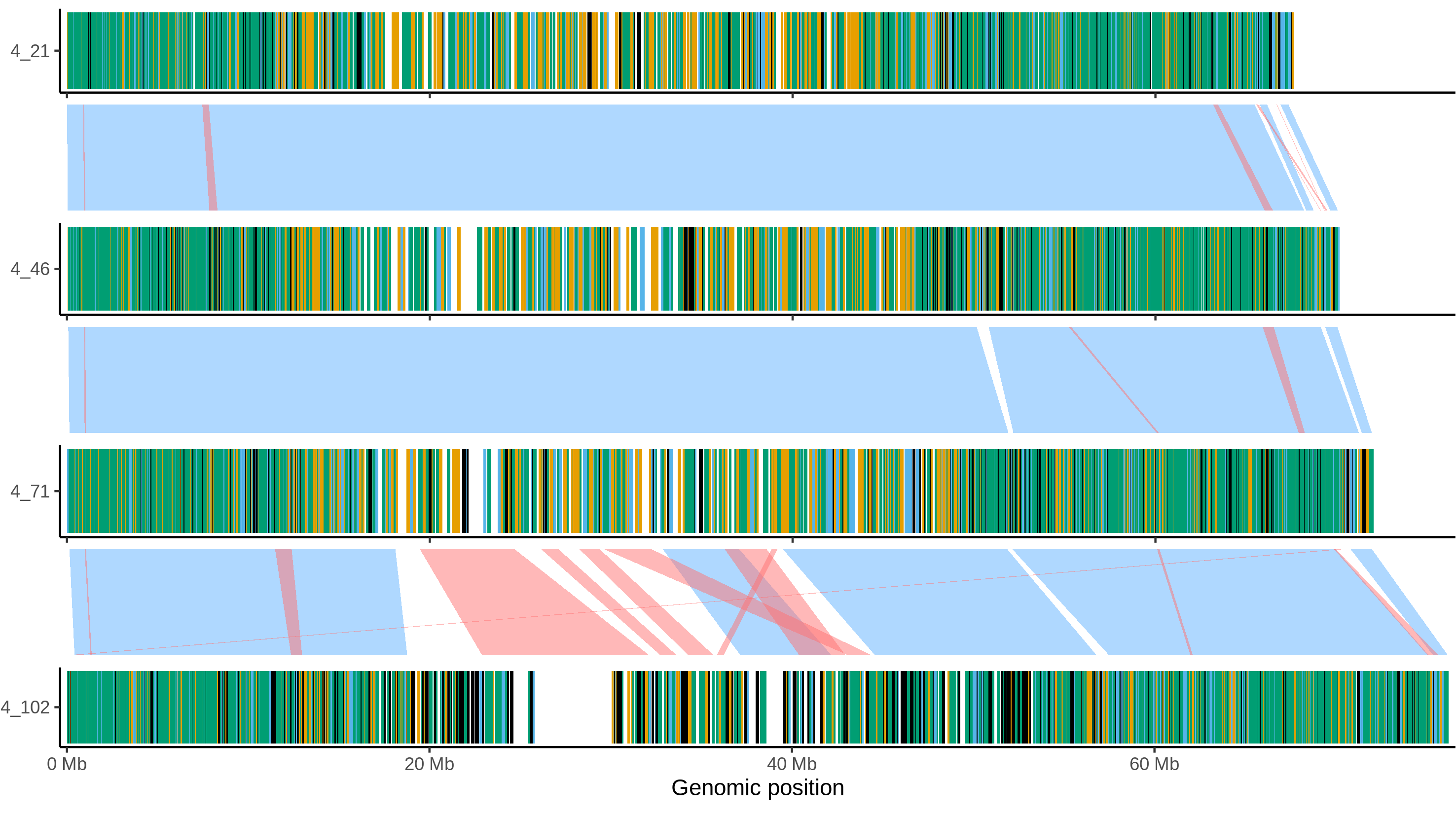

### Otava_chr_10.png

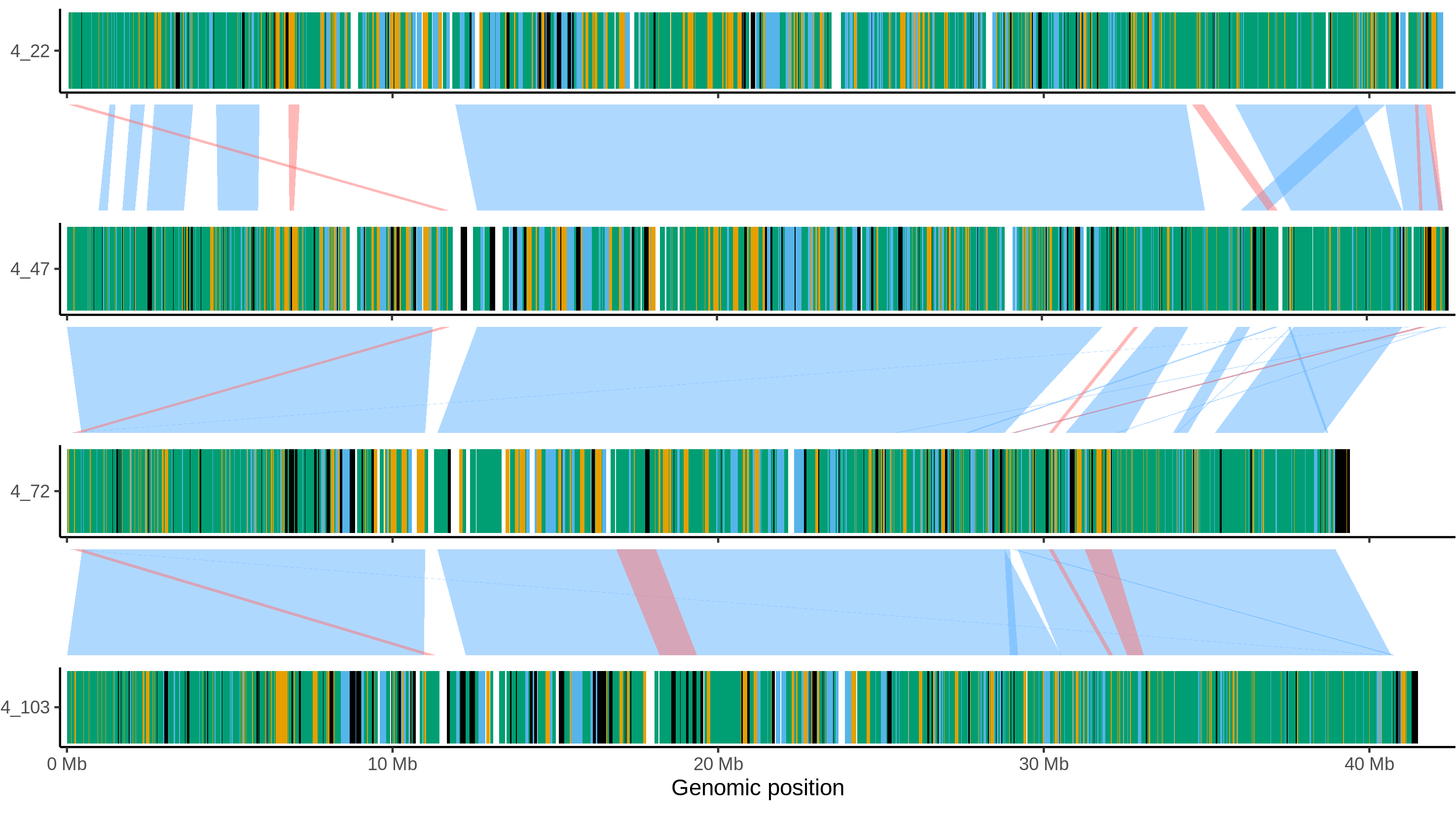

### Otava_chr_11.png

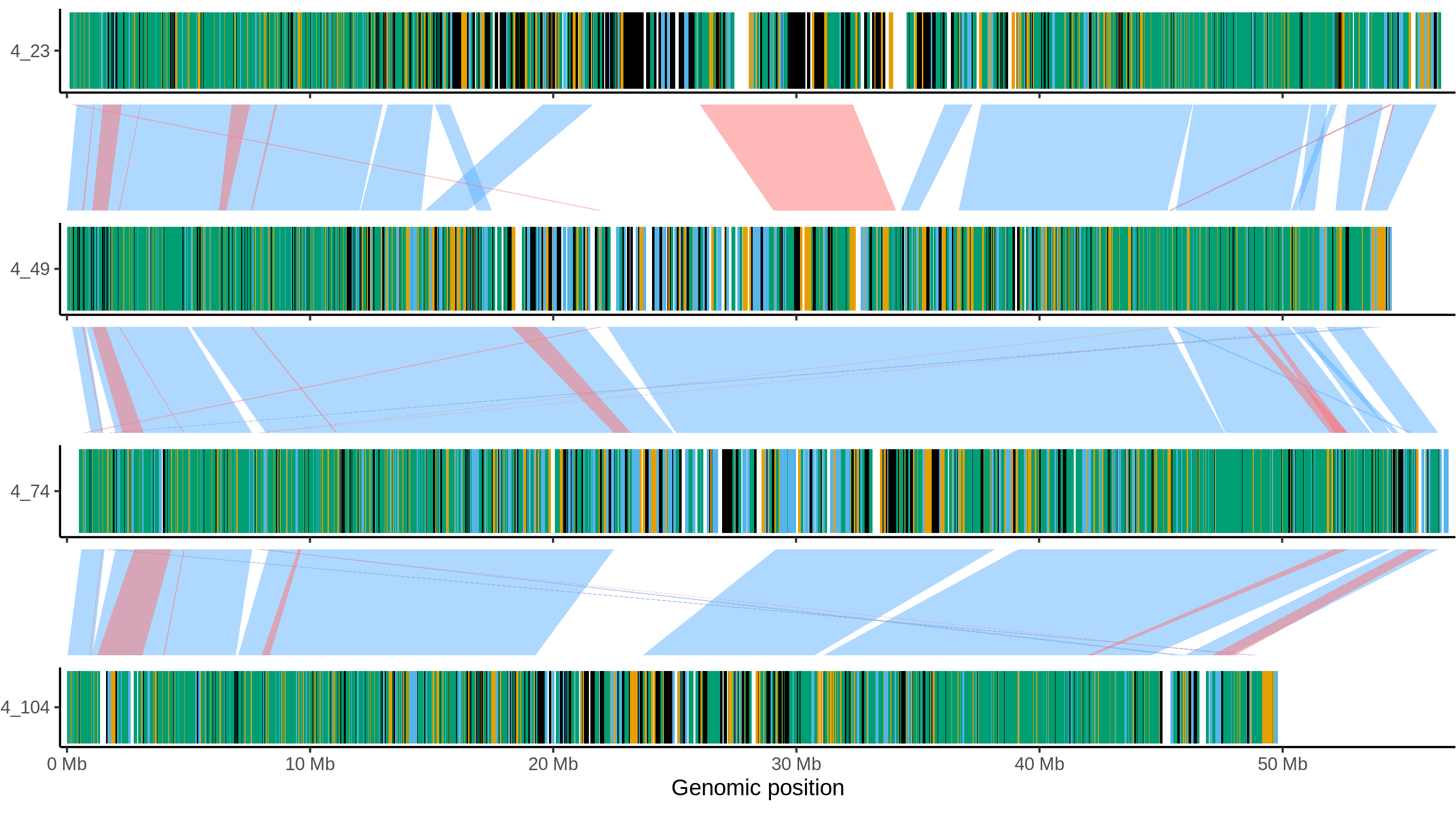

### Otava_chr_12.png

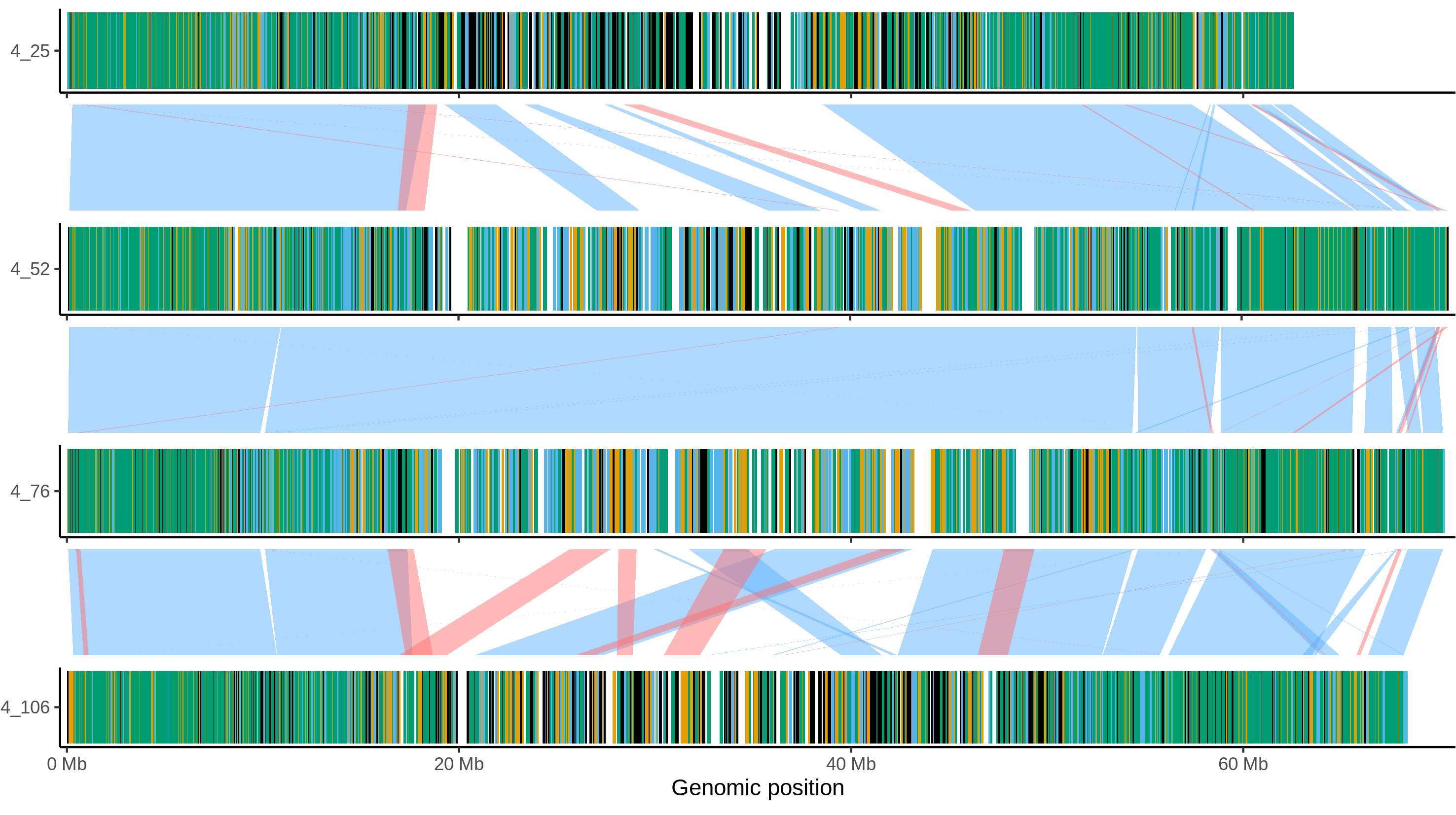
